# Distinct energetic blueprints diversify function of conserved protein folds

**DOI:** 10.1101/2025.04.02.646877

**Authors:** Malcolm L. Wells, Chenlin Lu, Daniel Sultanov, Kyle C. Weber, Erin Ahern, Zhen Gong, Ethan Chen, Anum Glasgow

## Abstract

While the ongoing revolution in structural biology offers an unprecedented understanding of the relationship between protein structure and function, it also confirms a puzzling, widely applicable principle: protein domains with highly conserved three-dimensional folds can perform radically disparate biochemical functions. To gain insight to this fundamental structural enigma, we mapped the energetic landscapes of a family of bacterial transcription factors and their anciently diverged structural homologs, the periplasmic binding proteins. Using hydrogen exchange/mass spectrometry, bioinformatics, X-ray crystallography, and molecular dynamics, we uncovered an unexpected contrast: despite binding the same sugars, the two families evolved unique “energetic blueprints” to support their distinct functional requirements. To test if differences in energetic ensembles have functional consequences, we rationally redesigned the protein fold for tunable ligand-driven transcriptional responses. Strikingly, energy-driven protein engineering produced synthetic transcription factors with the theoretically anticipated ligand-induced transcriptional outputs. Thus, decoding energetic blueprints among conserved protein folds provides a novel explanation for diverse functional adaptations, paves an alternative roadmap for protein design, and offers a new approach for engineering challenging drug targets.

---

Over 700 million natural proteins^1^ fold into only a few thousand spatial arrangements of secondary structures, or protein folds.^2–6^ Extant protein folds have been maintained over billions of years despite low sequence identity.^7^ To evolve a new function, a protein typically samples sequence changes that preserve its fold while varying its conformational dynamics.^8^ Sequence changes to a protein that disrupt essential allosteric regulation are disfavored.^9^ Thus, proteins are robust to mutations that do not disrupt essential dynamics or function, enabling sequence diversity while preserving the basic topology.

Unfortunately, methodological constraints in observing energetic relationships in proteins limit our capacity to identify the functionally important elements of the fold for most proteins. For example, protein functions may depend on perturbation-driven changes in the local stability of specific residues, secondary structures, intramolecular interactions, and quaternary interfaces. Consequently, we do not generally understand how new functions emerge from old folds.

Using a high-resolution integrative structural biology strategy, we determined the evolutionarily conserved, functionally important energetic relationships in two bacterial protein families that share the Venus Flytrap (VFT) fold. VFT-fold proteins are found across the domains of life.^7,10,11^ Five proteins from *E. coli* serve as our model system for probing energetic redistribution in the families: three dimeric LacI/GalR transcription factors (TFs) and two monomeric periplasmic binding proteins (PBPs). The TFs regulate sugar metabolism genes in the cytoplasm, while the PBPs are periplasmic transporters for the same sugars.^12,13^ The PBPs and TFs most likely evolved ligand binding specificity independently after a DNA binding domain (DBD) acquisition event that occurred before the divergence of eubacteria.^7,14^

Unlike the TFs, which are dimeric or tetrameric proteins where each protomer consists of a VFT domain attached to a DBD, the PBPs are monomers consisting of a single VFT domain. Upon binding ligands, PBPs undergo a pronounced conformational change in which the VFT fold subdomains close around the ligand via three flexible hinge loops.^15^ PBPs bind cognate ABC transporters to facilitate ligand transport from periplasm to cytoplasm, but also play a role in chemotaxis, where they bind a common chemotactic receptor, Trg.^16,17^

Despite their shared topologies, in experimentally solved structures, the sugar-induced conformational changes in the VFT domains of TFs are subtler than in PBPs due to the dimer state of the TFs.^12,13,18^ The family’s namesake TF, the *lac* repressor (LacI), reweights specific intramolecular contacts to switch between its transcriptionally repressed and active functional states without dramatic changes to the VFT domain structure.^19,20^ The DBD of each TF has a helix-turn-helix (HTH) motif that is connected by a hinge helix to the VFT domain. TFs switch between their repressed and active states by allosterically restructuring their DBDs in response to binding inducer molecules in their VFT domains. We hypothesized that the VFT-HTH architecture might be so strongly conserved in the TFs because transcriptional regulation requires domain coupling via a minimal set of evolutionarily conserved energetic relationships in the family. Because the PBPs have different functional requirements, they may have evolved different energetic relationships within the constraints of the same fold, despite interactions with the same ligands. We reasoned that discovering the energetic basis for TF-wide function would provide a map for engineering customized allosteric transcriptional regulators.

## Sequence conservation in VFT folds

To investigate whether evolution of separate functions in PBPs versus LacI/GalR TFs precipitated family-specific sequence changes in the VFT fold, we built a gene tree based on 2,222 diverse LacI/GalR TF and PBP sequences using only their VFT domains (Extended Data Fig. 1A, Fig. S1). Although the large number of sequences and the small number of inspected amino acid sites limit the resolution of the more internal branches, all TFs form a separate clade from all PBPs with high confidence (bootstrap support 100%), corroborating a previous study^7^ and suggesting independent evolution of the VFT domain in the two lineages.

Comparing conservation scores (see **Methods**) for each protein family revealed distinct patterns of sequence conservation (Extended Data Fig. 1B). While the PBPs are uniformly highly conserved, the TFs exhibit “valleys” of higher sequence diversity. As expected, there is high conservation in the DBD. However, the N-terminal lobe of the VFT domain is more permissive to mutations than the C-terminal lobe (Extended Data Fig. 1B). The conservation pattern reflects experimental phenotypic outcomes in LacI mutants (Extended Data Fig. 1C).^21^

Interestingly, we found substantial sequence variability in the TF dimer interface, reflecting the biochemical diversity of interfaces across many TFs (Extended Data Fig. 1E, Fig. S2).^18,22–24^ Although several other helices in the TFs are also sequence-diverse, the variability in the dimer interface is surprising because the LacI/GalR TFs are obligate homodimers, thus limiting the solvent accessibility in this region. Despite the low sequence conservation, a comparison of experimentally solved structures of TFs show that the dimer interface exhibits an extensive shared network of interactions (Fig. S3), corroborating previous findings^25^, with more abundant hydrophobic interactions ensuring the strength of the dimer interface and with more TF-specific polar interactions contributing to interface specificity. Of note, there are also more interactions in the N-terminal lobe than in the C-terminal lobe in X-ray crystal structures.

By contrast, the same region is highly conserved in PBPs despite their high solvent accessibility (Extended Data Fig. 1D), likely due to interactions with nutrient transporters.^26^ Virtually all PBP sequences from our gene tree are annotated as sugar-binding proteins, which are known to interact with Type I ABC nutrient transporters.^27^ Since the transporters are structurally conserved and their mode of interaction with PBPs is similar across different complexes, this pattern results in the higher sequence conservation in the PBP-transporter interface. Differences in sequence conservation in the two families reflect their distinct functional constraints while sharing a fold.

## TF structure across functional states

Bioinformatics-derived sequence-level differences do not reveal the structural basis for ligand regulation in the TFs, which requires comparing high-resolution X-ray crystal structures of related TFs in different functional states.^13,18,24,28–31^ Of the 41 experimentally solved structures of full-length LacI/GalR TFs, 31 are solved to <3 Å resolution (Table S1). Among all 100 published TF structures that include at least the VFT domain, 83 are solved to <3 Å resolution. Overall, all TF VFT domain structures have <4 Å pairwise root-mean-square deviation (RMSD) across their protomers (Fig. S4). We are interested in TFs that activate transcription by binding small molecules, and of this subset, only the LacI VFT domain structure is available in both operator-and inducer-bound states.^18,28–30^ In summary, an evolutionarily conserved activation mechanism in the LacI/GalR family cannot be gleaned from the available structural information.

To compare LacI with another sugar-inducible TF, we solved X-ray crystal structures of the ribose repressor (RbsR) in its inducer- and operator-bound forms at 2.1 Å resolution (Extended Data Fig. 2, Fig. S5, Table S2). RbsR is induced by ribose and has 33% sequence identity with LacI.^32^ While LacI controls lactose metabolism by binding the *lac* operator DNA (*lacO1*), RbsR controls ribose metabolism by binding the *rbs* operator DNA (*rbsO*).

Like the LacI complexes with inducer isopropyl β-D-1-thiogalactopyranoside (IPTG) and *lacO1*, RbsR-ribose and RbsR-*rbsO* have 1.58 Å backbone RMSD in the VFT domain (Extended Data Fig. 2A). The C-terminal VFT lobe is almost identical between the operator- and inducer-bound states of both TFs (0.40 Å and 0.61 Å RMSD for RbsR and LacI, respectively).

The two TFs are also highly structurally similar, with 1.6 Å RMSD in the VFT domain. The DNA-bound states for both TFs show the hinge helices buried in the minor groove of the operator, which is bent at 45° (Extended Data Fig. 2C, D). Notably, the extra base insertion between the palindromic repeats in *lacO1* compared to *rbsO* does not change the crystallographic DNA or protein conformation.

## Measuring local protein stabilities

For transcriptional regulation to occur, the TFs must sample different conformations with different probabilities in each state, resulting in *conformational ensembles* that are energetically distinct and not fully represented in crystal structures.^33^ To determine how the energy landscape is conserved in TFs (the “energetic blueprint”), we performed hydrogen exchange/mass spectrometry (HX/MS) on three paralogous sugar-inducible LacI/GalR TFs in their apo, DNA-bound, and inducer-bound states: the functional dimeric form of LacI (residues 1-331), RbsR, and the galactose repressor (GalR) (Figs. 1A, S6-S8, Tables S3-S5, Appendices 1, 2).^34^ GalR induces transcription of the *gal* operon by binding galactose and has 25% sequence identity to LacI and 31% to RbsR.^35^ While the LacI/GalR TFs sample both monomeric and dimeric states, we confirmed that our LacI and RbsR samples predominantly exist as dimers at protein concentrations comparable to those used in the HX/MS experiments (3 µM) and down to 10 nM (Fig. S9). Moreover, the presence of their respective inducers (IPTG for LacI and ribose for RbsR) did not observably alter this equilibrium.

**Figure 1.**
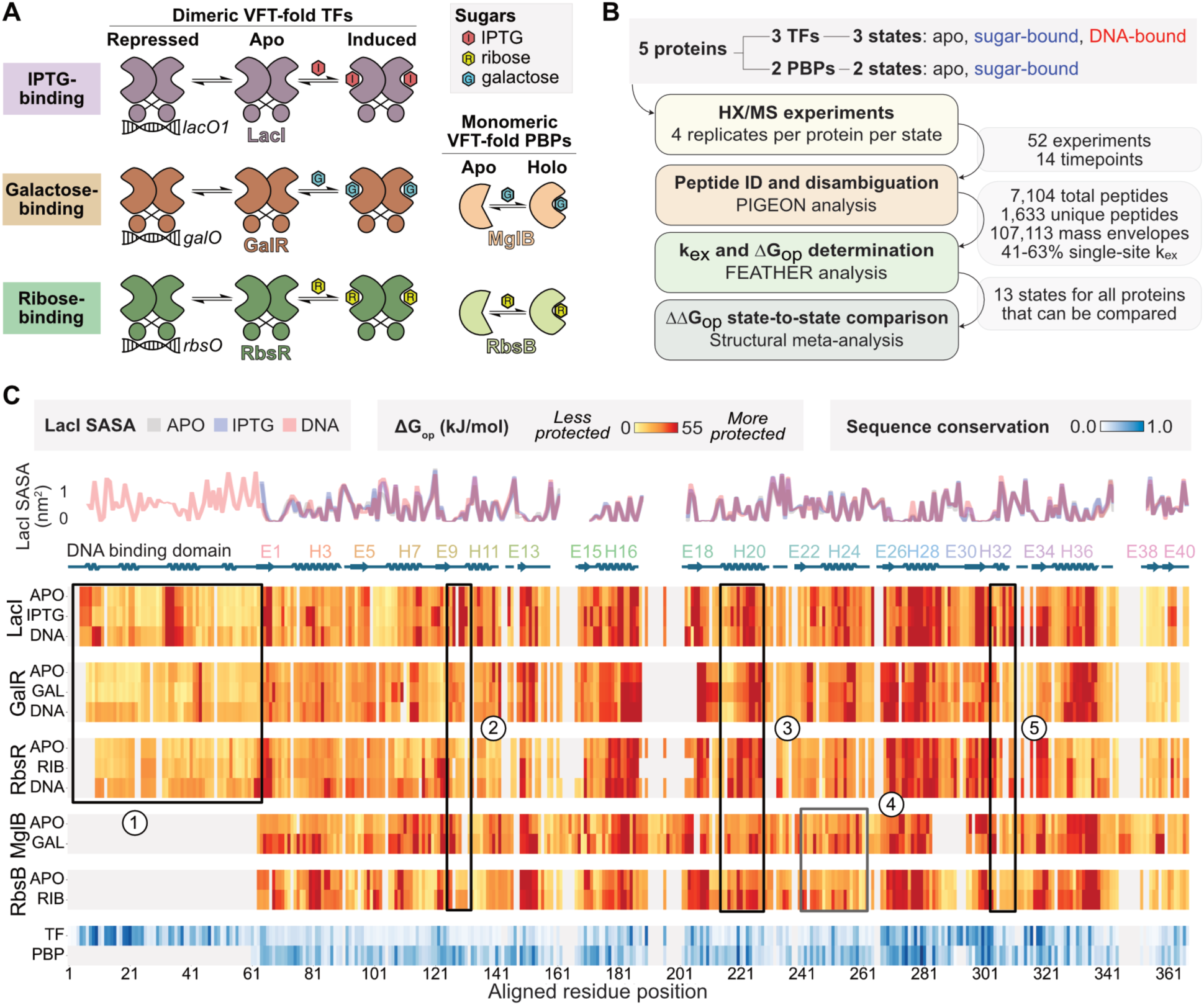
High-resolution HX/MS. **(A)** We chose two PBPs and three TFs to study by HX/MS, including two PBP/TF pairs that respond to the same ligands. **(B)** Experimental design and data summary. **(C)** ΔG_op_ heatmap for the proteins ordered by structurally aligned residues, compared to LacI solvent-accessible surface area (SASA) and TF and PBP sequence conservation in blue. Gray marks an alignment gap. *Right*: VFT-fold structure schematic. Several regions are marked by matching numbers on the heatmap and schematic. (1) The DBD has higher ΔG_op_ in the operator-bound state than in apo or sugar-bound states in the TFs. (2) A binding pocket loop responds differently to sugar binding in each protein. (3) Sugar binding stabilizes helix 20 in each protein. (4) Sugar binding stabilizes a C-terminal VFT lobe segment in the PBPs. (5) Sugar binding stabilizes the N-terminal region of helix 36 for all proteins. Secondary structure annotation: H, helix; E, strand.

To account for differences in hydrogen exchange among the TFs that might arise from binding different sugars, we also performed HX/MS on two PBPs that bind the same sugars as RbsR and GalR: ribose binding protein (RbsB) and galactose binding protein (MglB) (Figs. 1A, S10, Tables S6, S7). We observed EX2 regime behavior in all peptides collected for this study, indicating folded proteins. These five VFT proteins bind the sugars with affinities in the 100 nM to 1 mM range. Both TF/PBP pairs have 28% sequence identity in the VFT domain.

HX/MS measures the rate of exchange of backbone amide hydrogens with deuterium, revealing the relative local structuredness of regions of the protein in various functional states. Using PIGEON-FEATHER^20^, a high-resolution HX/MS analysis method, we determined the single-residue free energies of opening (ΔG_op_) for all three TFs in three states (apo, sugar-bound, and operator-bound) and both PBPs in two states (apo and sugar-bound) (Fig. 1B, Extended Data Fig. 3, Tables S8-S12). The ΔG_op_ is a measure of local stability; the highest ΔG_op_ values in a protein correspond with its global free energy of unfolding (for a more detailed summary, see Supplemental note 1).^36^ Ligand-driven shifts in ΔG_op_ report local energetic changes in the context of the protein ensemble. Because it is not possible to measure the sugar- and DNA-bound ternary complex of a TF directly^19^, allosteric energetic changes in the TF conformational ensemble are inferred from comparing the DNA- and inducer-bound states.

The ΔG_op_ datasets show local and global protein responses to binding sugars and operators (Figs. 1C, S11-S13, Supplemental note 1), demonstrating that some effects occur only in PBPs or TFs (Fig. 1C-1, 1C-4), while others are conserved in all VFT-fold proteins (Fig. 1C-3, 1C-5). Some regions of the VFT fold are differentially stabilized by ligand binding in each protein. For example, a structurally conserved binding pocket loop (Fig. 1C-2) responds differently to each sugar in each protein, suggesting protein-specific evolution of a functional role for the loop to allow *E. coli* to tune ligand binding affinity and specificity. We discuss state-, family-, and fold-specific energetic changes in the next sections.

**Figure 2.**
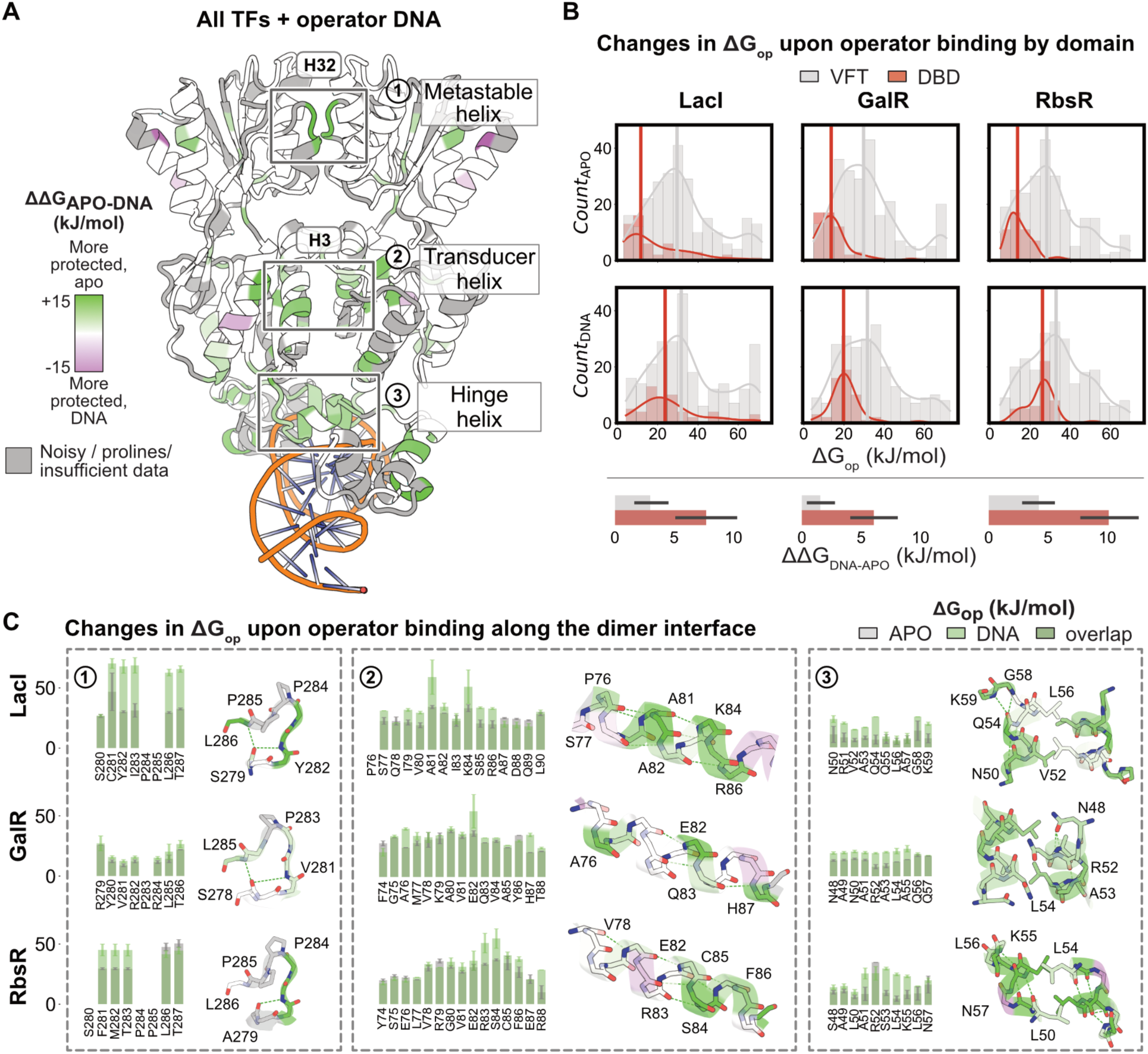
Conserved allosteric effects of DNA binding in TFs. **(A)** Conserved energetic effects of operator binding in the three TFs. **(B)** Although there is global restructuring in the TFs, operator binding causes a larger increase in average ΔG_op_ in the DBD than the VFT domain. Error bars show the standard deviation. **(C)** Conserved increase in ΔG_op_ for each TF in key operator-responsive regions in the dimer interface. Error bars reflect FEATHER bootstrapped uncertainties.

**Figure 3.**
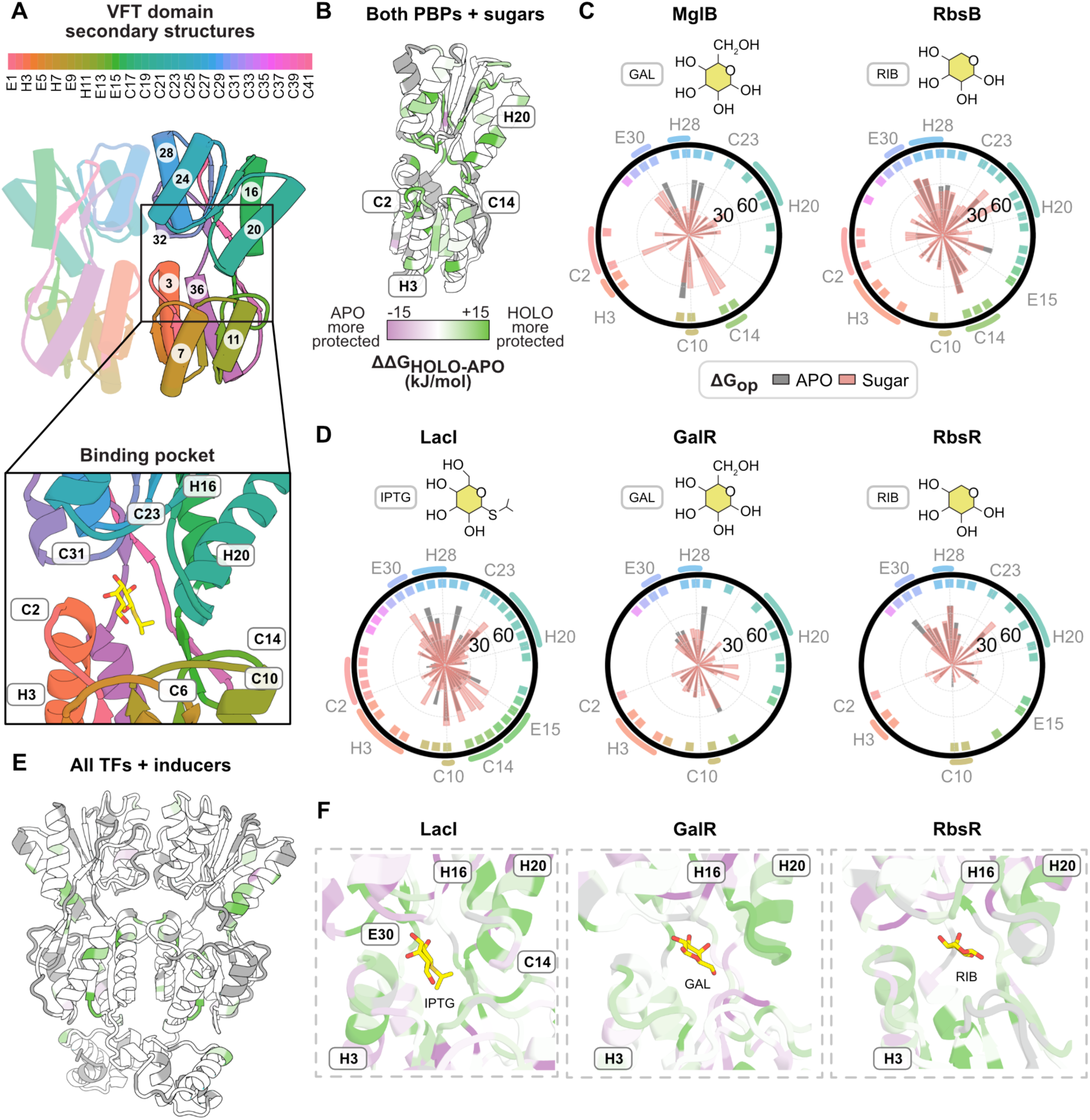
Ligand binding in PBPs and TFs. **(A)** Color-coded secondary structures of the VFT domain. Secondary structure annotation: H, helix; E, strand; C: coil. Gray: prolines, or noisy or insufficient data. **(B)** Conserved effects of sugar binding in PBPs. Green regions are stabilized by sugar binding in both PBPs. **(C)** Energetic redistribution in PBPs upon sugar binding. The radar plots show the ligand binding site. Each box at the radar plot edge, colored as in (A), corresponds to a residue lining the binding pocket. The spokes, colored by functional state, show their ΔG_op_: *e.g.*, residues in loop C14 in MglB are stabilized by sugar binding (increased ΔG_op_). This loop is green in (A). **(D)** Energetic redistribution in the TFs upon sugar binding, colored as in (C). **(E)** Conserved energetic effects of inducer binding in the three TFs. **(F)** Molecular details of the TF binding pockets. (E) and (F) are colored by ΔΔ_Gop, HOLO-APO_, as in (B).

## Allosteric effects of TF-DNA binding

To learn how the TFs respond to binding operator DNA, we compared their ΔG_op_ in operator-bound and apo states (ΔΔG_op, DNA-APO_) (Fig. 2A, Extended Data Fig. 4A). As residue-level ΔG_op_ values depend on the local structural context, absolute ΔG_op_ values are most comparable across proteins when their underlying closed-state baselines are equivalent. Therefore, we focus on intra-protein ΔΔG_op_ patterns to identify conserved energetic effects. As expected, the DBD of each TF has higher average ΔΔG_op, DNA-APO_ than the VFT domain due to direct interactions with DNA^37^ (Fig. 2B), but operator binding also structures the VFT domain, the hinge helix^38^, and several VFT-DBD interactions (Fig. 2B, C).

Remarkably, binding to operator DNA also stabilizes the TFs far beyond the DBD, an effect of the linked equilibrium between the DNA-bound and dimer states of LacI.^39^ One example is the structuring of a C-terminal dimer interface helix (the “metastable helix”, H32), which points to increased dimer stability in LacI/GalR TFs upon operator binding regardless of the DNA sequence, length, or symmetry (Fig. 2C-1, Supplemental note 2). This helix backbone is identical in crystal structures of LacI and RbsR in all functional states (Extended Data Fig. 2A, Fig. S5-3) and in models of GalR.^40^ The behavior of H32 exemplifies a functionally important, evolutionarily conserved shift in a protein’s ensemble that occurs without a crystallographic conformational change. While the regions flanking H32 are conserved in the LacI/GalR MSA (Extended Data Fig. 1E, Fig. S2), H32 itself is not, suggesting that evolution might tune this region to control DNA binding affinity and inducer sensitivity. In LacI, mutations to H32 affect the quaternary structure and ablate DNA binding.^41,42^ By contrast, the helix is well-conserved in PBPs (Extended Data Fig. 1D), likely because it interacts with membrane transporters.

We also observed other conserved effects of operator binding at the dimer interface, including stabilization in the “transducer helix” H3 (Figs. 2C-2, S5-2), which bridges the binding pocket with the DBD and experiences ligand-dependent shifts in ΔG_op_ that are discussed further in the next section. Operator binding also stabilizes the central residue of the N-terminal VFT beta strand E5 in each TF, together with its hydrogen bond partner in the neighboring strand and in the DBD (Extended Data Figure 4A, Fig. S5-4, Supplemental note 2). We used DNA constructs with two operator repeats for HX/MS in order to maximize bound species. To confirm this choice does not affect DNA binding behavior in these experiments, we also performed an experiment with LacI and a DNA construct with only one LacI binding site (Fig. S14).

X-ray crystal structures of LacI-*lacO1* and RbsR-*rbsO* showed that TF binding bends operator DNA (Extended Data Fig. 2D, Fig. S5-1). NMR spectroscopy on the LacI DBD corroborated this result and further demonstrated that DBD binding to off-target DNA maintains a linear DNA conformation.^43^ However, the structural explanation for LacI specificity for *lacO* remains mysterious because the behavior of the VFT domain in the presence of off-target DNA could not be studied using either technique. To explore LacI response to on- vs. off-target DNA in its VFT domain, we compared peptide-level HX/MS of LacI in solution with *lacO1* (on-target, pseudosymmetric), *rbsO* (off-target, symmetric), and *scrO*, a scrambled version of the *lacO1* sequence (off-target, not symmetric) (Fig. S15, Appendix 1). While *rbsO* produced attenuated effects in the DBD compared to *lacO1*, of the three DNA sequences tested, only *lacO1* could cause the energetic changes in the metastable helix H32, >50 Å from the DBD. (It is interesting that *rbsO* has a stronger effect on the LacI DBD than *scrO*, given that increased A-T content in off-target DNA improves LacI binding^44^, and *rbsO* has 15% *less* A-T content than *scrO.*) The structural rearrangements driven by *lacO1* in the VFT domain, which are not observed in LacI incubated with nonspecific (*rbsO*, *scrO*) DNA binding motifs, promote stronger binding to operator DNA as a demonstration of DNA sequence-induced protein allostery.

## VFT families differ in sugar response

We reasoned that comparing ΔG_op_ for structurally aligned regions would highlight similarities and differences in how binding sugars redistributes the energy in TFs vs. PBPs to mediate distinct sense-response behaviors (Fig. 3).^20^ Similarly, comparing ΔG_op_ for TFs and PBPs that bind the same ligands can reveal how the same fold can be adapted for different functional responses to the same ligand (Figs. 3, S16, Extended Data Fig. 4B). While sugar binding causes unique local effects in each protein due to each sugar’s specific biochemical characteristics (Fig. 3C, D), we also observed long-range pan-VFT and family-specific patterns (Figs. 3B, E, S16).

For example, family-specific energetic changes occur in the hinge loops of PBPs, which are stabilized in the sugar-bound state^13^ (Figs. 3B, S16, S17). By contrast, the TFs show no changes in these loops in apo vs. sugar-bound structures (Fig. 3E). Overall, although the PBPs and TFs bind the same sugars, they share few conserved effects by HX/MS (Figs. 3, S16C). All hinge loops are stabilized in both PBPs; some positions have ΔΔG_op, HOLO-APO_∼20 kJ/mol, on par with the strongest effects in the TFs that arise from protein-sugar interactions (Fig. 3B, C).

Each TF is globally stabilized by inducer binding (Fig. 3D-F), particularly in the binding pocket, pocket-periphery, and dimer interface. The N-terminal VFT dimer interface experiences pervasive inducer-triggered allosteric structuring: part of the transducer helix is stabilized, and strands E5, E1, and E9 unite to form a super-sheet with a hydrogen bond pattern distinct from the apo and operator-bound states (Figs. S5-4, S18). The differences in how inducer and operator binding stabilize this region suggest that TF induction includes a strand slipping mechanism that favors an energetically similar yet configurationally distinct beta sheet with a different strand register (Figs. S5-4, S18, Extended Data Fig. 4B, Supplemental note 2). Crystallographic evidence supports the HX/MS data: in LacI, strand slippage occurs through squeezing out a water molecule that is present in the operator-bound state; while in RbsR, hydrogen bond rearrangement occurs without a change in hydration (Fig. S18). Strand slippage is unlikely to occur spontaneously because it requires a substantial energetic disruption to other local structural elements but was previously implicated in mediating conformational changes in a membrane protein.^45^ Helices 16 and 20 at the binding pocket periphery are also critical to inducer binding, corroborating mutational studies of LacI (Fig. 3F).^19,21^

To understand the baseline requirements for sugar binding in the VFT fold, we collated conserved effects in all five proteins (Fig. S16C). All proteins are stabilized by sugar binding in the binding pocket helices (ΔΔG_op, HOLO-APO_>10 kJ/mol). However, while there are many family-specific energetic changes, we observed few pan-VFT effects, which underscores how the fold is uniquely co-opted by each family.

## Tuning TF inducer sensitivity by design

The HX/MS analysis revealed that the VFT-fold PBP and TF families have evolved unique energetic blueprints that enable their distinct functions. We hypothesized that these blueprints may be co-opted to rationally engineer VFT-fold sensors with tunable ligand responses, a central goal in protein engineering. Customized TFs in particular enable precise control over gene expression for optimized metabolic flux in biosynthetic pathways, reduced cellular toxicity, and dynamic regulation in synthetic gene circuits.^46^

Towards this end, we noted that although the metastable helix H32 has limited sequence conservation (Extended Data Fig. 1E, Fig. S2), it is stabilized in all operator-bound TFs compared to their inducer-bound states (Fig. 2). We hypothesized that computationally tuning the stability of this helix would result in controllable inducer sensitivity despite its 15 Å and 45 Å distance from the inducer binding pocket and DBD, respectively. To test this hypothesis, we computationally redesigned residues 276-282 in the LacI metastable helix^47^, generating 675 design models that we scored using Rosetta energy metrics^48^ (Fig. 4A). While maintaining the same design protocol and goal, we simultaneously ran three trials: in the first trial, the mutable residues included only the metastable helix; in the second trial, we additionally enabled five residues that interact with the metastable helix (S221, M223, L251, M254, R255) to mutate; and in the third trial, we allowed only metastable helix residues to mutate, but we applied a bias to favor the selection of polar residues. We then chose six designs using a Pareto front strategy to test experimentally in a cell-based fluorescence assay, in which a LacI variant is expressed constitutively from one plasmid, and controls the expression of a green fluorescent protein (GFP) from another plasmid (Figs. 4B, C, S19, Table S13).^49^ The GFP expression plasmid included one copy of the WT pseudosymmetric *lacO1* sequence upstream of the GFP sequence. The resulting cell fluorescence measurements were fit to the Hill equation as defined in Tack *et al.*^50^ (Table S14), with the limitation that this equation does not include a term quantifying the oligomerization state of the protein. Western blot analysis showed that all designed variants expressed within 6.5-fold of the WT LacI expression level (Fig. S20).

**Figure 4.**
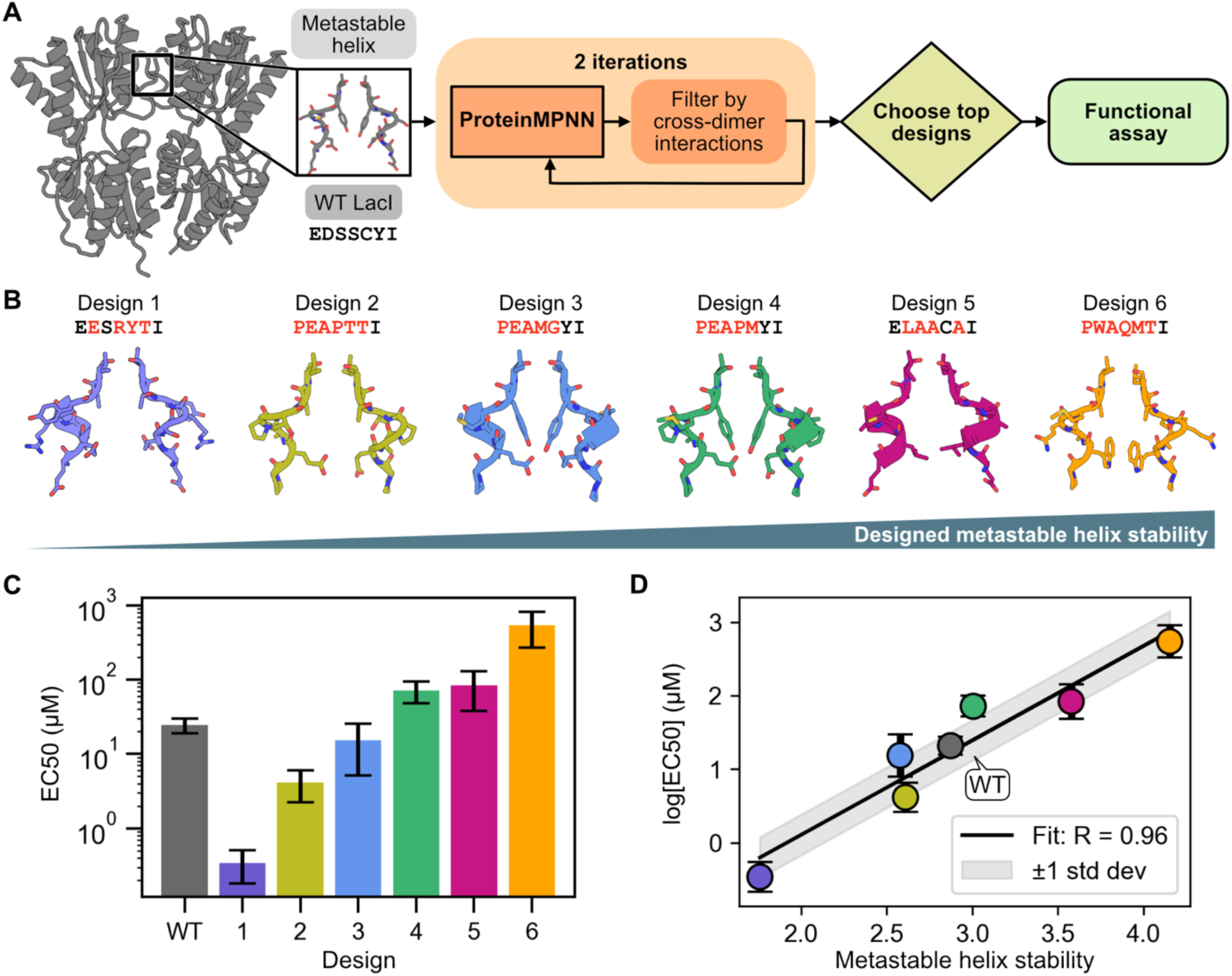
Computationally designed inducer sensitivity in LacI variants. **(A)** Design strategy for tuning metastable helix stability. **(B)** Top six design models. Mutations from WT sequence in red. **(C)** The designed LacI variants have IPTG EC_50_ values spanning four orders of magnitude. The *y-*axis is log-scaled. **(D)** Linear regression between the log(EC_50_) values and the metastable helix stability scores, as calculated by the composite energy function, indicates a strong correlation between our computational metrics and experimental results. Error bars indicate standard deviation.

Compared to the WT LacI IPTG EC_50_ of 24.5 µM, the variants had EC_50_ values ranging from 0.3 µM to 543 µM (Fig. 4C, Extended Data Fig. 5, Table S14). We confirmed by mass photometry that Design 1 remains dimeric at 10 nM and is not shifted by IPTG (Fig. S21), indicating that the altered inducer sensitivity does not arise from changes in dimerization. Remarkably, the variants’ EC_50_ values follow a simple composite energy function combining the selection metrics (Fig. 4D, Table S13) (see **Methods**). The ease with which one can tune the inducer sensitivity by modulating the metastable helix illustrates the utility of the energetic blueprint in protein engineering, especially considering the challenges in rationally designing predictable ligand responses in LacI and other allosteric proteins.^49,51,52^ We anticipate that the composite function can be used to redesign LacI/GalR TFs for which experimental data are not available to generate customized transcriptional regulators.

Interestingly, the designed variants included several mutations that ablate LacI function when introduced individually.^42,53^ This shows that the energetic effects of deleterious mutations can be compensated when a protein is designed with respect to the family’s energetic blueprint.

## The role of ordered waters in TF induction

HX/MS shows that inducer binding stabilizes the transducer helix H3 in the binding pocket of each TF (Fig. 3D), but crystal structures suggest that this requires water-mediated hydrogen bonds (Fig. S5-2).^18,19^ We were curious whether structural water molecules, defined as waters that form three or more hydrogen bonds with protein or inducer atoms, are evolutionarily conserved to maintain the induced TF ensemble.^19^ To explore this, we used RosettaECO (Explicit Consideration of cOordinated water)^54^ to model structural waters in the induced states of several sugar-inducible TFs. We identified a high-confidence water molecule in the binding pocket of the TFs near the dimer interface, bridging the sugar with the transducer helix (Fig. S22). In agreement with the models, this water is found with very low b-factors in RbsR-ribose (b=11.73) and LacI-IPTG (b=23.80) crystal structures (Fig. S5-2).^18^ As an interesting exception, in the trehalose repressor (TreR), the phosphoryl group of the inducer trehalose-6-phosphate acts as the bridge^22^ (Fig. S22C), whereas unphosphorylated trehalose is an anti-inducer that stabilizes binding on the operator DNA.^55^ The LacI anti-inducer *ortho*-nitrophenyl-β-D-fucoside (ONPF) similarly disrupts the water network established by IPTG.^19^

Using the crystal structures of LacI-IPTG and RbsR-ribose and our model for GalR-galactose, we performed atomistic MD simulations as an orthogonal method to explore how structural waters contribute to the induced TF ensembles (Fig. 5). We extracted the waters within 3.5 Å of the sugars and performed a clustering analysis to find clusters with >80% occupancy (Figs. 5A, B, S23). Echoing the RosettaECO results, the simulations identified the same water near the transducer helix to have high occupancy in all TFs (Fig. 5C). The highest duration/occupancy for this structural water was in LacI, due to two hydroxyl groups in IPTG positioned towards the dimer interface, rather than only one in ribose; in GalR, it has near 100% occupancy. The water does not appear in operator-bound TFs (Fig. S5-2)^29^, and we expect only transient waters in the binding pockets of apo TFs. These data support the hypothesis that the sugar-inducible TFs evolved to maintain not just the VFT fold or functionally important conformational changes, but also specific interactions with water that contribute to the induced state ensembles. PBPs also make a similar water-mediated interaction with sugars (Fig. S24), suggesting that the pan-VFT-fold stabilization of the N-terminus of the transducer helix upon sugar binding relies partially on a structured water molecule.

**Figure 5.**
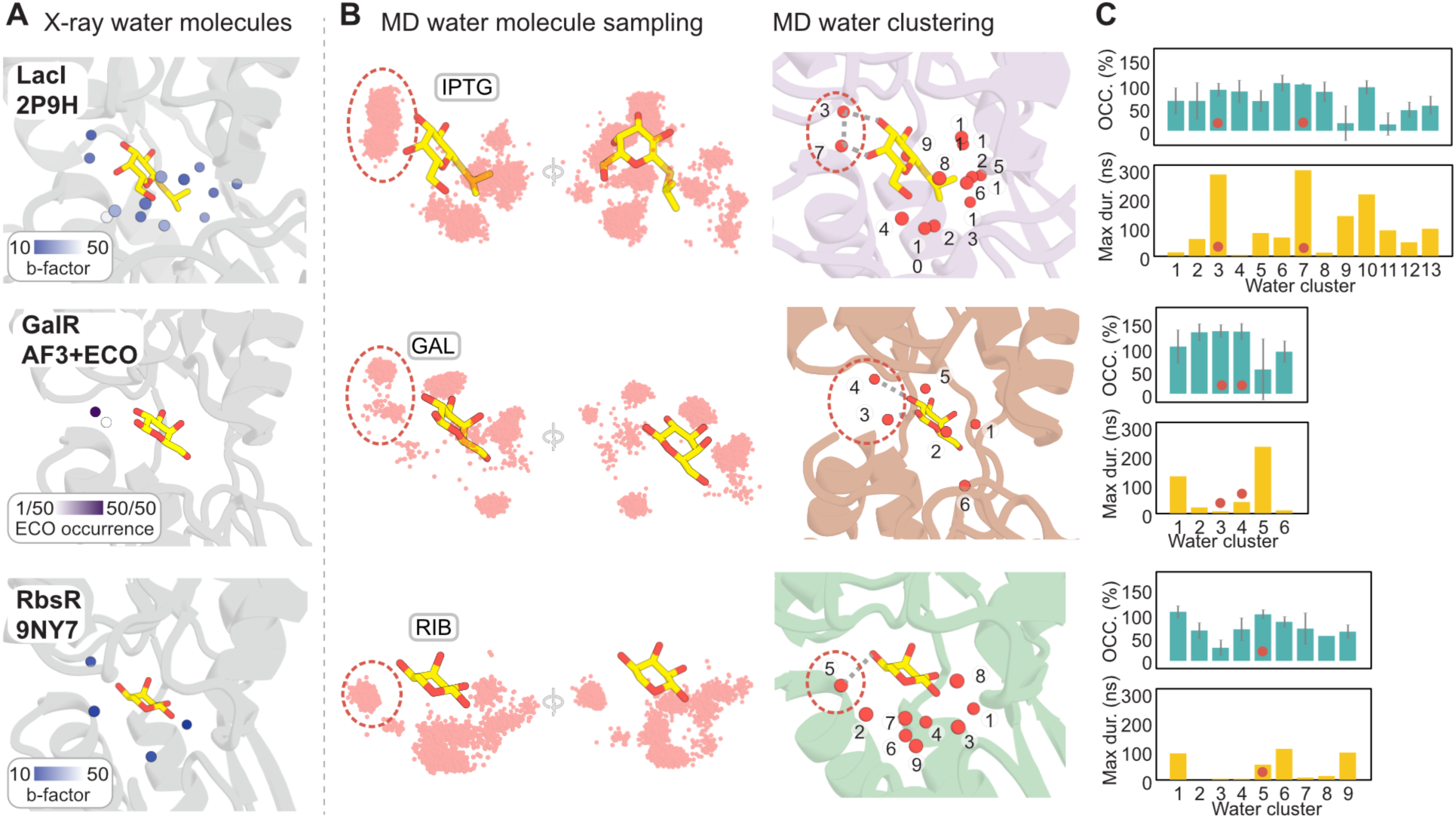
Structural waters in TF induction. **(A)** Binding pocket waters in TF structures and models. **(B)** Waters sampled in MD simulations cluster together. **(C)** Durations and occupancies for each water cluster in MD simulations. Red dots correspond to circled structural waters in (B).

## Molecular switches control TF states

PBPs respond to sugars with a hinge-mediated conformational change that is visible in structures (Fig. S17). Sugar binding stabilizes them in a “closed” conformational ensemble, with locked inter-lobe loops forming a perfect binding interface for sugar transporters. But in TFs, this region instead forms the dimer interface. The operator- and sugar-bound states of the TFs have incompatible energetic requirements along the dimer and VFT-DBD interfaces, and the transition between states is less pronounced in structural models and distinct from the PBPs. Ultimately, the TFs evolved an energetic blueprint that is not a smaller-scale version of the PBP response to sugars, but which is instead unique to the family (Figs. 6, S25).

**Figure 6.**
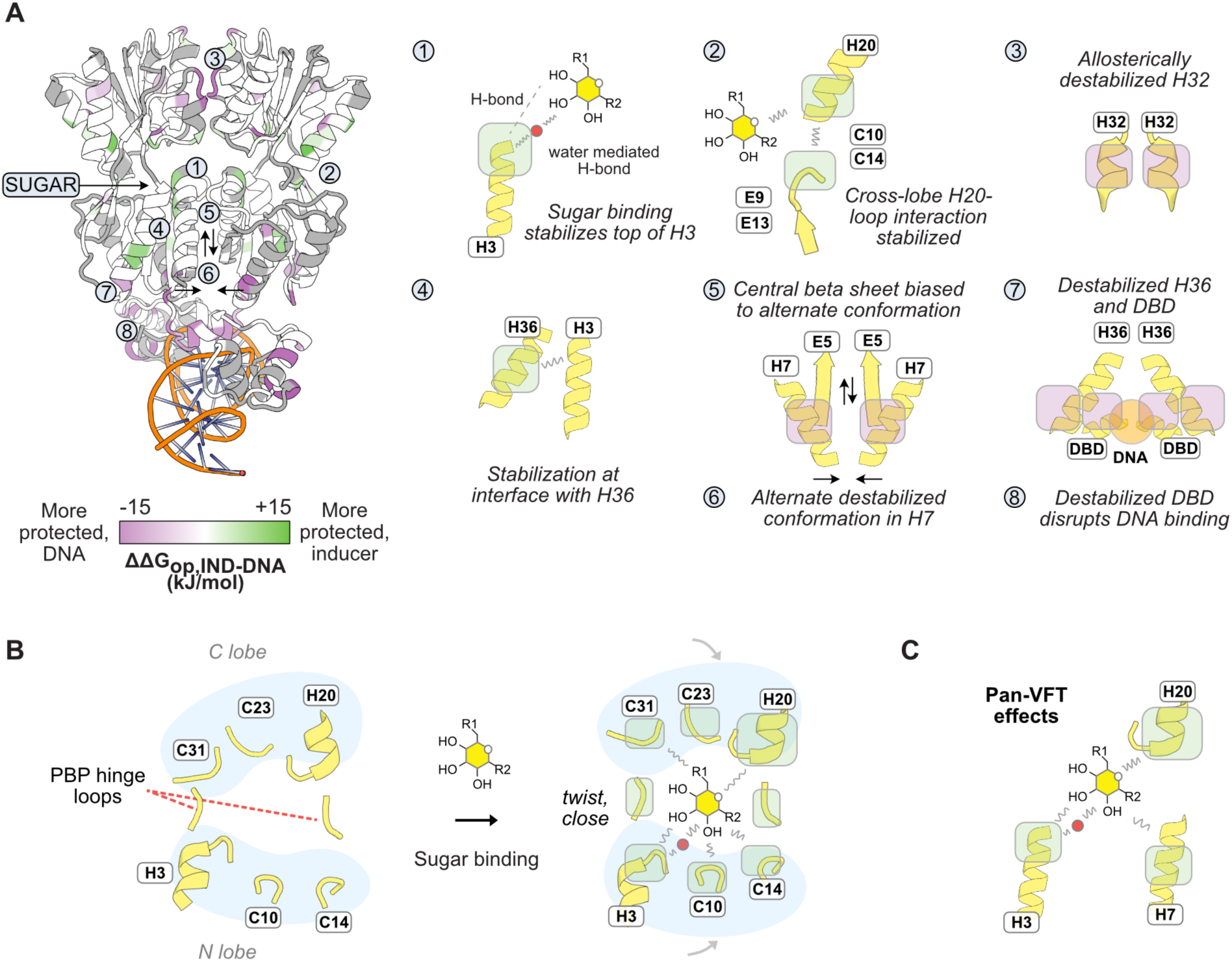
Model for the induction mechanism in LacI/GalR TFs. **(A)** The model is based on conserved ΔΔG_op, IND-DNA_ (*left*), with molecular switches labeled and schematized on the *right*. (**B)** Conserved mechanism of sugar response in both PBPs. **(C)** Pan-VFT conserved energetic relationships center on sugar interactions with binding pocket helices.

A set of “molecular switches” transition the LacI/GalR TFs to the induced state by restructuring the hinge helix^38^, but also, crucially, the VFT dimer interface, VFT cross-lobe interactions, and the VFT-DBD interface (Figs. 6A, S25, Supplemental note 2). Inducer- and operator-driven conformational changes at switch positions have substantial ΔΔG_op,DNA-ind_ of 10-20 kJ/mol. Based on the HX/MS data, we propose the following energetic changes in no specific order, as these experiments only report on energetic changes to the equilibrium state ensemble. Inducer binding to the DNA-bound TFs rigidifies the transducer helix at its N-terminus at the expense of the stability of the C-terminus, stabilizes inter-lobe interactions, and destabilizes the metastable helix, leading to changes in the local structure and monomer-dimer equilibrium and thereby reduced DNA binding. Simultaneously, the hydrogen bond network in the N-terminal VFT beta sheet is rearranged, and several helices adjacent to the DBD are restructured. In the VFT-DBD interface, inducer binding stabilizes H36 residues to reduce hydrophobic interactions with the loop that joins the hinge helix to the VFT domain, thereby destabilizing the DBD. Reduced hydrogen bonding between H7 and the hinge helices releases them from the minor groove of the operator DNA, linearizing it. In support of this mechanism, mutations in this region modulate LacI-operator DNA affinity.^21,56–58^ Destabilization of the DBD is conserved in the induced state, resulting in decreased ΔG_op_ at recognition helix positions key to operator specificity.^43^ Additionally, nonspecific charged interactions with DNA are destabilized, as are DBD helical packing interactions. While sugars stabilize some of the binding pocket helices in both PBPs and TFs, few of these long-range effects that define the energetic blueprint of TFs are manifested in PBPs (Fig. 6B, C).

In support of the functional role of these evolutionarily conserved energetic changes, mutationally sensitive positions identified by cellular phenotype assays of a LacI point mutant library^50^ are enriched in the set of residues that are stabilized by binding partners. By categorizing phenotypes into three classes, WT-like EC₅₀ (1.5–2.5), lower EC₅₀ (<1.5), and higher EC₅₀ (>2.5), we observed distinct enrichment patterns: mutations in the lower EC₅₀ class preferentially occur at residues with negative ΔΔG_op, (IPTG-DNA)_, whereas those in the higher EC₅₀ class cluster at residues with positive ΔΔG_op, (IPTG-DNA)_. These results indicate that destabilizing the DNA-bound state lowers the IPTG concentration required for induction, whereas perturbing the IPTG-bound state reduces IPTG sensitivity (Fig. S26).

## Discussion

Coordinating switching between operator DNA- and specific sugar-bound states for LacI/GalR TFs is a life-or-death necessity for bacteria. Here, we discovered how the distribution of ensemble energies in the LacI/GalR TFs is dramatically different when bound to operators vs. sugars, though their X-ray crystal structures are similar. We found specific residues in the TF family that behave as molecular switches by occupying different, mutually exclusive energies in different functional states. By the evolutionary conservation of these switches and their state-specific interactions with protein, ligand, and water molecules, the TFs undergo a shift in their conformational ensembles that forbids prolonged operator DNA binding simultaneously with inducer binding.

The VFT fold is not unique to LacI/GalR TFs. By comparing the TFs to their distant cousins, the PBPs, we determined how one protein fold performs two different functions in response to the same small molecule ligands. This yields a quantitative description of how specific energetic relationships persisted and adapted over the evolutionary timescale to protect functionally important conformational changes from being lost, even as new chemical triggers challenged the remarkable plasticity of the fold. We can use this blueprint to rationally engineer VFT-fold sensors, which has historically proven challenging for both TFs and PBPs despite their promise in biotechnology.^46,49–53,59–61^ Towards this goal, we determined a simple function for computationally designing TF variants with inducer sensitivities spanning four orders of magnitude by identifying the functional role of a crucial helix in the family’s energetic blueprint. This computational design strategy augments the limited toolbox for rationally engineering allosteric proteins, especially when more than 2-3 mutations are required simultaneously.

The story of conserved energetic relationships in a classic TF and its cousins demonstrates how regulators of the genome have evolved alongside the genome itself, and that the energetic blueprint for the family, buried in the evolutionary record, is key to designing transcriptional regulators *de novo*. Previous efforts to design proteins that bind to DNA sequences specifically and with high affinity are mired by the repetitive degeneracy of DNA structures and a bias towards modeling linear DNA, without considering that protein-DNA interactions can affect DNA conformation. For example, LacI/GalR TFs can bind multivalently to form loops in the genomic DNA.^62,63^ In one recent effort to design DNA-binding proteins, fourteen sequence-specific protein binders were identified from screening more than 140,000 design models, many of which were designed with an experimentally solved target DNA structure available.^64^ Perhaps designing transcription factors *de novo* requires considering both how the TF changes the conformational ensemble of DNA, and how different DNA sequences reshape the conformational ensemble of the TF.

## Acknowledgements

We thank members of the Glasgow Lab for discussions, Dr. Wayne Hendrickson for advice on X-ray crystallography, Dr. Naomi Latorraca for feedback on the manuscript, Dr. David Ross for advance access to deep mutational scanning data, and Dr. Rinat Abzalimov at the Advanced Science Research Center at the City University of New York in his role as the MS Facility Manager. We thank the staff at NYX beamline at NSLS-II for their support during data collection, Dr. Jerry Chang from Columbia’s Precision Biomolecular Characterization Facility for help with mass photometry experiments, as well as Drs. Fabiana Bahna and Larry Shapiro for help with SEC-MALS experiments using their instrument. We acknowledge support from the National Institutes of Health (R00GM135529 and R35GM157185 to AG). KCW was funded by a National Science Foundation (NSF) Graduate Research Fellowship. ECA was funded by the NSF RaMP in Biomolecular Structure Prediction and Design (award 2216011).

## Author contributions

* These authors made equal contributions. MLW collected the HX/MS data. MLW, CL, and AG analyzed the HX/MS data. CL performed MD simulations and analysis. DS performed the bioinformatics and computational solvation analysis. KCW grew the protein crystals and solved the X-ray crystal structures with help from CL and ZG. ECA designed the LacI variants with help from EC and AG, and experimentally assayed their behavior. AG conceived and supervised the study. MLW, CL, DS, KCW, ECA, and AG made the figures and wrote the manuscript with input from all authors.

## Data availability statement

The HX/MS deuterium uptake plots, FEATHER-derived centroid fits to HX/MS data, and a PyMOL session file showing protein-specific differences in ΔG_op_ are available in the Supplementary Materials. The HX/MS data have been deposited to the ProteomeXchange Consortium via the PRIDE^65^ partner repository with the dataset identifier PXD062370. The X-ray crystal structures are uploaded to the PDB with codes 9NY7 and 9NY8. The AlphaFold 3-RosettaECO models, MD trajectories, conservation score matrices, gene trees, MSA files, structural alignment and X-ray diffraction images are available in Zenodo (dataset identifier 10.5281/zenodo.15091462). Supplementary Information is available for this paper. Correspondence and requests for materials should be addressed to Anum Glasgow.

## Code availability statement

The SI includes all code used in this study. The PIGEON-FEATHER source code is available at https://github.com/glasgowlab/PIGEON-FEATHER, along with the synthetic datasets used for benchmarking, an example HX/MS dataset, and a tutorial.

## Methods

### Calculation of conservation scores

#### Multiple sequence alignment (MSA)

The MSA was generated using MMSeqs2^66^ separately for TFs (LacI as the query) and PBPs (RbsB as the query) with bacterial sequences from UniRef90 as the search database, using the following command line:

*mmseqs filtertaxseqdb ./Uniref90_DB UniRef90_bacteria --taxon-list 2 -s 6 --max-seqs 20000 -a 1 -e 0.01 --min-aln-len 150 --allow-deletion 1 --threads 60.*

#### Conservation scores

We used the global epistatic model GEMME^67^ which takes into account both MSA and evolutionary history of the sequences using the MSA generated above. We used the following command line with the provided docker:

*docker run -ti --rm --mount type=bind,source=$PWD,target=/project elodielaine/gemme:gemme then*
*python2.7 $GEMME_PATH/gemme.py MSA.fasta -r input -f MSA.fasta*

The combined prediction scores were normalized within each group of proteins (TFs or PBPs) such that the highest conservation score is 1.

### Gene tree construction

For the input sequence alignment, we combined MSAs from the previous step and added the sequence for LuxP (Uniprot P54300) to initially use as an outgroup^7^. We used only the VFT portion of the MSA and further excluded positions with >10% gaps. We used IQTree^68^ for the tree construction. We estimated a substitution model using the following command line:

*iqtree -s ./MSA.fasta -m MFP -B 1000 -bnni -T AUTO .*

We chose Q.pfam+R10^69^ as the best-fit model based on Bayesian information criterion (BIC)^70^ and ran the following command:

*iqtree -s ./MSA.fasta -m Q.pfam+R10 -B 1000 -bnni -T AUTO -safe -cptime 600.*

### Structural alignment of TFs

We collected experimentally solved structures of LacI/GalR TFs from the Protein Data Bank (PDB)^71^ using the following query:

*((Lineage Name = "GalR/LacI-like bacterial regulator" AND Annotation Type = "SCOP") OR (Lineage Name HAS ANY OF WORDS "LacI,GalR" AND Annotation Type = "ECOD") OR (Lineage Identifier = "IPR046335" AND Annotation Type = "InterPro")) AND Polymer Entity Sequence Length > 90*

The pairwise RMSD was calculated using USAlign.^72^

### Plasmid construction

DNA sequences encoding RbsR and GalR were synthesized as double-stranded genes by IDT DNA and cloned into a pET9a vector with an N-terminal 6× histidine tag and TEV protease sequence using a Golden Gate strategy with type IIs restriction enzyme BsaI. In the case of LacI, we used a previously described plasmid encoding the dimeric form of the protein (residues 1-331) with the same architecture.^19,49^ The MglB and RbsB expression plasmids were also constructed with N-terminal 6× histidine tag and TEV site using the same strategy in the same vector, but the genes were amplified by polymerase chain reaction from the *E. coli* DH10B genome. All genetic constructs were confirmed by sequencing. All gene sequences are available in Appendix 2.

### Protein expression and purification

#### LacI

We introduced a 6×his-LacI1-331 gene amplified from the *E. coli* DH10B genome into a modified pET9a plasmid via Golden Gate cloning using a BsaI restriction endonuclease (NEB) to produce the pET9a-6×his LacI1-331 plasmid, as described previously^19,73^. This construct encodes a dimeric LacI without the tetramerization domain and with an N-terminal 6×-histidine tag. We chemically transformed the pET9a-6×his-LacI1-331 plasmid into a BL21-AI Δ*lacI* strain and a BL21 LOBSTR strain. Cells were grown at 37 °C overnight shaking at 200 rpm in LB media with 50 µg/ml kanamycin, then subcultured 1/100 into 1 L LB media with 50 µg/ml kanamycin and grown at 37 °C until the optical density at 600 nm was measured between 0.2 and 0.4. Protein expression was induced by the addition of 0.2% L-arabinose by weight (for replicates 1 and 2) for BL21-AI cells or 1 mM isopropyl ß-D-1-thiogalactopyranoside (IPTG) for LOBTSR cells (replicates 3 and 4), and the cells were grown at 16 °C for 12-16 hrs. The culture was centrifuged at 6000 × *g* for 20 minutes and the cell pellet was frozen at -20 °C. After several weeks, the pellet was thawed in 30 ml lysis buffer: 50 mM Tris, 30 mM NaCl, 5 mM MgCl_2_, 1 mM MnCl_2_, 100 µM CaCl_2_, 2 mM EDTA, with a protease inhibitor cocktail (Pierce), 1 mg/ml lysozyme, and 2000 units/ml DNase I (Sigma-Aldrich). The mixture was lysed at 37 °C for 15 minutes prior to centrifugation at 27,000 × *g* for 30 minutes at 4 °C. The soluble fraction was filtered and applied to 2 ml Ni-NTA resin (Thermo Fisher) in 2-5 ml wash buffer (50 mM Tris, 150 mM NaCl, 20 mM imidazole, pH 8.0), and the mixture left to nutate overnight at 4 °C. The mixture was then added to a 20 ml benchtop gravity column, allowed to drain and settle, and washed with 50 ml of wash buffer. The protein was eluted in 15 ml of elution buffer (50 mM Tris, 150 mM NaCl, 250 mM imidazole, pH 8.0) and concentrated to 1.5 ml using a 15 ml ultracentrifugal filter column with a 10 KDa MWCO (MilliporeSigma). To remove the 6×his tag, 10 µl TEV protease (MilliporeSigma) was added to the concentrated protein solution, and the reaction was incubated at 30 °C for 1 hour. The sample was injected on a Bio-Rad NGC™ medium-pressure chromatography system and applied to a Cytiva S100 size-exclusion chromatography (SEC) column in sample buffer (50 mM Tris, 150 mM NaCl, pH 8.0). Fractions corresponding to an A280 peak of the correct size were collected, pooled, concentrated to 3.8 mg/ml, and flash-frozen in aliquots at -80 °C.

#### RbsR

We introduced a 6×his-RbsR gene into a modified pET9a plasmid via Golden Gate cloning using a BsaI restriction endonuclease (NEB) to produce the pET9a-6×his-RbsR plasmid. This construct encodes full-length RbsR with an N-terminal 6×-histidine tag. Protein expression, purification, and storage was conducted as described for LacI in LOBSTR cells for all replicates.

#### GalR

We introduced a 6×his-GalR gene into a modified pET9a plasmid via Golden Gate cloning using a BsaI restriction endonuclease (NEB) to produce the pET9a-6×his-GalR plasmid. This construct encodes full-length GalR with an N-terminal 6×-histidine tag. We chemically transformed the pET9a-6×his-GalR plasmid into a BL21-AI strain. Cells were grown at 37 °C overnight shaking at 200 rpm in LB media with 50 µg/ml kanamycin, then subcultured 1/100 into 1 L LB media with 50 µg/ml kanamycin and grown at 37 °C until the optical density at 600 nm was measured between 0.2 and 0.4. Protein expression was induced by the addition of 0.2% L-arabinose by weight and the cells were grown at 16 °C for 12-16 hrs. The culture was centrifuged at 6000 × *g* for 20 minutes and the cell pellet was frozen at -20 °C. After several weeks, the pellet was thawed in 30 ml lysis buffer: 50 mM Tris, 30 mM NaCl, 5 mM MgCl2, 1 mM MnCl2, 100 µM CaCl2, 2 mM EDTA, with a protease inhibitor cocktail (Pierce), 1 mg/ml lysozyme, and 2000 units/ml DNase I (Sigma-Aldrich). The mixture was lysed at 37 °C for 15 minutes prior to centrifugation at 27,000 × *g* for 30 minutes at 4 °C. The soluble fraction was filtered and applied to 2 ml Ni-NTA resin (Thermo Fisher) in 2-5 ml wash buffer (50 mM Tris, 600 mM NaCl, 20 mM imidazole, 0.5mM tris(2-carboxyethyl)phosphine (TCEP), pH 8.0), and the mixture left to nutate overnight at 4 °C. The mixture was then added to a 20 ml benchtop gravity column, allowed to drain and settle, and washed with 50 ml of wash buffer. The protein was eluted in 15 ml of elution buffer (50 mM Tris, 600 mM NaCl, 0.5MM TCEP, 250 mM imidazole, pH 8.0). Because GalR precipitates under spin concentration (even in the 600 mM NaCl buffer) we used this eluate in HX/MS experiments without further concentration or SEC, and without freezing.

#### RbsB

We introduced a 6×his-RbsB (residues 26-296) gene amplified from the *E. coli* DH10B genome by colony PCR into a modified pET9a plasmid via Gibson cloning using NcoI and SacI restriction endonucleases (NEB) to produce the pET9a-6×his-RbsB26-296 plasmid. This construct encodes RbsB without the N-terminal signal peptide and with an N-terminal 6×his tag. We chemically transformed the pET9a-6×his-RbsB26-296 plasmid into a BL21 LOBSTR strain. Cells were grown at 37 °C overnight shaking at 200 rpm in LB media with 50 µg/ml kanamycin, then subcultured 1/100 into 1 L LB media with 50 µg/ml kanamycin and grown at 37 °C until the OD at 600 nm was measured between 0.2 and 0.4. Protein expression was induced by the addition of 1 mM IPTG and the cells were grown at 16 °C for 12-16 hrs. The culture was centrifuged at 6000 × *g* for 20 minutes and the cell pellet was frozen at -20 °C. After several weeks, the pellet was thawed in 30 ml lysis buffer and purified as described above for LacI. The samples were then concentrated to 1.5 ml using a 15 ml ultracentrifugal filter column with a 10 KDa MWCO (MilliporeSigma). Concentrated eluate was injected on a Bio-Rad NGC™ medium-pressure chromatography system and applied to a Cytiva S100 SEC column in sample buffer (50 mM Tris, 150 mM NaCl, pH 8.0). Fractions corresponding to an A280 peak of the correct size were collected, pooled, concentrated to 0.83 mg/ml, and flash-frozen in aliquots at -80 °C.

#### MglB

We introduced a 6×his-MglB gene amplified from the *E. coli* DH10B genome into a modified pET9a plasmid via Golden Gate cloning to produce the pET9a-6×his-MglB plasmid and chemically transformed it into BL21 LOBSTR cells. Cells were grown and the protein was purified as described for LacI. SEC fractions corresponding to an A280 peak of the correct size were collected, pooled, concentrated to 2 mg/ml, and flash-frozen in aliquots at -80 °C.

### X-ray crystallography sample preparation

To prepare the *rbsO-*RbsR sample, self-complementary single-stranded 30-base-pair primers encoding *rbsO* (5′-GTGGGTCAGCGAAACGTTTCGCTGATGGAG-3′) were synthesized (IDT DNA) and annealed using a previously described protocol.^19^ RbsR dimer (10 mg/mL) was then incubated at 4 °C for 12 h with a two-fold molar excess of the annealed DNA. After incubation, the mixture was centrifuged (14,000 rpm, 4 °C, 10 min), and crystals were grown by hanging-drop vapor diffusion using 400 μL of the protein–DNA complex solution and 300 μL of reservoir solution containing 0.2 M sodium sulfate, 0.1 M bis-tris-propane (pH 6.5), and 20% PEG 3350 (PACT Premier 2-20) at 20 °C.

To prepare the ribose-RbsR sample, RbsR (10.1 mg/mL) was incubated with a six-fold molar excess of ribose (1.3 mM) for 12 hours at 4 °C and subsequently centrifuged (14,000 rpm, 4 °C, 10 min). The resulting mixture was crystallized by hanging-drop vapor diffusion using 400 μL of the protein–ribose complex solution and 300 μL of reservoir solution containing 0.2 M sodium malonate (pH 4.0) and 20% PEG 3350 (Pegion2 A5) at 20 °C.

### X-ray crystallography data collection and model building

Diffraction data were collected at the NYX beamline 19-ID of the National Synchrotron Light Source II (NSLS-II) at Brookhaven National Laboratory. The raw diffraction HDF5 images were processed using Xia2^74^ with DIALS^75^ pipeline, as implemented in CCP4 suite^76^, for ribose-bound RbsR structure and XDS^77^ for *rbsO-*bound structure to perform indexing, integration and initial scaling. Data scaling and merging were carried out using AIMLESS or STARANISO^78^. Molecular replacement was performed with Phaser^79^, using an AlphaFold2 model^80^ as the search model for the ribose-bound RbsR structure, and an AlphaFold3 model^40^ as the input model for the *rbsO-*bound structure. Coot^81^ was used for manual modelling. Structural refinement was conducted using both Phenix.refine^82^ and REFMAC.^83^ Final validation and refinement were completed for both using PDB-Redo.^84^

### Size-exclusion chromatography with multi-angle light scattering (SEC-MALS)

SEC-MALS analyses were performed using a Superdex 200 Increase 3.2/300 SEC column on an ÄKTA FPLC system (GE Healthcare). The column was connected in-line to a multi-angle light scattering detector (DAWN Heleos II, Wyatt Technology), a differential refractive index detector (Optilab rEX, Wyatt Technology), and a UV absorbance detector. Purified proteins were diluted to 75 µM in running buffer (50 mM Tris-HCl, pH 8.0, 150 mM NaCl), with 10 mM IPTG or ribose added as indicated. Samples of 200 µL volume were injected and eluted at a flow rate of 0.5 mL/min at 25 °C, yielding collected fractions at approximately 3 µM protein. All samples eluted as single peaks (Figure S9). Data were analyzed using ASTRA software (Wyatt Technology).

### Mass photometry

For each measurement, after instrument calibration, 2 µL of protein sample (pre-incubated with ligand as needed) at 100 nM was diluted into 18 µL running buffer (50 mM Tris-HCl, pH 8.0, 150 mM NaCl) supplemented with 10 mM IPTG or ribose where indicated. The diluted buffer was applied to a coverslip and placed in the chamber of a Refeyn TwoMP mass photometer (Refeyn Ltd., Oxford, UK). After autofocus stabilization, 2 µL of protein sample was added to the droplet on the cassette and mixed by pipetting three times. Subsequently, 60 s movies were recorded. Data acquisition was performed using AcquireMP (Refeyn Ltd.) and processed with DiscoverMP (Refeyn Ltd.).

### HX/MS sample preparation and experiment

Sample preparation was performed as described previously^20^, except for the following variables: sample buffer composition, protein stock composition, and quench composition, as specified in Tables S3-S7. All proteins except GalR were purified by size exclusion chromatography, concentrated to 5-10 µM, and flash-frozen in aliquots; all proteins were confirmed to be >95% pure by SDS-PAGE. HX/MS experiments were performed by diluting freshly purified proteins or thawed aliquots by tenfold in an equivalent Tris-NaCl D_2_O buffer at pH 8.0. GalR was stored in 600 mM NaCl, and HX/MS experiments were also conducted with 600 nM NaCl, unlike other proteins for which 150 mM NaCl was used. This is because GalR forms filamentous aggregates and precipitates at low salt concentrations^85^. Consequently, DNA association rates for GalR are lower than they would be at 150 mM NaCl^86^. Since GalR affinity for its operator has been measured in 300 mM KCl at 4 nM^87^, and twofold changes in salt concentration have less than order-of-magnitude effects on GalR k_a_ for its operator^86^, it is probable that the large majority of GalR remains bound to DNA in 600 mM NaCl at the µM concentrations used in these experiments. IPTG, galactose, and ribose were added to protein samples at 10 mM. Operator DNAs were added at twofold molar excess of TF dimer concentrations, except for the single-repeat control, where we used a fourfold molar excess to maintain the same concentration of operator sequence. HX experiments were performed as described previously for LacI^20^ except for the following variables: reaction and injection volumes, and protease columns, specified in Tables S3-S7. After incubation in D_2_O buffer, the exchange reaction was quenched by 1.9-fold sample dilution in quench buffer (3% acetonitrile, 1% formic acid in H_2_O, with GuHCl concentrations noted in Tables S3-S7 for all proteins, all MS-grade, Fisher Scientific).

### HX/MS fully deuterated sample preparation

Fully deuterated controls were prepared as previously described^20^ (Tables S3-S7). Of note, due to some observed aggregation, we removed GuHCl from the preparation of fully deuterated control samples for GalR samples. We allowed them to exchange at 37 °C for at least 24 hours and used a quench buffer containing GuHCl as in the other HX experiments. No fully deuterated control was used for the single-repeat DNA control.

The protein stock solutions were diluted ten-fold in deuterated buffer and exchanged at 15 °C for the specified time, then diluted in quench solution (75 µl, 3 M GuHCl, 3% acetonitrile, 1% formic acid in H_2_O, all MS-grade, Fisher Scientific) of equal volume to the deuterated buffer at 2 °C. All but 10 µl of quenched reaction was injected into the protease column at 7 °C. Solvent A (3% acetonitrile, 0.15% formic acid in H_2_O) was pumped (UltiMate 3000, ThermoFisher Scientific) at 150 μL/min through the protease column. Peptides eluted from the protease column were trapped on a 1×10 mm C18 column (Hypersil GOLD, 3 μM pore size, ThermoFisher Scientific).

### HX/MS data analysis

MS/MS peak picking and deconvolution, disambiguation by peak picking, peptide disambiguation by PIGEON, FEATHER rate fitting, PIGEON error analysis, and quality control of all fit spectra were performed as described previously for LacI.^20^ Conservation among FEATHER-derived ΔG_op_ was determined by treating as nonzero only state-state differences that exceeded our significance threshold and occurred in the same direction for all proteins in the comparison set. We calculated the mean of these positional ΔΔG_op_, wherever mean{ΔΔG_op_} > 2 kJ/mol. For a given protein, state-state differences at each site were considered significant if the “mini-peptide” (the set of residues including that site covered indistinguishably by a set of fit exchange rates) was the same for both states and satisfied *mean*{ΔG_op,*a*,i_-ΔG_op,*b*,i_} > *std*{ΔG_op,*a*,i_-ΔG_op,*b*,i_}, where the {ΔG_op,*a*,i_} are the sorted ΔG_op_ corresponding to that site for each exchange rate in the mini-peptide, in state *a*. In all structural models presented in this work that are colored by ΔΔG_op_ conserved among 2+ proteins, we calculated the average positional ΔΔG_op_ for those proteins, if the selection criteria described above were met. Otherwise, we colored those positions white to indicate that the exchange behavior was not conserved. Gray coloring indicates a lack of data for at least one protein or state, or high standard deviation (noise) in the significance calculation. The full sets of ΔG_op_, their standard deviations, and their resolutions are available for all proteins in Tables S8-S12. For the single LacI binding site DNA control experiment, FEATHER rate fitting was omitted and data were compared qualitatively at the peptide level.

### Preparation of LacI structure for design

To design the metastable helix of the LacI dimer, we used the 2.60 Å resolution X-ray structure of LacI bound to the operator and the anti-inducer ONPF (PDB ID 1EFA). This structure was downloaded from the PDB, and the DNA binding domain (residues 1-60), the DNA, and ONPF were manually removed from the structure in PyMOL.^88^ The truncated structure was relaxed in the Rosetta software suite with coordinate constraints on backbone and side chain heavy atoms and used as the input structure for computational design.

### Sequence design and filtering

The truncated structure of the LacI regulatory domain was mutated in seven residues within the metastable helix of the dimer interface (E277, D278, S279, S280, C281, Y282, I283), and mutations were paired across the homodimer. Two iterations of ProteinMPNN^89^ at sampling temperature 0.5 were used for sequence design in the metastable helix, with each iteration generating 30 designs. We retained the top 15 designs of each iteration as ranked by the Rosetta InterfaceAnalyzer metric dg_separated/dSASA to evaluate the stability of the dimer interface. After two iterations of design and filtering by dg_separated/dSASA, 225 designs were made and scored by Rosetta metrics interface_energy and a calculated per_residue_energy sum of metastable helix residues; this protocol was done 3 times producing 675 designs total. The top six designs to experimentally test for IPTG sensitivity were chosen based on the metrics, interface_energy, helix_energy, and number of hydrogen bonds in the dimer interface. Designs 3, 4, 5, and 6 were produced with the design protocol above. Design 2 was produced with the same design protocol except for the addition of a ProteinMPNN amino acid bias that favors polar residues. Design 1 was produced with the same design protocol above, but five residues that interact with the metastable helix (S221, M223, L251, M254, R255) were additionally allowed to mutate during ProteinMPNN iterations.

### Metastable helix stability calculation

The metastable helix stability score is the sum of four weighted Rosetta metrics:

*Stability score = -(0.3*interface_energy + 0.7*helix_energy)/(hbond_count + 10*unfavorable_energy_sum)*

The four metrics are defined as such:

1. interface_energy = sum of Rosetta pairwise residue energies across all residue pairs (R1, R2) where R1 is residue on LacI dimer chain A and R2 is a residue on LacI dimer chain B
2. helix_energy = sum of the per-residue Rosetta energies for the 14 metastable helix residues across the LacI dimer
3. hbond_count = total number of cross interface hydrogen bonds in LacI dimer interface scored by Rosetta
4. unfavorable_energy_sum = sum of the 5 highest pairwise residue energies that make up the interface energy

Since higher, more positive stability scores are best, score terms to be maximized (interface_energy and helix_energy) were placed in the numerator and converted to positive values. Score terms to be minimized (hbond_count and unfavorable_energy_sum) were placed in the denominator and kept as positive to ensure consistency in the direction of optimization. Weighting was assigned to both numerator and denominator terms: numerator weights assign helix_energy with higher significance to best score the stability of the metastable helix region. Denominator weights were assigned based on the order of magnitude of the raw metric values; the unfavorable_energy_sum was multiplied by 10 so as not to be overwhelmed by the hbond_count when evaluating the designs. The calculated values of each metric for the six chosen designs and WT LacI are available in Table S13.

### Cell culture assay to measure LacI response to IPTG

The assay was performed as previously described^19,49^ with the following modifications. Each of our LacI designs were chemically transformed into *E.coli* DH10B cells that contained a plasmid encoding superfolder GFP (sfGFP) under the control of *lacO* and plated on Lysogeny broth (LB) agar with 50 µg/ml kanamycin and spectinomycin. One 5 ml culture for each design was grown from a single colony overnight in LB medium with antibiotics at the same concentration. New 500 µl cultures were subcultured in LB-kanamycin-spectinomycin from the overnight culture and grown at 37 °C and shaking at 200 rpm for 2 hours in 96-well blocks. IPTG was then added to cultures to final concentrations ranging from 0.001 µM to 10,000 µM. After 6 hours, the cultures were centrifuged at 3,000 rpm for five minutes and the supernatant was decanted. Pellets were resuspended in 200 µl phosphate-buffered saline (PBS) and diluted 1:10 in 100 µl PBS in a clear-bottom 96-well microtiter plate for plate reading. A Biotek Synergy Neo2 plate reader was used with an excitation wavelength of 485 nm, an emission wavelength of 528 nm, and a gain setting of 100. The fluorescence was normalized by absorbance measured at OD600 before analysis. Replicates were performed on nine different days, subculturing from new overnight cultures on each day, with each IPTG concentration tested three to nine times. Data were pooled and fitted using the logistic curve fitting function in SciPy, as shown in Extended Data Figure 5.

### Structural water prediction using RosettaECO

The experimental structures or AlphaFold 3 (AF3) models^40^ were first relaxed using the Rosetta biomolecular modeling suite^90^. The experimental structures were relaxed with backbone and sidechain constraints using the following command line to produce three models for each input file:

*./relax.default.linuxgccrelease -s input.pdb -out:suffix.constrain.relax -nstruct 3 - relax:default_repeats 5 -out:path:pdb ./ -out:path:score ./ -in:file:extra_res_fa ./tre.params - constrain_relax_to_start_coords true -relax:coord_constrain_sidechains true*

Three models were produced for each input protein. The structural model with lowest score in Rosetta Energy Units (REU) was chosen as the best model. For the AF3 models, we performed unconstrained relaxation using the command:

*./relax.default.linuxgccrelease -s input.pdb -out:suffix .relax -nstruct 3 -relax:default_repeats 5 - out:path:pdb ./ -out:path:score ./ -in:file:extra_res_fa ./param_fle.params*

The ligand was specified through the “param_file.params” file where needed. To solvate the models using Rosetta Explicit Consideration of cOordinated water (RosettaECO)^54^, we used an .xml protocol (see **Data availability**) to generate 50 solvated models per input unsolvated relaxed model:

*./rosetta_scripts.default.linuxgccrelease -parser:protocol solvate.xml -s input.pdb -nstruct 50 - beta_nov16 -out:suffix .solvate -out:path:pdb ./ -in:file:extra_res_fa ./param_file.params*

Water molecules within 0.5 Å of each other were clustered and reported in the final solvated model (see **Data availability**; the number of waters in each model is reported as “b-factor” in the final .pdb file).

### Molecular dynamics simulations

Five sugar-bound VFT-fold systems (LacI-IPTG, GalR-galactose, RbsR-ribose, MglB-galactose, and RbsB-ribose) were built for MD simulations using the following structures by PDB ID: 2P9H, AF3 model, 9NY7, 1GLG, and 2DRI as the input models, respectively. All simulations were performed using the GROMACS 2022 software package with the CHARMM36 force field^91^.

Ligand parameters were generated using the CHARMM General Force Field (CGenFF)^92^. We energy minimized the systems using the steepest descent minimization method until the maximum force was less than 1000 kJ/mol/nm^2^, followed by 100 ps restrained MD simulations in NVT ensemble by constraining the heavy atoms of the protein and ligand at 298.15 K. The resulting simulation system was further equilibrated for a 1000 ps NPT simulation at 1 atm and 298.15 K with 1000 kJ/mol/nm^2^ position restraints on the protein and ligand. Finally, we conducted the NPT production MD simulations using the same conditions without restraints, with a time step of 2 fs, nonbonded cutoff of 12 Å, and particle-mesh Ewald long-range electrostatics. Five 300 ns simulation replicates were collected for each system. The MD trajectories were analyzed using MDAnalysis^93^, VMD^94^, or MDTraj^95^.

### Water clustering analysis

The trajectories from the production runs for each system were aligned to its initial structure based on the protein backbone atoms, and the coordinates of water molecules located within 3.5 Å of the ligands were extracted. These coordinates were subsequently clustered using the *k-means* algorithm implemented in the scikit-learn^96^ Python package. The number of the clusters was determined by the elbow method.

### LacI designed variants SDS-PAGE and western blot

Frozen LacI designed variant strains, the wild-type strain, and the control Δ*lacI* strain were cultured overnight in 5ml of LB media containing 50 µg/ml kanamycin and spectinomycin for the wild-type and the designed variants and only kanamycin for the control strain. The strains were grown overnight at 37 °C and 200 rpm. The next day, cultures were subcultured (1:100 dilution) in 5 ml LB media and grown for two hours (37 °C, 200 rpm), followed by induction with IPTG to a final concentration of 1 mM for an additional six hours. The optical density (OD) at 600 nm of all cultures was then measured and standardized to a final OD of 1.5. We collected 100 µl for each culture and spun it down at 3000 × *g* for 5 minutes. The supernatant was removed and the cell pellet was resuspended in 16 µl water mixed with 16 µl 2x Laemmli-buffer. The samples were placed at 95 °C for 10 minutes to promote cellular lysis and protein denaturation.

After lysis, 15 µl of each sample was loaded on two stain-free TGX gels (Bio-Rad) alongside the protein ladder. One of the gels was used for total protein visualization via Coomassie blue staining, while the second one was used for western blot analysis. These gels were run at 95-100 volts for about an hour in SDS buffer. Bands were transferred to a PVDF membrane via a Bio-Rad Turboblot system using the 7 minute mini gels, mixed MW settings. The membrane was soaked in 5% milk suspended in Tris buffered saline with tween (TBST) overnight at 4 °C with gentle shaking. The next day, the membrane was washed three times for five minutes in TBST. The membrane was then incubated in 5% milk-TBST with a 1:5000 dilution of a mouse anti-LacI antibody (clone 9A5-05-503-I, Fisher) for 1 hour at room temperature with gentle shaking. The membrane was then washed again 3 times with TBST. We then incubated the membrane for one hour at room temperature in 5% milk-TBST with a 1:5000 peroxidase-conjugated anti-mouse antibody and washed three times with TBST. The blot was imaged by chemiluminescence using the Pierce™ ECL Western Blotting Substrate (ThermoFisher).

---

**Extended Data Figure 1.**
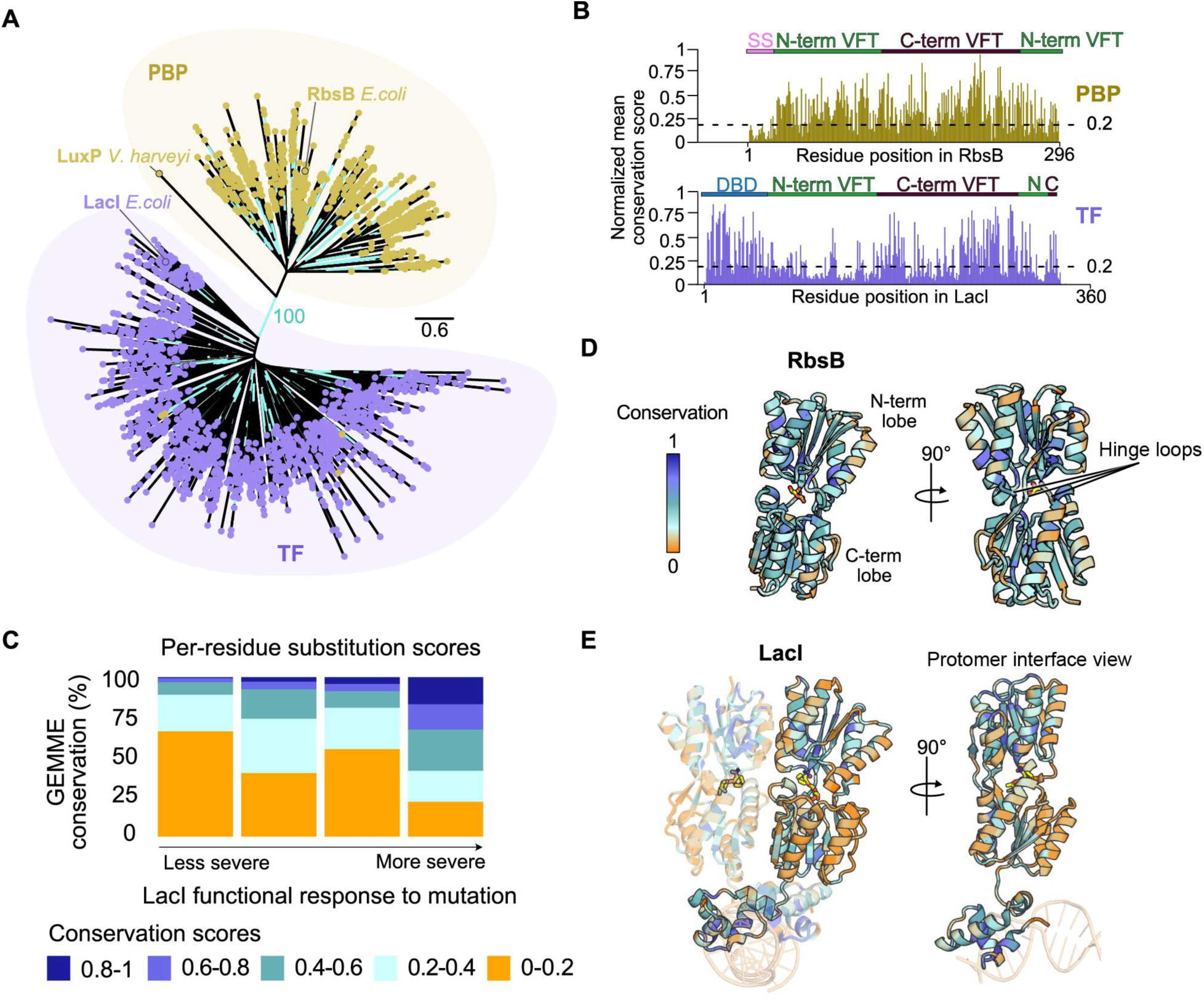
Sequence conservation in LacI/GalR TF and PBP families. **(A)** Reconstructed gene tree shows distinct clades of LacI/GalR TFs versus PBPs. Bootstrap values >90% are in turquoise. **(B)** Sequence conservation is higher and more uniform for PBPs than for TFs. **(C)** TF sequence conservation echoes mutational data for LacI: more conserved positions are less tolerant to mutations, causing severe changes to LacI function. **(D)** PBP sequence conservation mapped into the structure of RbsB. **(E)** The TF dimer interface is among the least conserved regions of the fold. LacI shown here.

**Extended Data Figure 2.**
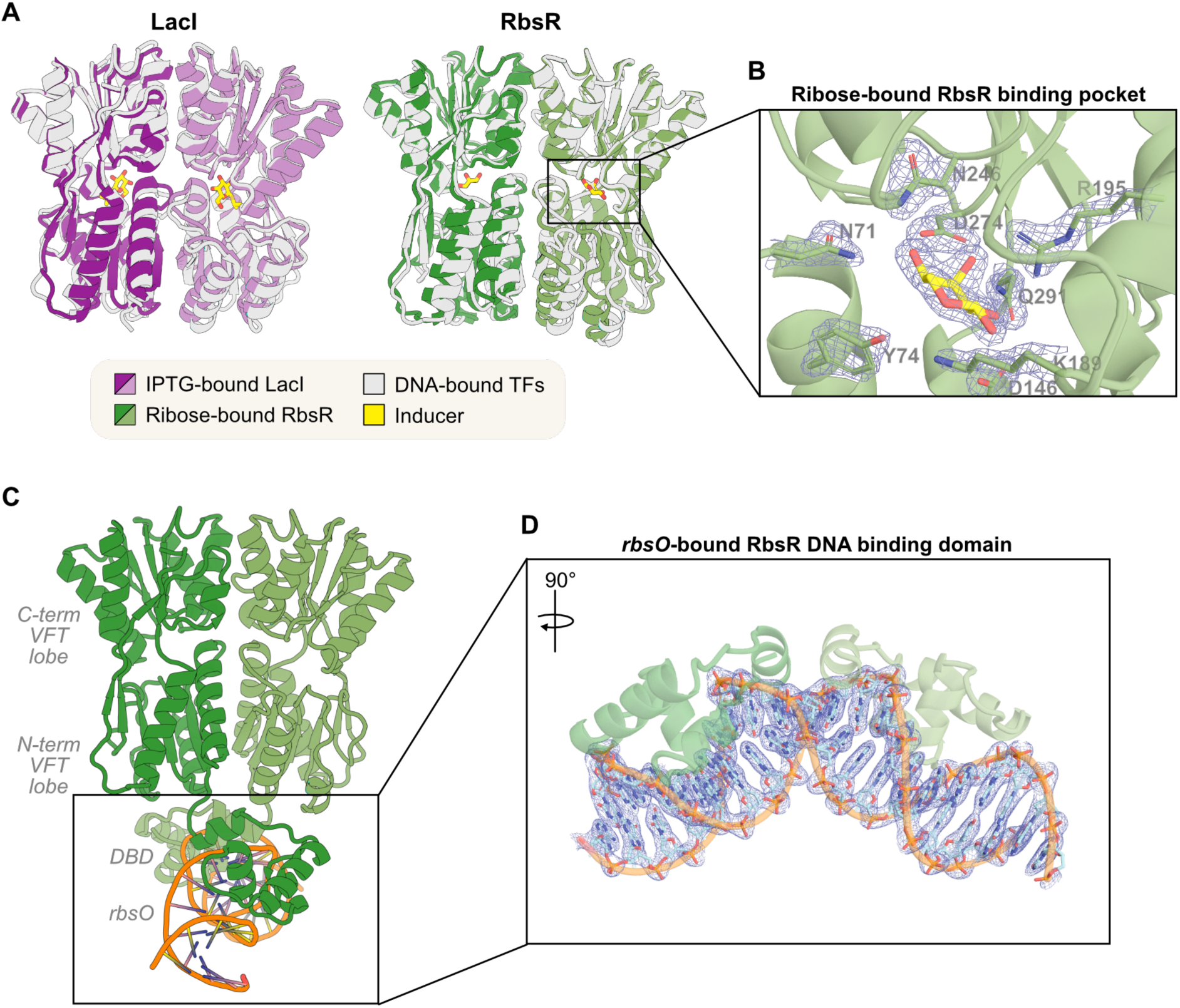
Structural conservation in LacI/GalR TFs. **(A)** We solved 2.1 Å X-ray crystal structures of RbsR complexed with ribose and its operator DNA *rbsO*. The inducer-bound states of LacI and RbsR VFT domains are structurally similar by X-ray crystallography. **(B)** Zoomed view of key RbsR interactions with ribose. **(C)** RbsR-*rbsO* X-ray crystal structure. **(D)** Zoomed view of *rbsO* interactions with the RbsR DBD. The 2Fo–Fc maps for sidechains, DNA, and ribose are shown at 1σ.

**Extended Data Figure 3.**
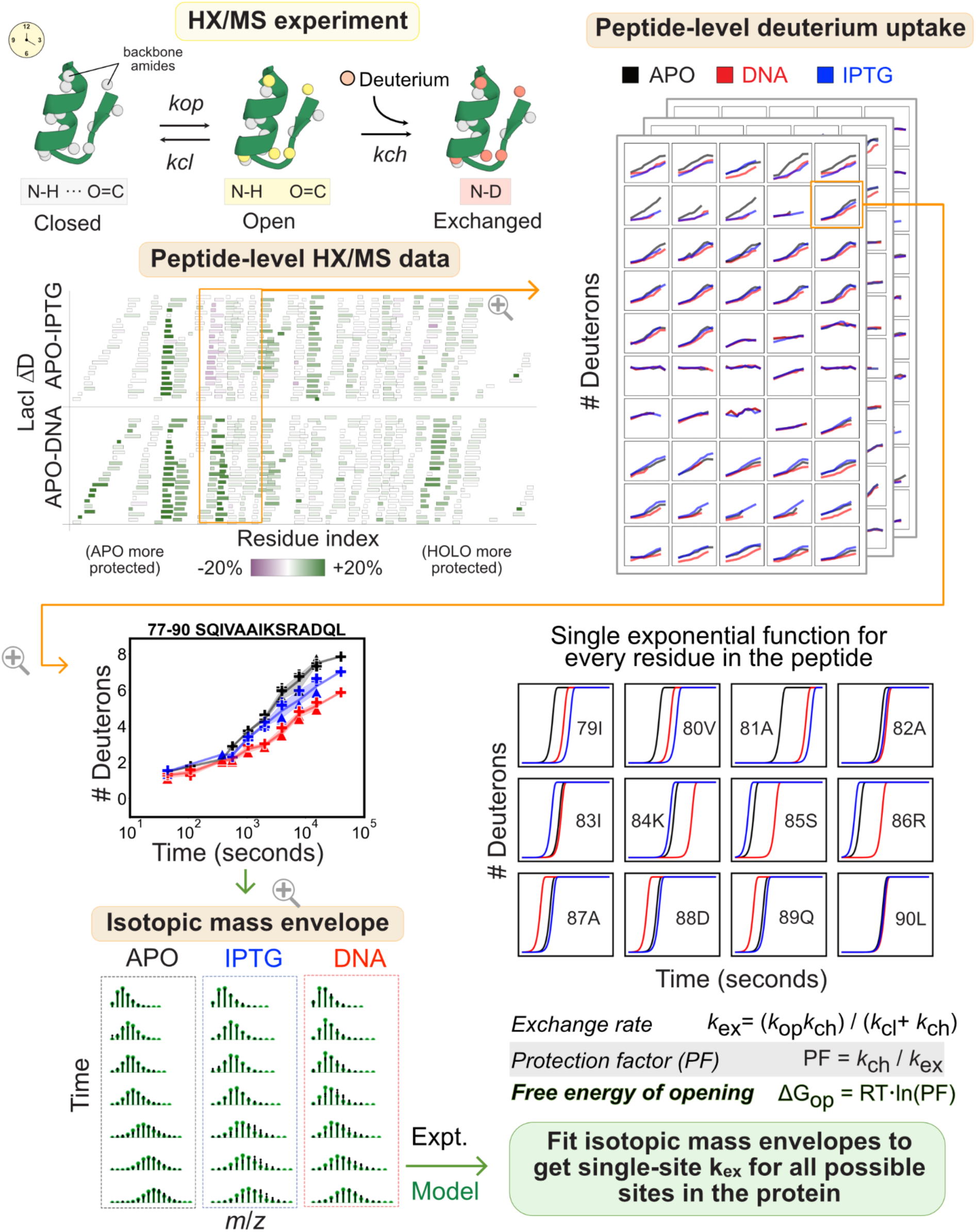
HX/MS strategy overview. HX/MS experiment and analysis using PIGEON-FEATHER, resulting in site-resolved ΔG_op_ for most residues.

**Extended Data Figure 4.**
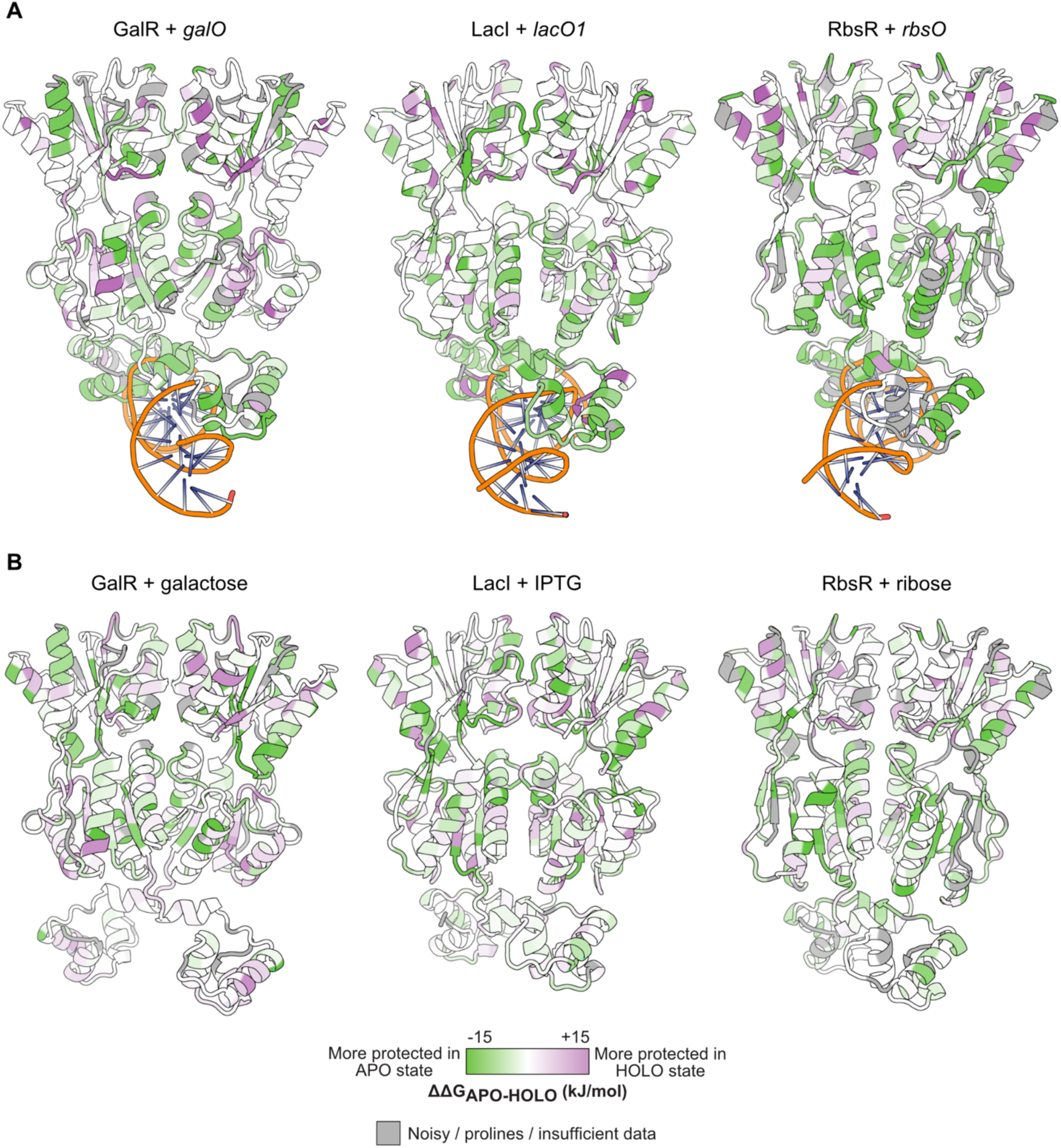
Ligand response in each TF ensemble. Each TF responds to binding **(A)** its own operator DNA and **(B)** its inducer molecule with global changes in local stability.

**Extended Data Figure 5.**
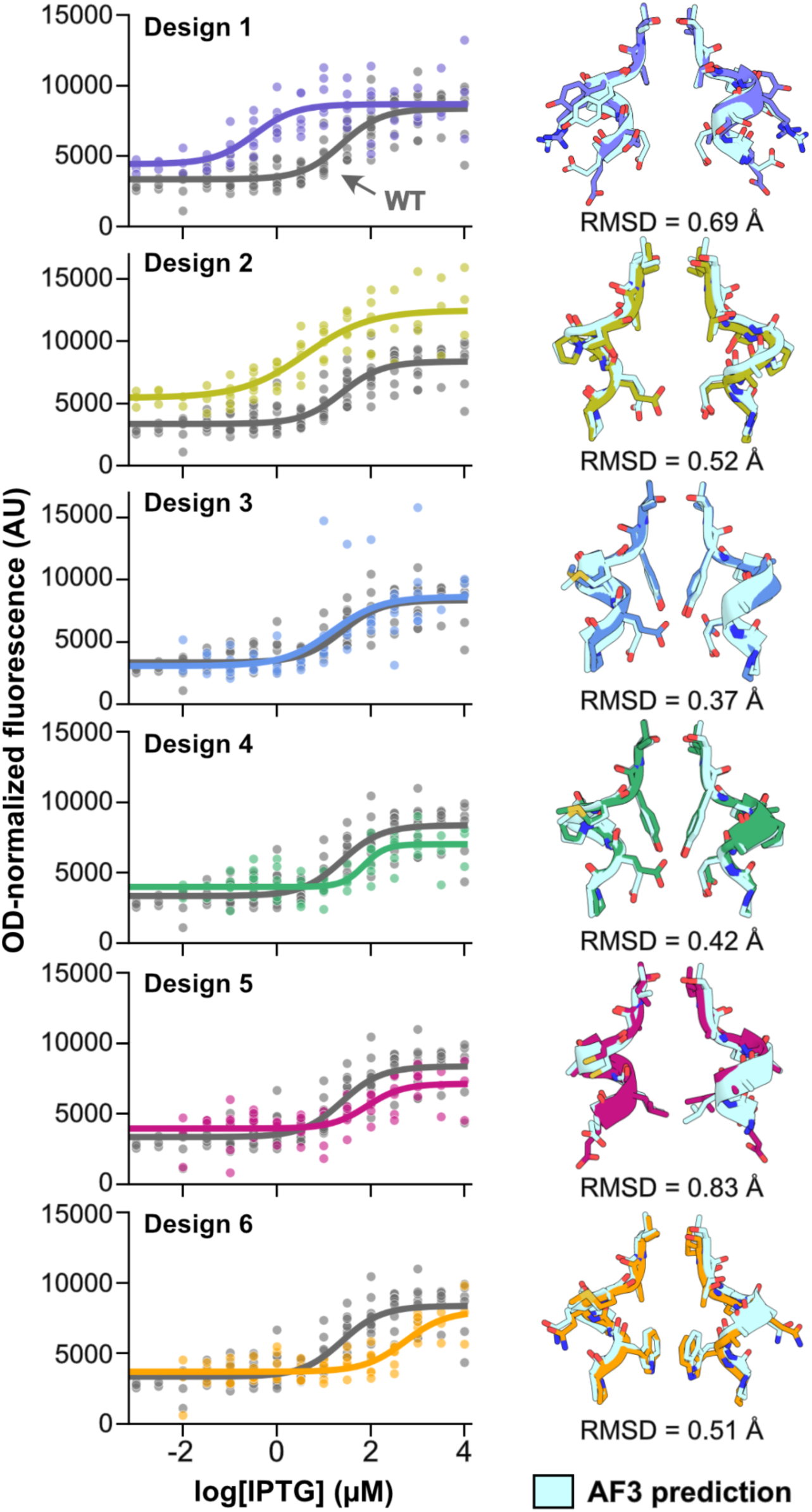
Designed LacI variant IPTG responses. *Left:* IPTG titrations for each variant. *Right:* AF3 structure predictions have <1 Å all-atom RMSD from design models.

## References

1. Lin, Z. et al. Evolutionary-scale prediction of atomic-level protein structure with a language model. Science 379, 1123–1130 (2023).

2. Chothia, C. One thousand families for the molecular biologist. Nature 357, 543–544 (1992).

3. Li, H., Tang, C. & Wingreen, N. S. Are protein folds atypical? Proceedings of the National Academy of Sciences 95, 4987–4990 (1998).

4. Cossio, P. et al. Exploring the Universe of Protein Structures beyond the Protein Data Bank. PLOS Computational Biology 6, e1000957 (2010).

5. Chitturi, B., Shi, S., Kinch, L. N. & Grishin, N. V. Compact Structure Patterns in Proteins. Journal of Molecular Biology 428, 4392–4412 (2016).

6. Minami, S. et al. Exploration of novel αβ-protein folds through de novo design. Nat Struct Mol Biol 30, 1132–1140 (2023).

7. Fukami-Kobayashi, K., Tateno, Y. & Nishikawa, K. Parallel evolution of ligand specificity between LacI/GalR family repressors and periplasmic sugar-binding proteins. Mol Biol Evol 20, 267–277 (2003).

8. Kaczmarski, J. A. et al. Altered conformational sampling along an evolutionary trajectory changes the catalytic activity of an enzyme. Nat Commun 11, 5945 (2020).

9. Torgeson, K. R. et al. Conserved conformational dynamics determine enzyme activity. Science Advances 8, eabo5546 (2022).

10. Cao, J. et al. Evolution of the class C GPCR Venus flytrap modules involved positive selected functional divergence. BMC Evolutionary Biology 9, 67 (2009).

11. Swint-Kruse, L. & Matthews, K. S. Allostery in the LacI/GalR family: variations on a theme. Current Opinion in Microbiology 12, 129–137 (2009).

12. Shilton, B. H., Flocco, M. M., Nilsson, M. & Mowbray, S. L. Conformational Changes of Three Periplasmic Receptors for Bacterial Chemotaxis and Transport: The Maltose-, Glucose/Galactose- and Ribose-binding Proteins. Journal of Molecular Biology 264, 350–363 (1996).

13. Mowbray, S. L. & Björkman, A. J. Conformational changes of ribose-binding protein and two related repressors are tailored to fit the functional need. J Mol Biol 294, 487–499 (1999).

14. Parente, D. J. & Swint-Kruse, L. Multiple Co-Evolutionary Networks Are Supported by the Common Tertiary Scaffold of the LacI/GalR Proteins. PLOS ONE 8, e84398 (2013).

15. Quiocho, F. A. & Ledvina, P. S. Atomic structure and specificity of bacterial periplasmic receptors for active transport and chemotaxis: variation of common themes. Molecular Microbiology 20, 17–25 (1996).

16. Binnie, R. A., Zhang, H., Mowbray, S. & Hermodson, M. A. Functional mapping of the surface of escherichia coli ribose-binding protein: Mutations that affect chemotaxis and transport. Protein Science 1, 1642–1651 (1992).

17. Kondoh, H., Ball, C. B. & Adler, J. Identification of a methyl-accepting chemotaxis protein for the ribose and galactose chemoreceptors of Escherichia coli. Proc. Natl. Acad. Sci. U.S.A. 76, 260–264 (1979).

18. Daber, R., Stayrook, S., Rosenberg, A. & Lewis, M. Structural Analysis of Lac Repressor Bound to Allosteric Effectors. Journal of Molecular Biology 370, 609–619 (2007).

19. Glasgow, A. et al. Ligand-specific changes in conformational flexibility mediate long-range allostery in the lac repressor. Nat Commun 14, 1179 (2023).

20. Lu, C., Wells, M.L., Reckers, A. et al. Site-resolved energetic information from HX–MS experiments. Nat Chem Biol (2025). 10.1038/s41589-025-02049-1

21. Markiewicz, P., Kleina, L. G., Cruz, C., Ehret, S. & Miller, J. H. Genetic studies of the lac repressor. XIV. Analysis of 4000 altered Escherichia coli lac repressors reveals essential and non-essential residues, as well as ‘spacers’ which do not require a specific sequence. J. Mol. Biol. 240, 421–433 (1994).

22. Hars, U., Horlacher, R., Boos, W., Welte, W. & Diederichs, K. Crystal structure of the effector-binding domain of the trehalose-repressor of escherichia coli, a member of the LacI family, in its complexes with inducer trehalose-6-phosphate and noninducer trehalose. Protein Science 7, 2511–2521 (1998).

23. Schumacher, M. A., Glasfeld, A., Zalkin, H. & Brennan, R. G. The X-ray Structure of the PurR-Guanine-purF Operator Complex Reveals the Contributions of Complementary Electrostatic Surfaces and a Water-mediated Hydrogen Bond to Corepressor Specificity and Binding Affinity *. Journal of Biological Chemistry 272, 22648–22653 (1997).

24. Schumacher, M. A. et al. Structural basis for allosteric control of the transcription regulator CcpA by the phosphoprotein HPr-Ser46-P. Cell 118, 731–741 (2004).

25. Swint-Kruse, L., Elam, C. R., Lin, J. W., Wycuff, D. R. & Matthews, K. S. Plasticity of quaternary structure: Twenty-two ways to form a LacI dimer. Protein Science 10, 262–276 (2001).

26. Oldham, M. L., Khare, D., Quiocho, F. A., Davidson, A. L. & Chen, J. Crystal structure of a catalytic intermediate of the maltose transporter. Nature 450, 515–521 (2007).

27. Thomas, C. & Tampé, R. Structural and Mechanistic Principles of ABC Transporters. Annu Rev Biochem 89, 605–636 (2020).

28. Friedman, A. M., Fischmann, T. O. & Steitz, T. A. Crystal structure of lac repressor core tetramer and its implications for DNA looping. Science 268, 1721–1727 (1995).

29. Lewis, M. et al. Crystal Structure of the Lactose Operon Repressor and Its Complexes with DNA and Inducer. Science 271, 1247–1254 (1996).

30. Bell, C. E. & Lewis, M. A closer view of the conformation of the Lac repressor bound to operator. Nat Struct Biol 7, 209–214 (2000).

31. Schumacher, M. A., Choi, K. Y., Lu, F., Zalkin, H. & Brennan, R. G. Mechanism of corepressor-mediated specific DNA binding by the purine repressor. Cell 83, 147–155 (1995).

32. Mauzy, C. a. & Hermodson, M. a. Structural and functional analyses of the repressor, RbsR, of the ribose operon of Escherichia coli. Protein Science 1, 831–842 (1992).

33. Hilser, V. J. An Ensemble View of Allostery. Science 327, 653–654 (2010).

34. von Wilcken-Bergmann, B. & Müller-Hill, B. Sequence of galR gene indicates a common evolutionary origin of lac and gal repressor in Escherichia coli. Proceedings of the National Academy of Sciences 79, 2427–2431 (1982).

35. Majumdar, A. & Adhya, S. Probing the structure of gal operator-repressor complexes. Conformation change in DNA. Journal of Biological Chemistry 262, 13258–13262 (1987).

36. Bai, Y., Sosnick, T. R., Mayne, L. & Englander, S. W. Protein Folding Intermediates: Native-State Hydrogen Exchange. Science 269, 192–197 (1995).

37. Kalodimos, C. G. et al. Structure and Flexibility Adaptation in Nonspecific and Specific Protein-DNA Complexes. Science 305, 386–389 (2004).

38. Spronk, C. A. E. M., Slijper, M., van Boom, J. H., Kaptein, R. & Boelens, R. Formation of the hinge helix in the lac represser is induced upon binding to the lac operator. Nat Struct Mol Biol 3, 916–919 (1996).

39. Chen, J. & Matthews, K. S. Subunit dissociation affects DNA binding in a dimeric lac repressor produced by C-terminal deletion. Biochemistry 33, 8728–8735 (1994).

40. Abramson, J. et al. Accurate structure prediction of biomolecular interactions with AlphaFold 3. Nature 630, 493–500 (2024).

41. Schmitz, A., Schmeissner, U. & Miller, J. H. Mutations affecting the quaternary structure of the lac repressor. Journal of Biological Chemistry 251, 3359–3366 (1976).

42. Suckow, J. et al. Genetic Studies of the Lac Repressor XV: 4000 Single Amino Acid Substitutions and Analysis of the Resulting Phenotypes on the Basis of the Protein Structure. Journal of Molecular Biology 261, 509–523 (1996).

43. Spronk, C. A. et al. The solution structure of Lac repressor headpiece 62 complexed to a symmetrical lac operator. Structure 7, 1483–1492 (1999).

44. Lin, S. & Riggs, A. D. lac represser binding to non-operator DNA: Detailed studies and a comparison of equilibrium and rate competition methods. Journal of Molecular Biology 72, 671–690 (1972).

45. Makabe, K., Yan, S., Tereshko, V., Gawlak, G. & Koide, S. β-Strand Flipping and Slipping Triggered by Turn Replacement Reveal the Opportunistic Nature of β-Strand Pairing. J. Am. Chem. Soc. 129, 14661–14669 (2007).

46. Lalwani, M. A. et al. Optogenetic control of the lac operon for bacterial chemical and protein production. Nat Chem Biol 17, 71–79 (2021).

47. Dauparas, J. et al. Robust deep learning–based protein sequence design using ProteinMPNN. Science 378, 49–56 (2022).

48. Alford, R. F. et al. The Rosetta All-Atom Energy Function for Macromolecular Modeling and Design. J. Chem. Theory Comput. 13, 3031–3048 (2017).

49. Taylor, N. D. et al. Engineering an allosteric transcription factor to respond to new ligands. Nat Meth 13, 177–183 (2016).

50. Tack, D. S. et al. The genotype-phenotype landscape of an allosteric protein. Mol Syst Biol 17, e10179 (2021).

51. Tungtur, S., Egan, S. M. & Swint-Kruse, L. Functional consequences of exchanging domains between LacI and PurR are mediated by the intervening linker sequence. Proteins: Structure, Function, and Bioinformatics 68, 375–388 (2007).

52. Hersey, A. N., Kay, V. E., Lee, S., Realff, M. J. & Wilson, C. J. Engineering allosteric transcription factors guided by the LacI topology. cels 14, 645–655 (2023).

53. Garruss, A. S., Collins, K. M. & Church, G. M. Deep representation learning improves prediction of LacI-mediated transcriptional repression. Proceedings of the National Academy of Sciences 118, e2022838118 (2021).

54. Pavlovicz, R. E., Park, H. & DiMaio, F. Efficient consideration of coordinated water molecules improves computational protein-protein and protein-ligand docking discrimination. PLOS Computational Biology 16, e1008103 (2020).

55. Horlacher, R. & Boos, W. Characterization of TreR, the major regulator of the Escherichia coli trehalose system. J Biol Chem 272, 13026–13032 (1997).

56. Falcon, C. M. & Matthews, K. S. Glycine Insertion in the Hinge Region of Lactose Repressor Protein Alters DNA Binding*. Journal of Biological Chemistry 274, 30849–30857 (1999).

57. Kleina, L. G. & Miller, J. H. Genetic studies of the *lac* repressor. Journal of Molecular Biology 212, 295–318 (1990).

58. Campitelli, P., Swint-Kruse, L. & Ozkan, S. B. Substitutions at Nonconserved Rheostat Positions Modulate Function by Rewiring Long-Range, Dynamic Interactions. Molecular Biology and Evolution 38, 201–214 (2021).

59. Kröger, P., Shanmugaratnam, S., Ferruz, N., Schweimer, K. & Höcker, B. A comprehensive binding study illustrates ligand recognition in the periplasmic binding protein PotF. Structure 29, 433–443.e4 (2021).

60. Dwyer, M. A. & Hellinga, H. W. Periplasmic binding proteins: a versatile superfamily for protein engineering. Current Opinion in Structural Biology 14, 495–504 (2004).

61. Richards, D. H., Meyer, S. & Wilson, C. J. Fourteen Ways to Reroute Cooperative Communication in the Lactose Repressor: Engineering Regulatory Proteins with Alternate Repressive Functions. ACS Synth Biol 6, 6–12 (2017).

62. Müller, J., Barker, A., Oehler, S. & Müller-Hill, B. Dimeric lac repressors exhibit phase-dependent co-operativity1. Journal of Molecular Biology 284, 851–857 (1998).

63. Meinhardt, S. et al. Novel insights from hybrid LacI/GalR proteins: family-wide functional attributes and biologically significant variation in transcription repression. Nucleic Acids Research 40, 11139–11154 (2012).

64. Glasscock, C.J. et al. Computational design of sequence-specific DNA-binding proteins. Nat Struct Mol Biol 32, 2252–2261 (2025). 10.1038/s41594-025-01669-4

65. Perez-Riverol, Y. et al. The PRIDE database resources in 2022: a hub for mass spectrometry-based proteomics evidences. Nucleic Acids Research 50, D543–D552 (2022).

66. Steinegger, M. & Söding, J. MMseqs2 enables sensitive protein sequence searching for the analysis of massive data sets. Nat Biotechnol 35, 1026–1028 (2017).

67. Laine, E., Karami, Y. & Carbone, A. GEMME: A Simple and Fast Global Epistatic Model Predicting Mutational Effects. Molecular Biology and Evolution 36, 2604–2619 (2019).

68. Minh, B. Q. et al. IQ-TREE 2: New Models and Efficient Methods for Phylogenetic Inference in the Genomic Era. Mol Biol Evol 37, 1530–1534 (2020).

69. Minh, B. Q., Dang, C. C., Vinh, L. S. & Lanfear, R. QMaker: Fast and Accurate Method to Estimate Empirical Models of Protein Evolution. Systematic Biology 70, 1046–1060 (2021).

70. Schwarz, G. Estimating the Dimension of a Model. The Annals of Statistics 6, 461–464 (1978).

71. Berman, H. M. et al. The Protein Data Bank. Nucleic Acids Research 28, 235–242 (2000).

72. Zhang, C., Shine, M., Pyle, A. M. & Zhang, Y. US-align: universal structure alignments of proteins, nucleic acids, and macromolecular complexes. Nat Methods 19, 1109–1115 (2022).

73. Lu, C., Wells, M. L., Reckers, A., McBride, S. K. & Glasgow, A. Site-resolved energetic information from HX–MS experiments. Nat Chem Biol 1–11 (2025) doi:10.1038/s41589-025-02049-1.

74. Winter, G. xia2: an expert system for macromolecular crystallography data reduction. Applied Crystallography 43, 186–190 (2010).

75. Winter, G. et al. DIALS: implementation and evaluation of a new integration package. Acta Cryst D 74, 85–97 (2018).

76. Agirre, J. et al. The CCP4 suite: integrative software for macromolecular crystallography. Acta Cryst D 79, 449–461 (2023).

77. Kabsch, W. xds. Biological crystallography 66, 125–132 (2010).

78. STARANISO anisotropy & Bayesian estimation server. https://staraniso.globalphasing.org/cgi-bin/staraniso.cgi.

79. McCoy, A. J. et al. Phaser crystallographic software. Applied Crystallography 40, 658–674 (2007).

80. Jumper, J. et al. Highly accurate protein structure prediction with AlphaFold. Nature 596, 583–589 (2021).

81. Emsley, P. & Cowtan, K. Coot: model-building tools for molecular graphics. Biological crystallography 60, 2126–2132 (2004).

82. Afonine, P. V. et al. Towards automated crystallographic structure refinement with phenix. refine. Biological crystallography 68, 352–367 (2012).

83. Skubák, P. et al. REFMAC5 for the refinement of macromolecular crystal structures. Acta Crystallographica. Section D: Biological Crystallography 67, (2011).

84. Joosten, R. P., Long, F., Murshudov, G. N. & Perrakis, A. The PDB_REDO server for macromolecular structure model optimization. IUCrJ 1, 213–220 (2014).

85. Agerschou, E. D. et al. The transcriptional regulator GalR self-assembles to form highly regular tubular structures. Sci Rep 6, 27672 (2016).

86. Hsieh, M. & Brenowitz, M. Comparison of the DNA Association Kinetics of the Lac Repressor Tetramer, Its Dimeric Mutant LacI adi, and the Native Dimeric Gal Repressor *. Journal of Biological Chemistry 272, 22092–22096 (1997).

87. Chatterjee, S., Zhou, Y.-N., Roy, S. & Adhya, S. Interaction of Gal repressor with inducer and operator: Induction of gal transcription from repressor-bound DNA. Proc Natl Acad Sci U S A 94, 2957–2962 (1997).

88. The PyMOL Molecular Graphics System, Version 2.0 Schrödinger, LLC.

89. Dauparas, J. et al. Robust deep learning–based protein sequence design using ProteinMPNN. Science 378, 49–56 (2022).

90. Alford, R. F. et al. The Rosetta All-Atom Energy Function for Macromolecular Modeling and Design. J. Chem. Theory Comput. 13, 3031–3048 (2017).

91. Brooks, B. R. et al. CHARMM: A program for macromolecular energy, minimization, and dynamics calculations. Journal of Computational Chemistry 4, 187–217 (1983).

92. Vanommeslaeghe, K. et al. CHARMM General Force Field (CGenFF): A force field for drug-like molecules compatible with the CHARMM all-atom additive biological force fields. J Comput Chem 31, 671–690 (2010).

93. Michaud-Agrawal, N., Denning, E. J., Woolf, T. B. & Beckstein, O. MDAnalysis: A toolkit for the analysis of molecular dynamics simulations. Journal of Computational Chemistry 32, 2319–2327 (2011).

94. Humphrey, W., Dalke, A. & Schulten, K. VMD: Visual molecular dynamics. Journal of Molecular Graphics 14, 33–38 (1996).

95. McGibbon, R. T. et al. MDTraj: A Modern Open Library for the Analysis of Molecular Dynamics Trajectories. Biophysical Journal 109, 1528–1532 (2015).

96. Pedregosa, F. et al. Scikit-learn: Machine Learning in Python. Journal of Machine Learning Research 12, 2825–2830 (2011).

