## Supplemental information for "Distinct energetic blueprints diversify function of conserved protein folds"

---

**Additional supplementary materials available with this study:**

- HX/MS deuterium uptake plots
- FEATHER-derived centroid fits to HX/MS data
- VFT_ddGs.pse: a PyMOL session file for exploring VFT protein $\Delta\Delta G_{op}$
- Sequence alignment file for five proteins

**Bioinformatics and modeling data uploaded to Zenodo (dataset identifier 10.5281/zenodo.15091462):**

- Conservation score matrices
- Gene trees
- MSA files
- AlphaFold 3-RosettaECO models
- MD trajectories

**X-ray crystal structures deposited to RCSB PDB:**

- RbsR-ribose (PDB ID 9NY7)
- RbsR-*rbsO* (PDB ID 9NY8)

**Raw HX/MS data available in PRIDE database (dataset identifier PXD062370)**

Table S2. X-ray crystallography summary statistics ... 38
Table S3. HX/MS replicate information for LacI ... 39
Table S4. HX/MS replicate information for GalR ... 40
Table S5. HX/MS replicate information for RbsR ... 41
Table S6. HX/MS replicate information for MglB ... 42
Table S7. HX/MS replicate information for RbsB ... 43
Table S8. $\Delta G_{op}$ values for LacI ... 44
Table S9. $\Delta G_{op}$ values for GalR ... 56
Table S10. $\Delta G_{op}$ values for RbsR ... 67
Table S11. $\Delta G_{op}$ values for MglB ... 78
Table S12. $\Delta G_{op}$ values for RbsB ... 89
Table S13. Rosetta metrics for LacI designs ... 99
Table S14. Hill equation constants for LacI designs ... 100
**Appendix 1. DNA operator sequences used in this study** ... 101
**Appendix 2. DNA sequences for TF and PBP genes** ... 102
**References** ... 104

#### Supplemental note 1: interpreting high-resolution HX/MS data

This work presents the first high-resolution HX/MS datasets analyzed at a multi-protein scale. Here we provide a brief overview about how the data were collected and analyzed, noting some key limitations to help the reader interpret the data responsibly.

**Data quality and completeness.** For all proteins except for GalR (for which we have triplicate data), we performed at least two technical replicates (experiments on different days) for each of at least two biological replicates (different protein preparations) to produce quadruplicate datasets. To maximize the number of unique peptides and the depth of the shared peptides, we used different proteases for each replicate. We pooled the data for each protein for PIGEON-FEATHER analysis.[1] This led to distinct peptides with differing sequence coverage for each protein (Tables S3-S7). We manually checked the quality of the PIGEON-FEATHER fits to the mass spectra and to the centroid-level data. Although this approach revealed the single-site exchange rates for the majority of protons in each protein, some dipeptides and tripeptides remained. For these, we calculated a set of rates for the set of protons but did not assign the rates. The rates were instead averaged and converted to $\Delta G_{op}$ values, and the average $\Delta G_{op}$ were painted on the protein structures in the figures alongside the site-resolved $\Delta G_{op}$. The full list of site-resolved $\Delta G_{op}$ are available in Tables S8-S12, with the PIGEON-FEATHER-derived bootstrapping standard deviations. More details about experimental data collection and HX/MS analysis are included in the **Methods** section.

**Back exchange corrections.** We assume that the main contribution to $\Delta G_{op}$ is the backbone amide exchange for each non-proline amino acid in each protein during the HX/MS experiment[2], and we make back-exchange corrections to account for the loss of deuterium that occurs during quenching, chromatography, and electrospray ionization. While these corrections allow the calculation of true deuterium incorporation, they introduce additional noise and variability. The correction relies on a measured back-exchange rate from fully deuterated control samples.

**Positions with very low and high $\Delta G_{op}$ values.** Benchmarking shows that $\Delta G_{op}$ ≥55 kJ/mol, the highest exchange rates corresponding to the protons most protected from hydrogen exchange in the conditions of the experiment, are less reliable than smaller $\Delta G_{op}$ because of the timescale limitations in the experimental data collection.[1] We graphed the region of precise -log($k_{ex}$) determination afforded by the timecourse for the FEATHER results in Figs. S11 and S12 for the five proteins in this study. We suggest interpreting the calculated $\Delta G_{op}$ for slower- and faster-exchanging protons, marked with a red dot, with more caution and more qualitatively than the other protons. It is reasonable to classify them as slow-/fast-exchanging or high-/low-$\Delta G_{op}$ protons.

**Cooperativity in oligomeric protein complexes.** PIGEON-FEATHER calculates an ensemble average $\Delta G_{op}$, so we cannot infer the behavior of individual protomers in oligomeric complexes.

#### Supplemental note 2: TF molecular switches

In Fig. 6, we propose a mechanism for the transition between the operator DNA-bound state and the inducer-bound state for the LacI/GalR TFs included in this study. We describe a change in the TF conformational ensemble such that all the effects described can be presumed to occur simultaneously, not in any particular order. The molecular switches are numbered according to the cartoon model in Fig. 6. Here we include the molecular details for each TF.

**(1) Stabilization of N-terminal dimer interface helix by inducer binding via structural water**
The N-terminal end of the transducer helix H3 is stabilized via inducer-water-H3 interactions.

**LacI:** A75 is stabilized in the first turn of H3 via a water bridged hydrogen bond between its backbone amide group, IPTG (with which it also makes hydrophobic interactions), and H28 N246, which also directly hydrogen bonds with IPTG. IPTG also makes direct hydrogen bonds with D274 and Q291, which are also stabilized.

**GalR:** H3 F73 and F74 are stabilized in the first turn of H3 via packing interactions with the sugar. Galactose also interacts directly with D71, F73, N245, D273, R194, and D190, which are also stabilized.

**RbsR:** C2 N71 makes water-bridged hydrogen bonds with ribose and the backbone amide of H28 A248. Ribose makes hydrophobic interactions with nearby H3 F73. All three residues are stabilized by inducer binding. Ribose makes several additional hydrogen bonds in the pocket with other stabilized residues: N71, Y74, K189, D274. It also packs against F220.

**(2) Stabilization of cross-VFT-lobe H20-loop interactions**
A sidechain interaction coupling the N-terminal VFT lobe central beta sheet and attached loops to one position in H20 is conserved, as is stabilization by inducer binding at both sites. Modification of sidechain interactions between functional states is also seen in crystal structures.

**LacI:** H20 S193 hydrogen bonds with C14 D149. Inducer binding strengthens this interaction to stabilize both residues.

**GalR:** The H20 D190 interaction with E9 K124 is stabilized by inducer binding. The role of strand E9 in GalR is unique; in LacI and RbsR, this role is filled by C14. K124 also hydrogen bonds directly with galactose. Other local residues in both H20 and E9 are also strongly stabilized, most notably all residues in C19.

**RbsR:** H20 P191 and C14 W147 make a stacking interaction in the operator-bound state that is abolished on inducer binding. W147 is stabilized by inducer binding.

**(3) Allosteric destabilization of C-terminal dimer interface helix**
DNA binding stabilizes H32, the metastable helix, in all three TFs, but inducer binding strongly destabilizes it. Despite the broad destabilization, the same interaction at the base of the

metastable helix is stabilized across the dimer interface in the inducer-bound state compared to the operator-bound state in each TF. Changes in the monomer-dimer equilibrium caused by unbinding DNA contribute to these ensemble effects.

**LacI:** H32 residues 276, 278, 279, and 281-283 are destabilized. E277 and A222 at the base of the helix are stabilized in a sidechain-backbone hydrogen bond across the interface in the IPTG state compared to the *lacO1*-bound state.

**GalR:** H32 residues 279-282 are destabilized. L276 at the base of the helix is stabilized by a hydrophobic interaction with L276 on the other protomer in the galactose-bound state compared to the *galO*-bound state.

**RbsR:** H32 residues 281-283 are destabilized. E277 and F222 at the base of the helix are stabilized through a hydrophobic interaction across the interface in the ribose-bound state compared to the *rbsO*-bound state.

**(4) Stabilization of helix-helix interactions in the N-terminal VFT lobe**

Inducer binding increases the stability of the N-terminal region of H3, the transducer helix. Compared to the operator-bound state, the C-terminal region of H3 is destabilized in each TF, perhaps as a direct result of decreased DNA affinity. The central residues in the neighboring helix H36 are stabilized by upweighting H3-H36 interhelical interactions and hydrophobic interactions with the outer N-terminal beta sheet strands, E13 and E58.

**LacI:** Hydrophobic interactions between H3 A82 and I79 and H36 Q298 are stabilized, as are contiguous residues in H36. Compared to the *lacO1*-bound state, S300 and D302 are stabilized in the IPTG-bound state.

**GalR:** Galactose binding stabilizes the first three turns of H36. Compared to the *galO* state, H36 residues 298 and 301-303 are strongly stabilized in the galactose-bound state.

**RbsR:** The second turn of H36 is strongly stabilized in response to ribose binding, including the H32 G80-H3 G297 interaction. The contiguous residues 299-304 are stabilized in RbsR-ribose compared to RbsR-*rbsO.*

**(5) Restructuring of N-terminal VFT lobe beta sheet**

Operator DNA binding and inducer binding both increase cross-interface beta sheet interactions relative to the apo state, but inducer binding shifts the hydrogen bonding pattern in E5, E1, and E9 of each TF from its DNA state configuration, radiating outwards from the dimer interface.

**LacI:** Hydrogen bonds in the N-terminal VFT lobe beta sheet strengthened by IPTG binding in E1, E5, and E9: V66-G66; G65-G121, V95-V95’. C8 is strongly stabilized. These elements are closer to the dimer interface. Farther from the dimer interface, E13 and E15 are stabilized in the *lacO1* state compared to the IPTG state.

**GalR:** Compared to the *galO*-bound state, galactose binding strongly stabilizes E5 and weakly stabilizes E1 and E9, partially due to the hydrogen bonds between L64-L93 and across the dimer interface involving L92 and L93.

**RbsR:** Ribose binding and *rbsO* binding both strongly stabilize the central beta sheet strands E5, E1, and E9. Compared to the *rbsO* state, RbsR-ribose has stronger hydrogen bonds in E1 and E9, but not E5.

**(6) Helical destabilization and displacement at the VFT-DBD interface**
H7 is destabilized and moves visibly in X-ray crystal structures to pinch inwards towards the dimer interface when comparing inducer- and DNA-bound states, changing VFT domain interactions with the hinge helix.

**LacI:** In response to IPTG-driven DNA unbinding and reduced *lacO1* interactions, the entirety of H7 is destabilized in the IPTG state compared to LacI-*lacO1*. As a result, the interaction with the hinge helix is also strongly destabilized. The H7 A116 backbone hydrogen bond with hinge helix-adjacent loop residue R51 is broken. The H7 N113 sidechain hydrogen bond with R51 sidechain is also broken. These VFT domain disruptions to interactions with the hinge helices also disrupt inter-hinge-helix interactions, *e.g.*, V52-V52' interactions across the dimer interface.

**GalR:** H7 residues 109-111 are destabilized by galactose binding. In the hinge helix-loop region, the hydrogen bond between R52 and E60 is broken. Inter-hinge helix interactions, like the L54-L54' hydrophobic interaction across the dimer interface, are also disrupted.

**RbsR:** H7 is broadly destabilized, and interactions between H7, the hinge helix, and the hinge helix-adjacent loop are also weakened upon ribose binding. Examples: S53-Q58 both backbone and sidechain interactions locking the hinge helix in place are broken upon ribose binding, releasing the hinge helix. The H7 Q113-hinge helix A49 backbone hydrogen bond is weakened. H7 K114 makes a cross-dimer-interface hydrogen bond with E1 T61, which is also weakened.

**(7) VFT-DBD interface destabilization distal to the dimer interface**
Inducer binding destabilizes the C-terminal ends of H36 and H11 even in comparison to the apo state. In comparison to the operator-bound state, it is further destabilized, reducing hydrophobic interactions with the loop that joins the hinge helix to the VFT domain.

**LacI:** The hydrogen bond between H36 S308 and hinge helix-adjacent loop residue L62 is weakened. H11 is destabilized, and interactions with the DBD are broken, *e.g.*, T141-N46 hydrophobic contact.

**GalR:** H11 Q137 hydrogen bonds with DBD loop residues H46 and S44 are broken. The C-terminal ends of H11 and H36 are destabilized.

**RbsR:** H11 is broadly destabilized in the ribose-bound state. H36 is stabilized except for the C-terminal turn which is strongly destabilized (residues 307-309).

**(8) Destabilized DBD disrupts interactions with operator DNA**
The hinge helices are destabilized at a conserved central alanine, releasing them from the minor groove. Interactions between the recognition helix and the operator DNA are abolished. Nonspecific DNA interactions and intra-DBD helical packing interactions are weakened.

**LacI:** The hinge helix is destabilized at A53 upon IPTG binding, and there is decreased $\Delta G_{op}$ at recognition helix positions 14-19. This is accompanied by a reduction in nonspecific charged interactions with DNA, *e.g.*, S16; in addition, helical packing interactions are reduced, *e.g.*, A10/V15 interactions, Y12/L45 interactions.

**GalR and RbsR:** The hinge helix is destabilized at A51 by inducer binding in both TFs. There is decreased $\Delta G_{op}$ at recognition helix residues 14-19 and a reduction in nonspecific charged interactions with DNA, *e.g.*, by S14. Helical packing interactions are reduced, *e.g.*, A8/V13 interactions, L10/L43 interactions.

### Supplemental figures

##### Figure S1. Pairwise sequence identity in LacI/GalR TFs and PBPs

Comparison of pairwise sequence identities among **A**, 1,109 LacI/GalR TF sequences and **B**, 831 PBPs sequences in the MSA.

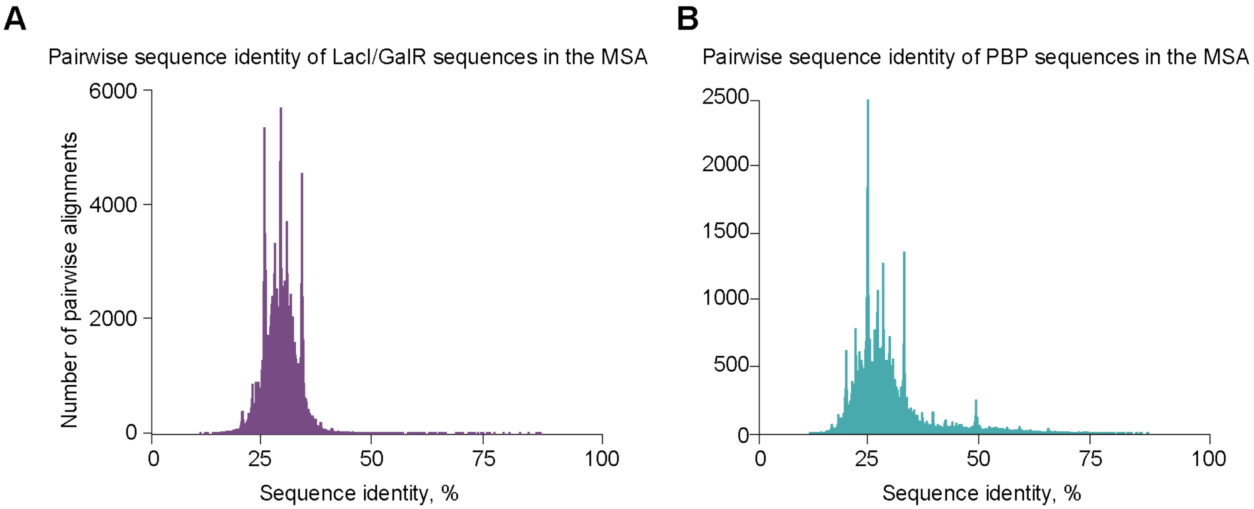

**Figure S2. Multiple sequence alignments (MSA) for LacI/GalR TFs and PBPs**

**A**, for LacI/GalR TFs (1,109 sequences)**, B**, for PBPs (831 sequences). The sequence logo was made using Weblogo[3], and the secondary structure assignment was made using SSDraw.[4]

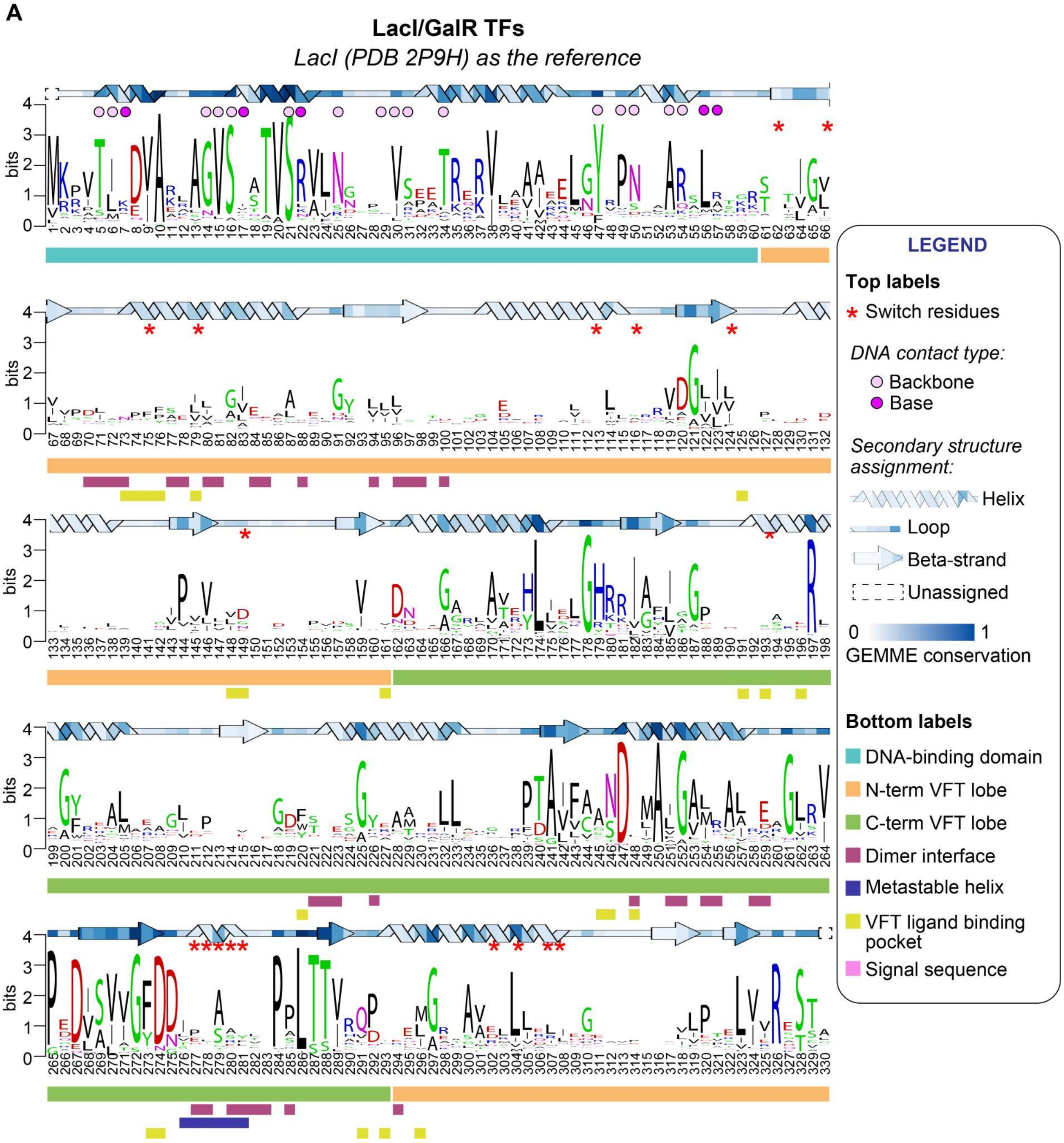

**B**

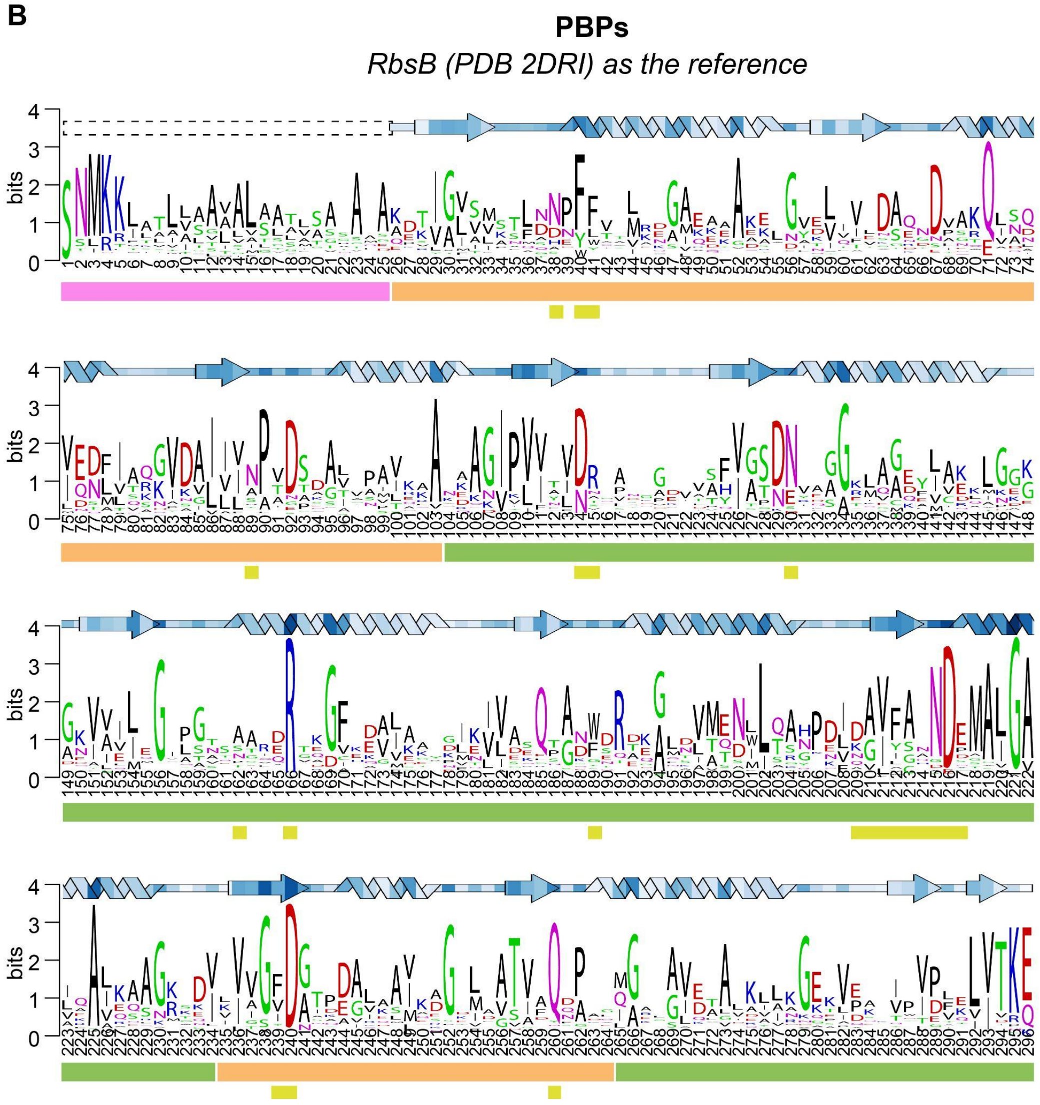

**Figure S3. Dimer interface comparison between three experimentally solved TFs**

Experimentally solved structures of three *E.coli* TFs in the ligand-bound form have varied dimer interfaces. Lowercase letters indicate conserved interacting residue positions.

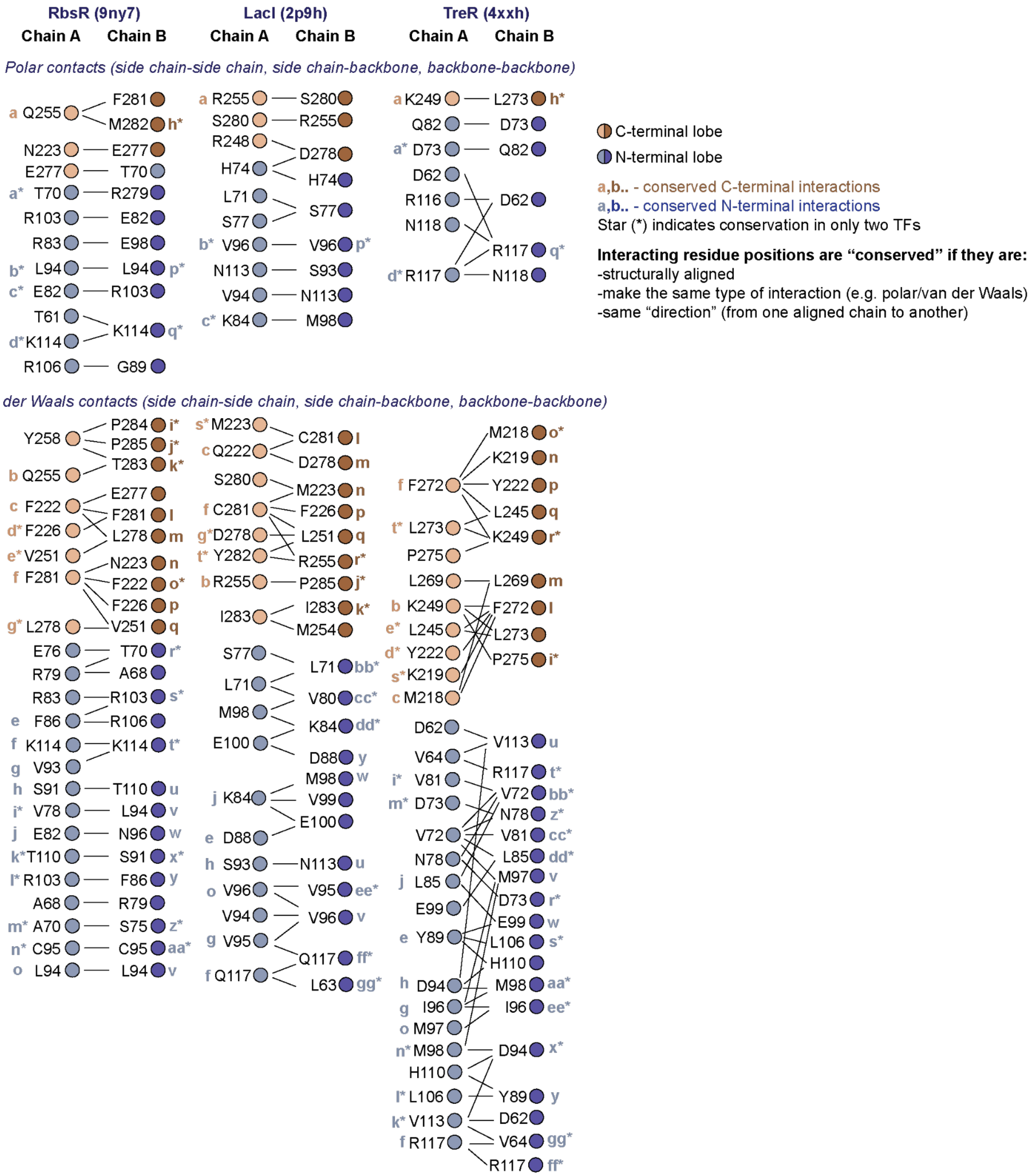

**Figure S4. X-ray crystal structures of LacI/GalR TFs are similar**

**A**, pairwise distribution of backbone RMSD values calculated using USAlign on structurally aligned protomers.[5] **B**, comparison of DNA-bound available experimental structures of LacI/GalR TFs.

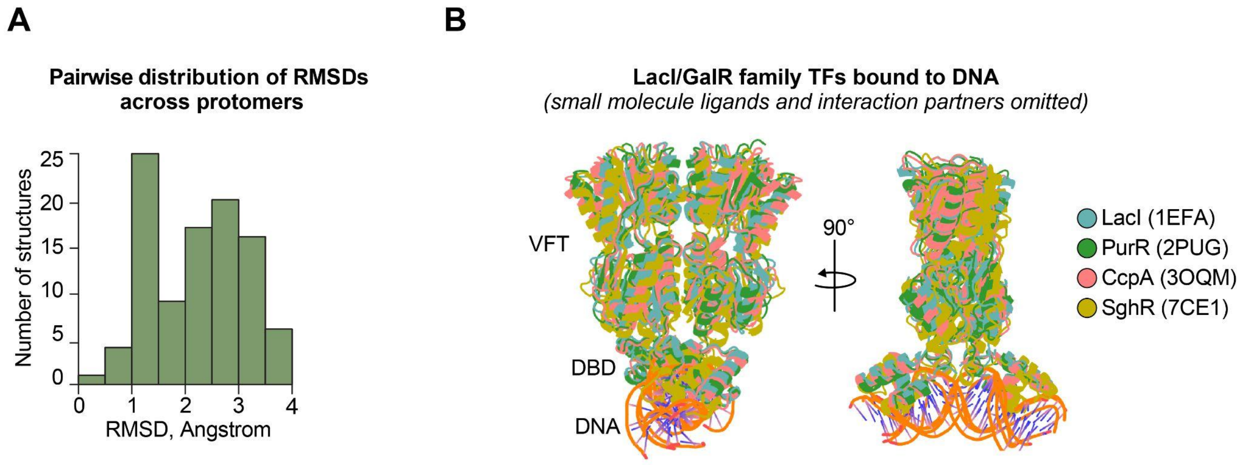

**Figure S5. Molecular details of RbsR-ribose and RbsR-*rbsO* X-ray crystal structures**

Five representative regions of RbsR in both states are highlighted with 2Fo–Fc maps at 1σ.

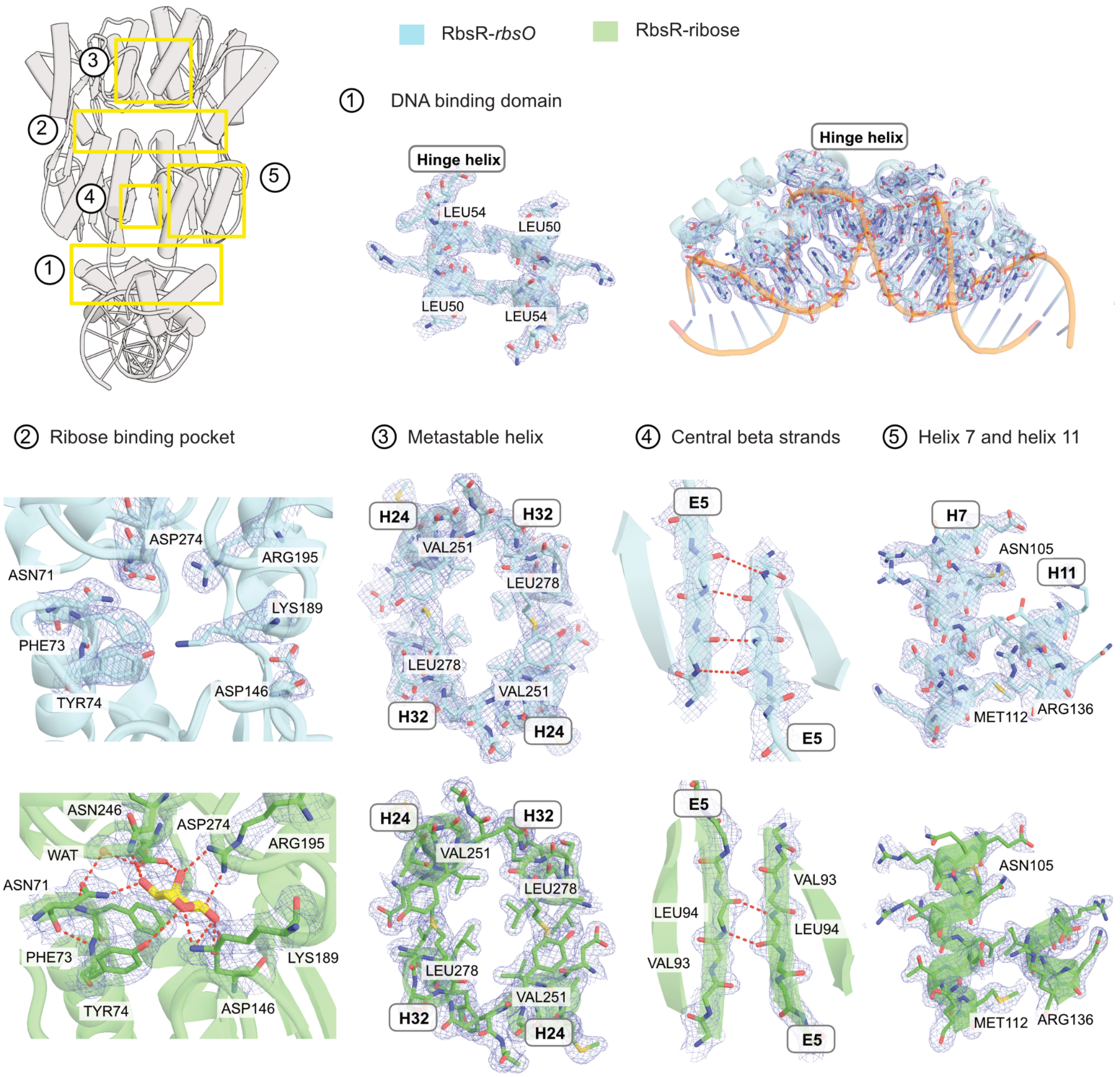

**Figure S6. Peptide level analysis of LacI datasets**

Following peptide list selection by PIGEON[1], we performed a conventional HX/MS data analysis for **A**, APO vs. IPTG (yellow sticks), **B**, APO vs. *lacO1*, and **C**, IPTG vs. *lacO1* states. Colors indicate average deuterium uptake across the timecourse, normalized by peptide length (average ΔD%).[6] *Left:* structural models painted by ΔD%. *Right:* Woods plots showing all the peptides colored by average ΔD%. All the deuterium uptake plots, which include all replicates, are provided as a separate supplemental file.

**A**

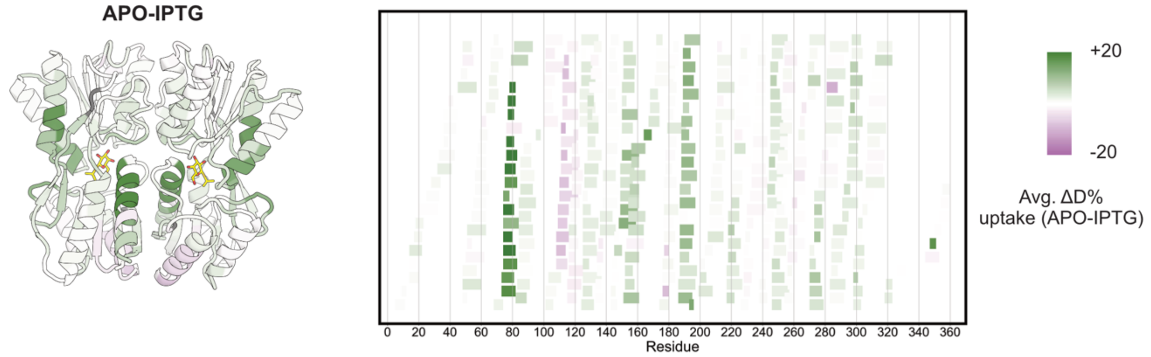

**B**

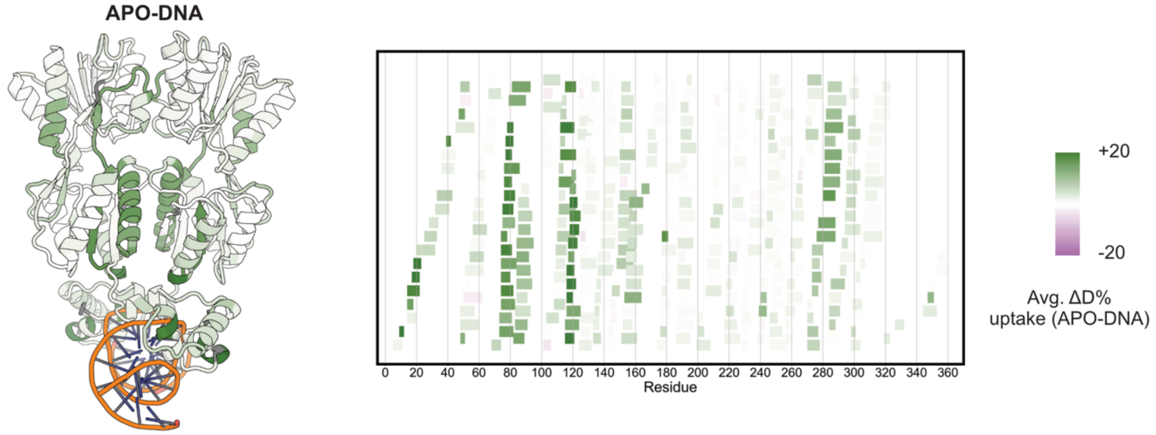

**C**

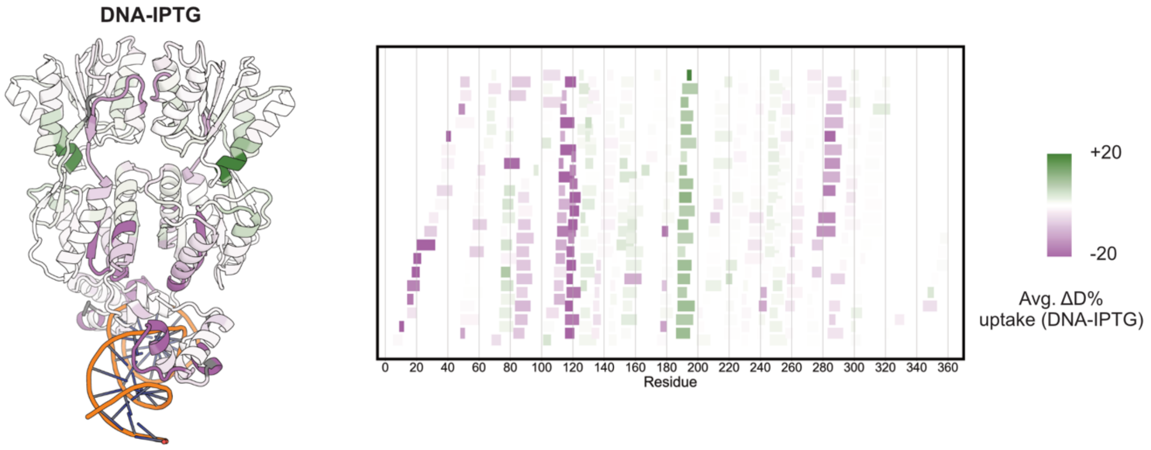

**Figure S7. Peptide level analysis of GalR datasets**

Following peptide list selection by PIGEON[1], we performed a conventional HX/MS data analysis for **A**, APO vs. galactose (yellow sticks), **B**, APO vs. *galO*, and **C**, galactose vs. *galO* states. Colors indicate average deuterium uptake across the timecourse, normalized by peptide length (average ∆D%).[6] *Left:* structural models painted by ∆D%. *Right:* Woods plots showing all the peptides colored by average ∆D%. All the deuterium uptake plots, which include all replicates, are provided as a separate supplemental file.

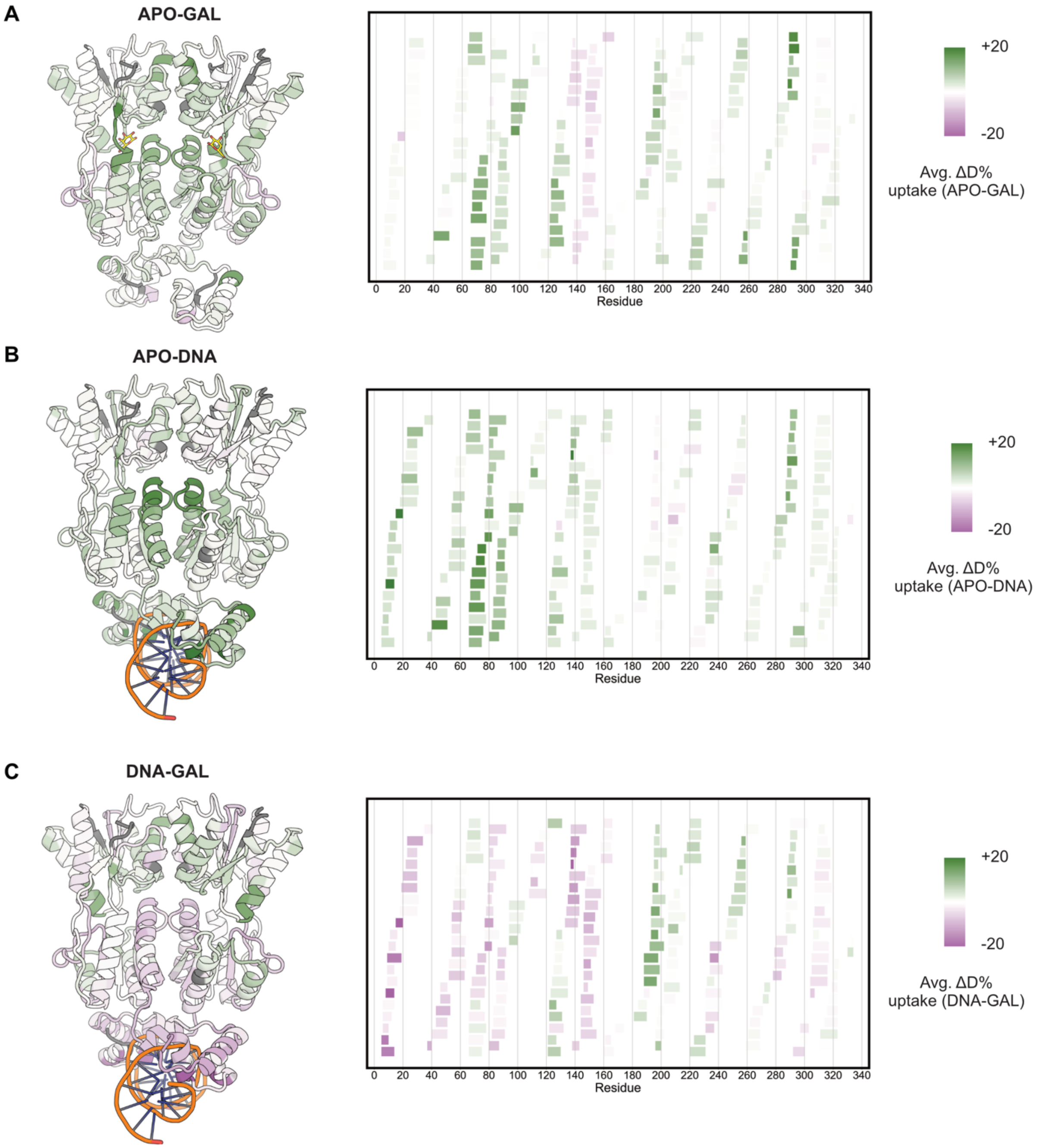

**Figure S8. Peptide level analysis of RbsR datasets**

Following peptide list selection by PIGEON[1], we performed a conventional HX/MS data analysis for **A**, APO vs. ribose (yellow sticks), **B**, APO vs. *rbsO*, and **C**, ribose vs. *rbsO* states. Colors indicate average deuterium uptake across the timecourse, normalized by peptide length (average ∆D%).[6] *Left:* structural models painted by ∆D%. *Right:* Woods plots showing all the peptides colored by average ∆D%.All the deuterium uptake plots, which include all replicates, are provided as a separate supplemental file.

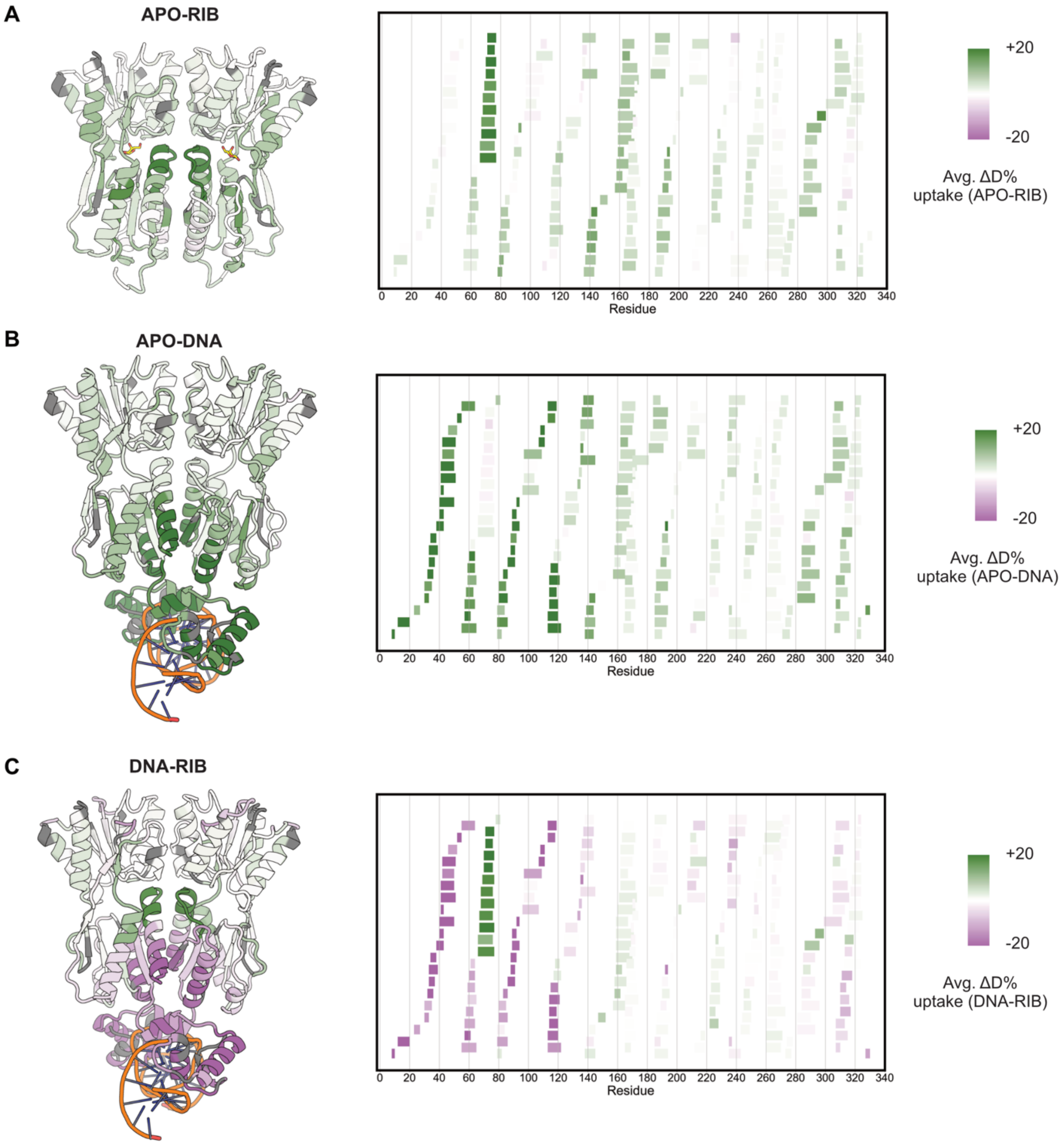

**Figure S9. SEC-MALS and mass photometry analyses of LacI and RbsR**

SEC–MALS measurements were performed at a protein concentration of 3 μM for both proteins, and mass photometry via Refeyn measurements were conducted at 10 nM. Both analyses were carried out in the absence and presence of their respective regulatory ligands as labeled in the plot titles.

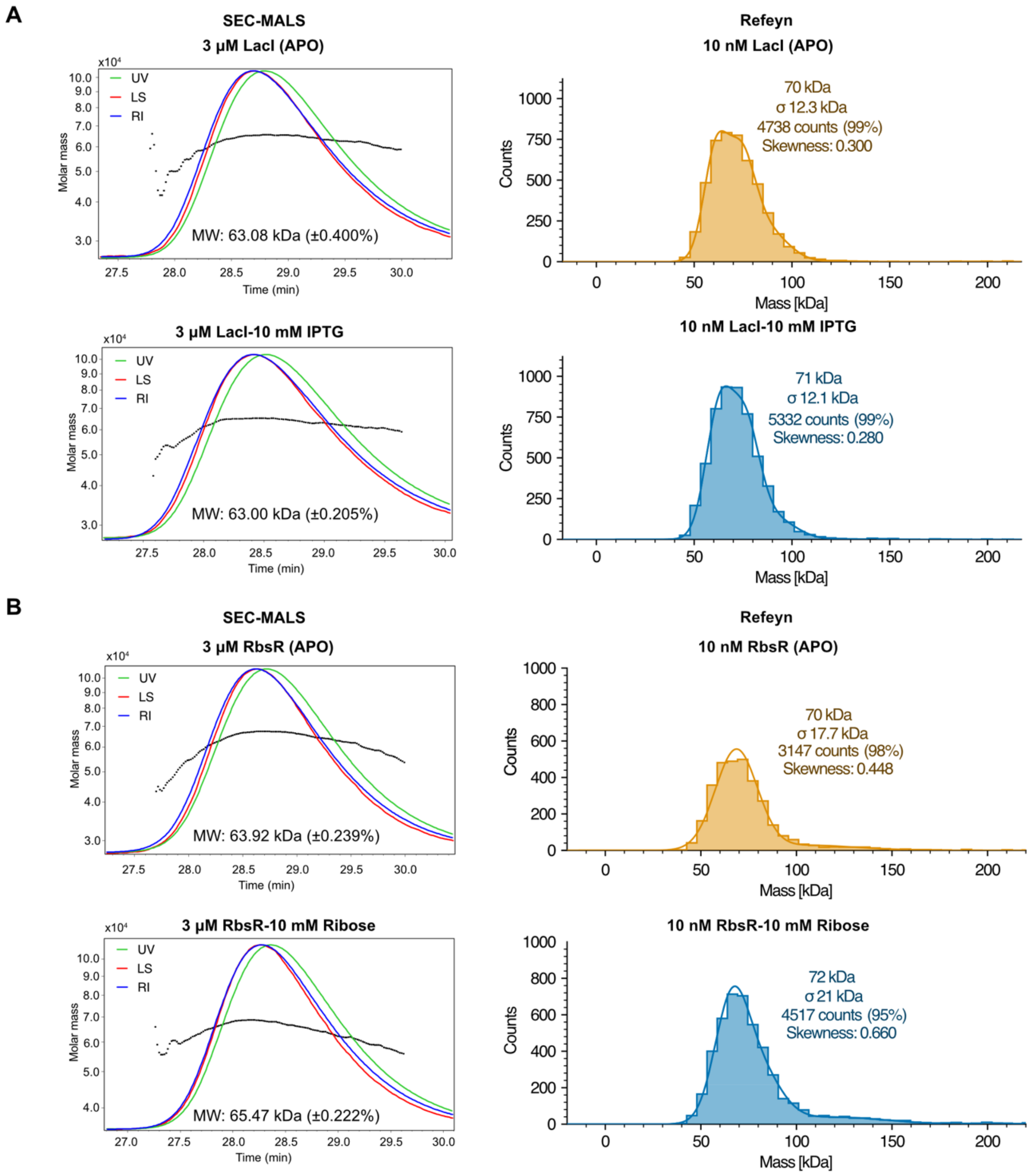

**Figure S10. Peptide level analysis of PBPs datasets**

Following peptide list selection by PIGEON[1], we performed a conventional HX/MS data analysis for **A**, APO vs. ribose in RbsB and **B**, APO vs. galactose in MglB. Colors indicate average deuterium uptake across the timecourse, normalized by peptide length (average ∆D%).[6] *Left:* structural models painted by ∆D%. Sugars are shown in yellow sticks. *Right:* Woods plots showing all the peptides colored by average ∆D%.All the deuterium uptake plots, which include all replicates, are provided as a separate supplemental file.

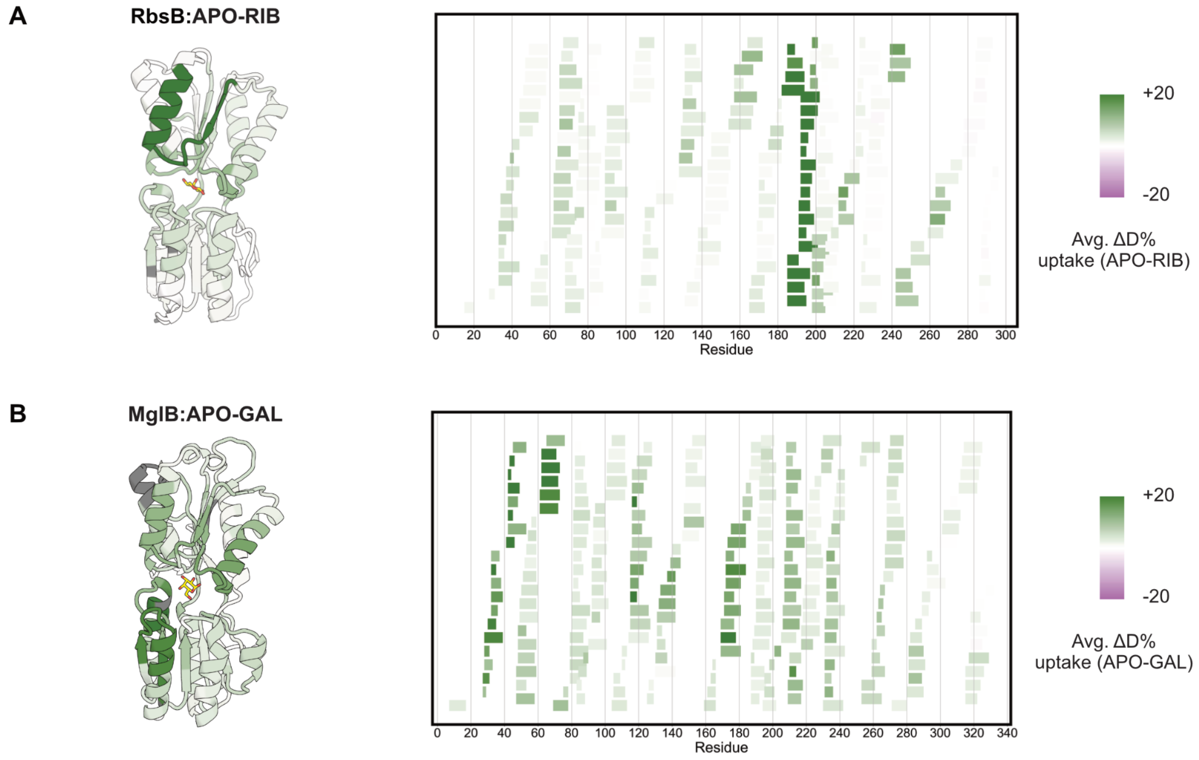

**Figure S11. FEATHER-derived exchange rates for LacI, GalR, and RbsR**

Error bars show bootstrapped uncertainties estimated by FEATHER. Dotted lines show the experimentally accessible $k_{ex}$ window, and red dots mark fast- and slow-exchanging residues outside this window. The blue-white bars underneath the protein sequences show peptide coverage at each residue. The ticks show where there is single-site coverage. The residue index follows the TFs-PBPs structural alignment (alignment file available in Supplemental Data).

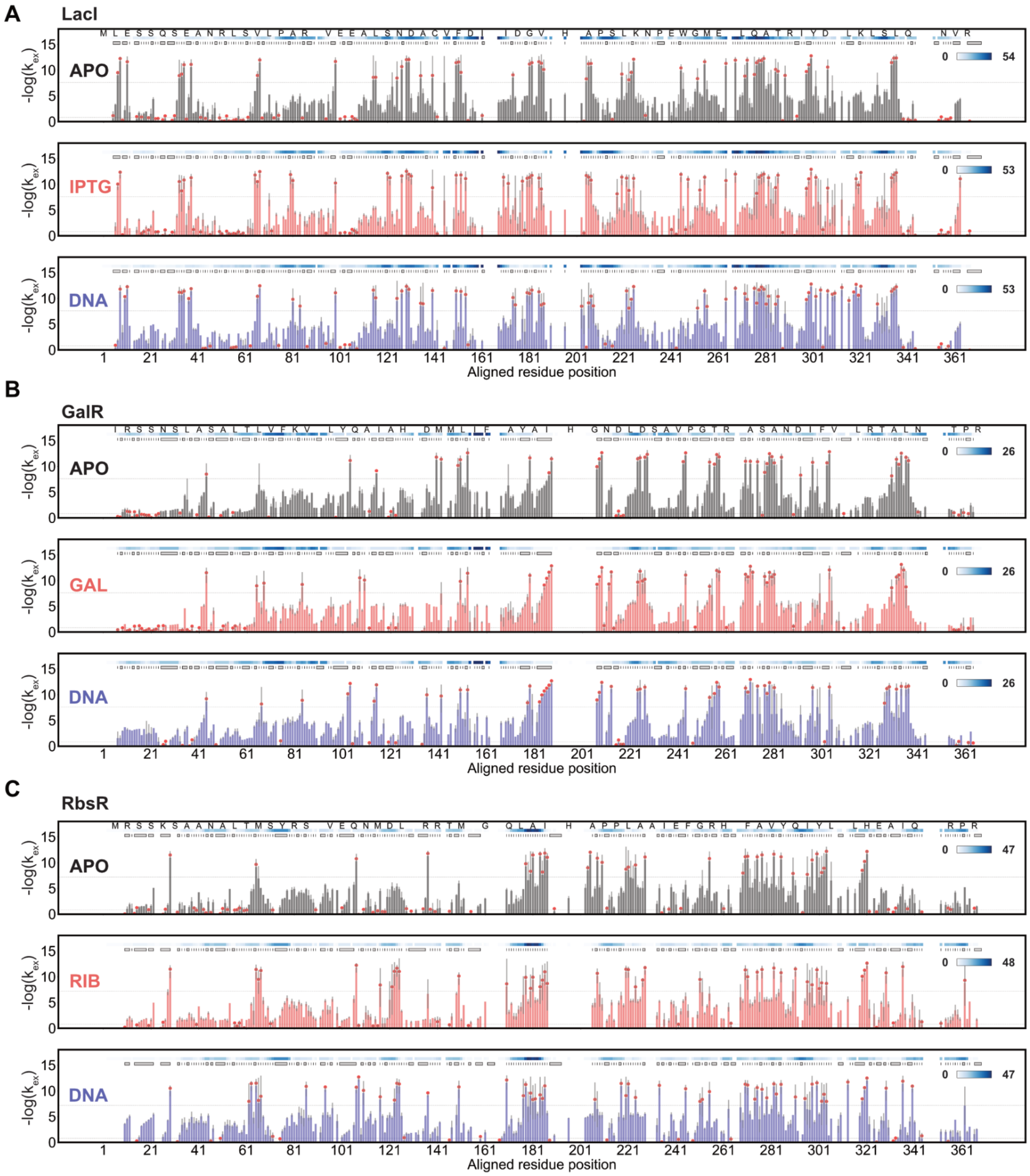

**Figure S12. FEATHER-derived exchange rates for MglB and RbsB**

The figure is formatted similarly to Fig. S11.

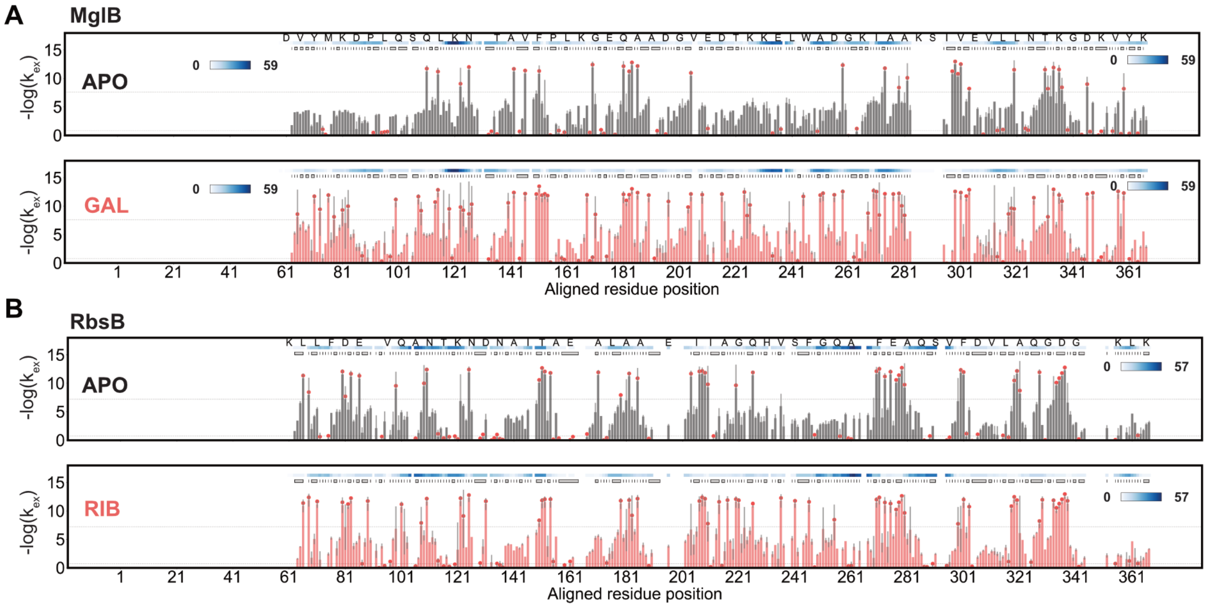

**Figure S13. RbsR and RbsB functional states colored by $\Delta G_{op}$**

**A**, Visual distribution of $\Delta G_{op}$ in apo, operator-bound, and ribose-bound RbsR, as examples of the heatplot in Fig. 1C painted on the protein's structure. **B**, Open (apo) and closed (ribose-bound) states of RbsB, colored also by $\Delta G_{op}$. Ribose shown in yellow sticks.

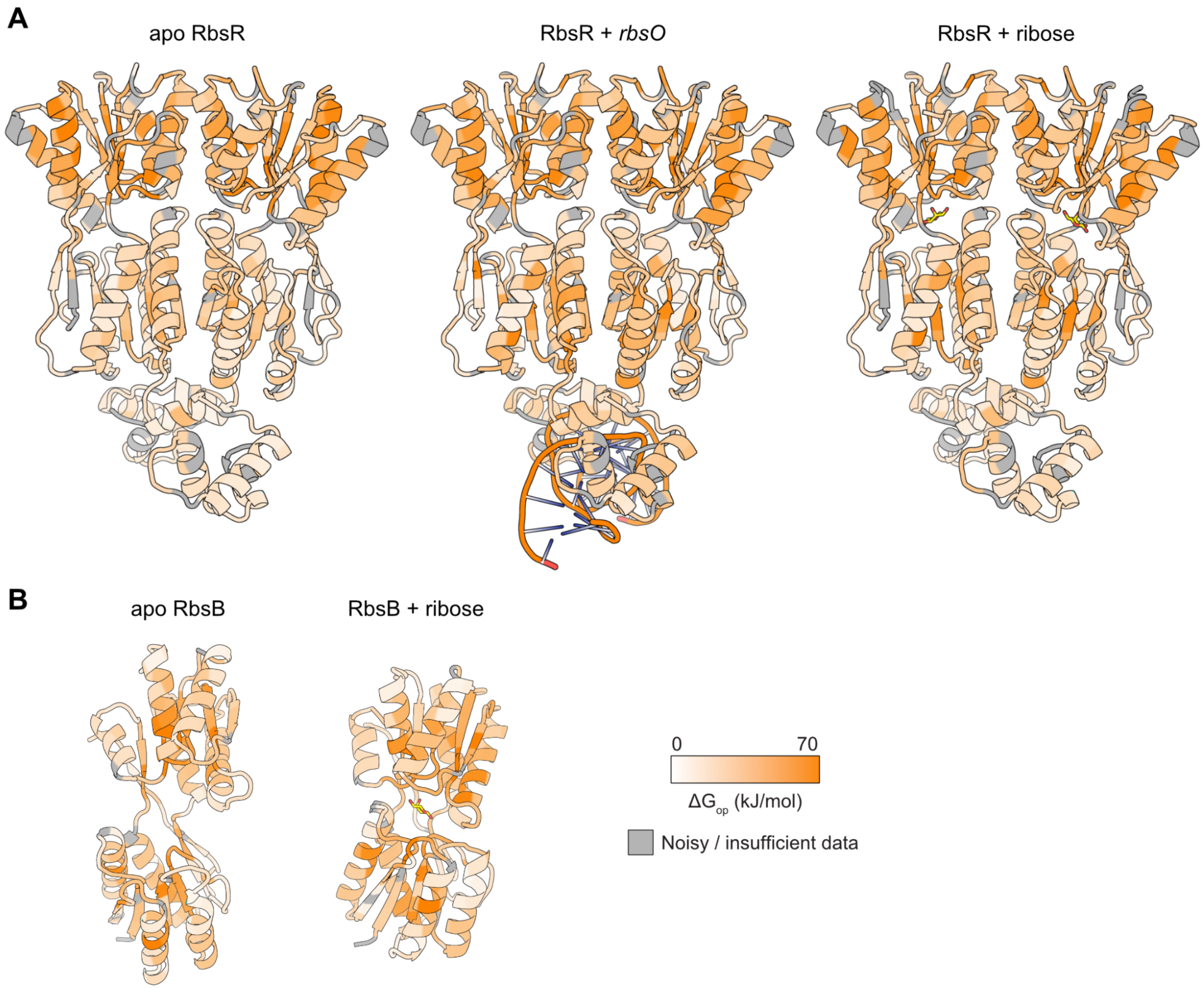

**Figure S14. HX/MS comparison of LacI for single-repeat and double-repeat operator DNA**

We compared replicate HX/MS data for LacI with a single binding site DNA to our main dataset, which was collected with two operator sequences per DNA. We found strong peptide-level agreement between these datasets. Below are representative deuterium uptake plots for peptides from regions where we found functionally important differential protection in the DNA state: **A,** the DNA-binding domain, **B,** peptides near the metastable helix (unfortunately peptide coverage of the helix itself in the singlicate experiment is not identical in the main double-repeat dataset peptides, but the protection in the DNA-bound state extends into adjacent residues where we found agreement), and **C**, helix 7. These regions are annotated on the LacI structure in **D**.

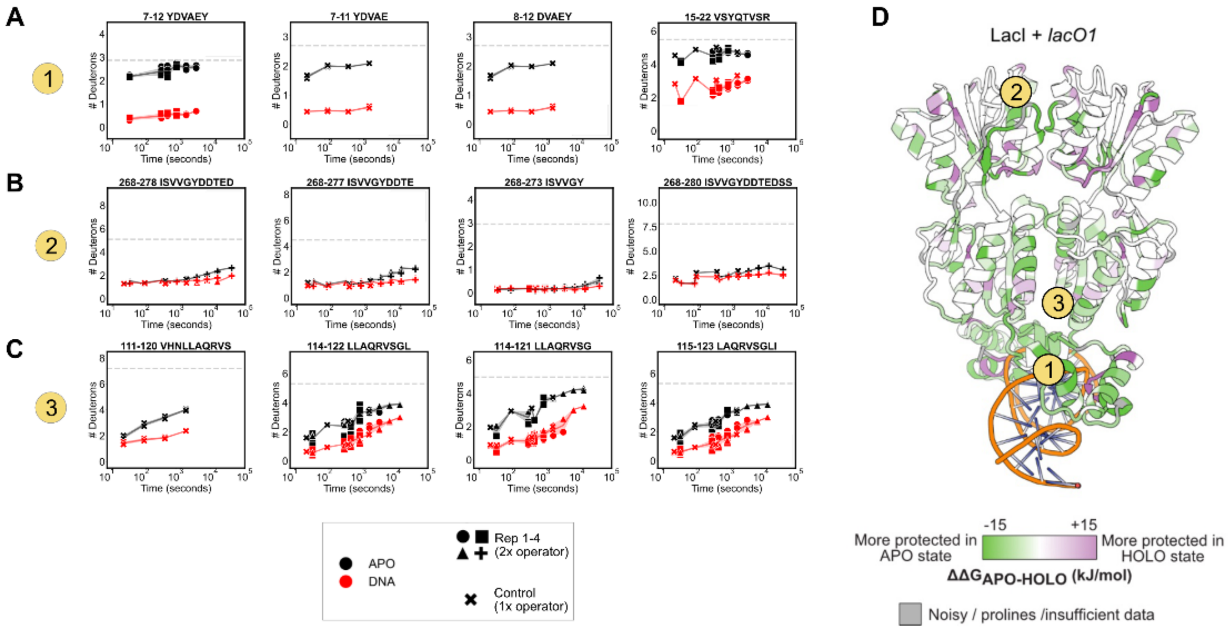

**Figure S15. Peptide-level HX/MS for LacI binding to off-target DNA**

LacI/GalR TFs bind nonspecifically to genomic DNA[7], but nonspecific binding does not bend the DNA.[8] While protection from HX is detected in the DBD for on- and off-target operator sequences (peptides 7-12 and 12-21, *lacO1* and *rbsO*), the (equally strong) protection observed in several key regions of the VFT domain in response to the cognate operator *lacO1* is not detected with other sequences (peptides 75-84, 114-121, 278-291).

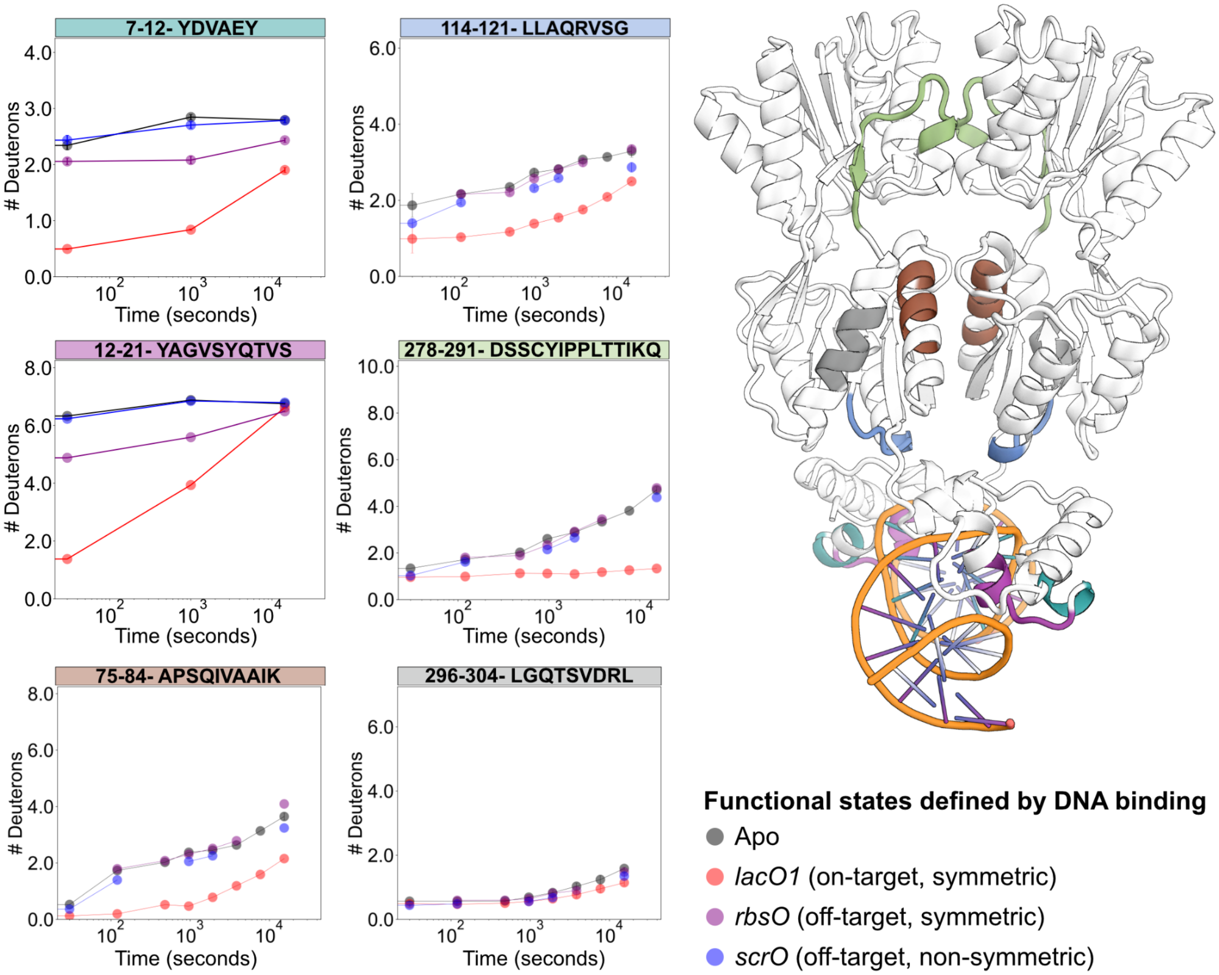

**Figure S16. Effects of sugar binding on TFs and PBPs**

**A**, PBPs respond to binding sugar ligands with global changes in local stability. **B**, TFs and PBPs that bind the same sugars share some patterns of energetic redistribution. **C**, Pan-VFT stabilization effects in binding pocket helices.

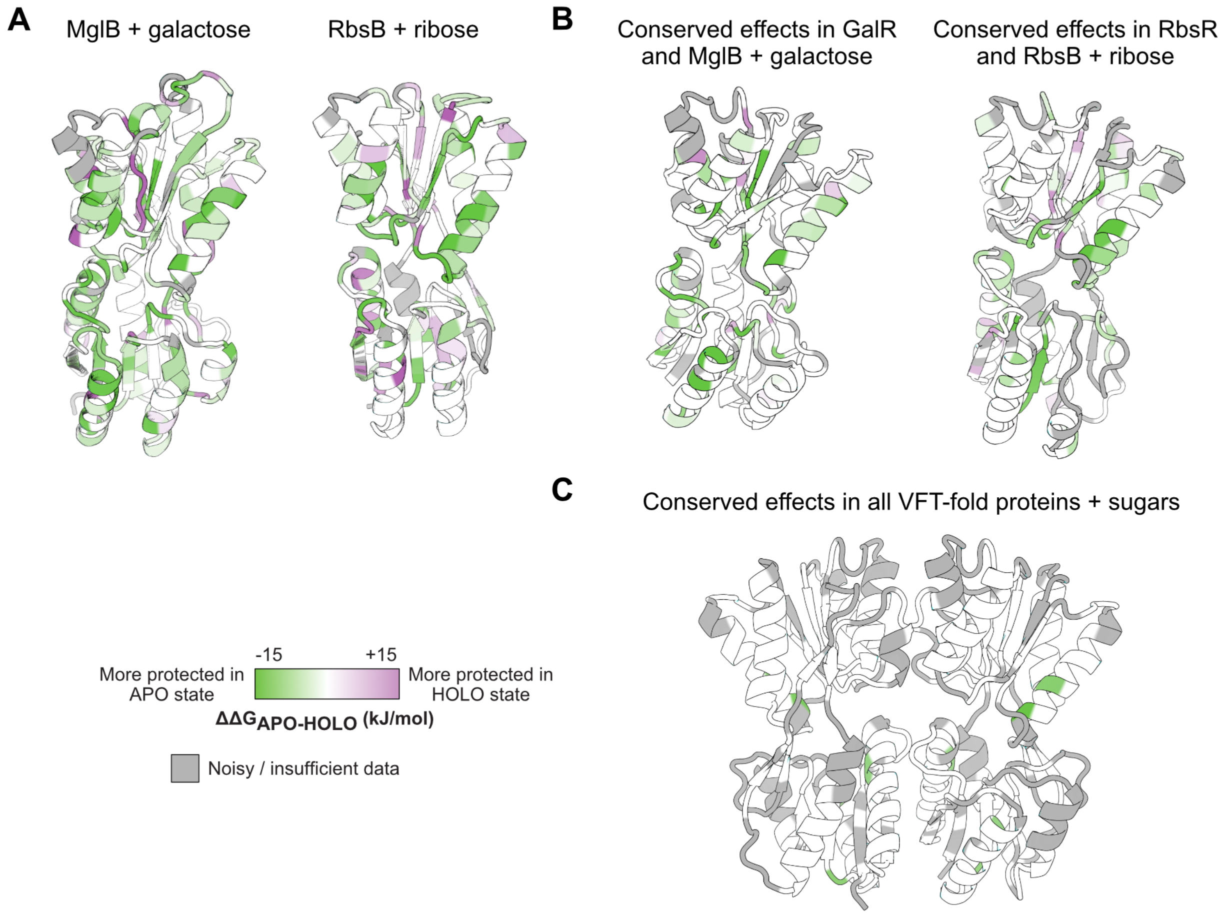

**Figure S17. Conformational changes in RbsB crystal structures**

PBPs like RbsB undergo more dramatic conformational changes in X-ray crystal structures via hinge motions than do LacI/GalR TFs, in which the C-terminal and N-terminal VFT lobes close around the ligand to stabilize the protein in its “closed” conformation.[9,10] Apo PBPs sample both open and closed states in solution.

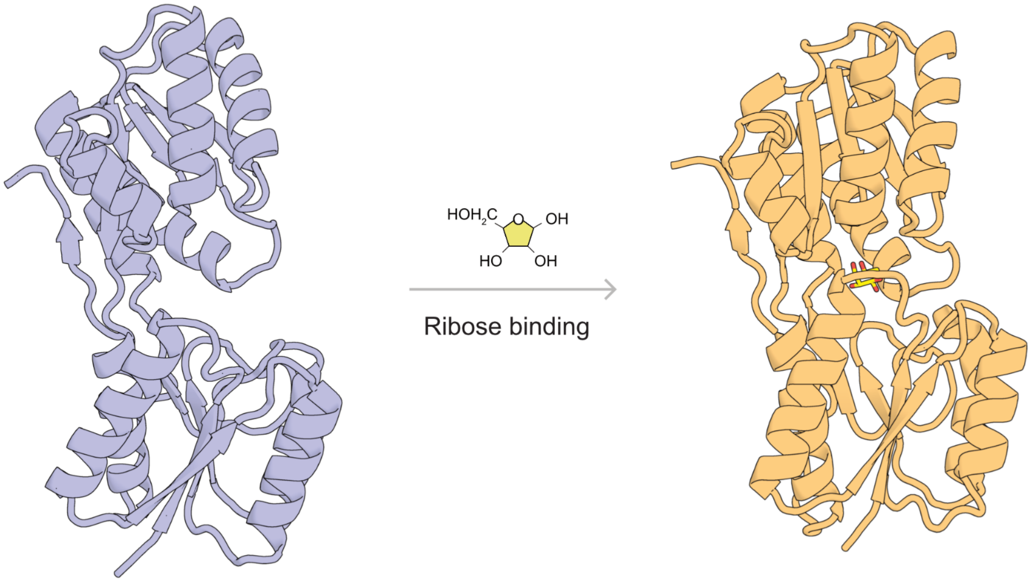

Open (PDB: 1URP)

Closed (PDB: 2DRI)

#### Figure S18. $\Delta\Delta G_{op}$ mapped onto the central β-sheet of LacI and RbsR

The hydrogen bond network in the N-terminal VFT lobe beta sheet is restructured upon ligand binding in both LacI (*left*) and RbsR (*right*), but in different ways. *Top row:* $\Delta\Delta G_{op}$ for DNA vs. apo states. *Bottom row:* $\Delta\Delta G_{op}$ for sugar vs. apo states. Green regions are stabilized in the holo state vs. apo state, purple regions are destabilized.

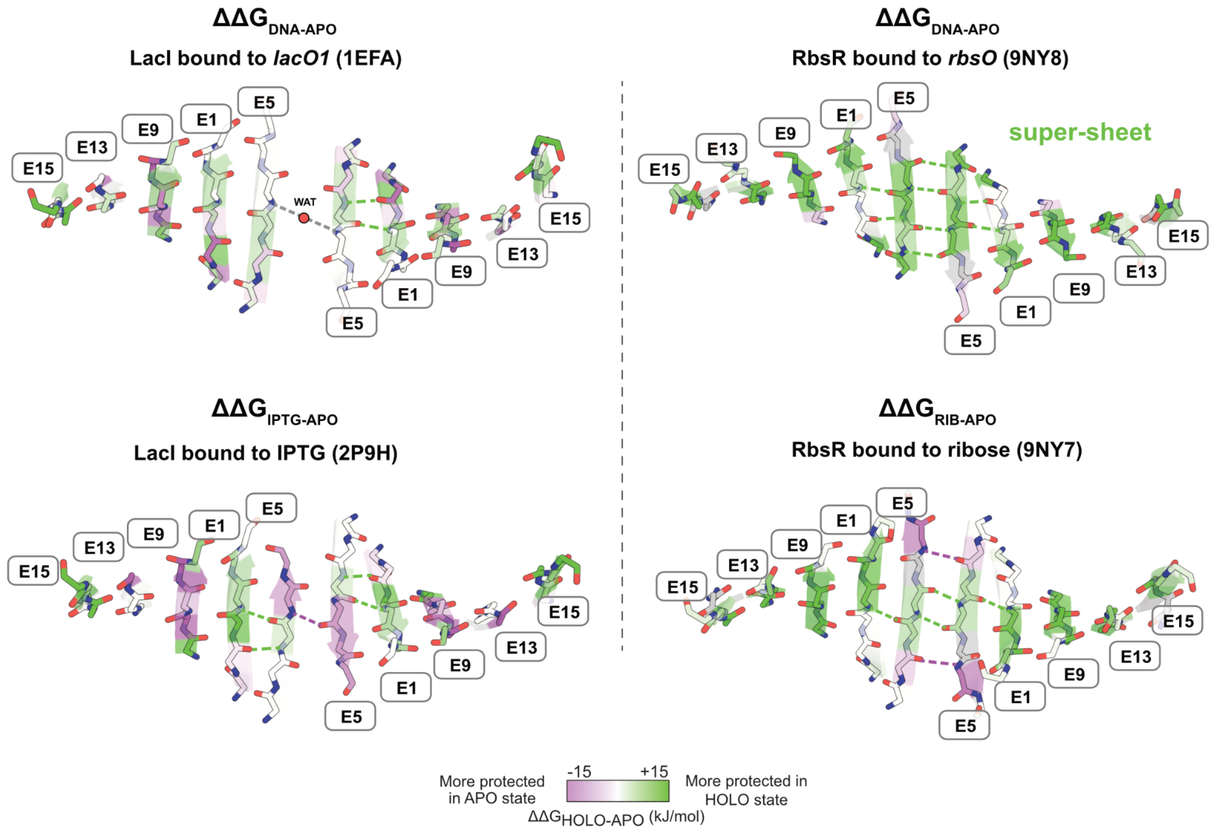

**Figure S19. Design selection strategy**

We selected six metastable helix designs to test experimentally from a pool of 675 designed sequences using a Pareto front optimization strategy that balanced four related energy metrics: interface energy, metastable helix total energy, number of cross-interface hydrogen bonds, and sum of the five least favorable cross-interface interactions (see **Methods**). Pareto fronts are shown as black lines. The colored points correspond to the chosen designs as shown in Fig. 4B-D and Extended Data Fig. 5. The transparent blue points correspond to the other designs.

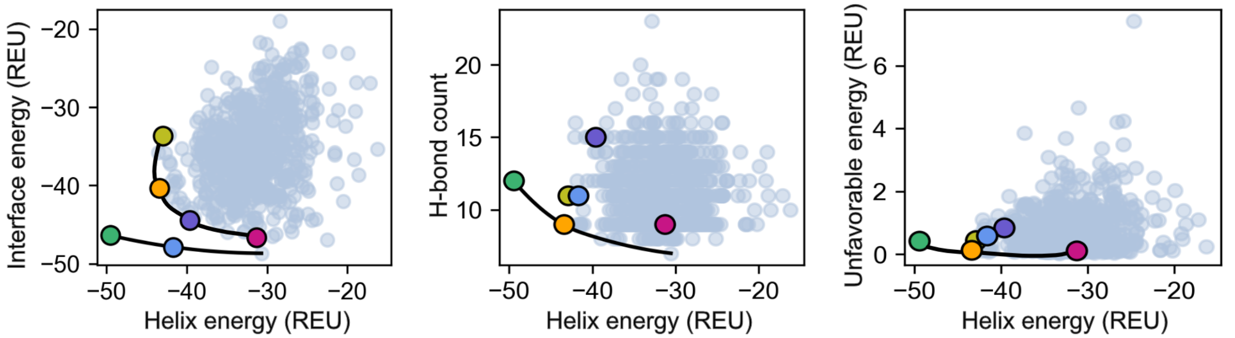

**Figure S20. Western blot and Coomassie blue staining of designed LacI variants**

Protein expression of LacI variants (Designs 1-6) alongside wild-type (WT) and non-transformed knockout (K/O) strains. **A,** Whole cell protein expression visualized by Coomassie blue staining of SDS-PAGE gel**.** Designs labeled D1-6 as in the main text. **B.** Western blotting for the same strains using a mouse anti-LacI primary antibody and an anti-mouse-peroxidase secondary antibody conjugated to horseradish peroxidase, with visualization by chemoluminescence.

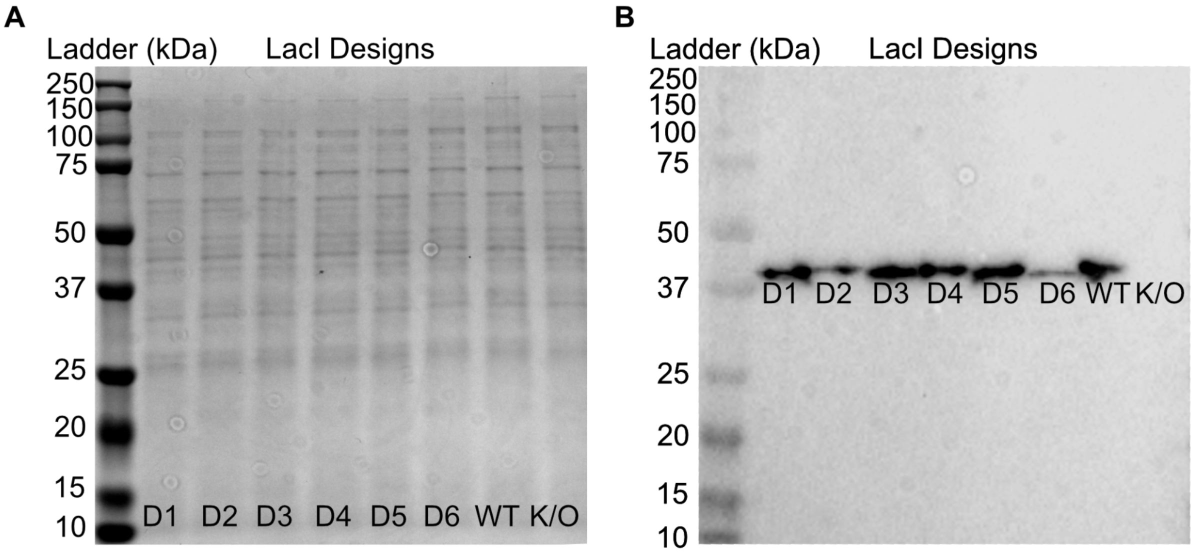

**Figure S21. Mass photometry analyses of LacI-Design 1**

Mass photometry via Refeyn measurements were conducted at 10 nM. The analyses were carried out in the absence and presence of 10 mM IPTG.

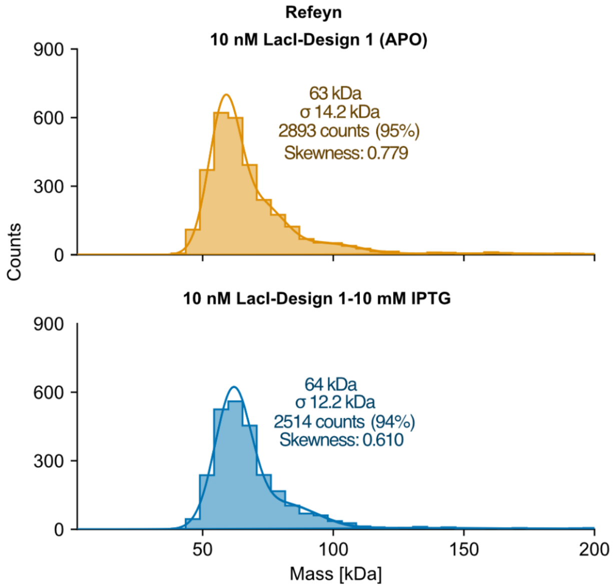

**Figure S22. Conserved structural waters in the binding pocket of sugar-inducible TFs**

**A**, RosettaECO workflow. **B**, The final set of TFs with references to original studies and crystal structures if available. **C**, Rosetta ECO models[11] with all waters shown as spheres. Some of the Kd values were approximated from experimental papers listed in **B** where the exact values were not known.

A

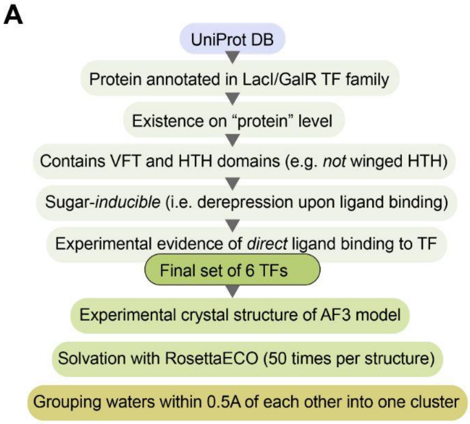

B

| TF | Uniprot ID | PDB ID | References (PMID) |
|---|---|---|---|
| GalS | P25748 | NA | *Geanacopoulos & Adhya, 1997* (8982002) |
| GalR | P03024 | NA | *Geanacopoulos & Adhya, 1997* (8982002) |
| CelR | O87590 | 5YSZ | *Spiridonov & Wilson, 1999* (10224066) |
| | | | *Fu et al., 2018* (30062758) |
| RbsR | P0ACQ0 | 9NY7 (This study) | *Mauzy & Hermodson*, 1992 (1304369) |
| LacI | P03023 | 2P9H | *Gilbert & Müller-Hill, 1966 (16591435)* |
| | | | *Daber et al., 2007* (17543986) |
| TreR | P36673 | 4XXH | *Horlacher & Boos, 1997* (9148912) |
| | | | *Hars et al., 1998* (9865945) |

C

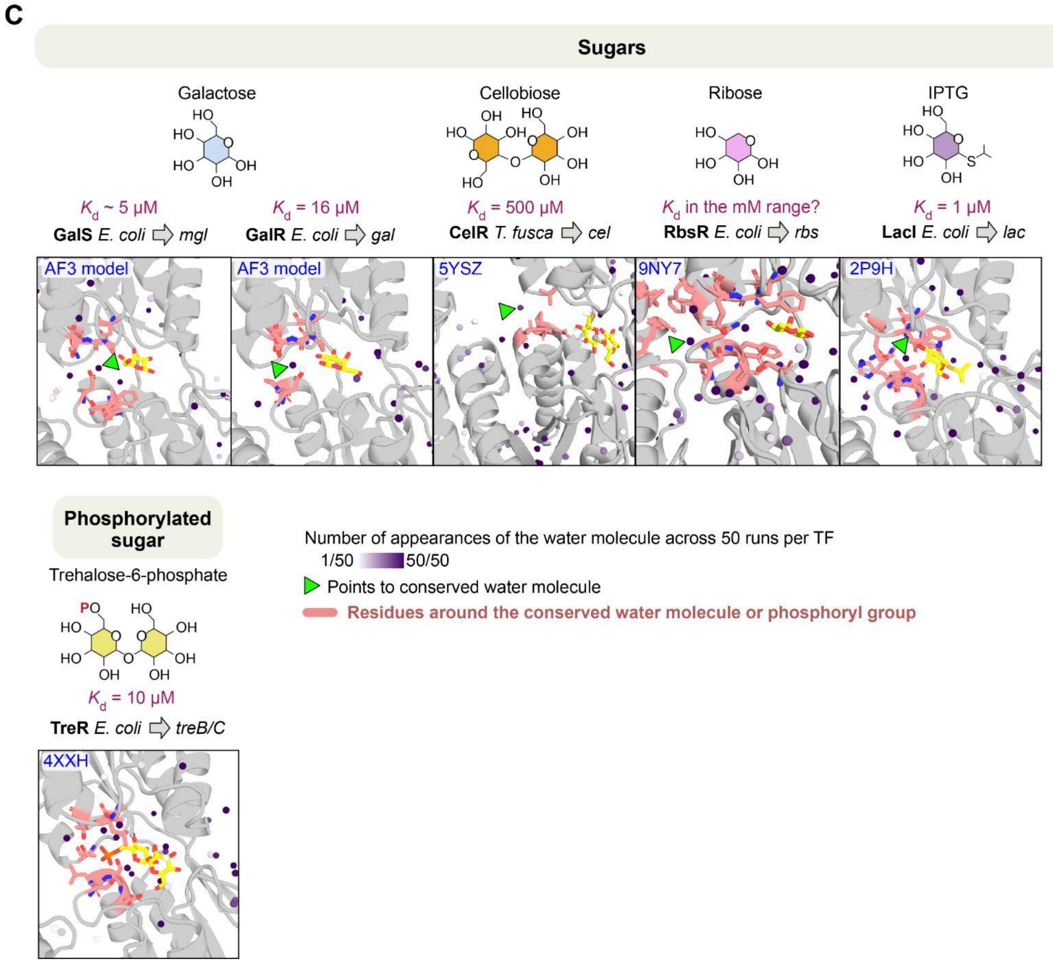

**Figure S23. Water sampling density from MD simulations**

Water clustering centers, identified using *k*-means, shown on the protein structure, and sampling density displayed in principal component analysis (PCA) projection space.

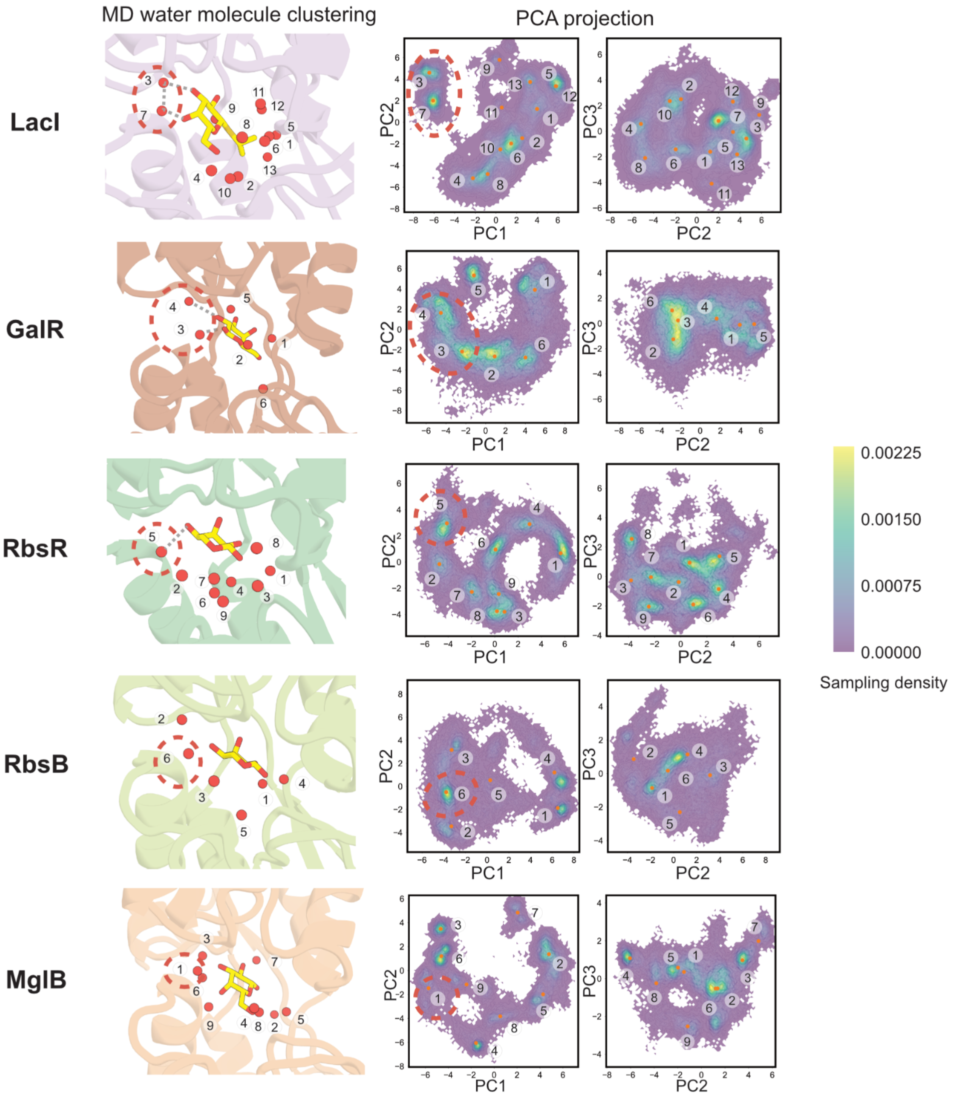

**Figure S24. Structural water analysis of RbsB and MglB**

MD simulations from X-ray crystal structures and water clustering analysis were performed for the PBPs as shown in Fig. 5. As in the TFs, we identified high-confidence structural waters at the same position near H3 (circled in red in **B**, marked by red dots in **C**).

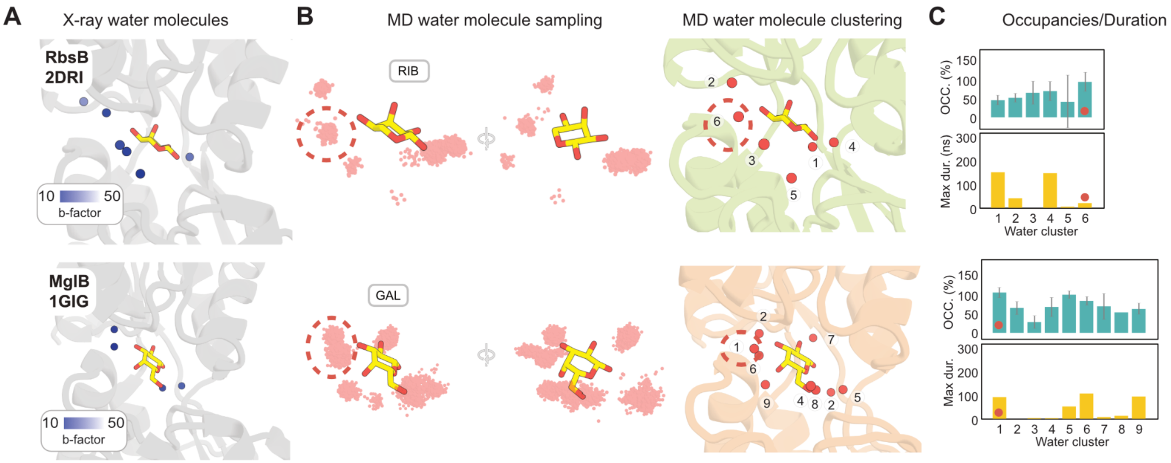

**Figure S25. Comparison of $\Delta G_{op}$ for TF inducer-and operator-bound states**

Comparison of how inducer binding energetically impacts each TF vs. operator binding. All the effects in the same direction with a magnitude above 2 kJ/mol in all the TFs were selected, and $\Delta\Delta G_{op}$ for these positions were averaged to determine conserved effects in TFs in Fig. 6. See **Methods: HX/MS data analysis**.

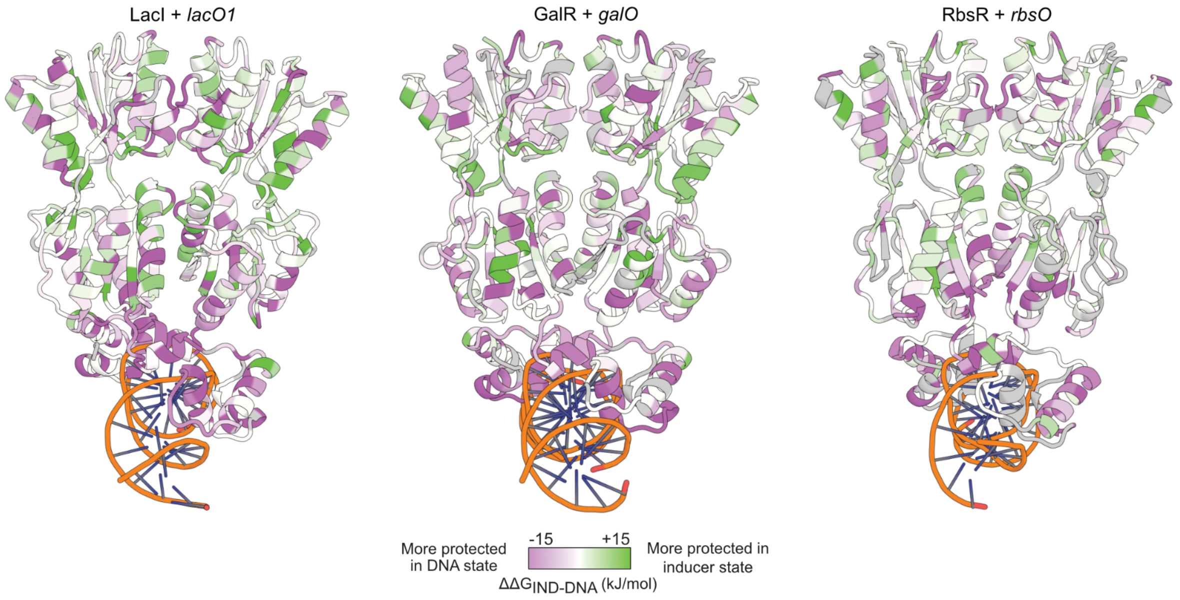

**Figure S26. Comparison of LacI HX/MS with mutational phenotype data**

$\Delta\Delta G_{op,\,(IPTG\text{-}DNA)}$ distribution for residues grouped by three phenotype classes from previous studies[12,13], shown at three mutation count thresholds. The mutation position counts were normalized using min–max scaling to obtain the normalized mutation count (NMC).

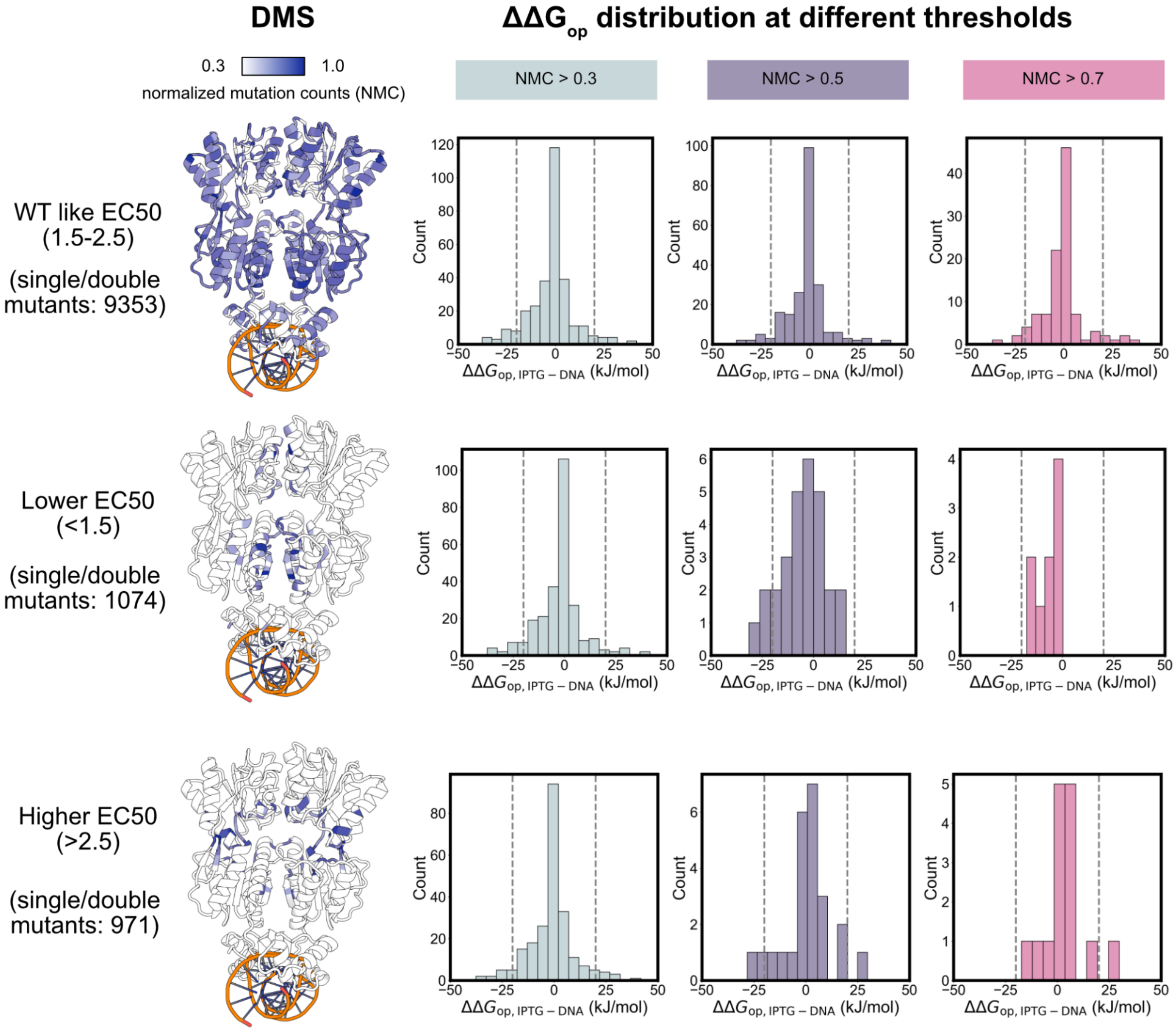

#### Supplemental tables

##### Table S1. Available crystal structures of full-length LacI/GalR TFs

| PDB id | TF | Ligand (PDB name) | DNA present? | Notes |
|---|---|---|---|---|
| 1bdh | PurR | HPA | yes | |
| 1bdi | PurR | HPA | yes | |
| 1efa | LacI | NPF | yes | ligand is anti-inducer |
| 1jfs | PurR | HPA | yes | |
| 1jft | PurR | HPA | yes | |
| 1jh9 | PurR | HPA | yes | |
| 1jwl | LacI | NPF | yes | ligand is anti-inducer |
| 1lbg | LacI | no | yes | |
| 1pnr | PurR | HPA | yes | |
| 1qp0 | PurR | HPA | yes | |
| 1qp4 | PurR | HPA | yes | |
| 1qp7 | PurR | HPA | yes | |
| 1qpz | PurR | HPA | yes | |
| 1qqa | PurR | HPA | yes | |
| 1qqb | PurR | HPA | yes | |
| 1rzr | CcpA | no | yes | with HPr |
| 1vpw | PurR | HPA | yes | |
| 1wet | PurR | GUN | yes | |
| 1zay | PurR | HPA | yes | |
| 1zvv | CcpA | no | yes | with HPr |
| 2hsg | CcpA | no | no | |
| 2jcg | CcpA | no | no | |
| 2pe5 | LacI | 145 | yes | |

| | | | | |
|---|---|---|---|---|
| 2pua | PurR | 6MP | yes | |
| 2pub | PurR | ADE | yes | |
| 2puc | PurR | GUN | yes | |
| 2pud | PurR | HPA | yes | |
| 2pue | PurR | ADE | yes | |
| 2puf | PurR | GUN | yes | |
| 2pug | PurR | HPA | yes | |
| 3gyb | transcriptional regulator from *C. glutamicum* | no | no | |
| 3kjx | transcriptional regulator from *S. pomeroyi* | no | no | |
| 3oqm | CcpA | no | yes | with HPr |
| 3oqn | CcpA | no | yes | with HPr |
| 3oqo | CcpA | no | yes | with HPr |
| 4fe4 | XylR | no | no | |
| 4fe7 | XylR | XYS | no | |
| 4rzs | LacI | no | no | engineered to bind sucralose |
| 4rzt | LacI | RRY_RRJ | no | engineered to bind sucralose |
| 5ysz | CelR | BGC_BGC | no | |
| 7ce1 | SghR | no | yes | |

**Table S2. X-ray crystallography summary statistics**

| | RbsR-ribose (PDB: 9NY7) | RbsR-*rbsO* (PDB: 9NY8) |
|---|---|---|
| **Data collection statistics** | | |
| Beamline | NSLS-II BEAMLINE 19-ID | NSLS-II BEAMLINE 19-ID |
| Wavelength (Å) | 0.979493 | 0.979493 |
| Space group | C2 | $P3_121$ |
| a, b, c (Å) | 177.46, 87.35, 130.94 | 125.53, 125.53, 122.31 |
| α, β, γ (°) | 90.00, 100.38, 90.00 | 90.00, 90.00, 120.00 |
| $Z_a$[a] / Solvent content (%) | 6/53.82 | 2/67.68 |
| Resolution range (Å) | 33.80 - 2.07 | 43.80 - 2.10 |
| Total reflections | 413544 | 1458741 |
| Unique reflections | 119865 | 47896 |
| Completeness (%) | 99.9 | 94.1 |
| Multiplicity | 3.5 | 30.5 |
| $CC_{1/2}$ | 0.968 | 0.999 |
| $R_{merge}$ | 0.063 | 0.245 |
| $R_{pim}$ | 0.228 | 0.045 |
| I/σ | 2.5 | 15.7 |
| **Refinement statistics** | | |
| Reflection used | 73891 | 47896 |
| R-factor | 0.192 | 0.184 |
| Free R-factor | 0.239 | 0.224 |
| RMSD from ideal | | |
| Bond length (Å) | 0.40 | 0.51 |
| Angle (°) | 0.64 | 0.81 |
| Ramachandran (%) | | |
| Favored | 97 | 97 |
| Allowed | 2 | 3 |

[a] Za is the number of subunits per asymmetric unit.

**Table S3. HX/MS replicate information for LacI**

Adapted from Lu *et al.*[1]. Abbreviations: FP, immobilized protease type XIII. Nep, nepenthesin 2. AP, alanyl aminopeptidase. Tables S3-S7 include all experimental reporting suggestions included in Masson *et al.*[14]

| Dataset | **LacI Rep1** | **LacI Rep2** | **LacI Rep3** | **LacI Rep4** |
|---|---|---|---|---|
| Date | 11/18/22 | 01/14/23 | 03/15/24 | 08/08/24 |
| Protease column | FP/pepsin | FP/pepsin | FP/pepsin | AP/pepsin |
| Protein states | APO/DNA/IPTG | APO/DNA/IPTG | APO/DNA/IPTG | APO/DNA/IPTG |
| Exchange conditions | 50 mM Tris, 150 mM NaCl, pH 8.0, temperature 15 °C | | | |
| Protein conc. (μM) | 1.3 | 2.0 | 1.7 | 1.6 |
| Quench solution GuHCl conc. (M) | 2 | 3 | | |
| Time course (s) | 4.5e1 to 3.9e3 | 4.5e1 to 1.5e4 | 4.5e1 to 1.5e4 | 4.5e1 to 4.0e4 |
| # of time points | 6 | 5 | 8 | 9 |
| Control samples | Maximally labeled sample (full-D) 3 M GuHCl | | | |
| Back-exchange (mean / IQR) | 0.20/0.12 | 0.20/0.11 | 0.23/0.10 | 0.24/0.10 |
| # of peptides | 189 | 187 | 240 | 240 |
| Sequence coverage | 0.81 | 0.79 | 0.91 | 0.87 |
| Average peptide length / Redundancy | 1.9 | 2.0 | 1.5 | 1.5 |
| Significant difference in $\Delta G_{op}$ | $mean\{\Delta G_{op,a,i} - \Delta G_{op,b,i}\} > std\{\Delta G_{op,a,i} - \Delta G_{op,b,i}\}$ | | | |

**Table S4. HX/MS replicate information for GalR**

Abbreviations: FP, immobilized protease type XIII. Nep, nepenthesin 2. AP, alanyl aminopeptidase.

| Dataset | **GalR Rep1** | **GalR Rep2** | **GalR Rep3** |
|---|---|---|---|
| Date | 05/13/23 | 12/05/23 | 11/27/24 |
| Protease column | FP/pepsin | FP/pepsin | AP/pepsin |
| Protein states | APO/DNA/GAL | APO/DNA/GAL | APO/DNA/GAL |
| Exchange conditions | 50 mM Tris, 150 mM NaCl, pH 8.0, temperature 15 °C | 50 mM Tris, 600 mM NaCl, 0.5 mM TCEP, pH 8.0, temperature 15 °C | |
| Protein conc. (μM) | 1.6 | 1.5 | 0.7 |
| Quench solution GuHCl conc. (M) | 3 | | |
| Time course (s) | 4.8e1 to 7.8e3 | 4.6e1 to 1.5e4 | 5.5e1 to 4.0e4 |
| # of time points | 8 | 8 | 7 |
| Control samples | Maximally labeled sample (full-D) 3 M GuHCl | | 0 M GuHCl |
| Back-exchange (mean / IQR) | 0.29/0.12 | 0.30/0.09 | 0.28/0.11 |
| # of peptides | 57 | 120 | 142 |
| Sequence coverage | 0.58 | 0.79 | 0.76 |
| Average peptide length / Redundancy | 6.2 | 2.9 | 2.5 |
| Significant difference in $\Delta G_{op}$ | $mean\{\Delta G_{op,a,i} - \Delta G_{op,b,i}\} > std\{\Delta G_{op,a,i} - \Delta G_{op,b,i}\}$ | | |

**Table S5. HX/MS replicate information for RbsR**

Abbreviations: FP, immobilized protease type XIII. Nep, nepenthesin 2. AP, alanyl aminopeptidase.

| Dataset | **RbsR Rep1** | **RbsR Rep2** | **RbsR Rep3** | **RbsR Rep4** |
|---|---|---|---|---|
| Date | 08/31/22 | 01/06/23 | 02/22/23 | 05/30/24 |
| Protease column | FP/pepsin | FP/pepsin | FP/pepsin | NP/pepsin |
| Protein states | APO/DNA/RIB | APO/DNA/RIB | APO/DNA/RIB | APO/DNA/RIB |
| Exchange conditions | 50 mM Tris, 150 mM NaCl, pH 8.0, temperature 15 °C | | | |
| Protein conc. (μM) | 0.6 | 1.6 | 1.4 | 5.2 |
| Quench solution GuHCl conc. (M) | 3 | | | |
| Time course (s) | 5.0e1 to 1.5e4 | 4.5e1 to 1.5e4 | 4.5e1 to 7.7e3 | 4.2e1 to 1.5e4 |
| # of time points | 8 | 8 | 8 | 8 |
| Control samples | Maximally labeled sample (full-D) 3 M GuHCl | | | |
| Back-exchange (mean / IQR) | 0.20/0.13 | 0.19/0.14 | 0.18/0.14 | 0.22/0.11 |
| # of peptides | 100 | 110 | 149 | 140 |
| Sequence coverage | 0.71 | 0.79 | 0.85 | 0.80 |
| Average peptide length / Redundancy | 2.4 | 3.1 | 2.4 | 2.4 |
| Significant difference in $\Delta G_{op}$ | $mean\{\Delta G_{op,a,i} - \Delta G_{op,b,i}\} > std\{\Delta G_{op,a,i} - \Delta G_{op,b,i}\}$ | | | |

**Table S6. HX/MS replicate information for MglB**

Abbreviations: FP, immobilized protease type XIII. Nep, nepenthesin 2. AP, alanyl aminopeptidase.

| Dataset | **MglB Rep1** | **MglB Rep2** | **MglB Rep3** | **MglB Rep4** |
|---|---|---|---|---|
| Date | 06/06/23 | 06/06/23 | 08/24/24 | 09/14/24 |
| Protease column | FP/pepsin | FP/pepsin | AP/pepsin | NP/pepsin |
| Protein states | APO/GAL | APO/GAL | APO/GAL | APO/GAL |
| Exchange conditions | 50 mM Tris, 150 mM NaCl, pH 8.0, temperature 15 °C | | | |
| Protein conc. (μM) | 4.1 | 4.1 | 5.0 | 5.5 |
| Quench solution GuHCl conc. (M) | 3 | | | |
| Time course (s) | 5.2e1 to 1.5e4 | 5.2e1 to 7.7e4 | 4.2e1 to 4.0e4 | 4.2e1 to 4.0e4 |
| # of time points | 9 | 7 | 7 | 7 |
| Control samples | Maximally labeled sample (full-D) 0M GuHCl | | | |
| Back-exchange (mean / IQR) | 0.23/0.09 | 0.23/0.09 | 0.21/0.09 | 0.22/0.12 |
| # of peptides | 147 | 146 | 99 | 165 |
| Sequence coverage | 0.83 | 0.85 | 0.58 | 0.85 |
| Average peptide length / Redundancy | 2.3 | 2.3 | 3.4 | 2.0 |
| Significant difference in $\Delta G_{op}$ | $mean\{\Delta G_{op,a,i} - \Delta G_{op,b,i}\} > std\{\Delta G_{op,a,i} - \Delta G_{op,b,i}\}$ | | | |

**Table S7. HX/MS replicate information for RbsB**

The RbsB protein sequence does not contain any tryptophans, so protein concentrations could not be measured by absorbance at 280 nM. Instead, following the determination of protein purity by SDS-PAGE with Coomassie staining, samples were diluted to an approximately 10 µM stock concentration and flash frozen for HX/MS experiments. Abbreviations: FP, immobilized protease type XIII. Nep, nepenthesin 2. AP, alanyl aminopeptidase.

| Dataset | **RbsB Rep1** | **RbsB Rep2** | **RbsB Rep3** | **RbsB Rep4** |
|---|---|---|---|---|
| Date | 10/18/22 | 10/25/22 | 03/11/23 | 04/23/23 |
| Protease column | FP/pepsin | FP/pepsin | FP/pepsin | FP/pepsin |
| Protein states | APO/RIB | APO/RIB | APO/RIB | APO/RIB |
| Exchange conditions | 50 mM Tris, 150 mM NaCl, pH 8.0, temperature 15 °C | | | |
| Protein conc. (µM) | n/a | n/a | n/a | n/a |
| Quench solution GuHCl conc. (M) | 1 | | 3 | |
| Time course (s) | 3.9e1 to 1.5e4 | 3.9e1 to 1.5e4 | 4.6e1 to 1.5e4 | 4.6e1 to 1.5e4 |
| # of time points | 9 | 9 | 8 | 8 |
| Control samples | Maximally labeled sample (full-D) 3M GuHCl | | | |
| Back-exchange (mean / IQR) | 0.23/0.09 | 0.22/0.10 | 0.27/0.07 | 0.26/0.09 |
| # of peptides | 95 | 169 | 120 | 172 |
| Sequence coverage | 0.74 | 0.86 | 0.72 | 0.84 |
| Average peptide length / Redundancy | 3.2 | 1.8 | 2.5 | 1.7 |
| Significant difference in $\Delta G_{op}$ | $mean\{\Delta G_{op,a,i} - \Delta G_{op,b,i}\} > std\{\Delta G_{op,a,i} - \Delta G_{op,b,i}\}$ | | | |

**Table S8. $\Delta G_{op}$ values for LacI**

All values in kJ/mol.

| Res. name | Res. number | APO | Std Dev | Single resolved | IPTG | Std Dev | Single resolved | *lacO1* | Std Dev | Single resolved |
|---|---|---|---|---|---|---|---|---|---|---|
| MET | 1 | | | | | | | | | |
| LYS | 2 | | | | | | | | | |
| PRO | 3 | | | | | | | | | |
| VAL | 4 | | | | | | | | | |
| THR | 5 | 40 | 2 | FALSE | 39 | 2 | FALSE | 24 | 2 | FALSE |
| LEU | 6 | 39 | 2 | FALSE | 38 | 2 | FALSE | 23 | 2 | FALSE |
| TYR | 7 | 38 | 2 | FALSE | 37 | 2 | FALSE | 23 | 2 | FALSE |
| ASP | 8 | 42 | 2 | FALSE | 41 | 2 | FALSE | 26 | 2 | FALSE |
| VAL | 9 | 27 | 3 | FALSE | 12 | 1 | FALSE | 50 | 3 | FALSE |
| ALA | 10 | 31 | 3 | FALSE | 16 | 1 | FALSE | 54 | 3 | FALSE |
| GLU | 11 | 32 | 3 | FALSE | 16 | 1 | FALSE | 54 | 3 | FALSE |
| TYR | 12 | 16 | 6 | TRUE | 14 | 0 | TRUE | 32 | 0 | TRUE |
| ALA | 13 | | | | | | | | | |
| GLY | 14 | 11 | 1 | FALSE | 15 | 1 | FALSE | 19 | 0 | FALSE |
| VAL | 15 | 8 | 1 | FALSE | 12 | 1 | FALSE | 16 | 0 | FALSE |
| SER | 16 | 12 | 1 | FALSE | 16 | 1 | FALSE | 20 | 0 | FALSE |
| TYR | 17 | <13 | 6 | TRUE | <13 | 5 | TRUE | 26 | 0 | TRUE |
| GLN | 18 | 15 | 5 | TRUE | <13 | 5 | TRUE | 30 | 4 | TRUE |
| THR | 19 | 20 | 0 | TRUE | 18 | 1 | TRUE | 19 | 1 | TRUE |
| VAL | 20 | <10 | 4 | TRUE | <10 | 5 | TRUE | 14 | 7 | TRUE |
| SER | 21 | 26 | 2 | FALSE | 28 | 1 | FALSE | 36 | 0 | FALSE |
| ARG | 22 | 27 | 2 | FALSE | 29 | 1 | FALSE | 37 | 0 | FALSE |
| VAL | 23 | <10 | 3 | TRUE | <10 | 4 | TRUE | 26 | 0 | TRUE |
| VAL | 24 | <8 | 2 | TRUE | <8 | 2 | TRUE | 26 | 0 | TRUE |
| ASN | 25 | <15 | 1 | FALSE | <15 | 1 | FALSE | 18 | 1 | FALSE |
| GLN | 26 | <15 | 1 | FALSE | <15 | 1 | FALSE | 19 | 1 | FALSE |
| ALA | 27 | <14 | 1 | FALSE | <14 | 1 | FALSE | 18 | 1 | FALSE |

| | | | | | | | | | | |
|---|---|---|---|---|---|---|---|---|---|---|
| SER | 28 | <15 | 1 | FALSE | 10 | 1 | FALSE | 17 | 1 | FALSE |
| HIS | 29 | <15 | 1 | FALSE | 10 | 1 | FALSE | 17 | 1 | FALSE |
| VAL | 30 | <10 | 1 | FALSE | 5 | 1 | FALSE | 12 | 1 | FALSE |
| SER | 31 | <14 | 1 | FALSE | 9 | 1 | FALSE | 16 | 1 | FALSE |
| ALA | 32 | 32 | 1 | TRUE | 31 | 2 | TRUE | 34 | 0 | TRUE |
| LYS | 33 | >50 | 20 | TRUE | >50 | 11 | TRUE | >50 | 8 | TRUE |
| THR | 34 | >50 | 19 | TRUE | >50 | 19 | TRUE | >50 | 11 | TRUE |
| ARG | 35 | >52 | 6 | TRUE | >52 | 6 | TRUE | >52 | 6 | TRUE |
| GLU | 36 | 38 | 4 | FALSE | 55 | 4 | FALSE | 50 | 2 | FALSE |
| LYS | 37 | 36 | 4 | FALSE | 53 | 4 | FALSE | 48 | 2 | FALSE |
| VAL | 38 | 34 | 4 | FALSE | 51 | 4 | FALSE | 46 | 2 | FALSE |
| GLU | 39 | 16 | 0 | TRUE | <11 | 7 | TRUE | 29 | 1 | TRUE |
| ALA | 40 | 24 | 1 | FALSE | 27 | 0 | FALSE | 32 | 0 | FALSE |
| ALA | 41 | 24 | 1 | FALSE | 27 | 0 | FALSE | 33 | 0 | FALSE |
| MET | 42 | <13 | 6 | TRUE | <13 | 5 | TRUE | 24 | 0 | TRUE |
| ALA | 43 | 25 | 4 | TRUE | 26 | 7 | TRUE | <13 | 6 | TRUE |
| GLU | 44 | <12 | 4 | TRUE | 12 | 5 | TRUE | <12 | 3 | TRUE |
| LEU | 45 | 14 | 3 | TRUE | 14 | 1 | TRUE | 18 | 0 | TRUE |
| ASN | 46 | <14 | 4 | TRUE | <14 | 5 | TRUE | <14 | 7 | TRUE |
| TYR | 47 | 16 | 10 | TRUE | 13 | 10 | TRUE | 28 | 0 | TRUE |
| ILE | 48 | 19 | 2 | FALSE | 20 | 3 | FALSE | 24 | 1 | FALSE |
| PRO | 49 | | | | | | | | | |
| ASN | 50 | <14 | 7 | TRUE | <14 | 4 | TRUE | 24 | 3 | TRUE |
| ARG | 51 | <15 | 4 | TRUE | <15 | 4 | TRUE | 20 | 1 | TRUE |
| VAL | 52 | <10 | 1 | FALSE | <10 | 1 | FALSE | 15 | 0 | FALSE |
| ALA | 53 | <12 | 1 | FALSE | <12 | 1 | FALSE | 17 | 0 | FALSE |
| GLN | 54 | <13 | 4 | TRUE | <13 | 4 | TRUE | 26 | 0 | TRUE |
| GLN | 55 | <14 | 4 | TRUE | <14 | 4 | TRUE | <14 | 4 | TRUE |
| LEU | 56 | <11 | 3 | TRUE | <11 | 3 | TRUE | <11 | 3 | TRUE |
| ALA | 57 | <11 | 3 | TRUE | <12 | 3 | TRUE | <12 | 4 | TRUE |
| GLY | 58 | 13 | 9 | TRUE | 9 | 5 | TRUE | 26 | 0 | TRUE |

| | | | | | | | | | | |
|---|---|---|---|---|---|---|---|---|---|---|
| LYS | 59 | <13 | 4 | TRUE | <13 | 4 | TRUE | 20 | 1 | TRUE |
| GLN | 60 | <14 | 4 | TRUE | <14 | 4 | TRUE | 14 | 5 | TRUE |
| SER | 61 | <16 | 2 | FALSE | 19 | 3 | FALSE | 24 | 3 | FALSE |
| LEU | 62 | <11 | 2 | FALSE | 15 | 3 | FALSE | 19 | 3 | FALSE |
| LEU | 63 | 14 | 10 | TRUE | 12 | 9 | TRUE | <8 | 2 | TRUE |
| ILE | 64 | 11 | 8 | TRUE | 10 | 6 | TRUE | 24 | 0 | TRUE |
| GLY | 65 | 40 | 0 | TRUE | >49 | 7 | TRUE | 38 | 0 | TRUE |
| VAL | 66 | >47 | 4 | FALSE | >47 | 2 | FALSE | >47 | 3 | FALSE |
| ALA | 67 | >49 | 4 | FALSE | >49 | 2 | FALSE | >49 | 3 | FALSE |
| THR | 68 | 10 | 2 | FALSE | 11 | 2 | FALSE | 16 | 0 | FALSE |
| SER | 69 | 14 | 2 | FALSE | 14 | 2 | FALSE | 20 | 0 | FALSE |
| SER | 70 | 21 | 1 | TRUE | 18 | 7 | TRUE | 23 | 1 | TRUE |
| LEU | 71 | 36 | 0 | TRUE | 44 | 11 | TRUE | 36 | 0 | TRUE |
| ALA | 72 | 14 | 3 | TRUE | <12 | 5 | TRUE | 25 | 1 | TRUE |
| LEU | 73 | <9 | 3 | TRUE | 16 | 0 | TRUE | <10 | 5 | TRUE |
| HIS | 74 | 28 | 2 | TRUE | 30 | 1 | TRUE | 31 | 1 | TRUE |
| ALA | 75 | <14 | 6 | TRUE | 22 | 0 | TRUE | <14 | 6 | TRUE |
| PRO | 76 | | | | | | | | | |
| SER | 77 | 23 | 3 | TRUE | 29 | 1 | TRUE | 32 | 0 | TRUE |
| GLN | 78 | 23 | 3 | TRUE | 28 | 0 | TRUE | 22 | 0 | TRUE |
| ILE | 79 | 22 | 1 | FALSE | 51 | 4 | FALSE | 32 | 1 | FALSE |
| VAL | 80 | 20 | 1 | FALSE | 49 | 4 | FALSE | 30 | 1 | FALSE |
| ALA | 81 | 35 | 1 | TRUE | >49 | 7 | TRUE | >49 | 19 | TRUE |
| ALA | 82 | 30 | 0 | TRUE | 33 | 1 | TRUE | 35 | 3 | TRUE |
| ILE | 83 | 23 | 4 | TRUE | 16 | 0 | TRUE | 25 | 4 | TRUE |
| LYS | 84 | 34 | 3 | TRUE | 30 | 2 | TRUE | >49 | 15 | TRUE |
| SER | 85 | 21 | 3 | FALSE | 17 | 3 | FALSE | 34 | 1 | FALSE |
| ARG | 86 | 20 | 3 | FALSE | 17 | 3 | FALSE | 33 | 1 | FALSE |
| ALA | 87 | 26 | 0 | FALSE | 29 | 1 | FALSE | 21 | 2 | FALSE |
| ASP | 88 | 25 | 0 | FALSE | 28 | 1 | FALSE | 20 | 2 | FALSE |
| GLN | 89 | 24 | 0 | FALSE | 27 | 1 | FALSE | 19 | 2 | FALSE |

| | | | | | | | | | | |
|---|---|---|---|---|---|---|---|---|---|---|
| LEU | 90 | 29 | 1 | TRUE | 30 | 3 | TRUE | 31 | 1 | TRUE |
| GLY | 91 | 27 | 2 | TRUE | 26 | 6 | TRUE | 31 | 2 | TRUE |
| ALA | 92 | 32 | 1 | TRUE | 37 | 1 | TRUE | 46 | 13 | TRUE |
| SER | 93 | 30 | 0 | TRUE | 30 | 2 | TRUE | 28 | 2 | TRUE |
| VAL | 94 | <10 | 3 | TRUE | <10 | 3 | TRUE | <10 | 5 | TRUE |
| VAL | 95 | 29 | 1 | TRUE | 28 | 4 | TRUE | 30 | 0 | TRUE |
| VAL | 96 | 37 | 3 | FALSE | 31 | 4 | FALSE | 29 | 4 | FALSE |
| SER | 97 | 43 | 3 | FALSE | 37 | 4 | FALSE | 35 | 4 | FALSE |
| MET | 98 | 43 | 3 | FALSE | 38 | 4 | FALSE | 35 | 4 | FALSE |
| VAL | 99 | | | | | | | | | |
| GLU | 100 | <11 | 3 | TRUE | <11 | 3 | TRUE | <11 | 3 | TRUE |
| ARG | 101 | <12 | 1 | FALSE | <12 | 2 | FALSE | <12 | 1 | FALSE |
| SER | 102 | <16 | 1 | FALSE | <16 | 2 | FALSE | <16 | 1 | FALSE |
| GLY | 103 | <14 | 1 | FALSE | <14 | 1 | FALSE | 12 | 1 | FALSE |
| VAL | 104 | <10 | 1 | FALSE | <10 | 1 | FALSE | 8 | 1 | FALSE |
| GLU | 105 | <11 | 1 | FALSE | <11 | 1 | FALSE | 9 | 1 | FALSE |
| ALA | 106 | <12 | 4 | TRUE | <12 | 5 | TRUE | 13 | 5 | TRUE |
| CYS | 107 | <16 | 5 | TRUE | <16 | 7 | TRUE | <16 | 5 | TRUE |
| LYS | 108 | 30 | 0 | TRUE | 28 | 3 | TRUE | 30 | 0 | TRUE |
| ALA | 109 | 26 | 0 | TRUE | 23 | 3 | TRUE | 22 | 0 | TRUE |
| ALA | 110 | 34 | 2 | TRUE | 31 | 1 | TRUE | 33 | 4 | TRUE |
| VAL | 111 | 27 | 1 | TRUE | 25 | 1 | TRUE | 28 | 4 | TRUE |
| HIS | 112 | 20 | 0 | TRUE | 22 | 3 | TRUE | 26 | 0 | TRUE |
| ASN | 113 | 34 | 1 | TRUE | 33 | 1 | TRUE | 41 | 1 | TRUE |
| LEU | 114 | >49 | 6 | TRUE | 37 | 2 | TRUE | >49 | 7 | TRUE |
| LEU | 115 | >46 | 18 | TRUE | 34 | 1 | TRUE | >46 | 18 | TRUE |
| ALA | 116 | 25 | 1 | TRUE | 22 | 4 | TRUE | 29 | 1 | TRUE |
| GLN | 117 | 27 | 1 | TRUE | 28 | 1 | TRUE | 34 | 0 | TRUE |
| ARG | 118 | 19 | 1 | TRUE | 18 | 1 | TRUE | 22 | 0 | TRUE |
| VAL | 119 | 22 | 3 | FALSE | 44 | 3 | FALSE | 29 | 2 | FALSE |
| SER | 120 | 26 | 3 | FALSE | 48 | 3 | FALSE | 33 | 2 | FALSE |

| | | | | | | | | | | |
|---|---|---|---|---|---|---|---|---|---|---|
| GLY | 121 | 33 | 6 | TRUE | >52 | 6 | TRUE | 38 | 3 | TRUE |
| LEU | 122 | 44 | 16 | TRUE | 29 | 3 | TRUE | 34 | 4 | TRUE |
| ILE | 123 | 19 | 2 | TRUE | 15 | 1 | TRUE | 26 | 0 | TRUE |
| ILE | 124 | >45 | 21 | TRUE | 30 | 1 | TRUE | >45 | 21 | TRUE |
| ASN | 125 | 20 | 1 | TRUE | 28 | 0 | TRUE | 23 | 2 | TRUE |
| TYR | 126 | >50 | 5 | FALSE | >50 | 2 | FALSE | >50 | 2 | FALSE |
| PRO | 127 | | | | | | | | | |
| LEU | 128 | >46 | 6 | TRUE | >46 | 6 | TRUE | >46 | 6 | TRUE |
| ASP | 129 | >50 | 5 | TRUE | >50 | 5 | TRUE | >50 | 5 | TRUE |
| ASP | 130 | 34 | 6 | TRUE | >50 | 7 | TRUE | 32 | 1 | TRUE |
| GLN | 131 | 17 | 1 | TRUE | 16 | 0 | TRUE | 14 | 3 | TRUE |
| ASP | 132 | 20 | 0 | TRUE | 20 | 0 | TRUE | 20 | 0 | TRUE |
| ALA | 133 | 34 | 4 | TRUE | 32 | 1 | TRUE | 33 | 2 | TRUE |
| ILE | 134 | >46 | 20 | TRUE | 35 | 4 | TRUE | >46 | 18 | TRUE |
| ALA | 135 | 38 | 5 | TRUE | 37 | 2 | TRUE | >49 | 21 | TRUE |
| VAL | 136 | 25 | 1 | FALSE | 25 | 0 | FALSE | 25 | 0 | FALSE |
| GLU | 137 | 27 | 1 | FALSE | 27 | 0 | FALSE | 28 | 0 | FALSE |
| ALA | 138 | 25 | 2 | TRUE | 26 | 0 | TRUE | 27 | 1 | TRUE |
| ALA | 139 | >50 | 22 | TRUE | >50 | 14 | TRUE | >50 | 13 | TRUE |
| CYS | 140 | 22 | 2 | TRUE | 23 | 2 | TRUE | 22 | 0 | TRUE |
| THR | 141 | <15 | 7 | TRUE | <1 | 5 | TRUE | 33 | 11 | TRUE |
| ASN | 142 | 49 | 30 | TRUE | 48 | 27 | TRUE | <17 | 5 | TRUE |
| VAL | 143 | 12 | 4 | TRUE | <11 | 5 | TRUE | 16 | 0 | TRUE |
| PRO | 144 | | | | | | | | | |
| ALA | 145 | 48 | 3 | FALSE | 48 | 6 | FALSE | 44 | 6 | FALSE |
| LEU | 146 | 46 | 3 | FALSE | 46 | 6 | FALSE | 42 | 6 | FALSE |
| PHE | 147 | >48 | 7 | TRUE | 30 | 3 | TRUE | 31 | 4 | TRUE |
| LEU | 148 | >47 | 21 | TRUE | >47 | 6 | TRUE | >47 | 6 | TRUE |
| ASP | 149 | 28 | 0 | TRUE | 33 | 1 | TRUE | 31 | 1 | TRUE |
| VAL | 150 | 33 | 1 | TRUE | >45 | 11 | TRUE | >45 | 12 | TRUE |
| SER | 151 | <14 | 4 | TRUE | <14 | 6 | TRUE | <14 | 6 | TRUE |

| | | | | | | | | | | |
|---|---|---|---|---|---|---|---|---|---|---|
| ASP | 152 | 16 | 8 | TRUE | 22 | 8 | TRUE | 20 | 7 | TRUE |
| GLN | 153 | 22 | 8 | TRUE | 22 | 5 | TRUE | 22 | 8 | TRUE |
| THR | 154 | 20 | 0 | TRUE | 20 | 2 | TRUE | 20 | 0 | TRUE |
| PRO | 155 | | | | | | | | | |
| ILE | 156 | 15 | 3 | FALSE | 21 | 2 | FALSE | 20 | 2 | FALSE |
| ASN | 157 | 22 | 3 | FALSE | 27 | 2 | FALSE | 27 | 2 | FALSE |
| SER | 158 | 33 | 2 | TRUE | 33 | 1 | TRUE | 32 | 1 | TRUE |
| ILE | 159 | 30 | 0 | TRUE | >48 | 7 | TRUE | 40 | 2 | TRUE |
| ILE | 160 | 25 | 1 | TRUE | 33 | 1 | TRUE | 31 | 1 | TRUE |
| PHE | 161 | 30 | 0 | TRUE | >48 | 12 | TRUE | 29 | 1 | TRUE |
| SER | 162 | 49 | 5 | FALSE | 33 | 3 | FALSE | 52 | 5 | FALSE |
| HIS | 163 | 49 | 5 | FALSE | 33 | 3 | FALSE | 51 | 5 | FALSE |
| GLU | 164 | 35 | 1 | TRUE | 51 | 23 | TRUE | >51 | 18 | TRUE |
| ASP | 165 | 18 | 0 | TRUE | 21 | 7 | TRUE | 18 | 0 | TRUE |
| GLY | 166 | 18 | 0 | TRUE | 33 | 22 | TRUE | 17 | 1 | TRUE |
| THR | 167 | 30 | 1 | TRUE | >51 | 7 | TRUE | 44 | 16 | TRUE |
| ARG | 168 | 37 | 0 | FALSE | >52 | 2 | FALSE | 49 | 8 | FALSE |
| LEU | 169 | 34 | 0 | FALSE | >48 | 2 | FALSE | 45 | 8 | FALSE |
| GLY | 170 | >49 | 12 | TRUE | >49 | 8 | TRUE | >49 | 9 | TRUE |
| VAL | 171 | >47 | 8 | TRUE | >47 | 7 | TRUE | >47 | 8 | TRUE |
| GLU | 172 | 29 | 0 | FALSE | 31 | 1 | FALSE | 31 | 1 | FALSE |
| HIS | 173 | 30 | 0 | FALSE | 31 | 1 | FALSE | 32 | 1 | FALSE |
| LEU | 174 | >48 | 7 | TRUE | >48 | 7 | TRUE | >48 | 7 | TRUE |
| VAL | 175 | >45 | 12 | TRUE | >45 | 20 | TRUE | >45 | 16 | TRUE |
| ALA | 176 | >49 | 12 | TRUE | 40 | 1 | TRUE | >49 | 14 | TRUE |
| LEU | 177 | 16 | 0 | TRUE | 16 | 0 | TRUE | 16 | 0 | TRUE |
| GLY | 178 | 32 | 0 | TRUE | 34 | 0 | TRUE | 33 | 2 | TRUE |
| HIS | 179 | 39 | 1 | TRUE | 38 | 0 | TRUE | 35 | 4 | TRUE |
| GLN | 180 | 30 | 0 | TRUE | 30 | 0 | TRUE | 52 | 22 | TRUE |
| GLN | 181 | <14 | 4 | TRUE | 10 | 5 | TRUE | 45 | 19 | TRUE |
| ILE | 182 | 47 | 28 | TRUE | >47 | 7 | TRUE | <10 | 3 | TRUE |

| | | | | | | | | | | |
|---|---|---|---|---|---|---|---|---|---|---|
| ALA | 183 | >49 | 6 | TRUE | >49 | 6 | TRUE | >49 | 22 | TRUE |
| LEU | 184 | >47 | 7 | TRUE | >47 | 11 | TRUE | >47 | 11 | TRUE |
| LEU | 185 | 34 | 0 | TRUE | 36 | 1 | TRUE | >46 | 16 | TRUE |
| ALA | 186 | 31 | 2 | TRUE | 36 | 2 | TRUE | 33 | 2 | TRUE |
| GLY | 187 | 28 | 5 | TRUE | 30 | 5 | TRUE | 30 | 7 | TRUE |
| PRO | 188 | | | | | | | | | |
| LEU | 189 | 17 | 6 | TRUE | 18 | 6 | TRUE | 17 | 3 | TRUE |
| SER | 190 | 23 | 3 | FALSE | 19 | 3 | FALSE | 19 | 4 | FALSE |
| SER | 191 | 26 | 3 | FALSE | 22 | 3 | FALSE | 22 | 4 | FALSE |
| VAL | 192 | 15 | 1 | TRUE | 20 | 10 | TRUE | 16 | 1 | TRUE |
| SER | 193 | <14 | 5 | TRUE | 29 | 1 | TRUE | <14 | 5 | TRUE |
| ALA | 194 | 20 | 0 | TRUE | 26 | 3 | TRUE | 20 | 0 | TRUE |
| ARG | 195 | 32 | 1 | TRUE | 50 | 18 | TRUE | 32 | 1 | TRUE |
| LEU | 196 | 29 | 1 | TRUE | 35 | 1 | TRUE | 32 | 1 | TRUE |
| ARG | 197 | >49 | 17 | TRUE | >49 | 7 | TRUE | 43 | 8 | TRUE |
| LEU | 198 | 30 | 4 | TRUE | 32 | 6 | TRUE | 28 | 4 | TRUE |
| ALA | 199 | 36 | 19 | TRUE | >49 | 7 | TRUE | >49 | 7 | TRUE |
| GLY | 200 | >49 | 21 | TRUE | >50 | 20 | TRUE | >50 | 22 | TRUE |
| TRP | 201 | >50 | 3 | FALSE | 57 | 5 | FALSE | >49 | 3 | FALSE |
| HIS | 202 | >50 | 3 | FALSE | 58 | 5 | FALSE | >50 | 3 | FALSE |
| LYS | 203 | 34 | 2 | TRUE | 36 | 0 | TRUE | 33 | 1 | TRUE |
| TYR | 204 | 27 | 10 | TRUE | 36 | 18 | TRUE | 37 | 1 | TRUE |
| LEU | 205 | 26 | 8 | TRUE | 34 | 23 | TRUE | 21 | 8 | TRUE |
| THR | 206 | 29 | 2 | TRUE | 29 | 3 | TRUE | 29 | 1 | TRUE |
| ARG | 207 | 15 | 8 | TRUE | 23 | 6 | TRUE | 18 | 1 | TRUE |
| ASN | 208 | 29 | 5 | TRUE | 21 | 1 | TRUE | 29 | 5 | TRUE |
| GLN | 209 | 26 | 3 | TRUE | 26 | 4 | TRUE | 28 | 3 | TRUE |
| ILE | 210 | 32 | 0 | TRUE | 32 | 0 | TRUE | 35 | 1 | TRUE |
| GLN | 211 | 24 | 0 | FALSE | 25 | 0 | FALSE | 24 | 0 | FALSE |
| PRO | 212 | | | | | | | | | |
| ILE | 213 | 19 | 0 | FALSE | 20 | 0 | FALSE | 19 | 0 | FALSE |

| | | | | | | | | | | |
|---|---|---|---|---|---|---|---|---|---|---|
| ALA | 214 | 23 | 0 | FALSE | 24 | 0 | FALSE | 23 | 0 | FALSE |
| GLU | 215 | 13 | 5 | TRUE | <12 | 5 | TRUE | 16 | 0 | TRUE |
| ARG | 216 | 30 | 0 | TRUE | 30 | 0 | TRUE | 31 | 1 | TRUE |
| GLU | 217 | 14 | 9 | TRUE | <13 | 4 | TRUE | <13 | 4 | TRUE |
| GLY | 218 | 45 | 6 | FALSE | 27 | 1 | FALSE | 31 | 3 | FALSE |
| ASP | 219 | 47 | 6 | FALSE | 30 | 1 | FALSE | 34 | 3 | FALSE |
| TRP | 220 | 39 | 18 | TRUE | 34 | 2 | TRUE | 31 | 3 | TRUE |
| SER | 221 | 21 | 14 | TRUE | <14 | 6 | TRUE | 24 | 8 | TRUE |
| ALA | 222 | 25 | 6 | TRUE | >52 | 6 | TRUE | 39 | 4 | TRUE |
| MET | 223 | 20 | 4 | TRUE | 19 | 4 | TRUE | 20 | 0 | TRUE |
| SER | 224 | 26 | 3 | TRUE | 24 | 4 | TRUE | 28 | 0 | TRUE |
| GLY | 225 | 42 | 3 | TRUE | 40 | 1 | TRUE | 38 | 0 | TRUE |
| PHE | 226 | >50 | 15 | TRUE | 38 | 3 | TRUE | >50 | 13 | TRUE |
| GLN | 227 | 39 | 1 | TRUE | >51 | 18 | TRUE | 40 | 2 | TRUE |
| GLN | 228 | 36 | 1 | TRUE | 40 | 3 | TRUE | 36 | 1 | TRUE |
| THR | 229 | >51 | 6 | TRUE | >51 | 5 | TRUE | >51 | 6 | TRUE |
| MET | 230 | 41 | 4 | TRUE | 36 | 3 | TRUE | >51 | 14 | TRUE |
| GLN | 231 | 32 | 0 | TRUE | 33 | 2 | TRUE | 32 | 0 | TRUE |
| MET | 232 | 38 | 0 | TRUE | 36 | 1 | TRUE | 36 | 0 | TRUE |
| LEU | 233 | 33 | 1 | TRUE | 31 | 1 | TRUE | 34 | 0 | TRUE |
| ASN | 234 | 37 | 1 | TRUE | 41 | 2 | TRUE | 38 | 1 | TRUE |
| GLU | 235 | 28 | 0 | TRUE | 28 | 0 | TRUE | 28 | 0 | TRUE |
| GLY | 236 | 18 | 0 | TRUE | 18 | 0 | TRUE | 16 | 0 | TRUE |
| ILE | 237 | 20 | 0 | TRUE | 20 | 0 | TRUE | 20 | 0 | TRUE |
| VAL | 238 | >45 | 7 | TRUE | >45 | 20 | TRUE | >45 | 22 | TRUE |
| PRO | 239 | | | | | | | | | |
| THR | 240 | >48 | 6 | TRUE | >48 | 6 | TRUE | >48 | 6 | TRUE |
| ALA | 241 | 22 | 0 | TRUE | 21 | 1 | TRUE | 22 | 0 | TRUE |
| MET | 242 | 35 | 2 | TRUE | 34 | 0 | TRUE | 35 | 2 | TRUE |
| LEU | 243 | 34 | 3 | TRUE | 36 | 2 | TRUE | 34 | 2 | TRUE |
| VAL | 244 | 48 | 18 | TRUE | 32 | 1 | TRUE | >45 | 13 | TRUE |

| | | | | | | | | | | |
|---|---|---|---|---|---|---|---|---|---|---|
| ALA | 245 | >49 | 6 | TRUE | 42 | 1 | TRUE | >49 | 19 | TRUE |
| ASN | 246 | 30 | 5 | TRUE | 35 | 6 | TRUE | 36 | 1 | TRUE |
| ASP | 247 | 26 | 4 | TRUE | 33 | 5 | TRUE | 28 | 0 | TRUE |
| GLN | 248 | >50 | 24 | TRUE | 49 | 19 | TRUE | >50 | 5 | TRUE |
| MET | 249 | >51 | 5 | TRUE | >51 | 13 | TRUE | >51 | 24 | TRUE |
| ALA | 250 | >51 | 5 | TRUE | >51 | 5 | TRUE | >51 | 5 | TRUE |
| LEU | 251 | >47 | 5 | TRUE | >47 | 6 | TRUE | >47 | 6 | TRUE |
| GLY | 252 | >49 | 6 | TRUE | >49 | 6 | TRUE | >49 | 6 | TRUE |
| ALA | 253 | 44 | 3 | TRUE | 41 | 2 | TRUE | >51 | 21 | TRUE |
| MET | 254 | 48 | 12 | TRUE | >50 | 6 | TRUE | >50 | 20 | TRUE |
| ARG | 255 | 48 | 18 | TRUE | 48 | 19 | TRUE | >51 | 20 | TRUE |
| ALA | 256 | 37 | 1 | TRUE | 45 | 16 | TRUE | 36 | 1 | TRUE |
| ILE | 257 | >46 | 13 | TRUE | >46 | 14 | TRUE | >46 | 17 | TRUE |
| THR | 258 | >49 | 13 | TRUE | >49 | 7 | TRUE | >49 | 12 | TRUE |
| GLU | 259 | 29 | 1 | TRUE | 28 | 0 | TRUE | 28 | 0 | TRUE |
| SER | 260 | <14 | 4 | TRUE | <14 | 5 | TRUE | <14 | 4 | TRUE |
| GLY | 261 | >52 | 13 | TRUE | 39 | 2 | TRUE | 49 | 14 | TRUE |
| LEU | 262 | 33 | 1 | TRUE | 32 | 0 | TRUE | 34 | 1 | TRUE |
| ARG | 263 | 33 | 2 | TRUE | 34 | 1 | TRUE | 35 | 1 | TRUE |
| VAL | 264 | 29 | 1 | TRUE | 28 | 2 | TRUE | 30 | 2 | TRUE |
| GLY | 265 | 14 | 1 | FALSE | 16 | 1 | FALSE | 16 | 1 | FALSE |
| ALA | 266 | 16 | 1 | FALSE | 17 | 1 | FALSE | 17 | 1 | FALSE |
| ASP | 267 | 16 | 1 | FALSE | 17 | 1 | FALSE | 17 | 1 | FALSE |
| ILE | 268 | 26 | 0 | TRUE | 25 | 1 | TRUE | 26 | 2 | TRUE |
| SER | 269 | 28 | 5 | TRUE | 29 | 2 | TRUE | 30 | 0 | TRUE |
| VAL | 270 | >48 | 1 | FALSE | >48 | 2 | FALSE | >48 | 2 | FALSE |
| VAL | 271 | >46 | 1 | FALSE | >46 | 2 | FALSE | >46 | 2 | FALSE |
| GLY | 272 | >49 | 1 | FALSE | >49 | 2 | FALSE | >49 | 2 | FALSE |
| TYR | 273 | 35 | 1 | TRUE | 38 | 1 | TRUE | 38 | 0 | TRUE |
| ASP | 274 | 47 | 15 | TRUE | >51 | 6 | TRUE | >51 | 11 | TRUE |
| ASP | 275 | 31 | 4 | TRUE | 46 | 23 | TRUE | 18 | 1 | TRUE |

| | | | | | | | | | | |
|---|---|---|---|---|---|---|---|---|---|---|
| THR | 276 | 15 | 1 | TRUE | 39 | 28 | TRUE | >49 | 6 | TRUE |
| GLU | 277 | 17 | 1 | TRUE | <13 | 6 | TRUE | <13 | 5 | TRUE |
| ASP | 278 | 33 | 3 | TRUE | 23 | 6 | TRUE | 32 | 1 | TRUE |
| SER | 279 | >51 | 6 | TRUE | >51 | 13 | TRUE | >51 | 5 | TRUE |
| SER | 280 | 27 | 1 | TRUE | 26 | 3 | TRUE | 27 | 1 | TRUE |
| CYS | 281 | 45 | 14 | TRUE | 42 | 18 | TRUE | >55 | 8 | TRUE |
| TYR | 282 | 30 | 1 | TRUE | 43 | 21 | TRUE | >52 | 5 | TRUE |
| ILE | 283 | 32 | 6 | TRUE | 41 | 19 | TRUE | >46 | 6 | TRUE |
| PRO | 284 | | | | | | | | | |
| PRO | 285 | | | | | | | | | |
| LEU | 286 | 29 | 0 | FALSE | 38 | 3 | FALSE | 40 | 2 | FALSE |
| THR | 287 | 32 | 0 | FALSE | 41 | 3 | FALSE | 43 | 2 | FALSE |
| THR | 288 | 35 | 0 | FALSE | 43 | 3 | FALSE | 46 | 2 | FALSE |
| ILE | 289 | 35 | 1 | FALSE | >47 | 4 | FALSE | >47 | 2 | FALSE |
| LYS | 290 | 37 | 1 | FALSE | >49 | 4 | FALSE | >49 | 2 | FALSE |
| GLN | 291 | 29 | 1 | TRUE | 38 | 0 | TRUE | 34 | 0 | TRUE |
| ASP | 292 | 20 | 1 | TRUE | 21 | 4 | TRUE | 21 | 2 | TRUE |
| PHE | 293 | 17 | 1 | TRUE | 18 | 4 | TRUE | 18 | 3 | TRUE |
| ARG | 294 | 20 | 0 | TRUE | 20 | 0 | TRUE | 20 | 0 | TRUE |
| LEU | 295 | 26 | 0 | TRUE | 24 | 0 | TRUE | 26 | 0 | TRUE |
| LEU | 296 | 31 | 1 | TRUE | 34 | 0 | TRUE | 34 | 0 | TRUE |
| GLY | 297 | 47 | 14 | TRUE | >49 | 12 | TRUE | >49 | 13 | TRUE |
| GLN | 298 | 36 | 1 | TRUE | 39 | 1 | TRUE | 38 | 3 | TRUE |
| THR | 299 | 28 | 0 | TRUE | 43 | 18 | TRUE | 45 | 18 | TRUE |
| SER | 300 | 44 | 2 | TRUE | >53 | 7 | TRUE | 45 | 17 | TRUE |
| VAL | 301 | 38 | 3 | TRUE | 31 | 7 | TRUE | 47 | 15 | TRUE |
| ASP | 302 | 32 | 0 | TRUE | 36 | 1 | TRUE | 30 | 3 | TRUE |
| ARG | 303 | >50 | 6 | TRUE | >50 | 6 | TRUE | >50 | 6 | TRUE |
| LEU | 304 | >48 | 5 | TRUE | >48 | 6 | TRUE | >48 | 5 | TRUE |
| LEU | 305 | >46 | 6 | TRUE | >46 | 7 | TRUE | >46 | 6 | TRUE |
| GLN | 306 | 32 | 0 | TRUE | 34 | 1 | TRUE | 33 | 1 | TRUE |

| | | | | | | | | | | |
|---|---|---|---|---|---|---|---|---|---|---|
| LEU | 307 | 16 | 0 | TRUE | 16 | 1 | TRUE | 17 | 1 | TRUE |
| SER | 308 | <14 | 5 | TRUE | <14 | 4 | TRUE | <14 | 6 | TRUE |
| GLN | 309 | <15 | 5 | TRUE | <15 | 8 | TRUE | 17 | 1 | TRUE |
| GLY | 310 | 18 | 2 | FALSE | 24 | 2 | FALSE | 23 | 2 | FALSE |
| GLN | 311 | 18 | 2 | FALSE | 25 | 2 | FALSE | 24 | 2 | FALSE |
| ALA | 312 | <14 | 4 | TRUE | <14 | 4 | TRUE | <14 | 4 | TRUE |
| VAL | 313 | <9 | 1 | FALSE | <9 | 1 | FALSE | <9 | 1 | FALSE |
| LYS | 314 | <12 | 1 | FALSE | <12 | 1 | FALSE | <12 | 1 | FALSE |
| GLY | 315 | <13 | 1 | FALSE | <13 | 1 | FALSE | <13 | 1 | FALSE |
| ASN | 316 | <16 | 4 | TRUE | <16 | 4 | TRUE | <16 | 5 | TRUE |
| GLN | 317 | <15 | 4 | TRUE | <15 | 4 | TRUE | <15 | 4 | TRUE |
| LEU | 318 | <11 | 3 | TRUE | <11 | 3 | TRUE | <11 | 4 | TRUE |
| LEU | 319 | <8 | 2 | TRUE | 14 | 0 | TRUE | 14 | 0 | TRUE |
| PRO | 320 | | | | | | | | | |
| VAL | 321 | 25 | 0 | FALSE | 38 | 2 | FALSE | 27 | 1 | FALSE |
| SER | 322 | 31 | 0 | FALSE | 45 | 2 | FALSE | 33 | 1 | FALSE |
| LEU | 323 | 28 | 0 | FALSE | 42 | 2 | FALSE | 31 | 1 | FALSE |
| VAL | 324 | | | | | | | | | |
| LYS | 325 | | | | | | | | | |
| ARG | 326 | 13 | 0 | FALSE | 16 | 0 | FALSE | 13 | 0 | FALSE |
| LYS | 327 | 13 | 0 | FALSE | 15 | 0 | FALSE | 13 | 0 | FALSE |
| THR | 328 | 12 | 0 | FALSE | 15 | 0 | FALSE | 12 | 0 | FALSE |
| THR | 329 | 13 | 0 | FALSE | 15 | 0 | FALSE | 13 | 0 | FALSE |
| LEU | 330 | 10 | 0 | FALSE | 12 | 0 | FALSE | 10 | 0 | FALSE |
| ALA | 331 | 11 | 0 | FALSE | 13 | 0 | FALSE | 11 | 0 | FALSE |
| PRO | 332 | | | | | | | | | |
| ASN | 333 | | | | | | | | | |
| THR | 334 | | | | | | | | | |
| GLN | 335 | | | | | | | | | |
| THR | 336 | | | | | | | | | |
| ALA | 337 | | | | | | | | | |

| | | | | | | | | | | |
|---|---|---|---|---|---|---|---|---|---|---|
| SER | 338 | | | | | | | | | |
| PRO | 339 | | | | | | | | | |
| ARG | 340 | | | | | | | | | |
| ALA | 341 | | | | | | | | | |
| LEU | 342 | | | | | | | | | |
| ALA | 343 | | | | | | | | | |
| ASP | 344 | 37 | 4 | FALSE | 17 | 1 | FALSE | 37 | 3 | FALSE |
| SER | 345 | 37 | 4 | FALSE | 17 | 1 | FALSE | 37 | 3 | FALSE |
| LEU | 346 | 35 | 4 | FALSE | 15 | 1 | FALSE | 35 | 3 | FALSE |
| MET | 347 | 14 | 2 | FALSE | 22 | 2 | FALSE | 18 | 2 | FALSE |
| GLN | 348 | 16 | 2 | FALSE | 25 | 2 | FALSE | 20 | 2 | FALSE |
| LEU | 349 | 13 | 2 | FALSE | 21 | 2 | FALSE | 17 | 2 | FALSE |
| ALA | 350 | 36 | 7 | FALSE | 47 | 6 | FALSE | 29 | 8 | FALSE |
| ARG | 351 | 37 | 7 | FALSE | 48 | 6 | FALSE | 30 | 8 | FALSE |
| GLN | 352 | >53 | 12 | TRUE | 49 | 14 | TRUE | 52 | 20 | TRUE |
| VAL | 353 | 16 | 1 | TRUE | 25 | 18 | TRUE | 26 | 22 | TRUE |
| SER | 354 | >51 | 22 | TRUE | >51 | 20 | TRUE | 44 | 22 | TRUE |
| ARG | 355 | 41 | 1 | FALSE | 41 | 1 | FALSE | 40 | 2 | FALSE |
| LEU | 356 | 37 | 1 | FALSE | 37 | 1 | FALSE | 35 | 2 | FALSE |
| GLU | 357 | 37 | 1 | FALSE | 37 | 1 | FALSE | 36 | 2 | FALSE |
| SER | 358 | 40 | 1 | FALSE | 40 | 1 | FALSE | 39 | 2 | FALSE |
| GLY | 359 | 41 | 1 | FALSE | 40 | 1 | FALSE | 39 | 2 | FALSE |
| GLN | 360 | 25 | 2 | TRUE | 33 | 22 | TRUE | 23 | 2 | TRUE |

**Table S9. $\Delta G_{op}$ values for GalR**

All values in kJ/mol.

| Res. name | Res. number | Apo | Std Dev | Single resolved | GAL | Std Dev | Single resolved | *galO* | Std Dev | Single resolved |
|---|---|---|---|---|---|---|---|---|---|---|
| MET | 1 | | | | | | | | | |
| ALA | 2 | | | | | | | | | |
| THR | 3 | | | | | | | | | |
| ILE | 4 | | | | | | | | | |
| LYS | 5 | <12 | 3 | TRUE | <12 | 4 | TRUE | 15 | 7 | TRUE |
| ASP | 6 | 12 | 2 | FALSE | <15 | 1 | FALSE | 22 | 1 | FALSE |
| VAL | 7 | 6 | 2 | FALSE | <8 | 1 | FALSE | 16 | 1 | FALSE |
| ALA | 8 | 17 | 1 | TRUE | 16 | 0 | TRUE | 28 | 0 | TRUE |
| ARG | 9 | <14 | 6 | TRUE | <14 | 4 | TRUE | 27 | 1 | TRUE |
| LEU | 10 | <11 | 5 | TRUE | <11 | 3 | TRUE | 24 | 0 | TRUE |
| ALA | 11 | <12 | 1 | FALSE | <12 | 1 | FALSE | 24 | 0 | FALSE |
| GLY | 12 | <13 | 1 | FALSE | <13 | 1 | FALSE | 25 | 0 | FALSE |
| VAL | 13 | 9 | 1 | FALSE | <10 | 1 | FALSE | 24 | 0 | FALSE |
| SER | 14 | 13 | 1 | FALSE | <15 | 1 | FALSE | 28 | 0 | FALSE |
| VAL | 15 | <11 | 2 | TRUE | <11 | 2 | TRUE | 18 | 0 | TRUE |
| ALA | 16 | <13 | 3 | TRUE | <13 | 3 | TRUE | 26 | 0 | TRUE |
| THR | 17 | <13 | 4 | TRUE | <13 | 4 | TRUE | 21 | 14 | TRUE |
| VAL | 18 | <11 | 2 | TRUE | <11 | 3 | TRUE | 20 | 9 | TRUE |
| SER | 19 | <15 | 3 | TRUE | <15 | 4 | TRUE | 23 | 7 | TRUE |
| ARG | 20 | <15 | 4 | TRUE | <15 | 5 | TRUE | 30 | 0 | TRUE |
| VAL | 21 | <11 | 3 | TRUE | <11 | 3 | TRUE | 20 | 0 | TRUE |
| ILE | 22 | <9 | 2 | TRUE | <9 | 1 | TRUE | 8 | 4 | TRUE |
| ASN | 23 | 18 | 0 | FALSE | 15 | 0 | FALSE | 15 | 0 | FALSE |
| ASN | 24 | 21 | 0 | FALSE | 18 | 0 | FALSE | 18 | 0 | FALSE |
| SER | 25 | 20 | 0 | FALSE | 18 | 0 | FALSE | 18 | 0 | FALSE |
| PRO | 26 | | | | | | | | | |
| LYS | 27 | 15 | 0 | FALSE | 12 | 0 | FALSE | 12 | 0 | FALSE |

| | | | | | | | | | | |
|---|---|---|---|---|---|---|---|---|---|---|
| ALA | 28 | 17 | 0 | FALSE | 15 | 0 | FALSE | 15 | 0 | FALSE |
| SER | 29 | 18 | 0 | FALSE | 16 | 0 | FALSE | 16 | 0 | FALSE |
| GLU | 30 | 17 | 0 | FALSE | 15 | 0 | FALSE | 15 | 0 | FALSE |
| ALA | 31 | <12 | 3 | TRUE | <12 | 4 | TRUE | 12 | 5 | TRUE |
| SER | 32 | 27 | 0 | FALSE | 23 | 2 | FALSE | 22 | 1 | FALSE |
| ARG | 33 | 27 | 0 | FALSE | 23 | 2 | FALSE | 22 | 1 | FALSE |
| LEU | 34 | 26 | 22 | TRUE | <11 | 3 | TRUE | 18 | 1 | FALSE |
| ALA | 35 | 14 | 1 | FALSE | <12 | 1 | FALSE | <12 | 1 | FALSE |
| VAL | 36 | 12 | 1 | FALSE | <9 | 1 | FALSE | <9 | 1 | FALSE |
| HIS | 37 | 11 | 1 | FALSE | <13 | 1 | FALSE | 20 | 1 | FALSE |
| SER | 38 | 14 | 1 | FALSE | <17 | 1 | FALSE | 23 | 1 | FALSE |
| ALA | 39 | 13 | 1 | FALSE | <15 | 1 | FALSE | 22 | 1 | FALSE |
| MET | 40 | 31 | 1 | TRUE | 36 | 0 | TRUE | 34 | 0 | TRUE |
| GLU | 41 | 36 | 0 | TRUE | 37 | 1 | TRUE | 38 | 3 | TRUE |
| SER | 42 | >51 | 14 | TRUE | >51 | 8 | TRUE | >51 | 10 | TRUE |
| LEU | 43 | 15 | 1 | FALSE | 17 | 2 | FALSE | 22 | 1 | FALSE |
| SER | 44 | 17 | 1 | FALSE | 19 | 2 | FALSE | 25 | 1 | FALSE |
| TYR | 45 | 17 | 1 | FALSE | 18 | 2 | FALSE | 24 | 1 | FALSE |
| HIS | 46 | 12 | 0 | FALSE | 12 | 0 | FALSE | 19 | 1 | FALSE |
| PRO | 47 | | | | | | | | | |
| ASN | 48 | 13 | 0 | FALSE | 13 | 0 | FALSE | 19 | 1 | FALSE |
| ALA | 49 | 14 | 0 | FALSE | 13 | 0 | FALSE | 20 | 1 | FALSE |
| ASN | 50 | 15 | 0 | FALSE | 14 | 0 | FALSE | 21 | 1 | FALSE |
| ALA | 51 | 14 | 0 | FALSE | 13 | 0 | FALSE | 20 | 1 | FALSE |
| ARG | 52 | <14 | 1 | FALSE | <14 | 1 | FALSE | 19 | 0 | FALSE |
| ALA | 53 | <15 | 1 | FALSE | <15 | 1 | FALSE | 20 | 0 | FALSE |
| LEU | 54 | 12 | 0 | TRUE | <10 | 4 | TRUE | 21 | 2 | TRUE |
| ALA | 55 | 14 | 0 | TRUE | 14 | 0 | TRUE | 23 | 3 | TRUE |
| GLN | 56 | 17 | 1 | TRUE | 17 | 1 | TRUE | 19 | 1 | TRUE |
| GLN | 57 | 17 | 0 | FALSE | 12 | 1 | FALSE | 17 | 0 | FALSE |
| THR | 58 | 16 | 0 | FALSE | 11 | 1 | FALSE | 17 | 0 | FALSE |

| | | | | | | | | | | |
|---|---|---|---|---|---|---|---|---|---|---|
| THR | 59 | 16 | 0 | FALSE | 11 | 1 | FALSE | 17 | 0 | FALSE |
| GLU | 60 | 16 | 0 | FALSE | 11 | 1 | FALSE | 16 | 0 | FALSE |
| THR | 61 | 16 | 0 | TRUE | 16 | 0 | TRUE | 16 | 0 | TRUE |
| VAL | 62 | 23 | 1 | FALSE | 34 | 7 | FALSE | 29 | 1 | FALSE |
| GLY | 63 | 25 | 1 | FALSE | 36 | 7 | FALSE | 31 | 1 | FALSE |
| LEU | 64 | 41 | 22 | TRUE | 38 | 0 | TRUE | 34 | 7 | TRUE |
| VAL | 65 | 32 | 1 | TRUE | 29 | 6 | TRUE | >45 | 20 | TRUE |
| VAL | 66 | 32 | 1 | TRUE | >45 | 15 | TRUE | 29 | 1 | TRUE |
| GLY | 67 | 27 | 3 | TRUE | 31 | 1 | TRUE | 33 | 1 | TRUE |
| ASP | 68 | 20 | 2 | FALSE | 27 | 1 | FALSE | 25 | 1 | FALSE |
| VAL | 69 | 14 | 2 | FALSE | 20 | 1 | FALSE | 19 | 1 | FALSE |
| SER | 70 | 20 | 2 | FALSE | 27 | 1 | FALSE | 25 | 1 | FALSE |
| ASP | 71 | 21 | 6 | TRUE | 32 | 1 | TRUE | 37 | 1 | TRUE |
| PRO | 72 | | | | | | | | | |
| PHE | 73 | 26 | 2 | FALSE | 25 | 1 | FALSE | 18 | 2 | FALSE |
| PHE | 74 | 27 | 2 | FALSE | 27 | 1 | FALSE | 20 | 2 | FALSE |
| GLY | 75 | 32 | 0 | TRUE | 34 | 0 | TRUE | 33 | 1 | TRUE |
| ALA | 76 | 24 | 0 | TRUE | 24 | 0 | TRUE | 39 | 1 | TRUE |
| MET | 77 | 27 | 2 | TRUE | 34 | 3 | TRUE | 33 | 3 | TRUE |
| VAL | 78 | 30 | 6 | TRUE | 29 | 4 | TRUE | 32 | 4 | TRUE |
| LYS | 79 | 31 | 4 | TRUE | 27 | 1 | TRUE | 34 | 0 | TRUE |
| ALA | 80 | 37 | 1 | FALSE | 36 | 2 | FALSE | 39 | 1 | FALSE |
| VAL | 81 | 32 | 1 | FALSE | 32 | 2 | FALSE | 35 | 1 | FALSE |
| GLU | 82 | 35 | 2 | TRUE | >48 | 15 | TRUE | >48 | 14 | TRUE |
| GLN | 83 | 28 | 0 | TRUE | 32 | 0 | TRUE | 32 | 0 | TRUE |
| VAL | 84 | 29 | 1 | TRUE | 33 | 1 | TRUE | 32 | 0 | TRUE |
| ALA | 85 | 24 | 0 | TRUE | 26 | 1 | TRUE | 26 | 0 | TRUE |
| TYR | 86 | 34 | 0 | TRUE | 32 | 0 | TRUE | 29 | 1 | TRUE |
| HIS | 87 | 20 | 0 | TRUE | 18 | 0 | TRUE | 34 | 1 | TRUE |
| THR | 88 | 21 | 0 | FALSE | 22 | 0 | FALSE | 23 | 0 | FALSE |
| GLY | 89 | 21 | 0 | FALSE | 22 | 0 | FALSE | 23 | 0 | FALSE |

| | | | | | | | | | | |
|---|---|---|---|---|---|---|---|---|---|---|
| ASN | 90 | 23 | 0 | FALSE | 24 | 0 | FALSE | 26 | 0 | FALSE |
| PHE | 91 | 30 | 0 | TRUE | 34 | 0 | TRUE | 23 | 0 | FALSE |
| LEU | 92 | 31 | 2 | TRUE | 34 | 0 | TRUE | 30 | 0 | TRUE |
| LEU | 93 | 20 | 0 | TRUE | 31 | 10 | TRUE | 24 | 0 | TRUE |
| ILE | 94 | 26 | 1 | TRUE | 30 | 2 | TRUE | 33 | 1 | FALSE |
| GLY | 95 | 31 | 2 | FALSE | 31 | 0 | FALSE | 37 | 1 | FALSE |
| ASN | 96 | 36 | 2 | FALSE | 36 | 0 | FALSE | 42 | 1 | FALSE |
| GLY | 97 | 34 | 2 | FALSE | 34 | 0 | FALSE | 40 | 1 | FALSE |
| TYR | 98 | 32 | 2 | FALSE | 32 | 0 | FALSE | 38 | 1 | FALSE |
| HIS | 99 | 33 | 2 | FALSE | 33 | 0 | FALSE | 39 | 1 | FALSE |
| ASN | 100 | 36 | 2 | FALSE | 36 | 0 | FALSE | 43 | 1 | FALSE |
| GLU | 101 | 22 | 1 | FALSE | 19 | 0 | FALSE | 15 | 4 | FALSE |
| GLN | 102 | 20 | 1 | FALSE | 17 | 0 | FALSE | 13 | 4 | FALSE |
| LYS | 103 | 29 | 2 | FALSE | 44 | 7 | FALSE | 27 | 1 | FALSE |
| GLU | 104 | 28 | 2 | FALSE | 43 | 7 | FALSE | 26 | 1 | FALSE |
| ARG | 105 | 34 | 4 | TRUE | 31 | 1 | TRUE | 21 | 1 | TRUE |
| GLN | 106 | 18 | 0 | TRUE | >51 | 8 | TRUE | | | |
| ALA | 107 | | | | | | | | | |
| ILE | 108 | <9 | 6 | TRUE | <9 | 3 | TRUE | <9 | 3 | TRUE |
| GLU | 109 | 42 | 1 | FALSE | 35 | 0 | FALSE | 52 | 3 | FALSE |
| GLN | 110 | 43 | 1 | FALSE | 36 | 0 | FALSE | 53 | 3 | FALSE |
| LEU | 111 | 41 | 1 | FALSE | 34 | 0 | FALSE | 51 | 3 | FALSE |
| ILE | 112 | 20 | 1 | FALSE | 20 | 1 | FALSE | 19 | 2 | FALSE |
| ARG | 113 | 24 | 1 | FALSE | 24 | 1 | FALSE | 23 | 2 | FALSE |
| HIS | 114 | 27 | 1 | FALSE | 27 | 1 | FALSE | 25 | 2 | FALSE |
| ARG | 115 | 11 | 1 | FALSE | 16 | 1 | FALSE | 14 | 1 | FALSE |
| CYS | 116 | 14 | 1 | FALSE | 19 | 1 | FALSE | 17 | 1 | FALSE |
| ALA | 117 | 13 | 1 | FALSE | 17 | 1 | FALSE | 16 | 1 | FALSE |
| ALA | 118 | 10 | 1 | FALSE | 14 | 1 | FALSE | 13 | 1 | FALSE |
| LEU | 119 | <10 | 2 | TRUE | <10 | 3 | TRUE | <10 | 3 | TRUE |
| VAL | 120 | 20 | 0 | TRUE | 13 | 6 | TRUE | 10 | 5 | TRUE |

| | | | | | | | | | | |
|---|---|---|---|---|---|---|---|---|---|---|
| VAL | 121 | 33 | 3 | TRUE | 33 | 1 | TRUE | 34 | 1 | TRUE |
| HIS | 122 | 38 | 1 | TRUE | 37 | 1 | TRUE | 40 | 1 | TRUE |
| ALA | 123 | 31 | 7 | TRUE | 39 | 1 | TRUE | 21 | 1 | TRUE |
| LYS | 124 | 25 | 6 | TRUE | 36 | 0 | TRUE | 34 | 0 | TRUE |
| MET | 125 | 25 | 6 | TRUE | 26 | 0 | TRUE | 36 | 0 | TRUE |
| ILE | 126 | 25 | 6 | TRUE | 32 | 0 | TRUE | 24 | 0 | TRUE |
| PRO | 127 | | | | | | | | | |
| ASP | 128 | 23 | 2 | FALSE | 18 | 1 | FALSE | 29 | 5 | FALSE |
| ALA | 129 | 23 | 2 | FALSE | 18 | 1 | FALSE | 29 | 5 | FALSE |
| ASP | 130 | 25 | 2 | FALSE | 20 | 1 | FALSE | 30 | 5 | FALSE |
| LEU | 131 | 32 | 0 | TRUE | 34 | 0 | TRUE | 35 | 1 | TRUE |
| ALA | 132 | 20 | 5 | TRUE | 18 | 0 | TRUE | 28 | 1 | TRUE |
| SER | 133 | 32 | 1 | TRUE | 38 | 0 | TRUE | 29 | 1 | TRUE |
| LEU | 134 | >48 | 7 | TRUE | 34 | 0 | TRUE | 32 | 0 | TRUE |
| MET | 135 | 22 | 0 | TRUE | 20 | 0 | TRUE | 20 | 0 | TRUE |
| LYS | 136 | >50 | 5 | TRUE | 50 | 12 | TRUE | >50 | 14 | TRUE |
| GLN | 137 | 20 | 0 | TRUE | 14 | 7 | TRUE | 22 | 0 | TRUE |
| MET | 138 | 28 | 0 | FALSE | 26 | 0 | FALSE | 34 | 0 | FALSE |
| PRO | 139 | | | | | | | | | |
| GLY | 140 | 41 | 7 | FALSE | 28 | 1 | FALSE | 36 | 3 | FALSE |
| MET | 141 | 43 | 7 | FALSE | 30 | 1 | FALSE | 38 | 3 | FALSE |
| VAL | 142 | >46 | 7 | TRUE | >46 | 16 | TRUE | >46 | 6 | TRUE |
| LEU | 143 | 37 | 1 | TRUE | 33 | 1 | TRUE | 34 | 0 | TRUE |
| ILE | 144 | 53 | 4 | FALSE | 47 | 6 | FALSE | 46 | 5 | FALSE |
| ASN | 145 | 59 | 4 | FALSE | 54 | 6 | FALSE | 53 | 5 | FALSE |
| ARG | 146 | 39 | 2 | TRUE | 21 | 12 | TRUE | 27 | 12 | TRUE |
| ILE | 147 | 20 | 12 | TRUE | 21 | 12 | TRUE | 20 | 9 | TRUE |
| LEU | 148 | 31 | 1 | TRUE | 32 | 1 | TRUE | 35 | 4 | TRUE |
| PRO | 149 | | | | | | | | | |
| GLY | 150 | 26 | 1 | FALSE | 26 | 0 | FALSE | 28 | 1 | FALSE |
| PHE | 151 | 27 | 1 | FALSE | 27 | 0 | FALSE | 29 | 1 | FALSE |

| | | | | | | | | | | |
|---|---|---|---|---|---|---|---|---|---|---|
| GLU | 152 | 18 | 0 | TRUE | 18 | 0 | TRUE | 17 | 1 | TRUE |
| ASN | 153 | 34 | 0 | TRUE | 29 | 7 | TRUE | 35 | 2 | TRUE |
| ARG | 154 | 21 | 1 | TRUE | 26 | 9 | TRUE | 20 | 0 | TRUE |
| CYS | 155 | 39 | 2 | TRUE | 40 | 0 | TRUE | 39 | 1 | TRUE |
| ILE | 156 | 42 | 22 | TRUE | 29 | 8 | TRUE | 34 | 3 | TRUE |
| ALA | 157 | 28 | 2 | TRUE | 30 | 9 | TRUE | 30 | 1 | TRUE |
| LEU | 158 | 28 | 1 | TRUE | 23 | 8 | TRUE | 35 | 1 | TRUE |
| ASP | 159 | 18 | 0 | TRUE | 18 | 0 | TRUE | 18 | 0 | TRUE |
| ASP | 160 | 20 | 0 | TRUE | 24 | 5 | TRUE | 20 | 0 | TRUE |
| ARG | 161 | 41 | 2 | FALSE | 41 | 1 | FALSE | 43 | 2 | FALSE |
| TYR | 162 | 41 | 2 | FALSE | 42 | 1 | FALSE | 43 | 2 | FALSE |
| GLY | 163 | 42 | 2 | FALSE | 42 | 1 | FALSE | 44 | 2 | FALSE |
| ALA | 164 | 42 | 2 | FALSE | 43 | 1 | FALSE | 44 | 2 | FALSE |
| TRP | 165 | 39 | 2 | FALSE | 40 | 1 | FALSE | 41 | 2 | FALSE |
| LEU | 166 | 30 | 2 | TRUE | 25 | 6 | TRUE | 43 | 16 | TRUE |
| ALA | 167 | 18 | 0 | TRUE | 17 | 1 | TRUE | 18 | 0 | TRUE |
| THR | 168 | 46 | 1 | FALSE | 54 | 2 | FALSE | 60 | 2 | FALSE |
| ARG | 169 | 47 | 1 | FALSE | 56 | 2 | FALSE | 62 | 2 | FALSE |
| HIS | 170 | 47 | 1 | FALSE | 56 | 2 | FALSE | 62 | 2 | FALSE |
| LEU | 171 | 44 | 1 | FALSE | 52 | 2 | FALSE | 59 | 2 | FALSE |
| ILE | 172 | 41 | 1 | FALSE | 49 | 2 | FALSE | 56 | 2 | FALSE |
| GLN | 173 | 45 | 1 | FALSE | 53 | 2 | FALSE | 60 | 2 | FALSE |
| GLN | 174 | 47 | 1 | FALSE | 56 | 2 | FALSE | 62 | 2 | FALSE |
| GLY | 175 | | | | | | | | | |
| HIS | 176 | | | | | | | | | |
| THR | 177 | | | | | | | | | |
| ARG | 178 | | | | | | | | | |
| ILE | 179 | | | | | | | | | |
| GLY | 180 | | | | | | | | | |
| TYR | 181 | >49 | 1 | FALSE | >49 | 1 | FALSE | >49 | 1 | FALSE |
| LEU | 182 | >47 | 1 | FALSE | >47 | 1 | FALSE | >47 | 1 | FALSE |

| | | | | | | | | | | |
|---|---|---|---|---|---|---|---|---|---|---|
| CYS | 183 | >52 | 1 | FALSE | >52 | 1 | FALSE | >52 | 1 | FALSE |
| SER | 184 | 31 | 1 | FALSE | 49 | 2 | FALSE | 40 | 2 | FALSE |
| ASN | 185 | 30 | 1 | FALSE | 48 | 2 | FALSE | 39 | 2 | FALSE |
| HIS | 186 | 28 | 1 | FALSE | 46 | 2 | FALSE | 37 | 2 | FALSE |
| SER | 187 | 29 | 1 | FALSE | 47 | 2 | FALSE | 38 | 2 | FALSE |
| ILE | 188 | <11 | 1 | FALSE | 16 | 2 | FALSE | <11 | 1 | FALSE |
| SER | 189 | <14 | 1 | FALSE | 19 | 2 | FALSE | <14 | 1 | FALSE |
| ASP | 190 | <16 | 1 | FALSE | 21 | 2 | FALSE | <16 | 1 | FALSE |
| ALA | 191 | <12 | 2 | TRUE | 21 | 4 | TRUE | <12 | 4 | TRUE |
| GLU | 192 | <13 | 3 | TRUE | 21 | 1 | TRUE | <13 | 4 | TRUE |
| ASP | 193 | 20 | 0 | TRUE | 32 | 0 | TRUE | 20 | 0 | TRUE |
| ARG | 194 | 31 | 1 | TRUE | 36 | 1 | TRUE | 29 | 1 | TRUE |
| LEU | 195 | 32 | 0 | TRUE | 37 | 1 | TRUE | 32 | 0 | TRUE |
| GLN | 196 | 33 | 1 | TRUE | 40 | 2 | TRUE | 34 | 1 | TRUE |
| GLY | 197 | 40 | 2 | TRUE | 41 | 2 | TRUE | 38 | 2 | TRUE |
| TYR | 198 | >49 | 5 | TRUE | >49 | 12 | TRUE | >49 | 7 | TRUE |
| TYR | 199 | >48 | 6 | TRUE | >48 | 5 | TRUE | >48 | 12 | TRUE |
| ASP | 200 | 38 | 0 | TRUE | >51 | 10 | TRUE | 38 | 0 | TRUE |
| ALA | 201 | >49 | 4 | TRUE | >49 | 12 | TRUE | >49 | 5 | TRUE |
| LEU | 202 | >47 | 7 | TRUE | 36 | 0 | TRUE | 38 | 1 | TRUE |
| ALA | 203 | 34 | 0 | TRUE | 35 | 4 | TRUE | 32 | 0 | TRUE |
| GLU | 204 | 19 | 1 | TRUE | 32 | 3 | TRUE | 18 | 0 | TRUE |
| SER | 205 | 20 | 0 | FALSE | 19 | 0 | FALSE | 19 | 0 | FALSE |
| GLY | 206 | 20 | 0 | FALSE | 19 | 0 | FALSE | 20 | 0 | FALSE |
| ILE | 207 | 15 | 0 | FALSE | 15 | 0 | FALSE | 15 | 0 | FALSE |
| ALA | 208 | 17 | 0 | FALSE | 16 | 0 | FALSE | 17 | 0 | FALSE |
| ALA | 209 | 24 | 1 | FALSE | 24 | 0 | FALSE | 25 | 1 | FALSE |
| ASN | 210 | 27 | 1 | FALSE | 27 | 0 | FALSE | 27 | 1 | FALSE |
| ASP | 211 | 21 | 1 | TRUE | 20 | 0 | TRUE | 19 | 1 | TRUE |
| ARG | 212 | 18 | 0 | TRUE | 18 | 0 | TRUE | 17 | 1 | TRUE |
| LEU | 213 | 18 | 0 | TRUE | 18 | 0 | TRUE | 16 | 0 | TRUE |

| | | | | | | | | | | |
|---|---|---|---|---|---|---|---|---|---|---|
| VAL | 214 | 30 | 1 | FALSE | 31 | 0 | FALSE | 29 | 1 | FALSE |
| THR | 215 | 34 | 1 | FALSE | 35 | 0 | FALSE | 33 | 1 | FALSE |
| PHE | 216 | >49 | 2 | FALSE | 43 | 9 | FALSE | 53 | 5 | FALSE |
| GLY | 217 | >50 | 2 | FALSE | 43 | 9 | FALSE | 53 | 5 | FALSE |
| GLU | 218 | 20 | 1 | TRUE | 22 | 0 | TRUE | 20 | 0 | TRUE |
| PRO | 219 | | | | | | | | | |
| ASP | 220 | 21 | 1 | FALSE | 16 | 2 | FALSE | 20 | 1 | FALSE |
| GLU | 221 | 20 | 1 | FALSE | 16 | 2 | FALSE | 20 | 1 | FALSE |
| SER | 222 | 23 | 1 | FALSE | 18 | 2 | FALSE | 22 | 1 | FALSE |
| GLY | 223 | 35 | 2 | FALSE | 40 | 1 | FALSE | 38 | 1 | FALSE |
| GLY | 224 | 34 | 2 | FALSE | 39 | 1 | FALSE | 37 | 1 | FALSE |
| GLU | 225 | 33 | 2 | FALSE | 39 | 1 | FALSE | 37 | 1 | FALSE |
| GLN | 226 | 33 | 2 | FALSE | 38 | 1 | FALSE | 36 | 1 | FALSE |
| ALA | 227 | 34 | 2 | FALSE | 39 | 1 | FALSE | 38 | 1 | FALSE |
| MET | 228 | 35 | 1 | TRUE | 48 | 17 | TRUE | 50 | 23 | TRUE |
| THR | 229 | >50 | 2 | FALSE | 36 | 2 | FALSE | >50 | 2 | FALSE |
| GLU | 230 | >50 | 2 | FALSE | 36 | 2 | FALSE | >50 | 2 | FALSE |
| LEU | 231 | >46 | 7 | TRUE | >46 | 7 | TRUE | >46 | 7 | TRUE |
| LEU | 232 | 23 | 1 | FALSE | 24 | 0 | FALSE | 22 | 1 | FALSE |
| GLY | 233 | 26 | 1 | FALSE | 27 | 0 | FALSE | 25 | 1 | FALSE |
| ARG | 234 | 19 | 0 | FALSE | 17 | 1 | FALSE | 20 | 0 | FALSE |
| GLY | 235 | 18 | 0 | FALSE | 17 | 1 | FALSE | 19 | 0 | FALSE |
| ARG | 236 | 19 | 0 | FALSE | 17 | 1 | FALSE | 20 | 0 | FALSE |
| ASN | 237 | 21 | 0 | FALSE | 20 | 1 | FALSE | 22 | 0 | FALSE |
| PHE | 238 | 46 | 4 | FALSE | 48 | 2 | FALSE | 37 | 1 | TRUE |
| THR | 239 | 45 | 4 | FALSE | 47 | 2 | FALSE | 55 | 3 | FALSE |
| ALA | 240 | 46 | 4 | FALSE | 48 | 2 | FALSE | 56 | 3 | FALSE |
| VAL | 241 | 49 | 3 | FALSE | >46 | 2 | FALSE | >46 | 2 | FALSE |
| ALA | 242 | 52 | 3 | FALSE | >49 | 2 | FALSE | >49 | 2 | FALSE |
| CYS | 243 | 38 | 0 | TRUE | >53 | 4 | TRUE | 46 | 33 | TRUE |
| TYR | 244 | | | | | | | | | |

| | | | | | | | | | | |
|---|---|---|---|---|---|---|---|---|---|---|
| ASN | 245 | >53 | 6 | TRUE | 40 | 0 | TRUE | >53 | 4 | TRUE |
| ASP | 246 | 23 | 15 | TRUE | 41 | 3 | TRUE | 45 | 18 | TRUE |
| SER | 247 | 32 | 8 | FALSE | 43 | 3 | FALSE | 47 | 5 | FALSE |
| MET | 248 | 33 | 8 | FALSE | 43 | 3 | FALSE | 48 | 5 | FALSE |
| ALA | 249 | >50 | 2 | FALSE | >50 | 2 | FALSE | >50 | 3 | FALSE |
| ALA | 250 | >50 | 2 | FALSE | >50 | 2 | FALSE | >50 | 3 | FALSE |
| GLY | 251 | >50 | 4 | TRUE | >50 | 6 | TRUE | 42 | 31 | TRUE |
| ALA | 252 | >51 | 6 | TRUE | >51 | 7 | TRUE | >51 | 6 | TRUE |
| MET | 253 | 45 | 20 | TRUE | 38 | 1 | TRUE | 43 | 21 | TRUE |
| GLY | 254 | 22 | 0 | TRUE | 26 | 0 | TRUE | 24 | 1 | TRUE |
| VAL | 255 | >47 | 6 | TRUE | 36 | 2 | TRUE | 45 | 16 | TRUE |
| LEU | 256 | 25 | 3 | TRUE | 28 | 3 | TRUE | 29 | 1 | FALSE |
| ASN | 257 | 32 | 0 | TRUE | 36 | 0 | TRUE | 34 | 1 | FALSE |
| ASP | 258 | 36 | 0 | TRUE | 40 | 4 | TRUE | 32 | 4 | TRUE |
| ASN | 259 | 32 | 4 | TRUE | 36 | 4 | TRUE | 34 | 3 | TRUE |
| GLY | 260 | <15 | 2 | TRUE | <15 | 5 | TRUE | 27 | 7 | TRUE |
| ILE | 261 | 19 | 3 | TRUE | 26 | 9 | TRUE | 15 | 2 | TRUE |
| ASP | 262 | 34 | 6 | FALSE | 38 | 1 | FALSE | 29 | 5 | FALSE |
| VAL | 263 | 30 | 6 | FALSE | 34 | 1 | FALSE | 25 | 5 | FALSE |
| PRO | 264 | | | | | | | | | |
| GLY | 265 | 33 | 2 | FALSE | 21 | 1 | FALSE | 39 | 2 | FALSE |
| GLU | 266 | 35 | 2 | FALSE | 23 | 1 | FALSE | 41 | 2 | FALSE |
| ILE | 267 | 30 | 2 | FALSE | 18 | 1 | FALSE | 36 | 2 | FALSE |
| SER | 268 | 35 | 2 | FALSE | 24 | 1 | FALSE | 41 | 2 | FALSE |
| LEU | 269 | 34 | 2 | TRUE | 32 | 8 | TRUE | 28 | 6 | TRUE |
| ILE | 270 | 37 | 18 | TRUE | 32 | 4 | TRUE | 45 | 19 | TRUE |
| GLY | 271 | 18 | 3 | TRUE | 48 | 21 | TRUE | 42 | 28 | TRUE |
| PHE | 272 | 44 | 2 | FALSE | 39 | 2 | FALSE | 26 | 2 | FALSE |
| ASP | 273 | 46 | 2 | FALSE | 40 | 2 | FALSE | 28 | 2 | FALSE |
| ASP | 274 | 44 | 2 | FALSE | 39 | 2 | FALSE | 26 | 2 | FALSE |
| VAL | 275 | 40 | 2 | FALSE | 34 | 2 | FALSE | 22 | 2 | FALSE |

| | | | | | | | | | | |
|---|---|---|---|---|---|---|---|---|---|---|
| LEU | 276 | 13 | 1 | TRUE | 38 | 25 | TRUE | 33 | 26 | TRUE |
| VAL | 277 | | | | | | | | | |
| SER | 278 | 19 | 2 | TRUE | 21 | 1 | TRUE | 18 | 12 | TRUE |
| ARG | 279 | 26 | 0 | TRUE | 22 | 9 | TRUE | 27 | 6 | TRUE |
| TYR | 280 | 12 | 1 | FALSE | 13 | 1 | FALSE | 16 | 1 | FALSE |
| VAL | 281 | 9 | 1 | FALSE | 10 | 1 | FALSE | 12 | 1 | FALSE |
| ARG | 282 | 12 | 1 | FALSE | 13 | 1 | FALSE | 16 | 1 | FALSE |
| PRO | 283 | 0 | 0 | FALSE | 0 | 0 | FALSE | 0 | 0 | FALSE |
| ARG | 284 | 12 | 1 | FALSE | 13 | 1 | FALSE | 15 | 1 | FALSE |
| LEU | 285 | 14 | 3 | TRUE | 15 | 5 | TRUE | 20 | 6 | TRUE |
| THR | 286 | 22 | 0 | TRUE | 22 | 1 | TRUE | 27 | 3 | TRUE |
| THR | 287 | 30 | 1 | TRUE | 32 | 0 | TRUE | 34 | 0 | TRUE |
| VAL | 288 | 28 | 0 | TRUE | 36 | 0 | TRUE | 29 | 3 | TRUE |
| ARG | 289 | 30 | 0 | TRUE | 34 | 0 | TRUE | 34 | 0 | TRUE |
| TYR | 290 | 21 | 3 | TRUE | 34 | 0 | TRUE | 25 | 6 | TRUE |
| PRO | 291 | | | | | | | | | |
| ILE | 292 | 17 | 1 | FALSE | 12 | 0 | FALSE | 19 | 1 | FALSE |
| VAL | 293 | 18 | 1 | FALSE | 12 | 0 | FALSE | 19 | 1 | FALSE |
| THR | 294 | 28 | 3 | TRUE | 32 | 0 | TRUE | 31 | 1 | TRUE |
| MET | 295 | 36 | 0 | TRUE | 40 | 0 | TRUE | >51 | 14 | TRUE |
| ALA | 296 | 32 | 0 | TRUE | 34 | 0 | TRUE | >50 | 4 | TRUE |
| THR | 297 | 44 | 0 | TRUE | 46 | 0 | TRUE | >49 | 6 | TRUE |
| GLN | 298 | >51 | 12 | TRUE | >51 | 11 | TRUE | 46 | 1 | TRUE |
| ALA | 299 | 53 | 6 | FALSE | 52 | 2 | FALSE | 54 | 5 | FALSE |
| ALA | 300 | 52 | 6 | FALSE | 51 | 2 | FALSE | 53 | 5 | FALSE |
| GLU | 301 | >49 | 2 | FALSE | >49 | 2 | FALSE | 53 | 8 | FALSE |
| LEU | 302 | >46 | 2 | FALSE | >46 | 2 | FALSE | 50 | 8 | FALSE |
| ALA | 303 | 42 | 2 | TRUE | >49 | 6 | TRUE | 44 | 0 | TRUE |
| LEU | 304 | >47 | 4 | TRUE | >47 | 19 | TRUE | >47 | 7 | TRUE |
| ALA | 305 | >49 | 6 | TRUE | 36 | 0 | TRUE | >49 | 5 | TRUE |
| LEU | 306 | 28 | 0 | TRUE | 28 | 0 | TRUE | 30 | 0 | TRUE |

| | | | | | | | | | | |
|---|---|---|---|---|---|---|---|---|---|---|
| ALA | 307 | 24 | 0 | TRUE | 22 | 0 | TRUE | 26 | 0 | TRUE |
| ASP | 308 | 20 | 0 | TRUE | 20 | 0 | TRUE | 19 | 1 | TRUE |
| ASN | 309 | <15 | 5 | TRUE | 18 | 0 | TRUE | 18 | 0 | TRUE |
| ARG | 310 | 18 | 0 | TRUE | 21 | 1 | TRUE | 19 | 2 | TRUE |
| PRO | 311 | | | | | | | | | |
| LEU | 312 | 15 | 0 | FALSE | 14 | 0 | FALSE | 15 | 0 | FALSE |
| PRO | 313 | | | | | | | | | |
| GLU | 314 | 16 | 0 | TRUE | 14 | 0 | TRUE | 17 | 1 | TRUE |
| ILE | 315 | 12 | 0 | TRUE | 10 | 0 | TRUE | 12 | 0 | TRUE |
| THR | 316 | <12 | 4 | TRUE | <12 | 3 | TRUE | 14 | 0 | TRUE |
| ASN | 317 | <17 | 5 | TRUE | <17 | 5 | TRUE | <17 | 5 | TRUE |
| VAL | 318 | <11 | 3 | TRUE | <11 | 3 | TRUE | <11 | 3 | TRUE |
| PHE | 319 | 16 | 0 | TRUE | 15 | 1 | TRUE | 18 | 0 | TRUE |
| SER | 320 | <16 | 7 | TRUE | 17 | 9 | TRUE | <16 | 2 | TRUE |
| PRO | 321 | | | | | | | | | |
| THR | 322 | 14 | 1 | FALSE | 15 | 1 | FALSE | 15 | 1 | FALSE |
| LEU | 323 | 14 | 1 | FALSE | 15 | 1 | FALSE | 15 | 1 | FALSE |
| VAL | 324 | 10 | 5 | TRUE | 11 | 8 | TRUE | <8 | 0 | TRUE |
| ARG | 325 | | | | | | | | | |
| ARG | 326 | | | | | | | | | |
| HIS | 327 | | | | | | | | | |
| SER | 328 | | | | | | | | | |
| VAL | 329 | | | | | | | | | |
| SER | 330 | 17 | 2 | FALSE | 17 | 2 | FALSE | 12 | 3 | FALSE |
| THR | 331 | 17 | 2 | FALSE | 17 | 2 | FALSE | 12 | 3 | FALSE |
| PRO | 332 | | | | | | | | | |
| SER | 333 | 18 | 2 | FALSE | 14 | 2 | FALSE | 14 | 1 | FALSE |
| LEU | 334 | 16 | 2 | FALSE | 12 | 2 | FALSE | 11 | 1 | FALSE |
| GLU | 335 | 19 | 4 | TRUE | 24 | 0 | TRUE | 24 | 0 | TRUE |

**Table S10. $\Delta G_{op}$ values for RbsR**

All values in kJ/mol.

| Res. name | Res. number | Apo | Std Dev | Single resolved | RIB | Std Dev | Single resolved | *rbsO* | Std Dev | Single resolved |
|---|---|---|---|---|---|---|---|---|---|---|
| MET | 1 | | | | | | | | | |
| ALA | 2 | | | | | | | | | |
| THR | 3 | | | | | | | | | |
| MET | 4 | | | | | | | | | |
| LYS | 5 | | | | | | | | | |
| ASP | 6 | | | | | | | | | |
| VAL | 7 | | | | | | | | | |
| ALA | 8 | 12 | 1 | FALSE | 14 | 1 | FALSE | 32 | 0 | FALSE |
| ARG | 9 | 13 | 1 | FALSE | 15 | 1 | FALSE | 33 | 0 | FALSE |
| LEU | 10 | 11 | 1 | FALSE | 13 | 1 | FALSE | 31 | 0 | FALSE |
| ALA | 11 | 13 | 5 | TRUE | 16 | 1 | TRUE | | | |
| GLY | 12 | 14 | 0 | FALSE | 13 | 0 | FALSE | 24 | 0 | FALSE |
| VAL | 13 | 11 | 0 | FALSE | 10 | 0 | FALSE | 21 | 0 | FALSE |
| SER | 14 | 16 | 0 | FALSE | 14 | 0 | FALSE | 25 | 0 | FALSE |
| THR | 15 | 16 | 0 | FALSE | 14 | 0 | FALSE | 25 | 0 | FALSE |
| SER | 16 | 18 | 0 | FALSE | 16 | 0 | FALSE | 27 | 0 | FALSE |
| THR | 17 | 16 | 0 | FALSE | 14 | 0 | FALSE | 25 | 0 | FALSE |
| VAL | 18 | 18 | 1 | FALSE | 18 | 1 | FALSE | 21 | 0 | FALSE |
| SER | 19 | 22 | 1 | FALSE | 22 | 1 | FALSE | 25 | 0 | FALSE |
| HIS | 20 | 23 | 1 | FALSE | 23 | 1 | FALSE | 26 | 0 | FALSE |
| VAL | 21 | | | | | | | | | |
| ILE | 22 | | | | | | | | | |
| ASN | 23 | 22 | 1 | FALSE | 31 | 1 | FALSE | 28 | 1 | FALSE |
| LYS | 24 | 22 | 1 | FALSE | 32 | 1 | FALSE | 28 | 1 | FALSE |
| ASP | 25 | 22 | 1 | FALSE | 31 | 1 | FALSE | 28 | 1 | FALSE |
| ARG | 26 | 20 | 1 | FALSE | 29 | 1 | FALSE | 26 | 1 | FALSE |
| PHE | 27 | 21 | 1 | FALSE | 30 | 1 | FALSE | 27 | 1 | FALSE |

| | | | | | | | | | | |
|---|---|---|---|---|---|---|---|---|---|---|
| VAL | 28 | | | | | | | | | |
| SER | 29 | | | | | | | | | |
| GLU | 30 | 15 | 2 | FALSE | 19 | 1 | FALSE | 12 | 2 | FALSE |
| ALA | 31 | 13 | 2 | FALSE | 17 | 1 | FALSE | 11 | 2 | FALSE |
| ILE | 32 | 10 | 7 | TRUE | 13 | 1 | TRUE | 23 | 6 | TRUE |
| THR | 33 | 16 | 0 | TRUE | 16 | 0 | TRUE | 28 | 5 | TRUE |
| ALA | 34 | 11 | 2 | FALSE | 22 | 1 | FALSE | 27 | 2 | FALSE |
| LYS | 35 | 10 | 2 | FALSE | 20 | 1 | FALSE | 25 | 2 | FALSE |
| VAL | 36 | 19 | 4 | TRUE | 17 | 3 | TRUE | 32 | 5 | TRUE |
| GLU | 37 | <12 | 4 | TRUE | 16 | 1 | TRUE | 27 | 6 | TRUE |
| ALA | 38 | <12 | 5 | TRUE | <12 | 4 | TRUE | 30 | 3 | TRUE |
| ALA | 39 | 32 | 0 | TRUE | 32 | 0 | TRUE | 30 | 2 | TRUE |
| ILE | 40 | 16 | 3 | TRUE | 15 | 1 | TRUE | 32 | 3 | TRUE |
| LYS | 41 | <12 | 5 | TRUE | 16 | 1 | TRUE | 28 | 1 | TRUE |
| GLU | 42 | 14 | 3 | FALSE | 22 | 1 | FALSE | 33 | 1 | FALSE |
| LEU | 43 | 10 | 3 | FALSE | 18 | 1 | FALSE | 29 | 1 | FALSE |
| ASN | 44 | 16 | 2 | FALSE | 20 | 0 | FALSE | 17 | 3 | FALSE |
| TYR | 45 | 14 | 2 | FALSE | 18 | 0 | FALSE | 15 | 3 | FALSE |
| ALA | 46 | 10 | 1 | FALSE | 17 | 0 | FALSE | 14 | 1 | FALSE |
| PRO | 47 | | | | | | | | | |
| SER | 48 | 11 | 1 | FALSE | 18 | 0 | FALSE | 15 | 1 | FALSE |
| ALA | 49 | 12 | 1 | FALSE | 19 | 0 | FALSE | 16 | 1 | FALSE |
| LEU | 50 | 7 | 1 | FALSE | 14 | 0 | FALSE | 11 | 1 | FALSE |
| ALA | 51 | <12 | 6 | TRUE | 15 | 1 | TRUE | 26 | 5 | TRUE |
| ARG | 52 | 36 | 0 | TRUE | 36 | 0 | TRUE | 27 | 6 | TRUE |
| SER | 53 | <16 | 2 | FALSE | <16 | 1 | FALSE | 31 | 1 | FALSE |
| LEU | 54 | <12 | 2 | FALSE | <12 | 1 | FALSE | 26 | 1 | FALSE |
| LYS | 55 | <12 | 3 | TRUE | <12 | 3 | TRUE | 24 | 1 | TRUE |
| LEU | 56 | <11 | 6 | TRUE | <11 | 5 | TRUE | 16 | 0 | TRUE |
| ASN | 57 | 17 | 7 | TRUE | 20 | 0 | TRUE | 21 | 2 | TRUE |
| GLN | 58 | <15 | 6 | TRUE | <15 | 4 | TRUE | 29 | 5 | TRUE |

| | | | | | | | | | | |
|---|---|---|---|---|---|---|---|---|---|---|
| THR | 59 | <14 | 5 | TRUE | 18 | 0 | TRUE | 18 | 0 | TRUE |
| HIS | 60 | 20 | 0 | TRUE | 20 | 0 | TRUE | >49 | 21 | TRUE |
| THR | 61 | 34 | 2 | TRUE | 35 | 1 | TRUE | >48 | 5 | TRUE |
| ILE | 62 | 44 | 5 | FALSE | 45 | 8 | FALSE | 49 | 4 | FALSE |
| GLY | 63 | 45 | 5 | FALSE | 46 | 8 | FALSE | 50 | 4 | FALSE |
| MET | 64 | 47 | 13 | TRUE | >48 | 12 | TRUE | >48 | 26 | TRUE |
| LEU | 65 | 34 | 0 | TRUE | >45 | 6 | TRUE | >45 | 24 | TRUE |
| ILE | 66 | 24 | 0 | TRUE | 33 | 3 | TRUE | 31 | 1 | TRUE |
| THR | 67 | 18 | 0 | TRUE | 19 | 1 | TRUE | 15 | 4 | TRUE |
| ALA | 68 | 26 | 0 | TRUE | 31 | 2 | TRUE | 26 | 1 | TRUE |
| SER | 69 | 16 | 6 | TRUE | <15 | 6 | TRUE | 17 | 6 | TRUE |
| THR | 70 | <14 | 6 | TRUE | 17 | 1 | TRUE | <14 | 6 | TRUE |
| ASN | 71 | 24 | 1 | FALSE | 33 | 1 | FALSE | 22 | 1 | FALSE |
| PRO | 72 | | | | | | | | | |
| PHE | 73 | 18 | 1 | FALSE | 26 | 1 | FALSE | 16 | 1 | FALSE |
| TYR | 74 | 19 | 1 | FALSE | 28 | 1 | FALSE | 17 | 1 | FALSE |
| SER | 75 | 23 | 1 | FALSE | 31 | 1 | FALSE | 20 | 1 | FALSE |
| GLU | 76 | 22 | 1 | FALSE | 30 | 1 | FALSE | 19 | 1 | FALSE |
| LEU | 77 | 20 | 0 | TRUE | 27 | 2 | TRUE | 22 | 0 | TRUE |
| VAL | 78 | 29 | 2 | TRUE | 32 | 2 | TRUE | 32 | 2 | TRUE |
| ARG | 79 | 35 | 2 | TRUE | 33 | 3 | TRUE | 30 | 0 | TRUE |
| GLY | 80 | 30 | 1 | FALSE | 34 | 1 | FALSE | 34 | 3 | FALSE |
| VAL | 81 | 26 | 1 | FALSE | 30 | 1 | FALSE | 30 | 3 | FALSE |
| GLU | 82 | 30 | 2 | TRUE | 32 | 1 | TRUE | 35 | 8 | TRUE |
| ARG | 83 | 32 | 1 | FALSE | 27 | 1 | FALSE | 50 | 6 | FALSE |
| SER | 84 | 36 | 1 | FALSE | 30 | 1 | FALSE | 53 | 6 | FALSE |
| CYS | 85 | 33 | 6 | TRUE | 32 | 5 | TRUE | 40 | 3 | TRUE |
| PHE | 86 | 26 | 4 | TRUE | 28 | 4 | TRUE | 37 | 2 | TRUE |
| GLU | 87 | 20 | 4 | TRUE | 20 | 1 | TRUE | 20 | 0 | TRUE |
| ARG | 88 | <13 | 5 | TRUE | 18 | 1 | TRUE | 28 | 0 | TRUE |
| GLY | 89 | 30 | 1 | FALSE | 29 | 1 | FALSE | 41 | 2 | FALSE |

| | | | | | | | | | | |
|---|---|---|---|---|---|---|---|---|---|---|
| TYR | 90 | 28 | 1 | FALSE | 28 | 1 | FALSE | 40 | 2 | FALSE |
| SER | 91 | 31 | 1 | FALSE | 31 | 1 | FALSE | 43 | 2 | FALSE |
| LEU | 92 | 30 | 3 | TRUE | 27 | 4 | TRUE | 36 | 3 | TRUE |
| VAL | 93 | 23 | 2 | FALSE | 29 | 2 | FALSE | 35 | 3 | FALSE |
| LEU | 94 | 25 | 2 | FALSE | 30 | 2 | FALSE | 36 | 3 | FALSE |
| CYS | 95 | | | | | | | | | |
| ASN | 96 | 40 | 0 | TRUE | 30 | 6 | TRUE | 37 | 7 | TRUE |
| THR | 97 | 29 | 1 | FALSE | 29 | 1 | FALSE | 31 | 1 | FALSE |
| GLU | 98 | 29 | 1 | FALSE | 28 | 1 | FALSE | 30 | 1 | FALSE |
| GLY | 99 | 27 | 1 | FALSE | 26 | 1 | FALSE | 29 | 1 | FALSE |
| ASP | 100 | 30 | 1 | FALSE | 29 | 1 | FALSE | 31 | 1 | FALSE |
| GLU | 101 | 26 | 1 | FALSE | 26 | 1 | FALSE | 28 | 1 | FALSE |
| GLN | 102 | 28 | 1 | FALSE | 27 | 1 | FALSE | 29 | 1 | FALSE |
| ARG | 103 | 30 | 1 | FALSE | 29 | 1 | FALSE | 31 | 1 | FALSE |
| MET | 104 | <14 | 4 | TRUE | <14 | 5 | TRUE | 31 | 1 | FALSE |
| ASN | 105 | <16 | 2 | FALSE | 12 | 2 | FALSE | 42 | 7 | FALSE |
| ARG | 106 | <15 | 2 | FALSE | 11 | 2 | FALSE | 41 | 7 | FALSE |
| ASN | 107 | 19 | 8 | TRUE | 29 | 7 | TRUE | 34 | 1 | TRUE |
| LEU | 108 | 29 | 1 | TRUE | 21 | 5 | TRUE | 35 | 2 | FALSE |
| GLU | 109 | 15 | 8 | TRUE | 19 | 3 | TRUE | 34 | 2 | FALSE |
| THR | 110 | 11 | 2 | FALSE | 12 | 2 | FALSE | 28 | 1 | FALSE |
| LEU | 111 | 11 | 2 | FALSE | 11 | 2 | FALSE | 27 | 1 | FALSE |
| MET | 112 | <12 | 3 | TRUE | <12 | 3 | TRUE | | | |
| GLN | 113 | 34 | 2 | TRUE | >48 | 20 | TRUE | >48 | 13 | TRUE |
| LYS | 114 | <14 | 4 | TRUE | 14 | 6 | TRUE | 37 | 1 | TRUE |
| ARG | 115 | <14 | 6 | TRUE | 15 | 5 | TRUE | 28 | 1 | FALSE |
| VAL | 116 | 24 | 8 | TRUE | 16 | 1 | TRUE | 24 | 1 | FALSE |
| ASP | 117 | 26 | 8 | TRUE | 39 | 19 | TRUE | 37 | 2 | TRUE |
| GLY | 118 | 37 | 4 | TRUE | >46 | 26 | TRUE | 34 | 1 | TRUE |
| LEU | 119 | 31 | 2 | TRUE | >45 | 12 | TRUE | >45 | 15 | TRUE |
| LEU | 120 | 33 | 3 | TRUE | >43 | 6 | TRUE | >43 | 7 | TRUE |

| | | | | | | | | | | |
|---|---|---|---|---|---|---|---|---|---|---|
| LEU | 121 | 31 | 2 | TRUE | >43 | 16 | TRUE | >43 | 7 | TRUE |
| LEU | 122 | 22 | 6 | TRUE | 22 | 1 | TRUE | 32 | 3 | TRUE |
| CYS | 123 | 25 | 1 | TRUE | | | | <15 | 6 | TRUE |
| THR | 124 | <16 | 6 | TRUE | 20 | 0 | TRUE | 20 | 1 | TRUE |
| GLU | 125 | 9 | 1 | FALSE | 19 | 2 | FALSE | 19 | 0 | FALSE |
| THR | 126 | 7 | 1 | FALSE | 17 | 2 | FALSE | 17 | 0 | FALSE |
| HIS | 127 | 10 | 1 | FALSE | 20 | 2 | FALSE | 20 | 0 | FALSE |
| GLN | 128 | 29 | 1 | FALSE | 20 | 2 | FALSE | 35 | 1 | FALSE |
| PRO | 129 | | | | | | | | | |
| SER | 130 | 28 | 1 | FALSE | 19 | 2 | FALSE | 34 | 1 | FALSE |
| ARG | 131 | 30 | 1 | FALSE | 20 | 2 | FALSE | 35 | 1 | FALSE |
| GLU | 132 | 28 | 1 | FALSE | 19 | 2 | FALSE | 34 | 1 | FALSE |
| ILE | 133 | <8 | 3 | TRUE | 9 | 4 | TRUE | 8 | 4 | TRUE |
| MET | 134 | 12 | 6 | TRUE | 18 | 0 | TRUE | 18 | 1 | TRUE |
| GLN | 135 | <14 | 4 | TRUE | 19 | 1 | TRUE | 24 | 1 | TRUE |
| ARG | 136 | 16 | 10 | TRUE | <15 | 8 | TRUE | 17 | 10 | TRUE |
| TYR | 137 | 15 | 8 | TRUE | 18 | 1 | TRUE | 29 | 1 | TRUE |
| PRO | 138 | | | | | | | | | |
| THR | 139 | 13 | 2 | FALSE | 13 | 2 | FALSE | 14 | 4 | FALSE |
| VAL | 140 | 12 | 2 | FALSE | 12 | 2 | FALSE | 13 | 4 | FALSE |
| PRO | 141 | | | | | | | | | |
| THR | 142 | 33 | 1 | FALSE | 52 | 2 | FALSE | 50 | 5 | FALSE |
| VAL | 143 | 32 | 1 | FALSE | 51 | 2 | FALSE | 49 | 5 | FALSE |
| MET | 144 | 15 | 7 | TRUE | 17 | 1 | TRUE | 18 | 0 | TRUE |
| MET | 145 | 25 | 1 | TRUE | 34 | 0 | TRUE | 29 | 1 | TRUE |
| ASP | 146 | | | | | | | | | |
| TRP | 147 | 11 | 0 | FALSE | 17 | 0 | FALSE | 19 | 1 | FALSE |
| ALA | 148 | 14 | 0 | FALSE | 19 | 0 | FALSE | 22 | 1 | FALSE |
| PRO | 149 | | | | | | | | | |
| PHE | 150 | 12 | 0 | FALSE | 17 | 0 | FALSE | 9 | 1 | FALSE |
| ASP | 151 | 15 | 0 | FALSE | 21 | 0 | FALSE | 13 | 1 | FALSE |

| | | | | | | | | | | |
|---|---|---|---|---|---|---|---|---|---|---|
| GLY | 152 | 13 | 0 | FALSE | 19 | 0 | FALSE | 11 | 1 | FALSE |
| ASP | 153 | | | | | | | | | |
| SER | 154 | | | | | | | | | |
| ASP | 155 | | | | | | | | | |
| LEU | 156 | | | | | | | | | |
| ILE | 157 | 29 | 1 | TRUE | >42 | 29 | TRUE | >42 | 7 | TRUE |
| GLN | 158 | 18 | 0 | TRUE | 25 | 2 | TRUE | 24 | 1 | TRUE |
| ASP | 159 | 35 | 1 | TRUE | 36 | 1 | TRUE | 35 | 3 | TRUE |
| ASN | 160 | 21 | 1 | TRUE | 23 | 3 | TRUE | 19 | 2 | TRUE |
| SER | 161 | 28 | 4 | TRUE | 25 | 3 | TRUE | 22 | 2 | TRUE |
| LEU | 162 | 20 | 4 | TRUE | 27 | 4 | TRUE | 28 | 4 | TRUE |
| LEU | 163 | 20 | 3 | TRUE | 27 | 4 | TRUE | 23 | 3 | TRUE |
| GLY | 164 | 34 | 0 | TRUE | 43 | 20 | TRUE | >46 | 20 | TRUE |
| GLY | 165 | >48 | 12 | TRUE | >48 | 15 | TRUE | >48 | 6 | TRUE |
| ASP | 166 | 46 | 19 | TRUE | 38 | 2 | TRUE | 48 | 16 | TRUE |
| LEU | 167 | >43 | 24 | TRUE | >43 | 17 | TRUE | >43 | 20 | TRUE |
| ALA | 168 | >46 | 6 | TRUE | >46 | 12 | TRUE | >46 | 18 | TRUE |
| THR | 169 | 37 | 3 | TRUE | 38 | 3 | TRUE | >47 | 19 | TRUE |
| GLN | 170 | 40 | 1 | TRUE | 40 | 1 | TRUE | 40 | 2 | TRUE |
| TYR | 171 | >47 | 5 | TRUE | >47 | 15 | TRUE | >47 | 21 | TRUE |
| LEU | 172 | >44 | 14 | TRUE | >44 | 18 | TRUE | >44 | 15 | TRUE |
| ILE | 173 | >42 | 8 | TRUE | >42 | 11 | TRUE | >42 | 12 | TRUE |
| ASP | 174 | >47 | 7 | TRUE | >47 | 26 | TRUE | 39 | 2 | TRUE |
| LYS | 175 | 33 | 1 | FALSE | | | | 27 | 1 | FALSE |
| GLY | 176 | 34 | 1 | FALSE | | | | 28 | 1 | FALSE |
| HIS | 177 | 35 | 1 | FALSE | | | | 29 | 1 | FALSE |
| THR | 178 | 35 | 1 | FALSE | | | | 29 | 1 | FALSE |
| ARG | 179 | 36 | 1 | FALSE | | | | 29 | 1 | FALSE |
| ILE | 180 | 31 | 1 | FALSE | | | | 25 | 1 | FALSE |
| ALA | 181 | 25 | 1 | TRUE | 32 | 2 | TRUE | 36 | 2 | TRUE |
| CYS | 182 | 54 | 4 | FALSE | 53 | 4 | FALSE | >51 | 20 | TRUE |

| | | | | | | | | | | |
|---|---|---|---|---|---|---|---|---|---|---|
| ILE | 183 | 49 | 4 | FALSE | 49 | 4 | FALSE | 43 | 19 | TRUE |
| THR | 184 | 35 | 2 | TRUE | 43 | 13 | TRUE | 36 | 3 | TRUE |
| GLY | 185 | >48 | 10 | TRUE | 35 | 6 | TRUE | 39 | 3 | TRUE |
| PRO | 186 | | | | | | | | | |
| LEU | 187 | 13 | 2 | FALSE | 15 | 1 | FALSE | 11 | 2 | FALSE |
| ASP | 188 | 17 | 2 | FALSE | 19 | 1 | FALSE | 15 | 2 | FALSE |
| LYS | 189 | 26 | 5 | TRUE | 35 | 2 | TRUE | 27 | 5 | TRUE |
| THR | 190 | 16 | 1 | TRUE | 17 | 1 | TRUE | 16 | 2 | TRUE |
| PRO | 191 | | | | | | | | | |
| ALA | 192 | 30 | 2 | FALSE | 38 | 2 | FALSE | 51 | 4 | FALSE |
| ARG | 193 | 32 | 2 | FALSE | 40 | 2 | FALSE | 52 | 4 | FALSE |
| LEU | 194 | 29 | 2 | TRUE | 31 | 2 | TRUE | 33 | 2 | TRUE |
| ARG | 195 | >47 | 26 | TRUE | >47 | 6 | TRUE | >47 | 22 | TRUE |
| LEU | 196 | >45 | 20 | TRUE | >45 | 5 | TRUE | >45 | 6 | TRUE |
| GLU | 197 | 46 | 18 | TRUE | 36 | 2 | TRUE | 36 | 2 | TRUE |
| GLY | 198 | 45 | 13 | TRUE | 37 | 1 | TRUE | 36 | 1 | TRUE |
| TYR | 199 | >47 | 14 | TRUE | >47 | 14 | TRUE | >47 | 10 | TRUE |
| ARG | 200 | 38 | 2 | TRUE | 38 | 2 | TRUE | 40 | 1 | TRUE |
| ALA | 201 | 37 | 1 | TRUE | 41 | 2 | TRUE | 37 | 1 | TRUE |
| ALA | 202 | 51 | 5 | FALSE | >47 | 3 | FALSE | 37 | 1 | FALSE |
| MET | 203 | 51 | 5 | FALSE | >47 | 3 | FALSE | 37 | 1 | FALSE |
| LYS | 204 | | | | | | | | | |
| ARG | 205 | | | | | | | | | |
| ALA | 206 | | | | | | | | | |
| GLY | 207 | 27 | 2 | FALSE | 29 | 2 | FALSE | 46 | 5 | FALSE |
| LEU | 208 | 25 | 2 | FALSE | 27 | 2 | FALSE | 44 | 5 | FALSE |
| ASN | 209 | 13 | 2 | FALSE | 13 | 4 | FALSE | 13 | 2 | FALSE |
| ILE | 210 | 9 | 2 | FALSE | 9 | 4 | FALSE | 9 | 2 | FALSE |
| PRO | 211 | | | | | | | | | |
| ASP | 212 | 23 | 3 | FALSE | 26 | 1 | FALSE | 26 | 3 | FALSE |
| GLY | 213 | 23 | 3 | FALSE | 25 | 1 | FALSE | 25 | 3 | FALSE |

| | | | | | | | | | | |
|---|---|---|---|---|---|---|---|---|---|---|
| TYR | 214 | 32 | 0 | TRUE | 33 | 3 | TRUE | 31 | 1 | TRUE |
| GLU | 215 | 18 | 11 | TRUE | 22 | 9 | TRUE | 20 | 0 | TRUE |
| VAL | 216 | 17 | 2 | FALSE | 24 | 1 | FALSE | 30 | 2 | FALSE |
| THR | 217 | 20 | 2 | FALSE | 28 | 1 | FALSE | 34 | 2 | FALSE |
| GLY | 218 | 22 | 2 | FALSE | 30 | 1 | FALSE | 36 | 2 | FALSE |
| ASP | 219 | 23 | 2 | FALSE | 30 | 1 | FALSE | 36 | 2 | FALSE |
| PHE | 220 | 23 | 1 | TRUE | 25 | 7 | TRUE | 16 | 1 | TRUE |
| GLU | 221 | | | | | | | | | |
| PHE | 222 | 12 | 4 | TRUE | 18 | 0 | TRUE | <11 | 3 | TRUE |
| ASN | 223 | 30 | 1 | FALSE | 29 | 4 | FALSE | 34 | 4 | FALSE |
| GLY | 224 | 29 | 1 | FALSE | 28 | 4 | FALSE | 33 | 4 | FALSE |
| GLY | 225 | 28 | 1 | FALSE | 27 | 4 | FALSE | 32 | 4 | FALSE |
| PHE | 226 | 38 | 16 | TRUE | 46 | 14 | TRUE | >47 | 15 | TRUE |
| ASP | 227 | 31 | 2 | FALSE | 29 | 1 | FALSE | 28 | 0 | FALSE |
| ALA | 228 | 29 | 2 | FALSE | 27 | 1 | FALSE | 26 | 0 | FALSE |
| MET | 229 | >47 | 10 | TRUE | 45 | 7 | TRUE | >47 | 13 | TRUE |
| ARG | 230 | 29 | 0 | FALSE | 29 | 0 | FALSE | 30 | 0 | FALSE |
| GLN | 231 | 30 | 0 | FALSE | 30 | 0 | FALSE | 31 | 0 | FALSE |
| LEU | 232 | 26 | 0 | FALSE | 26 | 0 | FALSE | 27 | 0 | FALSE |
| LEU | 233 | | | | | | | | | |
| SER | 234 | 21 | 1 | TRUE | 20 | 0 | TRUE | 20 | 1 | TRUE |
| HIS | 235 | 25 | 3 | FALSE | 26 | 1 | FALSE | 33 | 3 | FALSE |
| PRO | 236 | | | | | | | | | |
| LEU | 237 | 18 | 3 | FALSE | 20 | 1 | FALSE | 26 | 3 | FALSE |
| ARG | 238 | 23 | 2 | FALSE | 25 | 5 | FALSE | 27 | 5 | FALSE |
| PRO | 239 | | | | | | | | | |
| GLN | 240 | 23 | 2 | FALSE | 25 | 5 | FALSE | 26 | 5 | FALSE |
| ALA | 241 | >49 | 13 | TRUE | 39 | 2 | TRUE | 33 | 8 | TRUE |
| VAL | 242 | >44 | 11 | TRUE | >44 | 7 | TRUE | >44 | 7 | TRUE |
| PHE | 243 | >45 | 7 | TRUE | >45 | 11 | TRUE | >45 | 7 | TRUE |
| THR | 244 | 37 | 20 | TRUE | 35 | 4 | TRUE | 38 | 4 | TRUE |

| | | | | | | | | | | |
|---|---|---|---|---|---|---|---|---|---|---|
| GLY | 245 | 37 | 5 | TRUE | 35 | 1 | TRUE | 34 | 0 | TRUE |
| ASN | 246 | 50 | 19 | TRUE | 42 | 3 | TRUE | >51 | 15 | TRUE |
| ASP | 247 | >50 | 5 | TRUE | >50 | 5 | TRUE | >50 | 5 | TRUE |
| ALA | 248 | 50 | 6 | FALSE | 51 | 6 | FALSE | 47 | 8 | FALSE |
| MET | 249 | 50 | 6 | FALSE | 52 | 6 | FALSE | 48 | 8 | FALSE |
| ALA | 250 | 41 | 3 | TRUE | 40 | 2 | TRUE | 38 | 2 | TRUE |
| VAL | 251 | 43 | 16 | TRUE | 35 | 2 | TRUE | >44 | 14 | TRUE |
| GLY | 252 | 35 | 1 | TRUE | 37 | 1 | TRUE | 38 | 0 | TRUE |
| VAL | 253 | 52 | 5 | FALSE | 52 | 4 | FALSE | 53 | 3 | FALSE |
| TYR | 254 | 53 | 5 | FALSE | 52 | 4 | FALSE | 54 | 3 | FALSE |
| GLN | 255 | 38 | 2 | TRUE | 42 | 3 | TRUE | >48 | 13 | TRUE |
| ALA | 256 | 52 | 8 | FALSE | 57 | 4 | FALSE | 54 | 4 | FALSE |
| LEU | 257 | 47 | 8 | FALSE | 53 | 4 | FALSE | 50 | 4 | FALSE |
| TYR | 258 | 33 | 1 | TRUE | 33 | 1 | TRUE | 33 | 2 | TRUE |
| GLN | 259 | 22 | 0 | TRUE | 25 | 4 | TRUE | 22 | 0 | TRUE |
| ALA | 260 | 29 | 4 | TRUE | 30 | 4 | TRUE | 28 | 3 | TRUE |
| GLU | 261 | 28 | 4 | TRUE | 26 | 4 | TRUE | 28 | 4 | TRUE |
| LEU | 262 | >43 | 19 | TRUE | 44 | 12 | TRUE | 34 | 0 | TRUE |
| GLN | 263 | 38 | 5 | TRUE | >47 | 5 | TRUE | >47 | 16 | TRUE |
| VAL | 264 | 18 | 0 | TRUE | 18 | 0 | TRUE | 18 | 0 | TRUE |
| PRO | 265 | | | | | | | | | |
| GLN | 266 | 6 | 2 | FALSE | 16 | 0 | FALSE | 16 | 0 | FALSE |
| ASP | 267 | 36 | 5 | TRUE | 39 | 3 | TRUE | >49 | 20 | TRUE |
| ILE | 268 | >42 | 21 | TRUE | >42 | 22 | TRUE | 39 | 16 | TRUE |
| ALA | 269 | 37 | 10 | TRUE | 45 | 24 | TRUE | 42 | 19 | TRUE |
| VAL | 270 | >44 | 14 | TRUE | >44 | 21 | TRUE | >44 | 11 | TRUE |
| ILE | 271 | 26 | 3 | TRUE | 28 | 2 | TRUE | 25 | 1 | TRUE |
| GLY | 272 | >46 | 6 | TRUE | >46 | 5 | TRUE | >46 | 5 | TRUE |
| TYR | 273 | 47 | 21 | TRUE | >47 | 14 | TRUE | 38 | 1 | TRUE |
| ASP | 274 | >48 | 20 | TRUE | >48 | 20 | TRUE | >48 | 22 | TRUE |
| ASP | 275 | >47 | 18 | TRUE | 38 | 5 | TRUE | >47 | 13 | TRUE |

| | | | | | | | | | | |
|---|---|---|---|---|---|---|---|---|---|---|
| ILE | 276 | >42 | 7 | TRUE | >42 | 17 | TRUE | >42 | 22 | TRUE |
| GLU | 277 | 16 | 0 | TRUE | 18 | 0 | TRUE | 14 | 0 | TRUE |
| LEU | 278 | 40 | 20 | TRUE | 31 | 1 | TRUE | 30 | 0 | TRUE |
| ALA | 279 | | | | | | | | | |
| SER | 280 | | | | | | | | | |
| PHE | 281 | 29 | 1 | FALSE | 31 | 1 | FALSE | 45 | 3 | FALSE |
| MET | 282 | 29 | 1 | FALSE | 31 | 1 | FALSE | 45 | 3 | FALSE |
| THR | 283 | 29 | 1 | FALSE | 31 | 1 | FALSE | 45 | 3 | FALSE |
| PRO | 284 | | | | | | | | | |
| PRO | 285 | | | | | | | | | |
| LEU | 286 | 47 | 1 | FALSE | 43 | 1 | FALSE | 41 | 1 | FALSE |
| THR | 287 | 50 | 1 | FALSE | 46 | 1 | FALSE | 44 | 1 | FALSE |
| THR | 288 | 53 | 1 | FALSE | 48 | 1 | FALSE | 47 | 1 | FALSE |
| ILE | 289 | 49 | 1 | FALSE | 45 | 1 | FALSE | 43 | 1 | FALSE |
| HIS | 290 | 51 | 1 | FALSE | 47 | 1 | FALSE | 45 | 1 | FALSE |
| GLN | 291 | 10 | 3 | FALSE | 31 | 9 | TRUE | 29 | 4 | FALSE |
| PRO | 292 | | | | | | | | | |
| LYS | 293 | 24 | 0 | TRUE | 21 | 12 | TRUE | 21 | 9 | TRUE |
| ASP | 294 | 22 | 1 | FALSE | 21 | 1 | FALSE | 20 | 2 | FALSE |
| GLU | 295 | 19 | 1 | FALSE | 18 | 1 | FALSE | 18 | 2 | FALSE |
| LEU | 296 | 16 | 1 | FALSE | 16 | 1 | FALSE | 15 | 2 | FALSE |
| GLY | 297 | 31 | 0 | FALSE | 47 | 8 | FALSE | 46 | 6 | FALSE |
| GLU | 298 | 32 | 0 | FALSE | 48 | 8 | FALSE | 48 | 6 | FALSE |
| LEU | 299 | <9 | 4 | TRUE | 30 | 3 | TRUE | 20 | 8 | TRUE |
| ALA | 300 | 13 | 1 | FALSE | 13 | 1 | FALSE | 10 | 1 | FALSE |
| ILE | 301 | 10 | 1 | FALSE | 10 | 1 | FALSE | 7 | 1 | FALSE |
| ASP | 302 | 13 | 1 | FALSE | 14 | 1 | FALSE | 10 | 1 | FALSE |
| VAL | 303 | 9 | 1 | FALSE | 9 | 1 | FALSE | 6 | 1 | FALSE |
| LEU | 304 | 10 | 1 | FALSE | 10 | 1 | FALSE | 7 | 1 | FALSE |
| ILE | 305 | 32 | 2 | TRUE | >42 | 7 | TRUE | >42 | 8 | TRUE |
| HIS | 306 | 19 | 3 | TRUE | 23 | 6 | TRUE | 24 | 0 | TRUE |

| | | | | | | | | | | |
|---|---|---|---|---|---|---|---|---|---|---|
| ARG | 307 | 25 | 2 | FALSE | 21 | 3 | FALSE | 39 | 2 | FALSE |
| ILE | 308 | 21 | 2 | FALSE | 17 | 3 | FALSE | 35 | 2 | FALSE |
| THR | 309 | 22 | 2 | FALSE | 18 | 3 | FALSE | 36 | 2 | FALSE |
| GLN | 310 | 21 | 9 | TRUE | 21 | 12 | TRUE | 31 | 1 | TRUE |
| PRO | 311 | | | | | | | | | |
| THR | 312 | 12 | 10 | TRUE | 15 | 11 | TRUE | 12 | 7 | TRUE |
| LEU | 313 | <11 | 3 | TRUE | <11 | 5 | TRUE | <11 | 5 | TRUE |
| GLN | 314 | 18 | 4 | TRUE | <12 | 5 | TRUE | 19 | 2 | TRUE |
| GLN | 315 | 19 | 3 | TRUE | 19 | 1 | TRUE | 21 | 2 | TRUE |
| GLN | 316 | <15 | 6 | TRUE | 15 | 6 | TRUE | <15 | 5 | TRUE |
| ARG | 317 | <15 | 6 | TRUE | 19 | 3 | TRUE | 22 | 1 | TRUE |
| LEU | 318 | 13 | 5 | TRUE | 18 | 3 | TRUE | 12 | 6 | TRUE |
| GLN | 319 | 31 | 1 | TRUE | 27 | 5 | TRUE | 32 | 0 | TRUE |
| LEU | 320 | 18 | 2 | TRUE | 21 | 5 | TRUE | 17 | 2 | TRUE |
| THR | 321 | 15 | 3 | TRUE | 17 | 1 | TRUE | 13 | 6 | TRUE |
| PRO | 322 | | | | | | | | | |
| ILE | 323 | 32 | 0 | FALSE | >42 | 12 | FALSE | 42 | 10 | FALSE |
| LEU | 324 | 14 | 0 | TRUE | 14 | 0 | TRUE | 15 | 1 | TRUE |
| MET | 325 | 24 | 0 | TRUE | 36 | 0 | TRUE | 24 | 0 | TRUE |
| GLU | 326 | | | | | | | | | |
| ARG | 327 | 22 | 0 | FALSE | 20 | 1 | FALSE | 33 | 2 | FALSE |
| GLY | 328 | 24 | 0 | FALSE | 22 | 1 | FALSE | 35 | 2 | FALSE |
| SER | 329 | 26 | 0 | FALSE | 24 | 1 | FALSE | 37 | 2 | FALSE |
| ALA | 330 | 15 | 0 | FALSE | 13 | 1 | FALSE | 25 | 2 | FALSE |

**Table S11. $\Delta G_{op}$ values for MglB**

All values in kJ/mol.

| Res. name | Res. number | Apo | Std Dev | Single resolved | GAL | Std Dev | Single resolved |
|---|---|---|---|---|---|---|---|
| MET | 1 | | | | | | |
| ASN | 2 | | | | | | |
| LYS | 3 | | | | | | |
| LYS | 4 | | | | | | |
| VAL | 5 | | | | | | |
| LEU | 6 | | | | | | |
| THR | 7 | 29 | 1 | FALSE | 31 | 0 | FALSE |
| LEU | 8 | 29 | 1 | FALSE | 30 | 0 | FALSE |
| SER | 9 | 32 | 1 | FALSE | 33 | 0 | FALSE |
| ALA | 10 | 33 | 1 | FALSE | 34 | 0 | FALSE |
| VAL | 11 | 27 | 1 | FALSE | 29 | 0 | FALSE |
| MET | 12 | 30 | 1 | FALSE | 32 | 0 | FALSE |
| ALA | 13 | 31 | 1 | FALSE | 33 | 0 | FALSE |
| SER | 14 | 33 | 1 | FALSE | 34 | 0 | FALSE |
| MET | 15 | 32 | 1 | FALSE | 34 | 0 | FALSE |
| LEU | 16 | 28 | 1 | FALSE | 30 | 0 | FALSE |
| PHE | 17 | 28 | 1 | FALSE | 30 | 0 | FALSE |
| GLY | 18 | | | | | | |
| ALA | 19 | | | | | | |
| ALA | 20 | | | | | | |
| ALA | 21 | | | | | | |
| HIS | 22 | | | | | | |
| ALA | 23 | | | | | | |
| ALA | 24 | | | | | | |
| ASP | 25 | | | | | | |
| THR | 26 | | | | | | |
| ARG | 27 | 20 | 0 | TRUE | 20 | 0 | TRUE |

| | | | | | | | |
|---|---|---|---|---|---|---|---|
| ILE | 28 | 28 | 0 | FALSE | 34 | 1 | TRUE |
| GLY | 29 | 29 | 0 | FALSE | >48 | 26 | TRUE |
| VAL | 30 | 27 | 0 | FALSE | 42 | 1 | FALSE |
| THR | 31 | 29 | 0 | FALSE | 44 | 1 | FALSE |
| ILE | 32 | 30 | 0 | FALSE | 39 | 1 | FALSE |
| TYR | 33 | 30 | 0 | FALSE | 39 | 1 | FALSE |
| LYS | 34 | 23 | 0 | FALSE | 45 | 5 | FALSE |
| TYR | 35 | 23 | 0 | FALSE | 44 | 5 | FALSE |
| ASP | 36 | 22 | 0 | TRUE | 20 | 0 | TRUE |
| ASP | 37 | 28 | 0 | TRUE | >49 | 22 | TRUE |
| ASN | 38 | <15 | 6 | TRUE | 15 | 4 | TRUE |
| PHE | 39 | <14 | 2 | TRUE | 28 | 0 | TRUE |
| MET | 40 | | | | >50 | 4 | TRUE |
| SER | 41 | 22 | 0 | TRUE | 33 | 7 | TRUE |
| VAL | 42 | 30 | 0 | TRUE | >47 | 22 | TRUE |
| VAL | 43 | 25 | 1 | FALSE | 34 | 1 | FALSE |
| ARG | 44 | 29 | 1 | FALSE | 38 | 1 | FALSE |
| LYS | 45 | 32 | 0 | TRUE | >51 | 13 | TRUE |
| ALA | 46 | 34 | 0 | TRUE | 48 | 24 | TRUE |
| ILE | 47 | 27 | 1 | TRUE | >46 | 18 | TRUE |
| GLU | 48 | 28 | 0 | TRUE | 32 | 2 | TRUE |
| GLN | 49 | 30 | 0 | TRUE | 36 | 4 | TRUE |
| ASP | 50 | 16 | 7 | TRUE | 25 | 3 | TRUE |
| ALA | 51 | 25 | 2 | TRUE | 23 | 3 | TRUE |
| LYS | 52 | 20 | 1 | TRUE | <13 | 6 | TRUE |
| ALA | 53 | 24 | 3 | TRUE | 27 | 3 | TRUE |
| ALA | 54 | 20 | 0 | FALSE | 14 | 4 | FALSE |
| PRO | 55 | | | | | | |
| ASP | 56 | 21 | 1 | FALSE | 26 | 1 | FALSE |
| VAL | 57 | 17 | 1 | FALSE | 22 | 1 | FALSE |
| GLN | 58 | 21 | 1 | FALSE | 26 | 1 | FALSE |

| | | | | | | | |
|---|---|---|---|---|---|---|---|
| LEU | 59 | <11 | 3 | TRUE | <11 | 3 | TRUE |
| LEU | 60 | <9 | 2 | TRUE | 23 | 11 | TRUE |
| MET | 61 | <12 | 3 | TRUE | 16 | 0 | TRUE |
| ASN | 62 | 30 | 0 | FALSE | 43 | 2 | FALSE |
| ASP | 63 | 29 | 0 | FALSE | 42 | 2 | FALSE |
| SER | 64 | 28 | 0 | FALSE | 41 | 2 | FALSE |
| GLN | 65 | 17 | 1 | FALSE | 26 | 0 | FALSE |
| ASN | 66 | 18 | 1 | FALSE | 28 | 0 | FALSE |
| ASP | 67 | 17 | 1 | FALSE | 26 | 0 | FALSE |
| GLN | 68 | 14 | 1 | FALSE | 23 | 0 | FALSE |
| SER | 69 | 28 | 2 | FALSE | 45 | 2 | FALSE |
| LYS | 70 | 26 | 2 | FALSE | 43 | 2 | FALSE |
| GLN | 71 | 26 | 2 | FALSE | 42 | 2 | FALSE |
| ASN | 72 | 34 | 0 | TRUE | >54 | 11 | TRUE |
| ASP | 73 | 35 | 1 | TRUE | 39 | 1 | TRUE |
| GLN | 74 | >49 | 5 | TRUE | 34 | 2 | TRUE |
| ILE | 75 | 34 | 1 | FALSE | 33 | 1 | FALSE |
| ASP | 76 | 37 | 1 | FALSE | 36 | 1 | FALSE |
| VAL | 77 | 48 | 4 | FALSE | >45 | 2 | FALSE |
| LEU | 78 | 49 | 4 | FALSE | >46 | 2 | FALSE |
| LEU | 79 | 24 | 0 | TRUE | 27 | 1 | TRUE |
| ALA | 80 | 39 | 1 | TRUE | 46 | 31 | TRUE |
| LYS | 81 | 32 | 0 | TRUE | 33 | 1 | TRUE |
| GLY | 82 | 40 | 4 | TRUE | >50 | 17 | TRUE |
| VAL | 83 | 18 | 0 | TRUE | <10 | 6 | TRUE |
| LYS | 84 | 16 | 0 | TRUE | 22 | 0 | TRUE |
| ALA | 85 | 28 | 0 | TRUE | 28 | 0 | TRUE |
| LEU | 86 | >47 | 18 | TRUE | >47 | 26 | TRUE |
| ALA | 87 | 40 | 4 | TRUE | >49 | 12 | TRUE |
| ILE | 88 | 31 | 1 | TRUE | 36 | 2 | TRUE |
| ASN | 89 | >51 | 2 | TRUE | >51 | 34 | TRUE |

| | | | | | | | |
|---|---|---|---|---|---|---|---|
| LEU | 90 | 37 | 2 | TRUE | >48 | 20 | TRUE |
| VAL | 91 | 22 | 0 | TRUE | 27 | 1 | TRUE |
| ASP | 92 | 33 | 1 | TRUE | 34 | 1 | TRUE |
| PRO | 93 | | | | | | |
| ALA | 94 | 10 | 1 | FALSE | 17 | 1 | FALSE |
| ALA | 95 | 11 | 1 | FALSE | 18 | 1 | FALSE |
| ALA | 96 | 11 | 1 | FALSE | 18 | 1 | FALSE |
| GLY | 97 | 11 | 1 | FALSE | 18 | 1 | FALSE |
| THR | 98 | <14 | 3 | TRUE | <14 | 5 | TRUE |
| VAL | 99 | 29 | 1 | TRUE | 30 | 0 | TRUE |
| ILE | 100 | 30 | 0 | TRUE | 30 | 0 | TRUE |
| GLU | 101 | 40 | 1 | TRUE | 38 | 1 | TRUE |
| LYS | 102 | 37 | 1 | TRUE | >49 | 12 | TRUE |
| ALA | 103 | 32 | 0 | TRUE | 29 | 5 | TRUE |
| ARG | 104 | >50 | 4 | TRUE | >50 | 3 | TRUE |
| GLY | 105 | 27 | 1 | FALSE | 30 | 1 | FALSE |
| GLN | 106 | 27 | 1 | FALSE | 30 | 1 | FALSE |
| ASN | 107 | 30 | 1 | FALSE | 32 | 1 | FALSE |
| VAL | 108 | 24 | 1 | FALSE | 26 | 1 | FALSE |
| PRO | 109 | | | | | | |
| VAL | 110 | 48 | 3 | FALSE | 46 | 1 | FALSE |
| VAL | 111 | 49 | 3 | FALSE | 47 | 1 | FALSE |
| PHE | 112 | 51 | 3 | FALSE | 49 | 1 | FALSE |
| PHE | 113 | 38 | 2 | TRUE | >49 | 3 | TRUE |
| ASN | 114 | 28 | 36 | TRUE | >53 | 2 | TRUE |
| LYS | 115 | 30 | 0 | TRUE | >51 | 3 | TRUE |
| GLU | 116 | 12 | 1 | FALSE | 11 | 1 | FALSE |
| PRO | 117 | | | | | | |
| SER | 118 | 13 | 1 | FALSE | 12 | 1 | FALSE |
| ARG | 119 | <15 | 7 | TRUE | 34 | 0 | TRUE |
| LYS | 120 | <14 | 3 | TRUE | <14 | 5 | TRUE |

| | | | | | | | |
|---|---|---|---|---|---|---|---|
| ALA | 121 | 20 | 3 | FALSE | 21 | 1 | FALSE |
| LEU | 122 | 16 | 3 | FALSE | 17 | 1 | FALSE |
| ASP | 123 | 31 | 2 | TRUE | 28 | 0 | TRUE |
| SER | 124 | 20 | 0 | TRUE | 24 | 0 | TRUE |
| TYR | 125 | 16 | 2 | TRUE | 18 | 0 | TRUE |
| ASP | 126 | 38 | 0 | TRUE | <14 | 4 | TRUE |
| LYS | 127 | 14 | 3 | TRUE | 26 | 0 | TRUE |
| ALA | 128 | 26 | 0 | TRUE | 33 | 3 | TRUE |
| TYR | 129 | 36 | 0 | TRUE | >48 | 4 | TRUE |
| TYR | 130 | <12 | 3 | TRUE | <12 | 3 | TRUE |
| VAL | 131 | >46 | 5 | TRUE | <10 | 4 | TRUE |
| GLY | 132 | 37 | 10 | TRUE | >49 | 20 | TRUE |
| THR | 133 | <14 | 1 | FALSE | 37 | 1 | FALSE |
| ASP | 134 | <15 | 1 | FALSE | 38 | 1 | FALSE |
| SER | 135 | 18 | 2 | FALSE | <14 | 2 | FALSE |
| LYS | 136 | 19 | 2 | FALSE | <15 | 2 | FALSE |
| GLU | 137 | 37 | 2 | TRUE | 40 | 1 | TRUE |
| SER | 138 | 32 | 2 | TRUE | 14 | 6 | TRUE |
| GLY | 139 | <15 | 4 | TRUE | 36 | 2 | TRUE |
| ILE | 140 | 42 | 3 | FALSE | 47 | 2 | FALSE |
| ILE | 141 | 40 | 3 | FALSE | 45 | 2 | FALSE |
| GLN | 142 | 44 | 3 | FALSE | 49 | 2 | FALSE |
| GLY | 143 | 59 | 6 | FALSE | >51 | 1 | FALSE |
| ASP | 144 | 60 | 6 | FALSE | >51 | 1 | FALSE |
| LEU | 145 | 54 | 6 | FALSE | >46 | 1 | FALSE |
| ILE | 146 | 20 | 0 | TRUE | 22 | 3 | TRUE |
| ALA | 147 | >48 | 4 | TRUE | >48 | 3 | TRUE |
| LYS | 148 | 35 | 1 | FALSE | 41 | 3 | FALSE |
| HIS | 149 | 36 | 1 | FALSE | 42 | 3 | FALSE |
| TRP | 150 | 34 | 1 | FALSE | 40 | 3 | FALSE |
| ALA | 151 | 35 | 1 | FALSE | 40 | 3 | FALSE |

| | | | | | | | |
|---|---|---|---|---|---|---|---|
| ALA | 152 | 9 | 1 | FALSE | 12 | 3 | FALSE |
| ASN | 153 | 12 | 1 | FALSE | 14 | 3 | FALSE |
| GLN | 154 | 12 | 1 | FALSE | 14 | 3 | FALSE |
| GLY | 155 | 10 | 1 | FALSE | 23 | 3 | TRUE |
| TRP | 156 | 22 | 2 | TRUE | 17 | 7 | TRUE |
| ASP | 157 | <13 | 3 | TRUE | 18 | 11 | TRUE |
| LEU | 158 | 32 | 0 | TRUE | >46 | 11 | TRUE |
| ASN | 159 | 24 | 0 | TRUE | 34 | 0 | TRUE |
| LYS | 160 | 34 | 0 | TRUE | 25 | 1 | TRUE |
| ASP | 161 | 33 | 1 | FALSE | 35 | 0 | FALSE |
| GLY | 162 | 31 | 1 | FALSE | 33 | 0 | FALSE |
| GLN | 163 | 36 | 8 | TRUE | 44 | 0 | TRUE |
| ILE | 164 | 45 | 2 | FALSE | 54 | 2 | FALSE |
| GLN | 165 | 47 | 2 | FALSE | 56 | 2 | FALSE |
| PHE | 166 | 48 | 2 | FALSE | 57 | 2 | FALSE |
| VAL | 167 | | | | | | |
| LEU | 168 | | | | | | |
| LEU | 169 | 31 | 1 | FALSE | 47 | 10 | FALSE |
| LYS | 170 | 34 | 1 | FALSE | 50 | 10 | FALSE |
| GLY | 171 | 37 | 1 | TRUE | 37 | 2 | TRUE |
| GLU | 172 | <13 | 6 | TRUE | 17 | 9 | TRUE |
| PRO | 173 | | | | | | |
| GLY | 174 | 17 | 0 | FALSE | 22 | 2 | FALSE |
| HIS | 175 | 33 | 0 | FALSE | 37 | 1 | FALSE |
| PRO | 176 | | | | | | |
| ASP | 177 | 31 | 0 | FALSE | 35 | 1 | FALSE |
| ALA | 178 | 24 | 2 | TRUE | 48 | 19 | TRUE |
| GLU | 179 | 28 | 0 | FALSE | 29 | 1 | FALSE |
| ALA | 180 | 28 | 0 | FALSE | 29 | 1 | FALSE |
| ARG | 181 | 29 | 0 | FALSE | 30 | 1 | FALSE |
| THR | 182 | 21 | 1 | FALSE | 25 | 0 | FALSE |

| | | | | | | | |
|---|---|---|---|---|---|---|---|
| THR | 183 | 21 | 1 | FALSE | 25 | 0 | FALSE |
| TYR | 184 | 28 | 0 | TRUE | 32 | 1 | TRUE |
| VAL | 185 | 34 | 0 | TRUE | >46 | 6 | TRUE |
| ILE | 186 | 34 | 2 | TRUE | >45 | 19 | TRUE |
| LYS | 187 | 34 | 1 | TRUE | >48 | 15 | TRUE |
| GLU | 188 | 41 | 4 | TRUE | 37 | 1 | TRUE |
| LEU | 189 | 31 | 2 | TRUE | 32 | 0 | TRUE |
| ASN | 190 | 34 | 0 | TRUE | 36 | 0 | TRUE |
| ASP | 191 | 22 | 0 | TRUE | 22 | 0 | TRUE |
| LYS | 192 | 36 | 2 | TRUE | 43 | 1 | TRUE |
| GLY | 193 | 35 | 6 | TRUE | 37 | 1 | TRUE |
| ILE | 194 | 26 | 5 | TRUE | 26 | 0 | TRUE |
| LYS | 195 | <12 | 3 | TRUE | <12 | 6 | TRUE |
| THR | 196 | 24 | 0 | TRUE | 24 | 0 | TRUE |
| GLU | 197 | 37 | 1 | TRUE | 40 | 0 | TRUE |
| GLN | 198 | <13 | 5 | TRUE | 14 | 7 | TRUE |
| LEU | 199 | 38 | 5 | TRUE | 38 | 2 | TRUE |
| GLN | 200 | 38 | 5 | TRUE | 47 | 20 | TRUE |
| LEU | 201 | 18 | 0 | TRUE | 17 | 1 | TRUE |
| ASP | 202 | 31 | 5 | TRUE | 38 | 1 | TRUE |
| THR | 203 | 15 | 0 | FALSE | 17 | 2 | FALSE |
| ALA | 204 | 17 | 0 | FALSE | 19 | 2 | FALSE |
| MET | 205 | 27 | 6 | TRUE | >50 | 6 | TRUE |
| TRP | 206 | 22 | 4 | TRUE | 26 | 0 | TRUE |
| ASP | 207 | 30 | 0 | TRUE | 36 | 0 | TRUE |
| THR | 208 | 11 | 6 | TRUE | 17 | 1 | TRUE |
| ALA | 209 | 36 | 0 | TRUE | 14 | 6 | TRUE |
| GLN | 210 | 21 | 3 | FALSE | 38 | 3 | FALSE |
| ALA | 211 | 22 | 3 | FALSE | 39 | 3 | FALSE |
| LYS | 212 | 22 | 13 | TRUE | >49 | 4 | TRUE |
| ASP | 213 | 34 | 0 | FALSE | 39 | 2 | FALSE |

| | | | | | | | |
|---|---|---|---|---|---|---|---|
| LYS | 214 | 31 | 0 | FALSE | 37 | 2 | FALSE |
| MET | 215 | 33 | 1 | TRUE | 40 | 0 | TRUE |
| ASP | 216 | 41 | 1 | TRUE | >51 | 4 | TRUE |
| ALA | 217 | 22 | 1 | TRUE | 27 | 2 | TRUE |
| TRP | 218 | 29 | 1 | TRUE | 27 | 4 | TRUE |
| LEU | 219 | >46 | 5 | TRUE | >46 | 4 | TRUE |
| SER | 220 | 34 | 6 | TRUE | 40 | 0 | TRUE |
| GLY | 221 | <15 | 4 | TRUE | <15 | 4 | TRUE |
| PRO | 222 | | | | | | |
| ASN | 223 | 9 | 1 | FALSE | 25 | 2 | FALSE |
| ALA | 224 | 10 | 1 | FALSE | 25 | 2 | FALSE |
| ASN | 225 | 15 | 4 | FALSE | 21 | 2 | FALSE |
| LYS | 226 | 14 | 4 | FALSE | 20 | 2 | FALSE |
| ILE | 227 | 29 | 2 | FALSE | 33 | 7 | FALSE |
| GLU | 228 | 30 | 2 | FALSE | 35 | 7 | FALSE |
| VAL | 229 | 35 | 2 | TRUE | 36 | 2 | TRUE |
| VAL | 230 | 37 | 4 | TRUE | >45 | 5 | TRUE |
| ILE | 231 | 36 | 1 | TRUE | >45 | 4 | TRUE |
| ALA | 232 | 37 | 1 | TRUE | >48 | 34 | TRUE |
| ASN | 233 | 63 | 3 | FALSE | 41 | 6 | FALSE |
| ASN | 234 | 65 | 3 | FALSE | 42 | 6 | FALSE |
| ASP | 235 | 22 | 0 | TRUE | 26 | 0 | TRUE |
| ALA | 236 | 26 | 0 | TRUE | 34 | 0 | TRUE |
| MET | 237 | 38 | 1 | TRUE | >50 | 3 | TRUE |
| ALA | 238 | 40 | 0 | TRUE | 43 | 1 | TRUE |
| MET | 239 | >50 | 14 | TRUE | >50 | 4 | TRUE |
| GLY | 240 | 40 | 1 | TRUE | >50 | 13 | TRUE |
| ALA | 241 | 41 | 10 | TRUE | >51 | 21 | TRUE |
| VAL | 242 | >46 | 16 | TRUE | 28 | 0 | TRUE |
| GLU | 243 | 31 | 4 | TRUE | 38 | 0 | TRUE |
| ALA | 244 | | | | | | |

| | | | | | | | |
|---|---|---|---|---|---|---|---|
| LEU | 245 | | | | | | |
| LYS | 246 | | | | | | |
| ALA | 247 | | | | | | |
| HIS | 248 | | | | | | |
| ASN | 249 | | | | | | |
| LYS | 250 | | | | | | |
| SER | 251 | | | | | | |
| SER | 252 | 28 | 0 | TRUE | 30 | 0 | TRUE |
| ILE | 253 | 52 | 1 | FALSE | 31 | 1 | FALSE |
| PRO | 254 | | | | | | |
| VAL | 255 | 49 | 1 | FALSE | 28 | 1 | FALSE |
| PHE | 256 | 52 | 1 | FALSE | 31 | 1 | FALSE |
| GLY | 257 | >50 | 2 | FALSE | 37 | 2 | FALSE |
| VAL | 258 | >47 | 2 | FALSE | 34 | 2 | FALSE |
| ASP | 259 | 39 | 1 | FALSE | 60 | 4 | FALSE |
| ALA | 260 | 39 | 1 | FALSE | 59 | 4 | FALSE |
| LEU | 261 | 36 | 1 | FALSE | 57 | 4 | FALSE |
| PRO | 262 | | | | | | |
| GLU | 263 | 16 | 0 | TRUE | 16 | 0 | TRUE |
| ALA | 264 | 33 | 1 | TRUE | 33 | 1 | TRUE |
| LEU | 265 | 23 | 2 | TRUE | 25 | 3 | TRUE |
| ALA | 266 | <12 | 3 | TRUE | 15 | 9 | TRUE |
| LEU | 267 | 23 | 4 | TRUE | 21 | 10 | TRUE |
| VAL | 268 | 25 | 6 | TRUE | 24 | 12 | TRUE |
| LYS | 269 | 23 | 8 | TRUE | 15 | 6 | TRUE |
| SER | 270 | 44 | 0 | TRUE | 50 | 36 | TRUE |
| GLY | 271 | 24 | 3 | FALSE | 26 | 4 | FALSE |
| ALA | 272 | 23 | 3 | FALSE | 26 | 4 | FALSE |
| LEU | 273 | 19 | 2 | FALSE | 18 | 1 | FALSE |
| ALA | 274 | 22 | 2 | FALSE | 20 | 1 | FALSE |
| GLY | 275 | 36 | 0 | TRUE | >50 | 13 | TRUE |

| | | | | | | | |
|---|---|---|---|---|---|---|---|
| THR | 276 | 38 | 1 | TRUE | >50 | 13 | TRUE |
| VAL | 277 | >47 | 11 | TRUE | >47 | 23 | TRUE |
| LEU | 278 | 16 | 0 | TRUE | 18 | 2 | TRUE |
| ASN | 279 | <15 | 3 | TRUE | 21 | 1 | TRUE |
| ASP | 280 | 34 | 0 | TRUE | 33 | 1 | TRUE |
| ALA | 281 | <12 | 6 | TRUE | 20 | 0 | TRUE |
| ASN | 282 | 27 | 2 | FALSE | 36 | 2 | FALSE |
| ASN | 283 | 28 | 2 | FALSE | 38 | 2 | FALSE |
| GLN | 284 | 26 | 2 | FALSE | 36 | 2 | FALSE |
| ALA | 285 | 49 | 2 | FALSE | 45 | 0 | FALSE |
| LYS | 286 | 48 | 2 | FALSE | 44 | 0 | FALSE |
| ALA | 287 | 49 | 2 | FALSE | 45 | 0 | FALSE |
| THR | 288 | 48 | 2 | FALSE | 44 | 0 | FALSE |
| PHE | 289 | >49 | 15 | TRUE | >49 | 33 | TRUE |
| ASP | 290 | 47 | 20 | TRUE | 38 | 0 | TRUE |
| LEU | 291 | >46 | 8 | TRUE | >46 | 3 | TRUE |
| ALA | 292 | 53 | 6 | FALSE | 57 | 3 | FALSE |
| LYS | 293 | 54 | 6 | FALSE | 58 | 3 | FALSE |
| ASN | 294 | >53 | 11 | TRUE | >53 | 6 | TRUE |
| LEU | 295 | 37 | 2 | TRUE | 37 | 1 | TRUE |
| ALA | 296 | 18 | 3 | FALSE | 21 | 3 | FALSE |
| ASP | 297 | 20 | 3 | FALSE | 23 | 3 | FALSE |
| GLY | 298 | 21 | 0 | FALSE | 19 | 0 | FALSE |
| LYS | 299 | 23 | 0 | FALSE | 21 | 0 | FALSE |
| GLY | 300 | 23 | 1 | TRUE | 23 | 1 | TRUE |
| ALA | 301 | <14 | 4 | TRUE | <14 | 6 | TRUE |
| ALA | 302 | 30 | 5 | FALSE | 41 | 4 | FALSE |
| ASP | 303 | 31 | 5 | FALSE | 41 | 4 | FALSE |
| GLY | 304 | 23 | 2 | FALSE | 38 | 2 | FALSE |
| THR | 305 | 25 | 2 | FALSE | 39 | 2 | FALSE |
| ASN | 306 | 16 | 1 | FALSE | 17 | 0 | FALSE |

| | | | | | | | |
|---|---|---|---|---|---|---|---|
| TRP | 307 | 12 | 1 | FALSE | 13 | 0 | FALSE |
| LYS | 308 | 11 | 1 | FALSE | 12 | 0 | FALSE |
| ILE | 309 | 9 | 1 | FALSE | 10 | 0 | FALSE |
| ASP | 310 | 11 | 1 | FALSE | 13 | 0 | FALSE |
| ASN | 311 | <15 | 3 | TRUE | <15 | 5 | TRUE |
| LYS | 312 | <15 | 6 | TRUE | 17 | 5 | TRUE |
| VAL | 313 | 26 | 6 | TRUE | 20 | 1 | TRUE |
| VAL | 314 | 35 | 1 | TRUE | >45 | 4 | TRUE |
| ARG | 315 | 29 | 9 | FALSE | 58 | 6 | FALSE |
| VAL | 316 | 26 | 9 | FALSE | 56 | 6 | FALSE |
| PRO | 317 | | | | | | |
| TYR | 318 | 18 | 2 | FALSE | 18 | 1 | FALSE |
| VAL | 319 | 17 | 2 | FALSE | 17 | 1 | FALSE |
| GLY | 320 | 20 | 2 | FALSE | 19 | 1 | FALSE |
| VAL | 321 | 12 | 2 | FALSE | 12 | 2 | FALSE |
| ASP | 322 | 15 | 2 | FALSE | 15 | 2 | FALSE |
| LYS | 323 | 35 | 1 | TRUE | 38 | 0 | TRUE |
| ASP | 324 | 24 | 3 | TRUE | 25 | 1 | TRUE |
| ASN | 325 | 24 | 0 | TRUE | 22 | 1 | TRUE |
| LEU | 326 | <12 | 2 | TRUE | <12 | 5 | TRUE |
| ALA | 327 | 9 | 2 | FALSE | 14 | 0 | FALSE |
| GLU | 328 | 10 | 2 | FALSE | 14 | 0 | FALSE |
| PHE | 329 | <11 | 3 | TRUE | 24 | 0 | TRUE |
| SER | 330 | 17 | 2 | FALSE | <15 | 1 | FALSE |
| LYS | 331 | 16 | 2 | FALSE | <15 | 1 | FALSE |
| LYS | 332 | 5 | 2 | FALSE | <4 | 1 | FALSE |

**Table S12. $\Delta G_{op}$ values for RbsB**

All values in kJ/mol.

| Res. name | Res. number | Apo | Std Dev | Single resolved | RIB | Std Dev | Single resolved |
|---|---|---|---|---|---|---|---|
| 1 | MET | | | | | | |
| 2 | ASN | | | | | | |
| 3 | MET | | | | | | |
| 4 | LYS | | | | | | |
| 5 | LYS | | | | | | |
| 6 | LEU | | | | | | |
| 7 | ALA | | | | | | |
| 8 | THR | | | | | | |
| 9 | LEU | | | | | | |
| 10 | VAL | | | | | | |
| 11 | SER | | | | | | |
| 12 | ALA | | | | | | |
| 13 | VAL | | | | | | |
| 14 | ALA | | | | | | |
| 15 | LEU | 41 | 1 | FALSE | 41 | 1 | FALSE |
| 16 | SER | 45 | 1 | FALSE | 45 | 1 | FALSE |
| 17 | ALA | 46 | 1 | FALSE | 45 | 1 | FALSE |
| 18 | THR | 43 | 1 | FALSE | 43 | 1 | FALSE |
| 19 | VAL | 41 | 1 | FALSE | 41 | 1 | FALSE |
| 20 | SER | 45 | 1 | FALSE | 45 | 1 | FALSE |
| 21 | ALA | | | | | | |
| 22 | ASN | | | | | | |
| 23 | ALA | | | | | | |
| 24 | MET | | | | | | |
| 25 | ALA | | | | | | |
| 26 | LYS | | | | | | |
| 27 | ASP | | | | | | |

| | | | | | | | |
|---|---|---|---|---|---|---|---|
| 28 | THR | 38 | 1 | FALSE | 27 | 1 | FALSE |
| 29 | ILE | 36 | 1 | FALSE | 26 | 1 | FALSE |
| 30 | ALA | 38 | 1 | FALSE | 28 | 1 | FALSE |
| 31 | LEU | 36 | 1 | FALSE | 26 | 1 | FALSE |
| 32 | VAL | | | | | | |
| 33 | VAL | >43 | 16 | TRUE | >43 | 6 | TRUE |
| 34 | SER | 31 | 2 | FALSE | 43 | 3 | FALSE |
| 35 | THR | 31 | 2 | FALSE | 43 | 3 | FALSE |
| 36 | LEU | 28 | 2 | FALSE | 39 | 3 | FALSE |
| 37 | ASN | <14 | 6 | TRUE | 18 | 8 | TRUE |
| 38 | ASN | <17 | 4 | TRUE | <17 | 4 | TRUE |
| 39 | PRO | | | | | | |
| 40 | PHE | <10 | 1 | TRUE | 19 | 1 | TRUE |
| 41 | PHE | 34 | 0 | TRUE | 36 | 1 | TRUE |
| 42 | VAL | 26 | 1 | FALSE | 9 | 3 | FALSE |
| 43 | SER | 31 | 1 | FALSE | 14 | 3 | FALSE |
| 44 | LEU | 34 | 2 | TRUE | 35 | 1 | TRUE |
| 45 | LYS | >46 | 4 | TRUE | >46 | 7 | TRUE |
| 46 | ASP | >49 | 12 | TRUE | 49 | 21 | TRUE |
| 47 | GLY | 53 | 4 | FALSE | >46 | 1 | FALSE |
| 48 | ALA | 55 | 4 | FALSE | >48 | 1 | FALSE |
| 49 | GLN | 27 | 1 | TRUE | 30 | 3 | TRUE |
| 50 | LYS | 51 | 4 | FALSE | 37 | 1 | FALSE |
| 51 | GLU | 51 | 4 | FALSE | 36 | 1 | FALSE |
| 52 | ALA | 27 | 2 | FALSE | 37 | 3 | FALSE |
| 53 | ASP | 28 | 2 | FALSE | 38 | 3 | FALSE |
| 54 | LYS | 26 | 2 | FALSE | 36 | 3 | FALSE |
| 55 | LEU | 37 | 2 | TRUE | 32 | 2 | TRUE |
| 56 | GLY | 18 | 12 | TRUE | 23 | 6 | TRUE |
| 57 | TYR | <12 | 1 | FALSE | 17 | 2 | FALSE |
| 58 | ASN | <16 | 1 | FALSE | 20 | 2 | FALSE |

| | | | | | | | |
|---|---|---|---|---|---|---|---|
| 59 | LEU | 25 | 9 | TRUE | <11 | 3 | TRUE |
| 60 | VAL | | | | | | |
| 61 | VAL | 9 | 4 | TRUE | <8 | 1 | TRUE |
| 62 | LEU | 19 | 12 | TRUE | 32 | 1 | TRUE |
| 63 | ASP | >47 | 16 | TRUE | 45 | 21 | TRUE |
| 64 | SER | 19 | 14 | TRUE | <14 | 4 | TRUE |
| 65 | GLN | 37 | 2 | TRUE | >49 | 5 | TRUE |
| 66 | ASN | 32 | 3 | TRUE | 47 | 21 | TRUE |
| 67 | ASN | 35 | 5 | TRUE | 37 | 5 | TRUE |
| 68 | PRO | | | | | | |
| 69 | ALA | 10 | 2 | FALSE | 11 | 2 | FALSE |
| 70 | LYS | 11 | 2 | FALSE | 12 | 2 | FALSE |
| 71 | GLU | 35 | 1 | TRUE | >47 | 18 | TRUE |
| 72 | LEU | >43 | 20 | TRUE | 36 | 4 | TRUE |
| 73 | ALA | >46 | 4 | TRUE | >46 | 3 | TRUE |
| 74 | ASN | 34 | 0 | TRUE | 36 | 0 | TRUE |
| 75 | VAL | 33 | 3 | TRUE | 36 | 6 | TRUE |
| 76 | GLN | 34 | 3 | TRUE | 33 | 3 | TRUE |
| 77 | ASP | <15 | 6 | TRUE | <15 | 5 | TRUE |
| 78 | LEU | 15 | 1 | TRUE | 16 | 0 | TRUE |
| 79 | THR | <11 | 3 | TRUE | <11 | 2 | TRUE |
| 80 | VAL | 18 | 0 | TRUE | 16 | 1 | TRUE |
| 81 | ARG | <13 | 3 | TRUE | <13 | 4 | TRUE |
| 82 | GLY | <14 | 1 | FALSE | <14 | 1 | FALSE |
| 83 | THR | <13 | 1 | FALSE | <13 | 1 | FALSE |
| 84 | LYS | <14 | 3 | TRUE | 27 | 26 | TRUE |
| 85 | ILE | 43 | 25 | TRUE | >44 | 4 | TRUE |
| 86 | LEU | 37 | 2 | TRUE | >43 | 28 | TRUE |
| 87 | LEU | 23 | 1 | TRUE | 24 | 0 | TRUE |
| 88 | ILE | >42 | 5 | TRUE | >42 | 3 | TRUE |
| 89 | ASN | 37 | 1 | TRUE | 40 | 0 | TRUE |

| | | | | | | | |
|---|---|---|---|---|---|---|---|
| 90 | PRO | 0 | 0 | FALSE | 0 | 0 | FALSE |
| 91 | THR | 11 | 1 | FALSE | 27 | 2 | FALSE |
| 92 | ASP | 14 | 1 | FALSE | 30 | 2 | FALSE |
| 93 | SER | 13 | 1 | FALSE | 29 | 2 | FALSE |
| 94 | ASP | 15 | 1 | FALSE | 31 | 2 | FALSE |
| 95 | ALA | <12 | 2 | TRUE | 13 | 7 | TRUE |
| 96 | VAL | <9 | 1 | FALSE | <9 | 1 | FALSE |
| 97 | GLY | <12 | 1 | FALSE | <12 | 1 | FALSE |
| 98 | ASN | <16 | 4 | TRUE | <16 | 6 | TRUE |
| 99 | ALA | <15 | 4 | TRUE | <15 | 7 | TRUE |
| 100 | VAL | 27 | 1 | FALSE | 27 | 0 | FALSE |
| 101 | LYS | 30 | 1 | FALSE | 29 | 0 | FALSE |
| 102 | MET | 30 | 0 | TRUE | 32 | 0 | TRUE |
| 103 | ALA | 26 | 0 | TRUE | 26 | 0 | TRUE |
| 104 | ASN | 33 | 1 | TRUE | 32 | 0 | TRUE |
| 105 | GLN | 31 | 0 | FALSE | 31 | 0 | FALSE |
| 106 | ALA | 30 | 0 | FALSE | 30 | 0 | FALSE |
| 107 | ASN | 22 | 0 | TRUE | 22 | 0 | TRUE |
| 108 | ILE | 35 | 1 | TRUE | 36 | 1 | TRUE |
| 109 | PRO | | | | | | |
| 110 | VAL | 56 | 2 | FALSE | 50 | 2 | FALSE |
| 111 | ILE | 56 | 2 | FALSE | 50 | 2 | FALSE |
| 112 | THR | 59 | 2 | FALSE | 53 | 2 | FALSE |
| 113 | LEU | >45 | 4 | TRUE | >45 | 5 | TRUE |
| 114 | ASP | 48 | 4 | FALSE | 50 | 5 | FALSE |
| 115 | ARG | 48 | 4 | FALSE | 50 | 5 | FALSE |
| 116 | GLN | <14 | 3 | TRUE | <14 | 6 | TRUE |
| 117 | ALA | 36 | 0 | TRUE | 40 | 0 | TRUE |
| 118 | THR | <12 | 3 | TRUE | 14 | 0 | FALSE |
| 119 | LYS | 11 | 0 | FALSE | 15 | 0 | FALSE |
| 120 | GLY | 11 | 0 | FALSE | 15 | 0 | FALSE |

| | | | | | | | |
|---|---|---|---|---|---|---|---|
| 121 | GLU | 11 | 0 | FALSE | 14 | 0 | FALSE |
| 122 | VAL | 6 | 0 | FALSE | 9 | 0 | FALSE |
| 123 | VAL | 6 | 0 | FALSE | 9 | 0 | FALSE |
| 124 | SER | 12 | 0 | FALSE | 15 | 0 | FALSE |
| 125 | HIS | 13 | 0 | FALSE | 16 | 0 | FALSE |
| 126 | ILE | 51 | 5 | FALSE | 36 | 1 | FALSE |
| 127 | ALA | 52 | 5 | FALSE | 37 | 1 | FALSE |
| 128 | SER | 34 | 0 | TRUE | 39 | 1 | TRUE |
| 129 | ASP | <15 | 4 | TRUE | 22 | 0 | TRUE |
| 130 | ASN | <15 | 4 | TRUE | <15 | 5 | TRUE |
| 131 | VAL | 16 | 5 | TRUE | 20 | 5 | TRUE |
| 132 | LEU | 33 | 1 | FALSE | 40 | 2 | FALSE |
| 133 | GLY | 35 | 1 | FALSE | 42 | 2 | FALSE |
| 134 | GLY | 38 | 1 | FALSE | 44 | 2 | FALSE |
| 135 | LYS | 37 | 1 | FALSE | 44 | 2 | FALSE |
| 136 | ILE | 32 | 1 | TRUE | 34 | 3 | TRUE |
| 137 | ALA | 46 | 20 | TRUE | 44 | 20 | TRUE |
| 138 | GLY | >47 | 4 | TRUE | >47 | 5 | TRUE |
| 139 | ASP | 39 | 3 | TRUE | >49 | 21 | TRUE |
| 140 | TYR | 34 | 1 | TRUE | 41 | 2 | TRUE |
| 141 | ILE | >44 | 12 | TRUE | >44 | 6 | TRUE |
| 142 | ALA | 22 | 0 | TRUE | 21 | 1 | TRUE |
| 143 | LYS | 31 | 1 | TRUE | 30 | 2 | TRUE |
| 144 | LYS | 21 | 2 | TRUE | 21 | 1 | TRUE |
| 145 | ALA | 23 | 1 | FALSE | 27 | 1 | FALSE |
| 146 | GLY | 22 | 1 | FALSE | 27 | 1 | FALSE |
| 147 | GLU | 23 | 1 | FALSE | 27 | 1 | FALSE |
| 148 | GLY | 21 | 1 | FALSE | 26 | 1 | FALSE |
| 149 | ALA | 23 | 1 | FALSE | 28 | 1 | FALSE |
| 150 | LYS | >47 | 5 | TRUE | 45 | 24 | TRUE |
| 151 | VAL | 45 | 3 | FALSE | 45 | 4 | FALSE |

| | | | | | | | |
|---|---|---|---|---|---|---|---|
| 152 | ILE | 43 | 3 | FALSE | 43 | 4 | FALSE |
| 153 | GLU | 46 | 3 | FALSE | 46 | 4 | FALSE |
| 154 | LEU | >43 | 6 | TRUE | >43 | 5 | TRUE |
| 155 | GLN | >47 | 4 | TRUE | >47 | 4 | TRUE |
| 156 | GLY | >48 | 20 | TRUE | >48 | 14 | TRUE |
| 157 | ILE | 12 | 2 | FALSE | 30 | 3 | FALSE |
| 158 | ALA | 14 | 2 | FALSE | 32 | 3 | FALSE |
| 159 | GLY | 15 | 2 | FALSE | 33 | 3 | FALSE |
| 160 | THR | 16 | 2 | FALSE | 34 | 3 | FALSE |
| 161 | SER | 13 | 2 | FALSE | 19 | 5 | FALSE |
| 162 | ALA | 11 | 2 | FALSE | 18 | 5 | FALSE |
| 163 | ALA | 34 | 2 | TRUE | >47 | 6 | TRUE |
| 164 | ARG | 25 | 4 | TRUE | 35 | 1 | TRUE |
| 165 | GLU | 25 | 6 | TRUE | 27 | 10 | TRUE |
| 166 | ARG | >47 | 22 | TRUE | >47 | 5 | TRUE |
| 167 | GLY | 36 | 1 | TRUE | >48 | 4 | TRUE |
| 168 | GLU | 23 | 7 | TRUE | 32 | 8 | TRUE |
| 169 | GLY | 34 | 1 | TRUE | 46 | 19 | TRUE |
| 170 | PHE | 36 | 2 | FALSE | 49 | 2 | FALSE |
| 171 | GLN | 37 | 2 | FALSE | 50 | 2 | FALSE |
| 172 | GLN | 38 | 2 | FALSE | 50 | 2 | FALSE |
| 173 | ALA | 31 | 1 | TRUE | 32 | 4 | TRUE |
| 174 | VAL | 30 | 1 | TRUE | 30 | 1 | TRUE |
| 175 | ALA | 25 | 5 | TRUE | 37 | 2 | TRUE |
| 176 | ALA | 33 | 2 | FALSE | 27 | 3 | FALSE |
| 177 | HIS | 33 | 2 | FALSE | 28 | 3 | FALSE |
| 178 | LYS | 20 | 0 | TRUE | 23 | 2 | TRUE |
| 179 | PHE | 29 | 3 | TRUE | 30 | 0 | TRUE |
| 180 | ASN | 26 | 6 | FALSE | 48 | 8 | FALSE |
| 181 | VAL | 21 | 6 | FALSE | 43 | 8 | FALSE |
| 182 | LEU | 17 | 11 | TRUE | 26 | 1 | TRUE |

| | | | | | | | |
|---|---|---|---|---|---|---|---|
| 183 | ALA | 23 | 4 | TRUE | 36 | 2 | TRUE |
| 184 | SER | 29 | 2 | TRUE | >49 | 3 | TRUE |
| 185 | GLN | 30 | 0 | FALSE | 43 | 2 | FALSE |
| 186 | PRO | | | | | | |
| 187 | ALA | 27 | 0 | FALSE | 40 | 2 | FALSE |
| 188 | ASP | 29 | 0 | FALSE | 42 | 2 | FALSE |
| 189 | PHE | 26 | 0 | FALSE | 39 | 2 | FALSE |
| 190 | ASP | 24 | 0 | TRUE | 34 | 2 | TRUE |
| 191 | ARG | <12 | 1 | TRUE | <12 | 3 | TRUE |
| 192 | ILE | 18 | 1 | FALSE | 19 | 0 | FALSE |
| 193 | LYS | 19 | 1 | FALSE | 21 | 0 | FALSE |
| 194 | GLY | 25 | 1 | TRUE | 46 | 20 | TRUE |
| 195 | LEU | 21 | 3 | TRUE | 33 | 2 | TRUE |
| 196 | ASN | 26 | 2 | TRUE | 35 | 4 | TRUE |
| 197 | VAL | 25 | 1 | TRUE | 33 | 5 | TRUE |
| 198 | MET | 28 | 0 | TRUE | >47 | 17 | TRUE |
| 199 | GLN | 28 | 0 | TRUE | 28 | 0 | TRUE |
| 200 | ASN | <17 | 9 | TRUE | <17 | 5 | TRUE |
| 201 | LEU | 24 | 0 | TRUE | 25 | 1 | TRUE |
| 202 | LEU | 29 | 1 | TRUE | 30 | 0 | TRUE |
| 203 | THR | <11 | 3 | TRUE | <11 | 3 | TRUE |
| 204 | ALA | <14 | 3 | TRUE | <14 | 6 | TRUE |
| 205 | HIS | 29 | 3 | TRUE | 29 | 3 | TRUE |
| 206 | PRO | | | | | | |
| 207 | ASP | 28 | 5 | TRUE | 24 | 9 | TRUE |
| 208 | VAL | 28 | 1 | TRUE | 29 | 2 | TRUE |
| 209 | GLN | 18 | 0 | TRUE | 24 | 1 | TRUE |
| 210 | ALA | 52 | 10 | FALSE | 46 | 11 | FALSE |
| 211 | VAL | 47 | 10 | FALSE | 41 | 11 | FALSE |
| 212 | PHE | >45 | 4 | TRUE | >45 | 3 | TRUE |
| 213 | ALA | 49 | 5 | FALSE | 51 | 6 | FALSE |

| | | | | | | | |
|---|---|---|---|---|---|---|---|
| 214 | GLN | 49 | 5 | FALSE | 51 | 6 | FALSE |
| 215 | ASN | 59 | 2 | FALSE | 52 | 8 | FALSE |
| 216 | ASP | 57 | 2 | FALSE | 50 | 8 | FALSE |
| 217 | GLU | 21 | 4 | TRUE | 37 | 2 | TRUE |
| 218 | MET | >47 | 1 | FALSE | >47 | 1 | FALSE |
| 219 | ALA | >48 | 1 | FALSE | >48 | 1 | FALSE |
| 220 | LEU | >44 | 1 | FALSE | >44 | 1 | FALSE |
| 221 | GLY | >46 | 23 | TRUE | >46 | 20 | TRUE |
| 222 | ALA | 35 | 2 | TRUE | 37 | 5 | TRUE |
| 223 | LEU | 39 | 4 | TRUE | 42 | 22 | TRUE |
| 224 | ARG | 18 | 0 | TRUE | 18 | 0 | TRUE |
| 225 | ALA | 38 | 0 | TRUE | 41 | 1 | TRUE |
| 226 | LEU | 32 | 0 | TRUE | 33 | 1 | TRUE |
| 227 | GLN | 20 | 0 | TRUE | 21 | 1 | TRUE |
| 228 | THR | <14 | 3 | TRUE | <14 | 4 | TRUE |
| 229 | ALA | 13 | 1 | FALSE | <14 | 3 | TRUE |
| 230 | GLY | 11 | 1 | FALSE | 11 | 1 | FALSE |
| 231 | LYS | 12 | 1 | FALSE | 12 | 1 | FALSE |
| 232 | SER | <16 | 3 | TRUE | 14 | 1 | FALSE |
| 233 | ASP | <15 | 4 | TRUE | <15 | 4 | TRUE |
| 234 | VAL | <8 | 1 | TRUE | 9 | 7 | TRUE |
| 235 | MET | 24 | 0 | TRUE | 24 | 0 | TRUE |
| 236 | VAL | 23 | 1 | TRUE | 23 | 1 | TRUE |
| 237 | VAL | 41 | 22 | TRUE | >43 | 18 | TRUE |
| 238 | GLY | >46 | 4 | TRUE | 45 | 22 | TRUE |
| 239 | PHE | >47 | 3 | TRUE | >47 | 5 | TRUE |
| 240 | ASP | 25 | 3 | FALSE | 55 | 4 | FALSE |
| 241 | GLY | 23 | 3 | FALSE | 52 | 4 | FALSE |
| 242 | THR | 17 | 0 | FALSE | 21 | 0 | FALSE |
| 243 | PRO | | | | | | |
| 244 | ASP | 16 | 0 | FALSE | 20 | 0 | FALSE |

| | | | | | | | |
|---|---|---|---|---|---|---|---|
| 245 | GLY | 15 | 0 | FALSE | 19 | 0 | FALSE |
| 246 | GLU | 17 | 0 | FALSE | 21 | 0 | FALSE |
| 247 | LYS | 15 | 0 | FALSE | 19 | 0 | FALSE |
| 248 | ALA | 23 | 0 | FALSE | 24 | 0 | FALSE |
| 249 | VAL | 19 | 0 | FALSE | 19 | 0 | FALSE |
| 250 | ASN | 16 | 7 | TRUE | 26 | 3 | TRUE |
| 251 | ASP | 16 | 8 | TRUE | 24 | 2 | TRUE |
| 252 | GLY | 11 | 1 | FALSE | 14 | 8 | TRUE |
| 253 | LYS | 13 | 1 | FALSE | 24 | 1 | FALSE |
| 254 | LEU | 10 | 1 | FALSE | 21 | 1 | FALSE |
| 255 | ALA | 35 | 3 | FALSE | 49 | 4 | FALSE |
| 256 | ALA | 36 | 3 | FALSE | 50 | 4 | FALSE |
| 257 | THR | 36 | 3 | FALSE | 50 | 4 | FALSE |
| 258 | ILE | >45 | 4 | TRUE | >45 | 1 | TRUE |
| 259 | ALA | >46 | 31 | TRUE | 41 | 36 | TRUE |
| 260 | GLN | 25 | 1 | TRUE | 36 | 0 | TRUE |
| 261 | LEU | 18 | 1 | FALSE | 20 | 0 | FALSE |
| 262 | PRO | | | | | | |
| 263 | ASP | 37 | 1 | FALSE | 42 | 2 | FALSE |
| 264 | GLN | 37 | 1 | FALSE | 42 | 2 | FALSE |
| 265 | ILE | 35 | 1 | FALSE | 40 | 2 | FALSE |
| 266 | GLY | 36 | 1 | FALSE | 41 | 2 | FALSE |
| 267 | ALA | 24 | 0 | TRUE | 44 | 2 | FALSE |
| 268 | LYS | <13 | 3 | TRUE | 26 | 0 | TRUE |
| 269 | GLY | 41 | 0 | FALSE | 38 | 0 | FALSE |
| 270 | VAL | 37 | 0 | FALSE | 35 | 0 | FALSE |
| 271 | GLU | 46 | 23 | TRUE | >46 | 4 | TRUE |
| 272 | THR | >46 | 1 | FALSE | >46 | 1 | FALSE |
| 273 | ALA | >49 | 1 | FALSE | >49 | 1 | FALSE |
| 274 | ASP | >48 | 1 | FALSE | >48 | 1 | FALSE |
| 275 | LYS | >46 | 1 | FALSE | >46 | 1 | FALSE |

| | | | | | | | |
|---|---|---|---|---|---|---|---|
| 276 | VAL | 41 | 19 | TRUE | >44 | 6 | TRUE |
| 277 | LEU | 29 | 3 | TRUE | 25 | 1 | TRUE |
| 278 | LYS | 23 | 0 | FALSE | 21 | 1 | FALSE |
| 279 | GLY | 24 | 0 | FALSE | 22 | 1 | FALSE |
| 280 | GLU | 14 | 1 | FALSE | 13 | 1 | FALSE |
| 281 | LYS | 13 | 1 | FALSE | 11 | 1 | FALSE |
| 282 | VAL | 11 | 1 | FALSE | 9 | 1 | FALSE |
| 283 | GLN | 19 | 2 | TRUE | 20 | 1 | TRUE |
| 284 | ALA | <14 | 3 | TRUE | <14 | 4 | TRUE |
| 285 | LYS | 12 | 7 | TRUE | 21 | 1 | TRUE |
| 286 | TYR | 22 | 0 | TRUE | 19 | 1 | TRUE |
| 287 | PRO | | | | | | |
| 288 | VAL | 10 | 4 | TRUE | 11 | 6 | TRUE |
| 289 | ASP | 13 | 8 | TRUE | <13 | 7 | TRUE |
| 290 | LEU | 30 | 0 | TRUE | 30 | 0 | TRUE |
| 291 | LYS | 24 | 0 | TRUE | 17 | 9 | TRUE |
| 292 | LEU | <10 | 5 | TRUE | <10 | 5 | TRUE |
| 293 | VAL | 9 | 6 | TRUE | 10 | 4 | TRUE |
| 294 | VAL | 18 | 1 | FALSE | 18 | 0 | FALSE |
| 295 | LYS | 21 | 1 | FALSE | 21 | 0 | FALSE |
| 296 | GLN | 13 | 1 | FALSE | 13 | 0 | FALSE |

**Table S13. Rosetta metrics for LacI designs**

For metric definitions, see **Methods: Metastable helix stability calculation**. Mutations to the wild-type sequence are shown in red.

| Design | Metastable helix sequence | Additional mutations | Interface energy | Helix energy | Cross hbond count | Unfavorable energy sum | Metastable helix stability score |
|---|---|---|---|---|---|---|---|
| WT | EDSSCYI | – | -51.53 | -40.61 | 13 | 0.23 | 2.87 |
| 1 | EESRYTI | S221T, M223E | -44.44 | -39.64 | 15 | 0.84 | 1.76 |
| 2 | PEAPTTI | None | -33.65 | -42.93 | 11 | 0.44 | 2.61 |
| 3 | PEAGMYI | None | -47.86 | -41.66 | 11 | 0.59 | 2.58 |
| 4 | PEAPMYI | None | -46.38 | -49.47 | 12 | 0.44 | 3.01 |
| 5 | ELAACAI | None | -46.65 | -31.28 | 9 | 0.10 | 3.58 |
| 6 | PWAQMTI | None | -40.33 | -43.40 | 9 | 0.12 | 4.15 |

**Table S14. Hill equation constants for LacI designs**

| Design | $G_0$ | $G_\infty$ | $EC_{50}$ | n | $G_0$ error | $G_\infty$ error | $EC_{50}$ error | n error |
|---|---|---|---|---|---|---|---|---|
| WT | 3348.34 | 8366.42 | 24.49 | 1.02 | 153.18 | 257.40 | 5.624 | 0.21 |
| 1 | 4430.90 | 8675.71 | 0.34 | 1.00 | 386.46 | 286.19 | 0.16 | 0.42 |
| 2 | 5447.80 | 12480.82 | 4.14 | 0.64 | 386.71 | 531.05 | 1.91 | 0.17 |
| 3 | 3064.36 | 8635.67 | 15.32 | 0.84 | 516.08 | 706.31 | 10.20 | 0.42 |
| 4 | 3992.73 | 7018.66 | 71.47 | 1.90 | 175.50 | 297.39 | 23.43 | 1.00 |
| 5 | 3952.40 | 7129.55 | 83.10 | 1.15 | 213.54 | 462.02 | 45.10 | 0.61 |
| 6 | 3695.15 | 7999.87 | 543.30 | 0.99 | 149.65 | 736.72 | 276.38 | 0.36 |

Curves were fit as in Tack *et al.*[15] per the following definition of the Hill equation:

$$G(L) = G_0 + \frac{G_\infty - G_0}{1 + (\frac{EC_{50}}{L})^n}$$

where $G_0$ is the basal gene expression in the absence of ligand, $G_\infty$ is the gene expression at saturating ligand concentrations, $EC_{50}$ is the concentration of ligand that results in gene expression halfway between $G_\infty$ and $G_0$, and the Hill coefficient, $n$, represents the steepness of the curve. L, the ligand concentration, is the independent variable.

#### Appendix 1. DNA operator sequences used in this study

Operator sequences are highlighted in yellow. Each operator sequence is repeated twice. The *scrO* sequence results from scrambling the *lacO1* sequence.

| Operator | DNA sequence |
|---|---|
| 2X-*lacO1* | AGTAGTggAATTGTGAGCGGATAACAATTGACATTGTGAGCGGATAACAAGATACTGAGC<br>GCTCAGTATCTTGTTATCCGCTCACAATGTCAATTGTTATCCGCTCACAATTccACTACT |
| 2X-*galO* | AATTAGTGGAATCGTTTACACAAGAAAATTAGTGGAATCGTTTACACAAGAA<br>TTCTTGTGTAAACGATTCCACTAATTTTCTTGTGTAAACGATTCCACTAATT |
| 2X-*rbsO* | GTGGGTCAGCGAAACGTTTCGCTGATGGAGGTGGGTCAGCGAAACGTTTCGCTGATGGAG<br>CTCCATCAGCGAAACGTTTCGCTGACCCACCTCCATCAGCGAAACGTTTCGCTGACCCAC |
| 2X-*scrO* | GCATGGGATAAGTGATGACAATTTATGGAAAACCTGACTGCAGTTATAAGCGGAGAAGTG<br>CACTTCTCCGCTTATAACTGCAGTCAGGTTTTCCATAAATTGTCATCACTTATCCCATGC |

#### Appendix 2. DNA sequences for TF and PBP genes

| Protein | DNA sequence |
|---|---|
| 6X-His-TEV-LacI | ATGCATCACCATCACCATCACGAAAACTTATATTTCCAATCTAAACCAGTAACGTTATA<br>CGATGTCGCAGAGTATGCCGGTGTCTCTTATCAGACCGTTTCCCGCGTGGTGAACCAGG<br>CCAGCCACGTTTCTGCGAAAACGCGGGAAAAAGTGGAAGCGGCGATGGCGGAGCTGAAT<br>TACATTCCCAACCGCGTGGCACAACAACTGGCGGGCAAACAGTCGTTGCTGATTGGCGT<br>TGCCACCTCCAGTCTGGCCCTGCACGCGCCGTCGCAAATTGTCGCGGCGATTAAATCTC<br>GCGCCGATCAACTGGGTGCCAGCGTGGTGGTGTCGATGGTAGAACGAAGCGGCGTCGAA<br>GCCTGTAAAGCGGCGGTGCACAATCTTCTCGCGCAACGCGTCAGTGGGCTGATCATTAA<br>CTATCCGCTGGATGACCAGGATGCCATTGCTGTGGAAGCTGCCTGCACTAATGTTCCGG<br>CGTTATTTCTTGATGTCTCTGACCAGACACCCATCAACAGTATTATTTTCTCCCATGAA<br>GACGGTACGCGACTGGGCGTGGAGCATCTGGTCGCATTGGGTCACCAGCAAATCGCGCT<br>GTTAGCGGGCCCATTAAGTTCTGTCTCGGCGCGTCTGCGTCTGGCTGGCTGGCATAAAT<br>ATCTCACTCGCAATCAAATTCAGCCGATAGCGGAACGGGAAGGCGACTGGAGTGCCATG<br>TCCGGTTTTCAACAAACCATGCAAATGCTGAATGAGGGCATCGTTCCCACTGCGATGCT<br>GGTTGCCAACGATCAGATGGCGCTGGGCGCAATGCGCGCCATTACCGAGTCCGGGCTGC<br>GCGTTGGTGCGGATATCTCGGTAGTGGGATACGACGATACCGAAGACAGCTCATGTTAT<br>ATCCCGCCGTCAACCACCATCAAACAGGATTTTCGCCTGCTGGGGCAAACCAGCGTGGA<br>CCGCTTGCTGCAACTCTCTCAGGGCCAGGCGGTGAAGGGCAATCAGCTGTTGCCCGTCT<br>CACTGGTGAAAAGAAAAACCACCCTGGCGCCCTAA |
| 6X-His-TEV-GalR | ATGCATCACCATCATCATCACGAAAACTTATATTTCCAATCTGCGACCATAAAGGATGT<br>AGCCCGACTGGCAGGCGTTTCAGTCGCCACCGTTTCCCGCGTCATTAATAATTCACCCA<br>AAGCCAGCGAAGCTTCCCGGCTGGCTGTGCATAGTGCAATGGAGTCTCTTAGCTATCAC<br>CCGAACGCCAACGCCCGTGCGCTGGCGCAGCAGACCACTGAAACGGTCGGTCTGGTCGT<br>TGGTGATGTTTCCGATCCGTTTTTCGGTGCAATGGTGAAAGCGGTCGAACAGGTGGCTT<br>ATCACACCGGTAATTTTTTATTGATTGGCAACGGTTACCACAACGAACAAAAAGAGCGT<br>CAGGCCATTGAGCAACTGATCCGCCATCGCTGTGCTGCGTTGGTCGTCCATGCCAAAAT<br>GATCCCGGATGCTGATTTAGCCTCATTAATGAAACAAATGCCCGGTATGGTGCTGATCA<br>ACCGTATCCTGCCTGGCTTTGAAAACCGTTGTATTGCTCTGGACGATCGTTACGGTGCC<br>TGGCTGGCAACGCGTCATTTAATTCAGCAAGGTCATACCCGCATTGGTTATCTGTGCTC<br>TAACCACTCTATTTCTGACGCCGAAGATCGTCTGCAAGGGTATTACGATGCCCTTGCTG<br>AAAGTGGTATTGCGGCCAATGACCGGCTGGTGACATTTGGCGAACCAGACGAAAGCGGC<br>GGCGAACAGGCAATGACCGAGCTTTTGGGACGAGGAAGAAATTTCACTGCGGTAGCCTG<br>TTATAACGATTCAATGGCGGCGGGTGCGATGGGCGTTCTCAATGATAATGGTATTGATG<br>TACCGGGTGAGATTTCGTTAATTGGCTTTGATGATGTGCTGGTGTCACGCTATGTGCGT<br>CCGCGCCTGACCACCGTGCGTTACCCAATCGTGACGATGGCGACCCAGGCTGCCGAACT<br>GGCTTTGGCGCTGGCGGATAATCGCCCTCTCCCGGAAATCACTAATGTCTTTAGTCCGA<br>CGCTGGTACGTCGTCATTCAGTGTCAACTCCGTCGCTGGAGGCAAGTCATCATGCAACC<br>AGCGACTAA |
| 6X-His-TEV-RbsR | ATGCATCACCATCATCATCACGAAAACTTATATTTCCAATCTGCTACAATGAAAGATGT<br>TGCCCGCCTGGCGGGCGTTTCTACCTCAACAGTTTCTCACGTTATCAATAAAGATCGCT<br>TCGTCAGTGAAGCGATTACCGCCAAAGTTGAAGCGGCGATTAAAGAACTCAATTACGCG<br>CCATCAGCTCTGGCGCGTAGCCTCAAACTCAATCAAACACATACCATTGGCATGTTGAT<br>CACTGCCAGTACCAATCCTTTCTATTCAGAACTGGTGCGTGGCGTTGAACGCAGCTGCT<br>TCGAACGCGGTTATAGTCTCGTCCTTTGCAATACCGAAGGCGATGAACAGCGGATGAAT<br>CGCAATCTGGAAACGCTGATGCAAAAACGCGTTGATGGCTTGCTGTTACTGTGCACCGA<br>AACGCATCAACCTTCGCGTGAAATCATGCAACGTTATCCGACAGTGCCTACTGTGATGA<br>TGGACTGGGCTCCGTTCGATGGCGACAGCGATCTTATTCAGGATAACTCGTTGCTGGGC<br>GGAGACTTAGCAACGCAATATCTGATCGATAAAGGTCATACCCGTATCGCCTGTATTAC<br>CGGCCCGCTGGATAAAACTCCGGCGCGCCTGCGGTTGGAAGGTTATCGGGCGGCGATGA |

| | |
|---|---|
| | AACGTGCGGGCCTTAACATTCCTGATGGCTATGAAGTCACTGGTGATTTTGAATTTAAC<br>GGCGGGTTTGACGCTATGCGCCAACTGCTATCACATCCGCTGCGTCCTCAGGCCGTCTT<br>TACCGGAAATGACGCTATGGCTGTTGGCGTTTACCAGGCGTTATATCAGGCAGAGTTAC<br>AGGTTCCGCAGGATATCGCGGTGATTGGCTATGACGATATCGAACTGGCAAGCTTTATG<br>ACGCCACCATTAACCACTATCCACCAACCGAAAGATGAACTGGGGGAGCTGGCGATTGA<br>TGTACTCATCCATCGGATAACCCAGCCGACCCTTCAGCAACAACGATTACAACTTACTC<br>CGATTCTGATGGAACGCGGTTCGGCTTAA |
| 6X-His-TEV-MglB | ATGCATCACCATCATCATCACGAAAACTTATATTTCCAATCTGCTGATACTCGCATTGG<br>TGTAACAATCTATAAGTACGACGATAACTTTATGTCTGTAGTGCGCAAGGCTATTGAGC<br>AAGATGCGAAAGCCGCGCCAGATGTTCAGCTGCTGATGAATGATTCTCAGAATGACCAG<br>TCCAAGCAGAACGATCAGATCGACGTATTGCTGGCGAAAGGGGTGAAGGCACTGGCAAT<br>CAACCTGGTTGACCCGGCAGCTGCGGGTACGGTGATTGAGAAAGCGCGTGGGCAAAACG<br>TGCCGGTGGTTTTCTTCAACAAAGAACCGTCTCGTAAGGCGCTGGATAGCTACGACAAA<br>GCCTACTACGTTGGCACTGACTCCAAAGAGTCCGGCATTATTCAAGGCGATTTGATTGC<br>TAAACACTGGGCGGCGAATCAGGGTTGGGATCTGAACAAAGACGGTCAGATTCAGTTCG<br>TACTGCTGAAAGGTGAACCGGGCCATCCGGATGCAGAAGCACGTACCACTTACGTGATT<br>AAAGAATTGAACGATAAAGGCATCAAAACTGAACAGTTACAGTTAGATACCGCAATGTG<br>GGACACCGCTCAGGCGAAAGATAAGATGGACGCCTGGCTGTCTGGCCCGAACGCCAACA<br>AAATCGAAGTGGTTATCGCCAACAACGATGCGATGGCAATGGGCGCGGTTGAAGCGCTG<br>AAAGCACACAACAAGTCCAGCATTCCGGTGTTTGGCGTCGATGCGCTGCCAGAAGCGCT<br>GGCGCTGGTGAAATCCGGTGCACTGGCGGCACCGTACTGAACGATGCTAACAACCAGG<br>CGAAAGCGACCTTTGATCTGGCGAAAAACCTGGCCGATGGTAAAGGTGCGGCTGATGGC<br>ACCAACTGGAAAATCGACAACAAAGTGGTCCGCGTACCTTATGTTGGCGTAGATAAAGA<br>CAACCTGGCTGAATTCAGCAAGAAATAA |
| 6X-His-TEV-RbsB | ATGCATCACCATCATCATCACGAAAACTTATATTTCCAATCTAAAGACACCATCGCGCT<br>GGTGGTCTCCACGCTTAACAACCCGTTTTTTGTATCGCTGAAAGATGGCGCGCAGAAAG<br>AGGCGGATAAACTTGGCTATAACTTGGTGGTGCTGGACTCCCAGAACAACCCGGCGAAA<br>GAGCTGGCGAACGTGCAGGACTTAACCGTTCGCGGCACAAAAATCCTGCTGATTAACCC<br>GACCGACTCCGACGCAGTGGGTAATGCTGTGAAGATGGCTAACCAAGCGAACATCCCGG<br>TTATCACTCTTGACCGCCAAGCAACGAAAGGTGAAGTGGTGAGCCACATTGCTTCTGAT<br>AACGTACTGGGCGGCAAAATCGCTGGTGATTACATCGCGAAGAAAGCGGGTGAAGGTGC<br>AAAAGTTATCGAGCTGCAAGGCATTGCTGGTACATCCGCAGCCCGTGAACGTGGCGAAG<br>GCTTCCAGCAGGCCGTTGCTGCTCACAAGTTTAATGTTCTTGCCAGCCAGCCAGCAGAT<br>TTTGATCGCATTAAAGGTTTGAACGTAATGCAGAACCTGTTGACCGCTCATCCGGATGT<br>TCAGGCTGTATTCGCGCAGAATGATGAAATGGCGCTGGGGGCGCTGCGCGCACTGCAAA<br>CTGCCGGTAAATCGGATGTGATGGTCGTCGGATTTGACGGTACACCGGATGGCGAAAAA<br>GCGGTGAATGATGGCAAACTGGCAGCGACTATCGCTCAGCTACCCGATCAGATTGGCGC<br>GAAAGGCGTCGAAACCGCAGATAAAGTGCTGAAAGGCGAGAAAGTTCAGGCTAAGTATC<br>CGGTTGATCTGAAACTGGTTGTTAAGCAGTAG |

#### References

1. Lu, C., Wells, M.L., Reckers, A. et al. Site-resolved energetic information from HX–MS experiments. *Nat Chem Biol* (2025). https://doi.org/10.1038/s41589-025-02049-1
2. Bai, Y., Milne, J. S., Mayne, L. & Englander, S. W. Primary structure effects on peptide group hydrogen exchange. *Proteins: Structure, Function, and Bioinformatics* **17**, 75–86 (1993).
3. Crooks, G. E., Hon, G., Chandonia, J.-M. & Brenner, S. E. WebLogo: A Sequence Logo Generator. *Genome Res.* **14**, 1188–1190 (2004).
4. Chen, E. A. & Porter, L. L. SSDraw: Software for generating comparative protein secondary structure diagrams. *Protein Science* **32**, e4836 (2023).
5. Zhang, C., Shine, M., Pyle, A. M. & Zhang, Y. US-align: universal structure alignments of proteins, nucleic acids, and macromolecular complexes. *Nat Methods* **19**, 1109–1115 (2022).
6. Weis, D. D. Recommendations for the Propagation of Uncertainty in Hydrogen Exchange-Mass Spectrometric Measurements. *J. Am. Soc. Mass Spectrom.* **32**, 1610–1617 (2021).
7. Stracy, M. *et al.* Transient non-specific DNA binding dominates the target search of bacterial DNA-binding proteins. *Molecular Cell* **81**, 1499-1514.e6 (2021).
8. Spronk, C. A. E. M. *et al.* Hinge-helix formation and DNA bending in various lac repressor–operator complexes. *The EMBO Journal* **18**, 6472–6480 (1999).
9. Björkman, A. J. & Mowbray, S. L. Multiple open forms of ribose-binding protein trace the path of its conformational change11Edited by R. Huber. *Journal of Molecular Biology* **279**, 651–664 (1998).
10. Björkman, A. J. *et al.* Probing protein-protein interactions. The ribose-binding protein in bacterial transport and chemotaxis. *J Biol Chem* **269**, 30206–30211 (1994).
11. Pavlovicz, R. E., Park, H. & DiMaio, F. Efficient consideration of coordinated water molecules improves computational protein-protein and protein-ligand docking discrimination.

*PLOS Computational Biology* **16**, e1008103 (2020).

12. Markiewicz, P., Kleina, L. G., Cruz, C., Ehret, S. & Miller, J. H. Genetic studies of the lac repressor. XIV. Analysis of 4000 altered Escherichia coli lac repressors reveals essential and non-essential residues, as well as 'spacers' which do not require a specific sequence. *J. Mol. Biol.* **240**, 421–433 (1994).
13. Suckow, J. *et al.* Genetic Studies of the Lac Repressor XV: 4000 Single Amino Acid Substitutions and Analysis of the Resulting Phenotypes on the Basis of the Protein Structure. *Journal of Molecular Biology* **261**, 509–523 (1996).
14. Masson, G. R. *et al.* Recommendations for performing, interpreting and reporting hydrogen deuterium exchange mass spectrometry (HDX-MS) experiments. *Nat Methods* **16**, 595–602 (2019).
15. Tack, D. S. *et al.* The genotype-phenotype landscape of an allosteric protein. *Mol Syst Biol* **17**, e10179 (2021).
