## Supplementary figures and images for "Distinct energetic blueprints diversify function of conserved protein folds"

### GalR_uptake_errors_APO_0.pdf

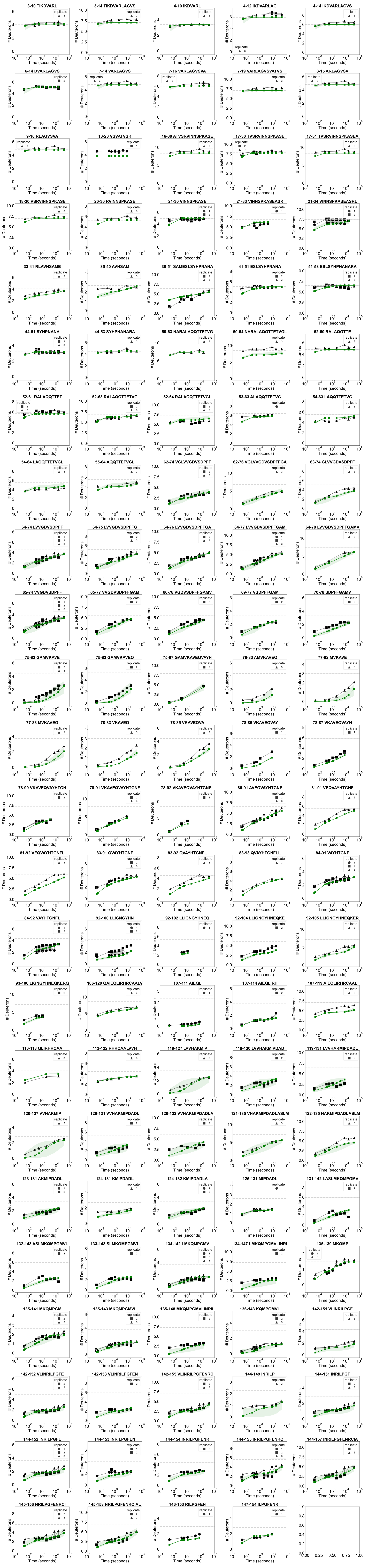

### GalR_uptake_errors_APO_1.pdf

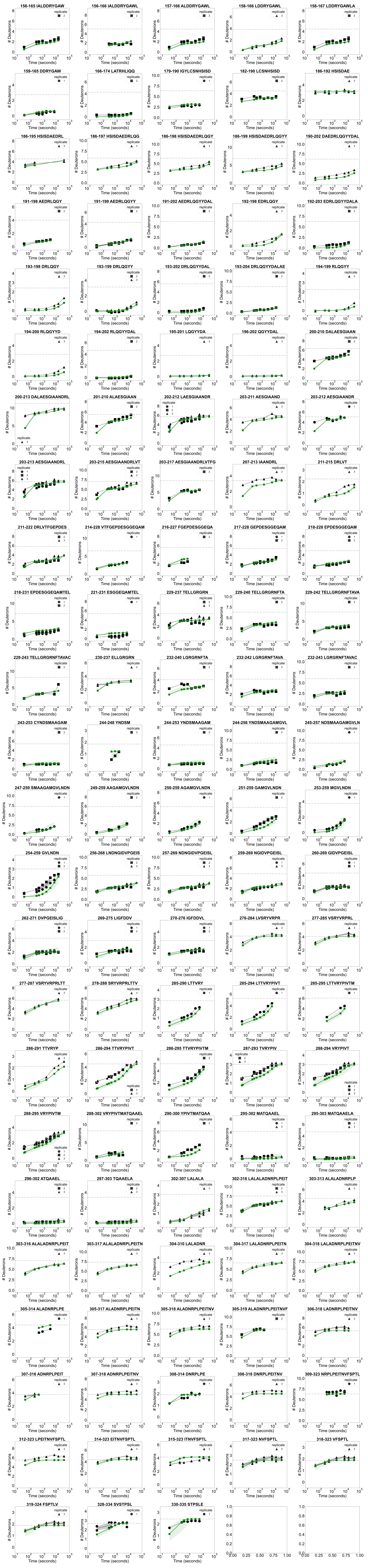

### GalR_uptake_errors_DNA_0.pdf

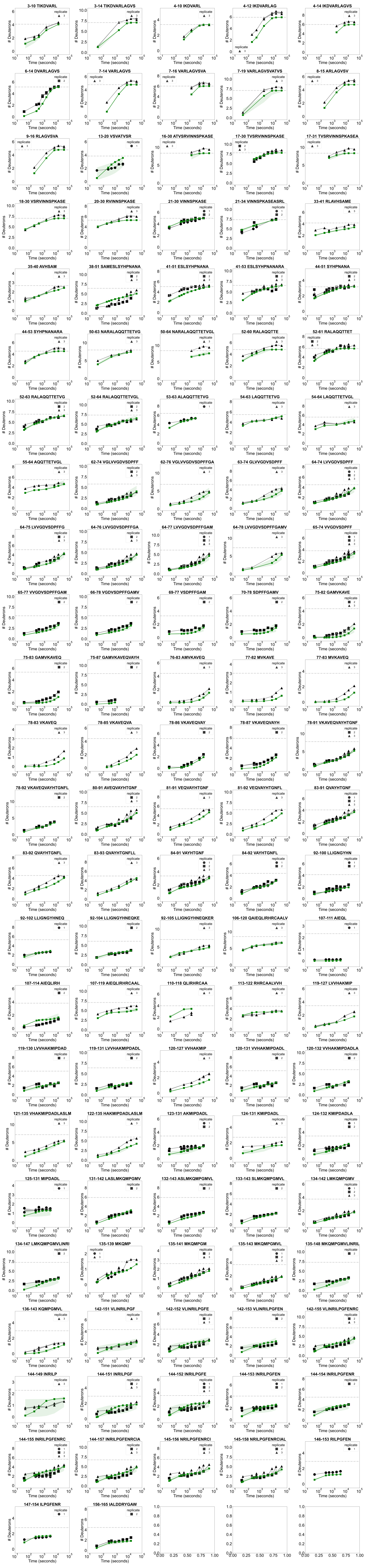

### GalR_uptake_errors_DNA_1.pdf

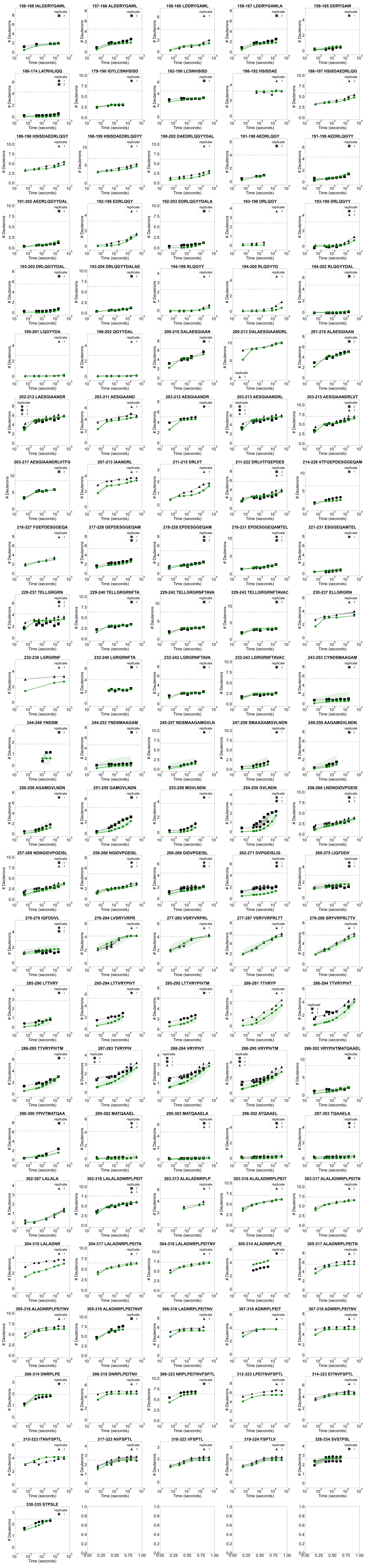

### GalR_uptake_errors_GAL_0.pdf

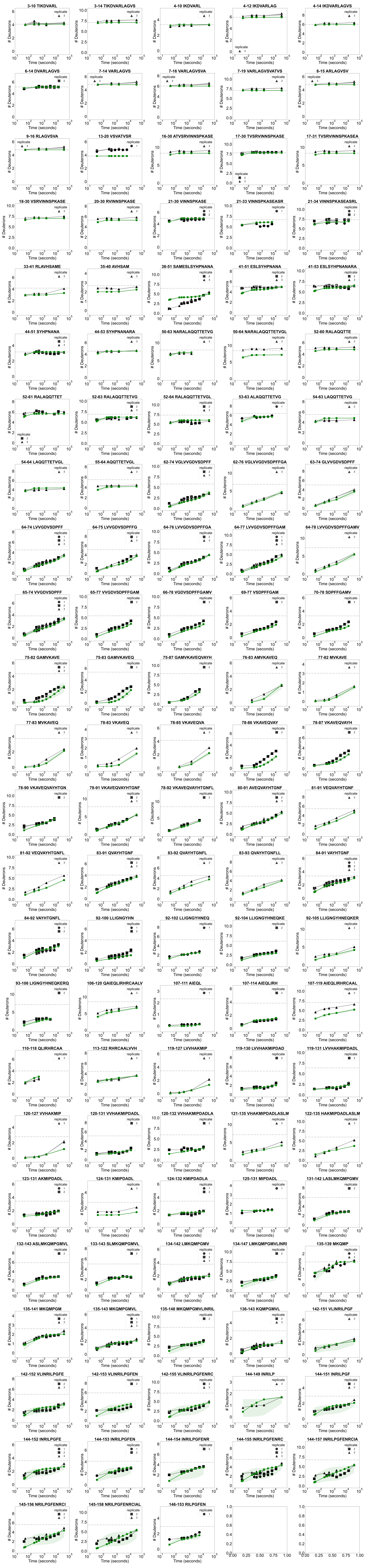

### GalR_uptake_errors_GAL_1.pdf

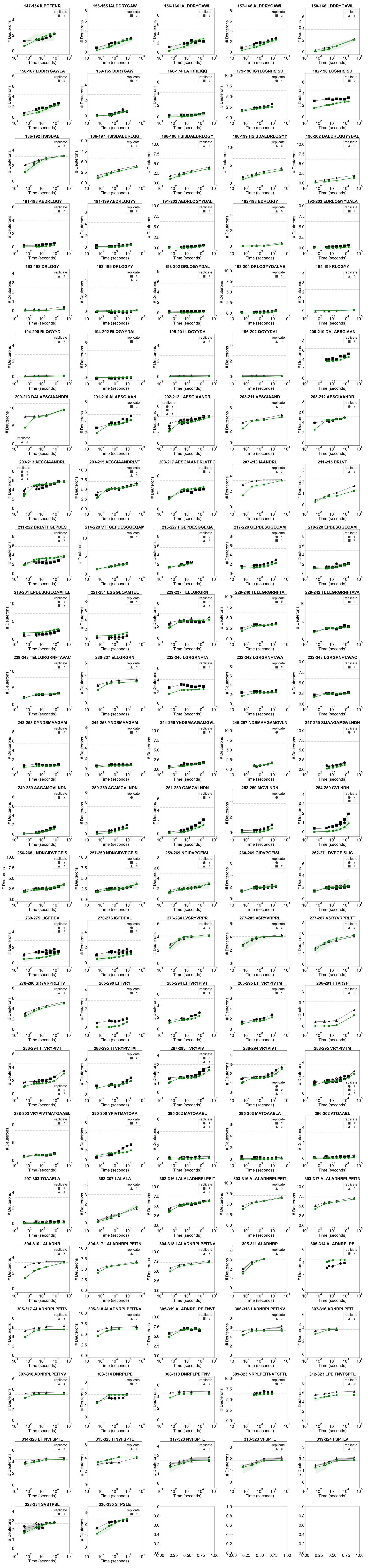

### LacI_exp_uptake_1.pdf

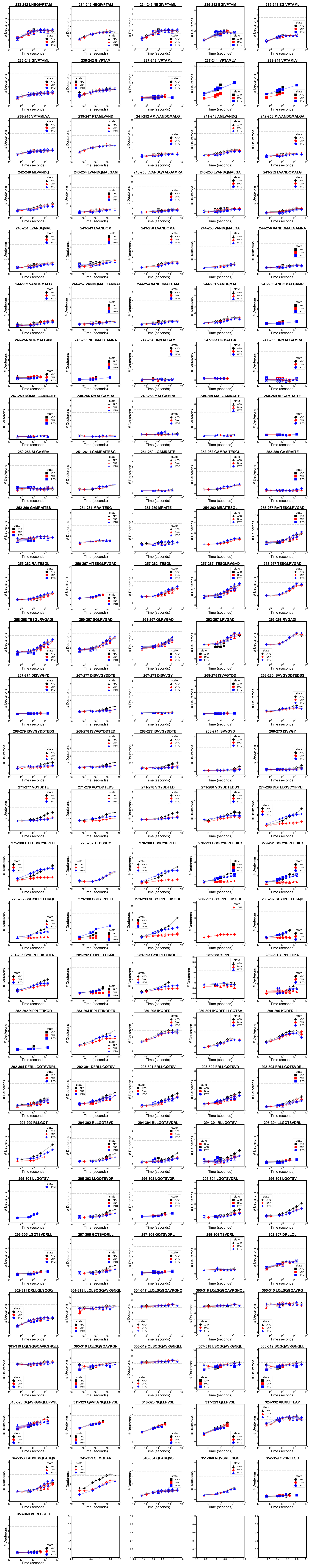

### MglB_uptake_errors_APO_0.pdf

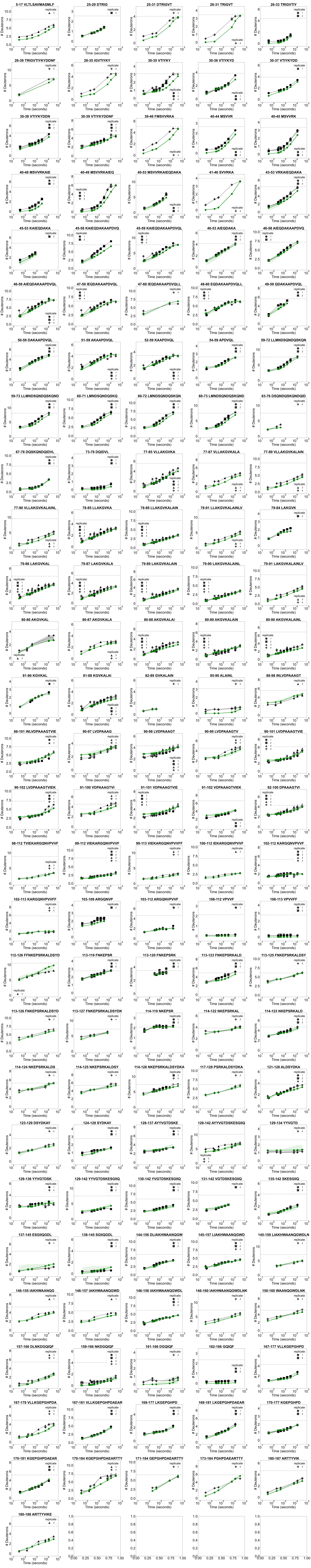

### MglB_uptake_errors_APO_1.pdf

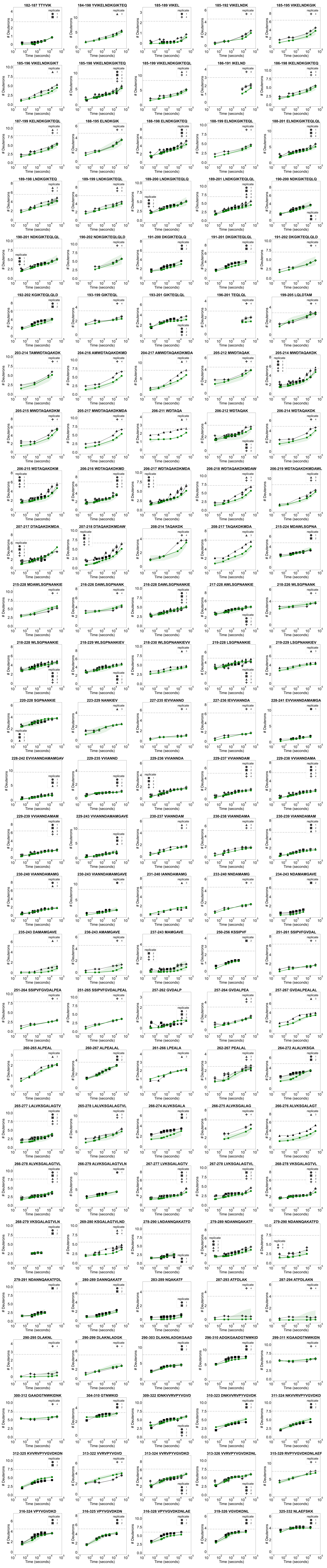

### MglB_uptake_errors_GAL_0.pdf

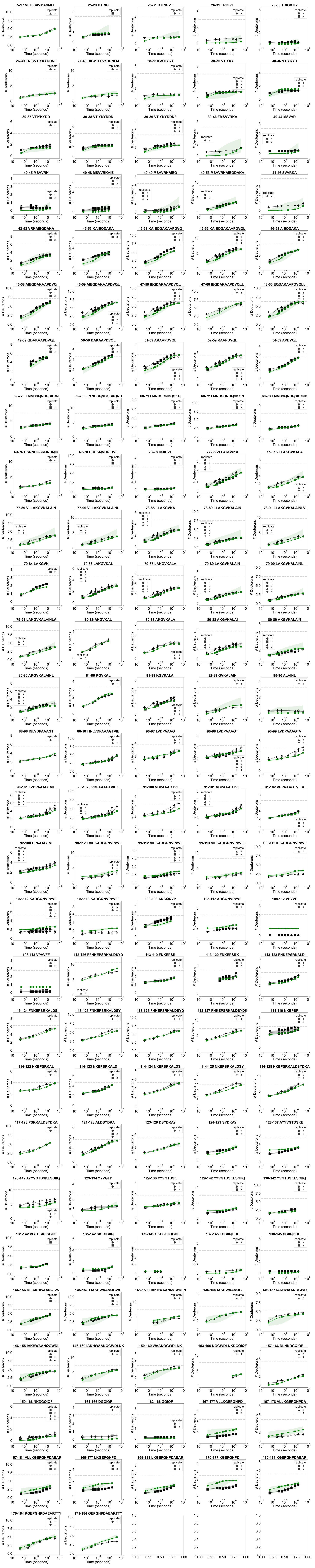

### MglB_uptake_errors_GAL_1.pdf

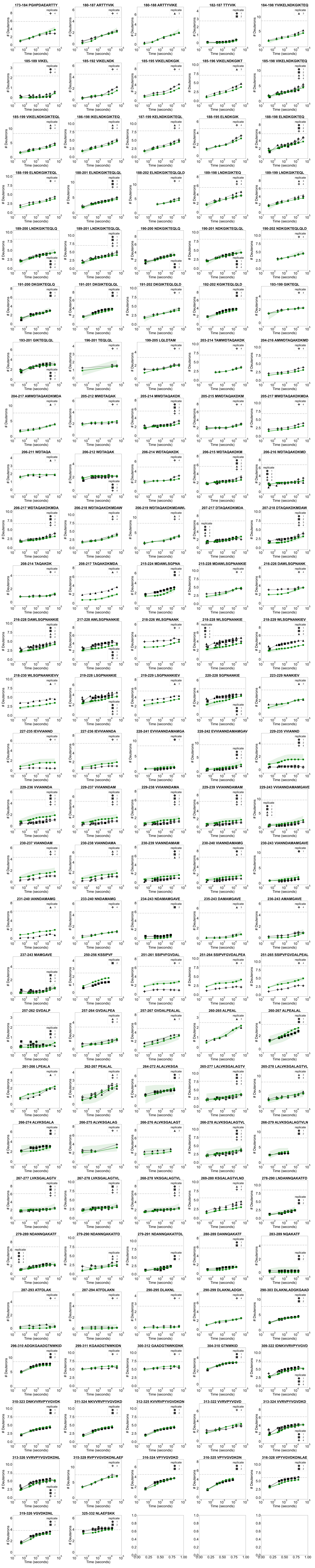

### RbsB_uptake_errors_APO_0.pdf

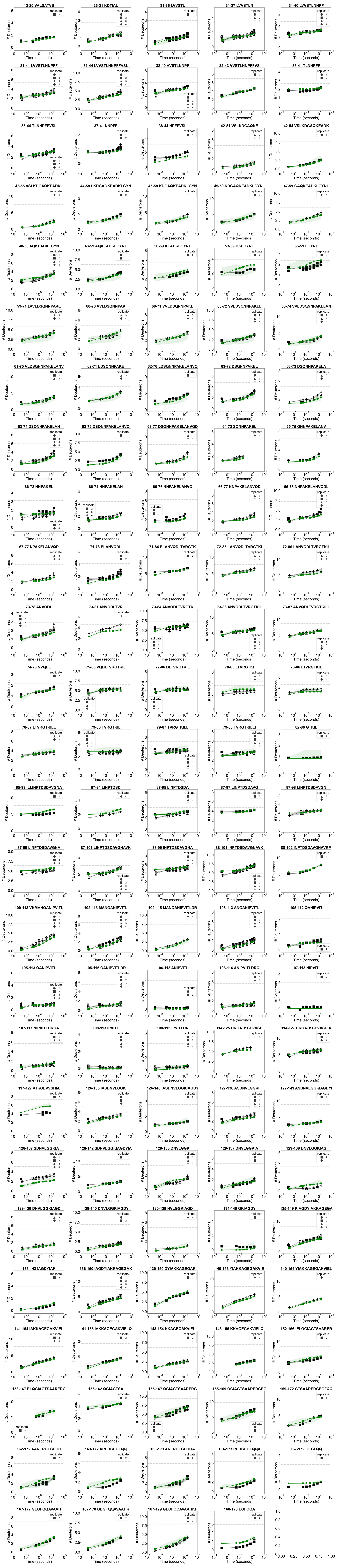

### RbsB_uptake_errors_APO_1.pdf

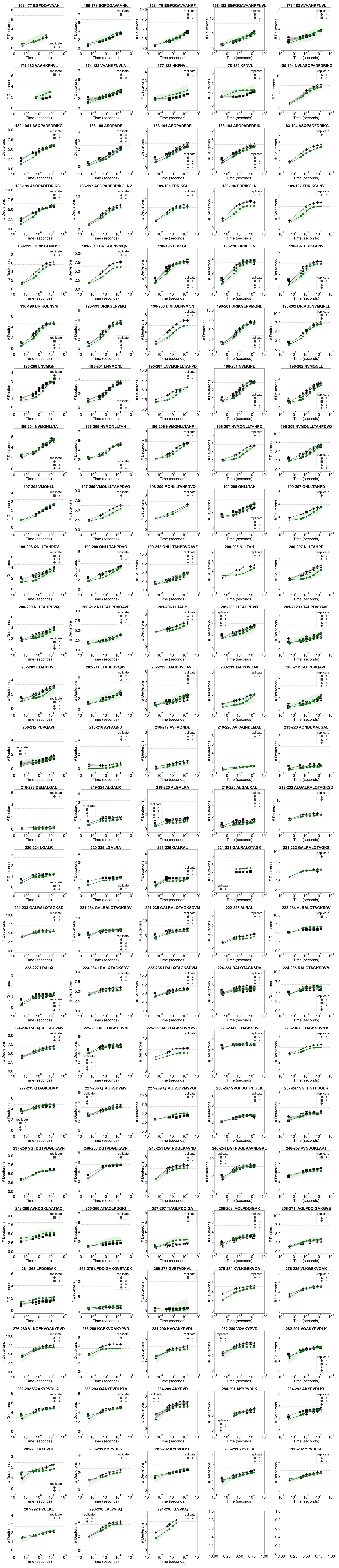

### RbsB_uptake_errors_RIB_0.pdf

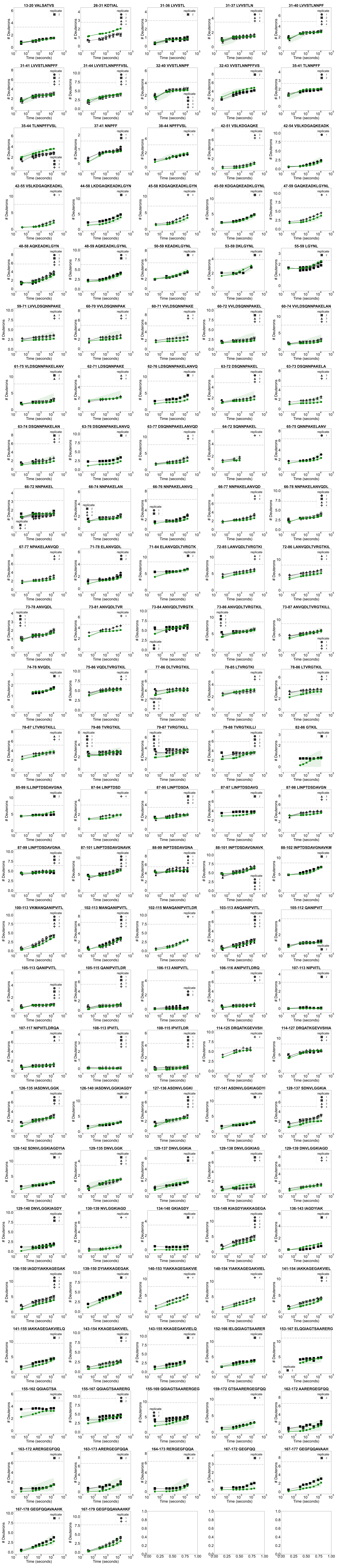

### RbsB_uptake_errors_RIB_1.pdf

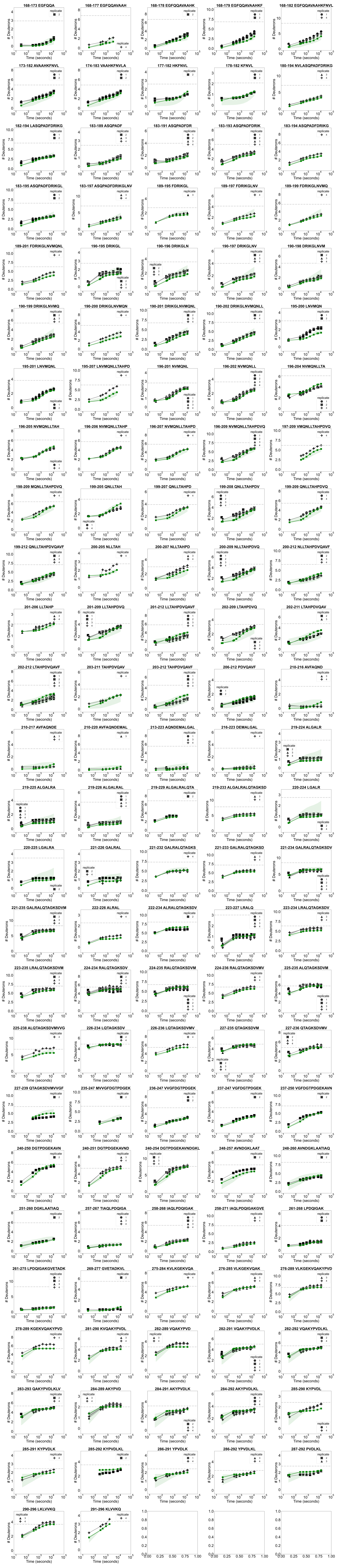

### RbsR_uptake_errors_APO_0.pdf

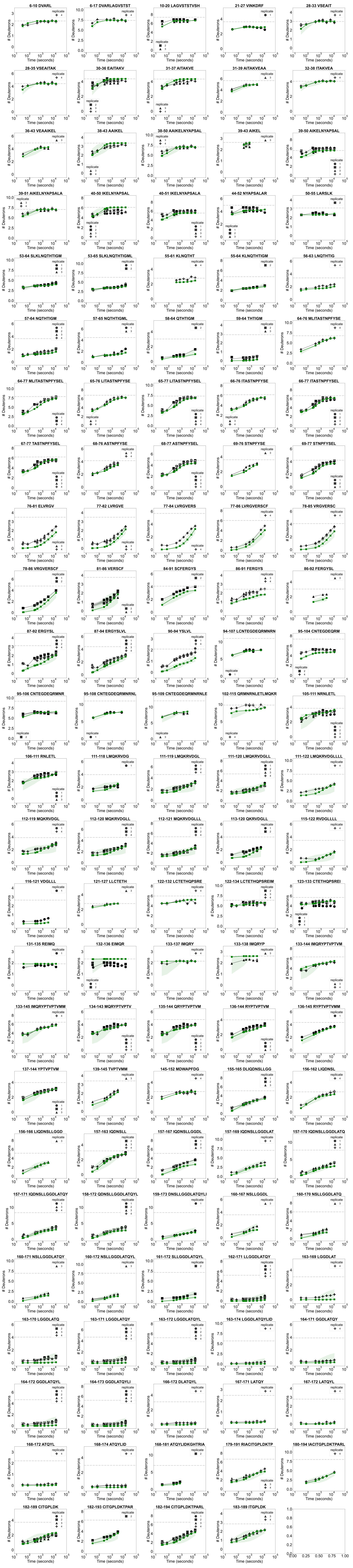

### RbsR_uptake_errors_APO_1.pdf

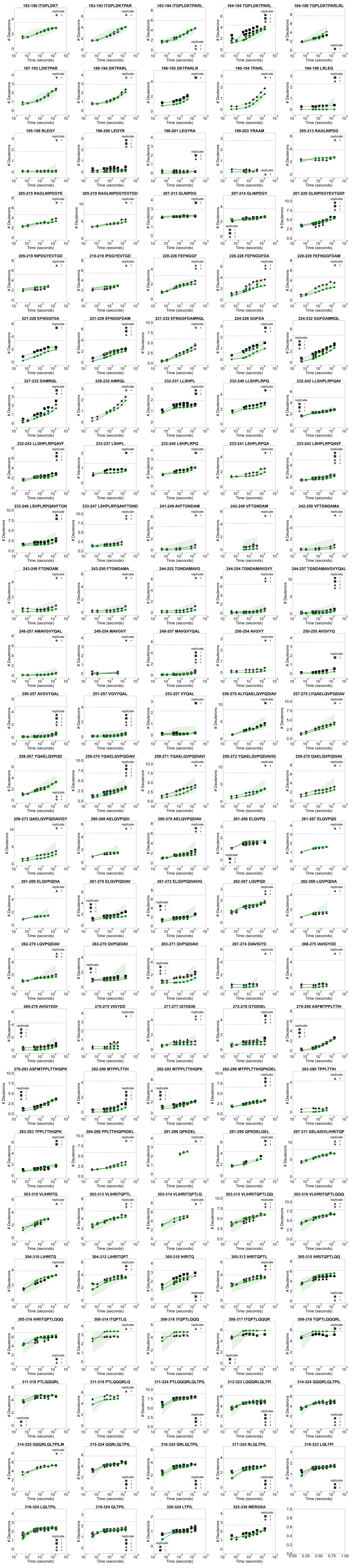

### RbsR_uptake_errors_DNA_0.pdf

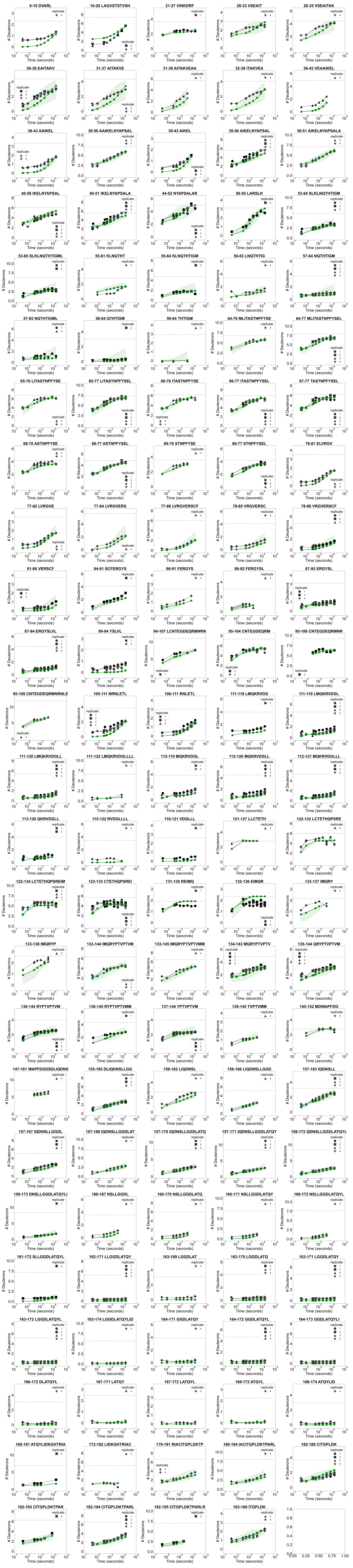

### RbsR_uptake_errors_DNA_1.pdf

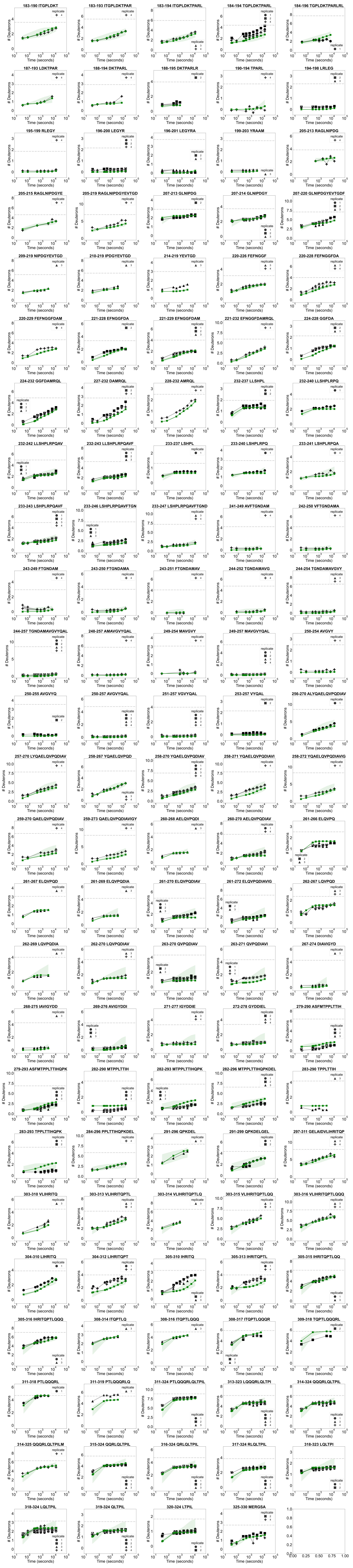

### RbsR_uptake_errors_RIB_0.pdf

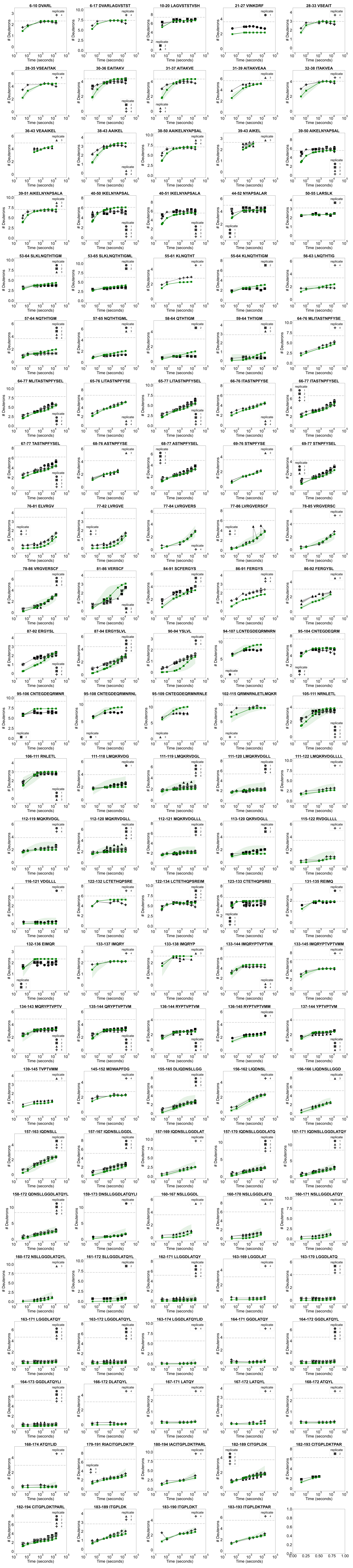

### RbsR_uptake_errors_RIB_1.pdf

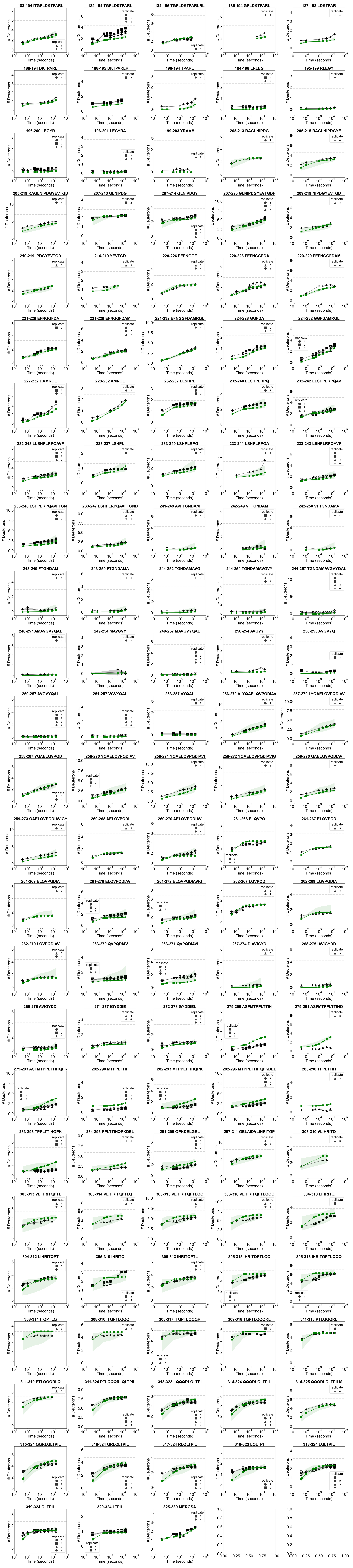
